# A supernumerary synthetic chromosome in *Komagataella phaffii* as a repository for extraneous genetic material

**DOI:** 10.1101/2023.10.05.561004

**Authors:** Dariusz Abramczyk, Maria del Carmen Sanchez Olmos, Adan Andres Ramirez Rojas, Daniel Schindler, Daniel Robertson, Stephen McColm, Adele L. Marston, Paul N. Barlow

## Abstract

**Background:** *Komagataella phaffii (Pichia pastoris*) is a methylotropic commercially important non-conventional species of yeast that grows in a fermentor to exceptionally high densities on simple media and secretes recombinant proteins efficiently. Genetic engineering strategies are being explored in this organism to facilitate cost-effective biomanufacturing. Small, stable artificial chromosomes in *K. phaffii* could offer unique advantages by accommodating multiple integrations of extraneous genes and their promoters without accumulating perturbations of native chromosomes or exhausting the availability of selection markers.

**Results:** Here, we describe a linear “nano”chromosome (of 15-25 kb) that, according to whole-genome sequencing, persists in *K. phaffii* over many generations with a copy number per cell of one, provided non-homologous end joining is compromised (by *KU70*-knockout). The nanochromosome includes a copy of the centromere from *K. phaffii* chromosome 3, a *K. phaffii*-derived autonomously replicating sequence on either side of the centromere, and a pair of *K. phaffii*-like telomeres. It contains, within its q arm, a landing zone in which genes of interest alternate with long (approx. 1-kb) non-coding DNA chosen to facilitate homologous recombination and serve as spacers. The landing zone can be extended along the nanochromosome, in an inch-worming mode of sequential gene integrations, accompanied by recycling of just two antibiotic-resistance markers. The nanochromosome was used to express *PDI*, a gene encoding protein disulfide isomerase. Co-expression with *PDI* allowed the production, from a genomically integrated gene, of secreted murine complement factor H, a plasma protein containing 40 disulfide bonds. As further proof-of-principle, we co-expressed, from a nanochromosome, both *PDI* and a gene for GFP-tagged human complement factor H under the control of *P_AOX1_* and demonstrated that the secreted protein was active as a regulator of the complement system.

**Conclusions:** We have added *K. phaffii* to the list of organisms that can produce human proteins from genes carried on a stable, linear, artificial chromosome. We envisage using nanochromosomes as repositories for numerous extraneous genes, allowing intensive engineering of *K. phaffii* without compromising its genome or weakening the resulting strain.

## BACKGROUND

Numerous platforms for producing heterologous proteins are deployed across the biotherapeutics-manufacturing sector. Often purpose-designed, proprietary and under continuous development, protocols chiefly rely on mammalian cells and can be expensive and complex. An economical, high-yielding, general-purpose microbial platform could lower barriers to entry and widen availability of vaccines and protein therapeutics globally [1]. Robust evidence of efficacy and versatility would, however, be needed for it to supersede the biomanufacturing strategies to which much of the industry is now committed [2]. Prerequisites include strong promoters of gene expression, high-density low-cost cell culture and, crucially, a toolbox for precise genetic manipulations of the protein-producing cells. We wondered if the “biotech” yeast *Komagataella phaffii (Pichia pastoris*) [3], supplemented with a synthetic chromosome dedicated to genetic engineering, could meet all of these criteria.

Engineered yeast cells possess multiple attributes considered essential for industrial protein production [4]. Model yeast species such as *Saccharomyces cerevisiae* and non-conventional yeast such as *Hansenula polymorpha*, *Yarrowia lipolytica* and *Kluyveromyces lacti* [5] have been used commercially but *K. phaffii* [6] [7] is the most prominent example. Advantages include its potential to achieve exceptionally high cell density in simple media, efficient protein-secretory machinery, ability to perform some post-translational modifications, strong natural promoters and track-record of pharmaceutical protein production [8]. Note that the *P. pastoris* genus was reclassified as *Komagataella*, which includes the subspecies *K. phaffii* and *K. pastoris*. Widely used strains are GS115 and *K. phaffii* CBS7435, as employed herein [9].

A hindrance to biomanufacture in *K. phaffii* and other yeasts is the need to integrate new genes into chromosomes. Hence sites within the native genome must serve as targets for the insertion of extraneous double-stranded DNA. This drawback is compounded in *K. phaffii* by a shortage of established tools for genetic editing compared to *S. cerevisiae* [10], and the nature of its DNA-repair machinery. In *K. phaffii,* unlike *S. cerevisiae*, the process of non-homologous end joining (NHEJ) for double-strand break (DSB) repair dominates over homologous recombination (HR) [11]. Compared to HR, NHEJ results in lower success rates for genetic deletions and insertions, and more off-target effects that can impact viability. On the positive side, lower efficiency of HR makes unintended recombination less likely [12].

One study [13] assessed the success of an HR-driven *GFP*-gene insertion designed to knock out *AOX1* (creating a Mut^S^ strain) in *K. phaffii* CBS7435. Fewer than half of 800 clones, were, in fact, Mut^S^ strains with a gene-copy number of 1. About 5% were multi-copy clones and some had a gene-copy number >10. Crucially, whole-genome sequencing (WGS) revealed non-canonical events including off-target gene disruption, co-integration of *E. coli* plasmid DNA and relocation of the *AOX1* target locus to another chromosome [14]. Such outcomes are acceptable for one-off gene insertions because screening numerous colonies is feasible while an elevated gene-copy number might boost yields. But, because clonal variability per manipulation is multiplied by the number of manipulations, attempts to insert multiple genes in *K. phaffii* are laborious and risky. Attempts have been reported to use 1000+ base-pair long HR-promoting regions (LHRs), antimicrobial-resistance (AMR) markers, and genotyping to screen multiple clones following each manipulation [11]. However, reliance on LHRs restricts genomic sites suitable for integration, the set of available selection markers for *K. phaffii* is limited, and genotyping may fail to spot accumulating off-target effects. Since damage-risk escalates with each inserted gene, resulting strains become progressively depleted of available selection markers and are less likely to grow quickly to high densities or yield as much product. These considerations hamper extensive engineering of the *K. phaffii* genome.

Numerous strategies for facilitating genetic engineering in *K. phaffii* have been reported [7]. Many involve deleting or overexpressing genes aimed at suppressing NHEJ and enhancing HR. Shifting the balance of DSB repair towards HR also improved the efficacy of CRISPR/Cas9-type protocols in *K. phaffii* [11]. Regardless of mechanisms exploited to improve the accuracy of editing and repair, genomic integration relies on identification of a suitable neutral chromosomal locus for insertion of each gene. This must be a region that is accessible to transcriptional apparatus and compatible with efficient gene expression, and which can be disrupted without adversely impacting cellular physiology or metabolism [15].

In *S. cerevisiae* [16] (bioRxiv doi: https://doi.org/10.1101/2022.10.03.510608) and *Y. lipolytica* [17] it was reported that multiple foreign genes could be incorporated into small, stable artificial chromosomes. The native 9.4-Mb *K. phaffii* genome consists of four chromosomes of 1.8, 2.3, 2.4 and 2.9 Mb with modular centromeres reminiscent of those in higher organisms [18], and telomeres consisting of 100 - 350 bp of homogenous repeated sequences [7]. We hypothesised that an additional or complementary route to facilitating genetic engineering in *K. phaffii*, while minimising damage to the native genome, would be to introduce a fifth, supernumerary chromosome, containing an appropriate origin of replication, centromere and telomeres, to serve as a dedicated receptacle for new genes.

Herein, we assembled a framework plasmid containing a centromere and an autonomously replicating sequence (ARS) (as well as a bacterial origin of replication, suitable for propagation in *Escherichia coli* cell culture). To this, we added a landing zone for new genes, and a pair of proto-telomeres (flanking an *I–Sce*I cleavage site), to create a precursor plasmid. We planned to transform *K. phaffii* cells with this precursor plasmid that could be *I-Sce*I-linearized *in vivo*, but subsequently reverted to *in vitro*, pre-transformation, *I-Sce*I linearization to create our tiny (hence “nano”) prototype chromosome. On a wild-type background strain, WGS revealed nanochromosome instability. This was largely resolved by adding a second ARS and switching to a NHEJ-compromised strain of *K. phaffii*. We eventually arrived at a stable chassis upon which a series of more elaborate nanochromosomes were constructed *in vivo* by swapping or adding genes within the integration array. Finally, we showed that genes overexpressed from nanochromosomes supported production of hard-to-manufacture human proteins.

## RESULTS

### Outline of the design and construction phase

Our goal was a stripped-down *K. phaffii* “nano”chromosome consisting only of a dedicated gene-landing zone plus those components essential for survival within the nucleus, for efficient and accurate duplication, and for reliable mitotic segregation. As reported below, we first ligated a *K. phaffii* autonomously replicating sequence (*ARS*) and a centromere part, along with an AMR gene, into pUC19 to create a framework plasmid (Fig. 1a and Additional file 1: Fig. S1). We added an initial version of the gene-landing zone and a proto-telomeres part, creating a nanochromosome-precursor plasmid. This was linearised by cleavage between the future telomeres, creating nanochromosome 1 (nChr 1) (Fig. 1b) that we used to transform *K. phaffii* cells. Subsequently we had to redesign our precursor plasmids (Fig. 1a) to improve nanochromosome stability, and switch to an NHEJ-deficient host strain of *K. phaffii*. In parallel, we established an assembly line for arrays of DNA segments (herein called integration and insertion arrays) designed to facilitate *in vivo* engineering of nanochromosomes through double-crossover HR.

**Figure 1.**
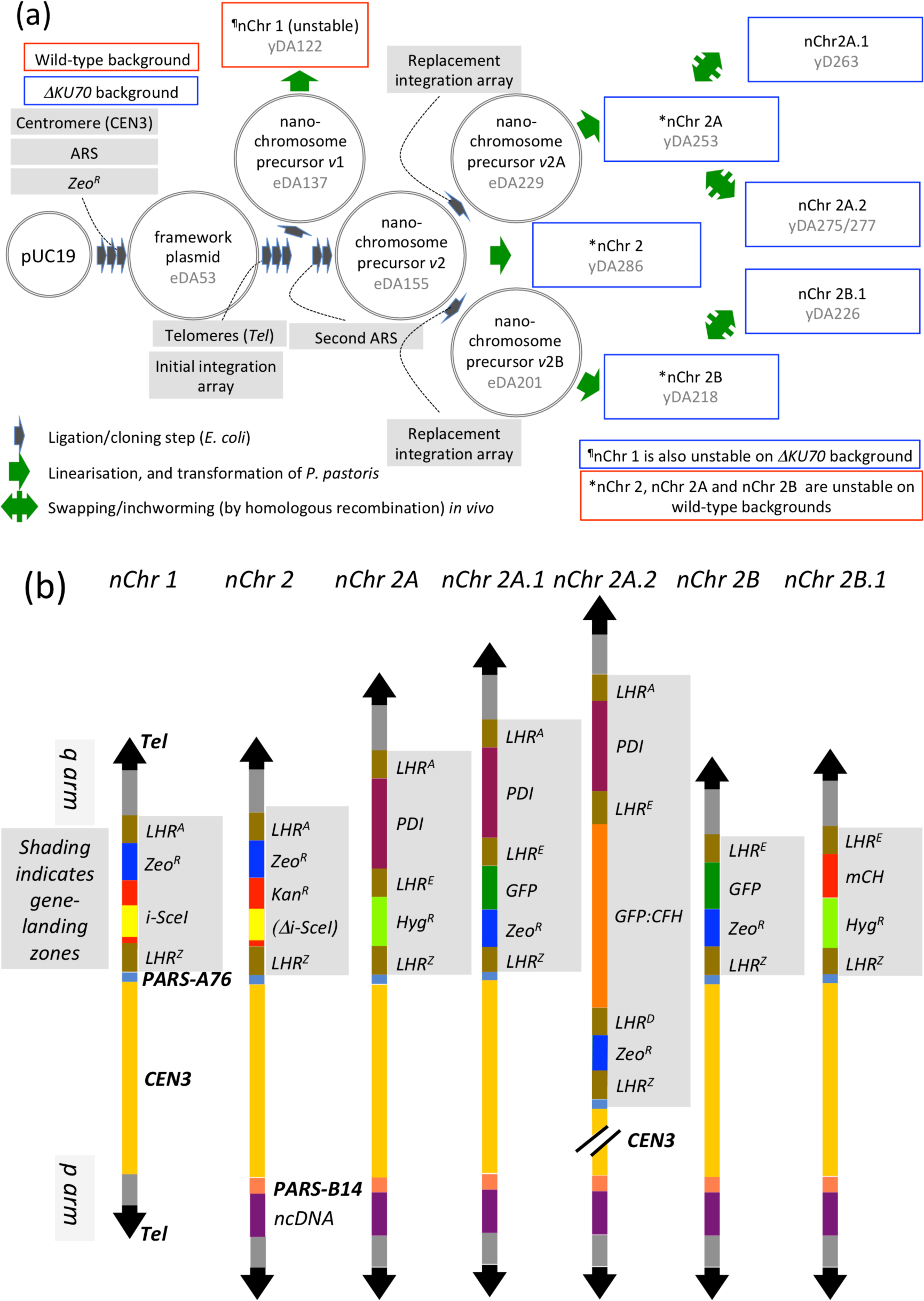
*Nanochromosome construction and engineering* (also see Additional file 1: Fig. S1). ***(a)*** *In vitro* and in *E. coli*, we assembled a centromeric, *K. phaffii* ARS-containing, framework. We added an initial gene-integration array (landing zone) and proto-telomeres, creating nanochromosome (nChr)-precursor version (*v*)1. We cleaved this between its proto-telomeres, and used the linear product - nChr 1 – to transform *K. phaffii* cells. To improve nChr stability, we extended nChr-precursor *v*1 and added a second ARS (for precursor *v*2), then replaced the initial integration array with new ones, creating precursors *v*2A and *v*2B. These yielded nChr 2A and nChr 2B, which were stable only in a *ΔKU70* strain. Finally we explored the feasibility of engineering nChr 2A and nChr 2B (to Chr 2A.1 *etc*.) *in vivo*. Plasmids are not drawn to scale. ***(b)*** Schematic representations (drawn approximately to scale) of the synthetic, linear nanochromosomes as constructed in the current study. Lists of oligos, plasmids and strains may be found in Additional file 2: Tables 1, 2 and 3 respectively. Promoters and terminators are not shown. *CEN* = centromere, *ncDNA* = non-coding DNA, *LHR* = long HR-compatible region, *mCH* = mCherry gene, *GFP:FH* = gene coding for a GFP-FH fusion.

### A centromeric, ARS-containing framework plasmid

We selected DNA corresponding to the centromere of *K. phaffii* Chr 3 [19] as the nanochromosome centromere (henceforth, *CEN3*). When creating the *EcoR*I-cleavage site, an additional nucleotide, G, was introduced in the *CEN3* core region (Fig. 2a and Additional file: Fig. S2). This subsequently aided distinction between the native centromere and (nanochromosomal) *CEN3* when analysing WGS results. Two segments were generated by PCR, from *K. phaffii* genomic (g)DNA, prior to combination by sequential cloning to create *CEN3* in plasmid pUC19, yielding plasmid eDA24 (Fig. 2a). We selected DNA corresponding to *PARS*-A76 on *K. phaffii* Chr 1 [20] to supply a yeast origin of replication, and inserted this into pUC19 creating eDA26. We amplified zeocin- (*Zeo^R^*, *ble*) and hygromycin-resistance (*Hyg^R^*, *hph*) genes from commercial plasmids and inserted them into eDA26, forming eDA40 (*Zeo^R^*) and eDA37 (*Hyg^R^*) (Additional file 1: Fig. S1). Gibson assembly of the following yielded the centromeric ARS-containing 10.5-kb framework plasmid, eDA53 (Fig. 2b): *(i)* a 1.4-kb *Zeo^R^* part; *(ii)* 0.51-kb *ARS-76*; and *(iii)* the linearized, 8.85-kb, centromeric, plasmid eDA24 that thereby contributes *CEN3*, some pUC19-derived scaffolding DNA, a bacterial origin of replication and an ampicillin resistance (*Amp^R^*, *bla*) gene.

**Figure 2.**
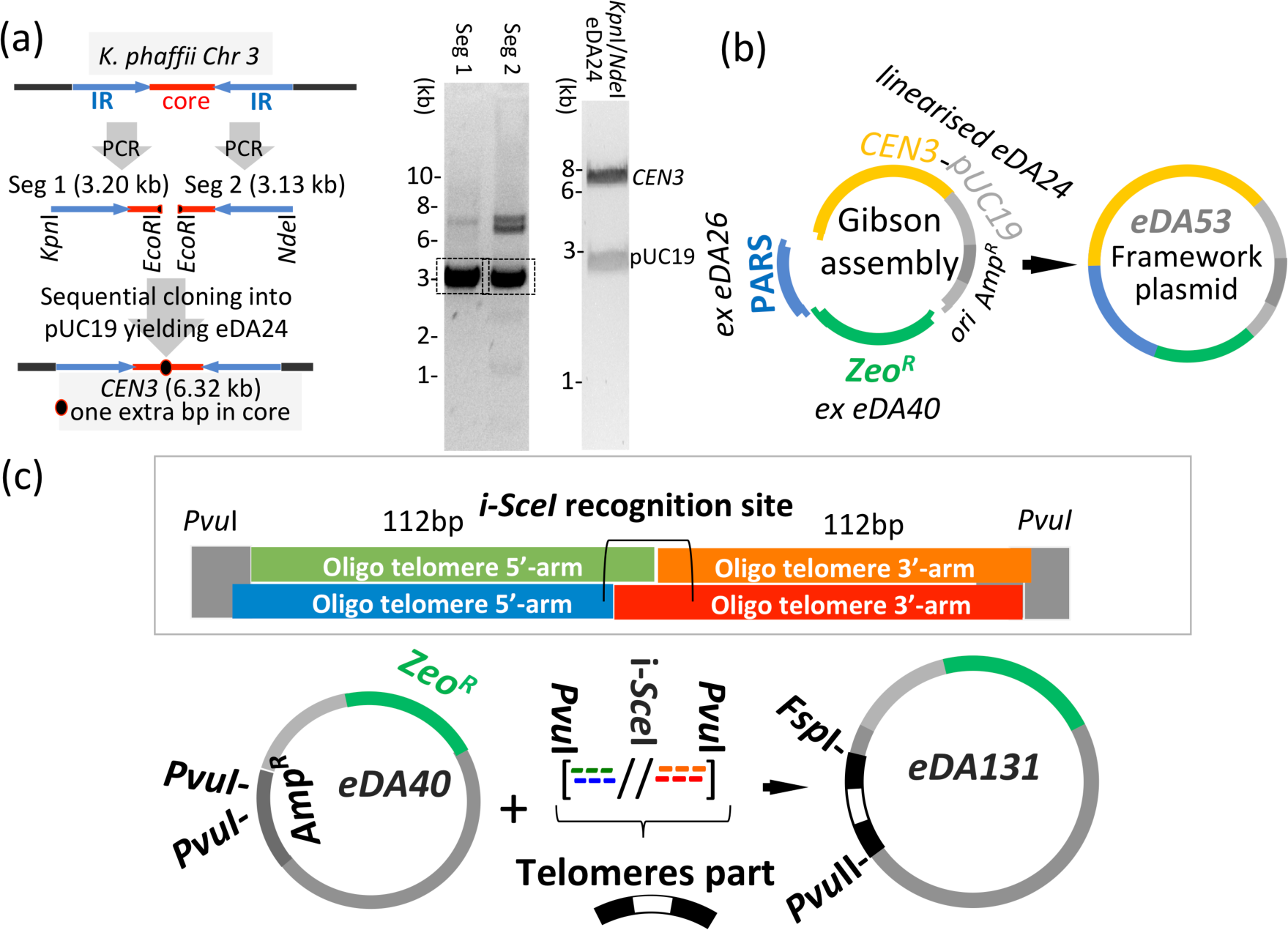
*Preparation of key parts. **(a)*** Preparation of *CEN3*, using *K. phaffii from* gDNA. Left-hand gel: PCR amplicons of segment (Seg) 1 (oligos 197/198), and Seg 2 (oligos 199/200). This approach introduces one extra, functionally silent, base pair (G:C) in the core region (see Additional file 1: Fig. S2). DNA bands of expected sizes (boxed) were extracted and digested with appropriate restriction enzymes before sequential cloning into pUC19 to yield eDA24. Right-hand gel: restriction digestion of eDA24 confirming presence of 6.3-kb *CEN3* (IR = inverted repeats). ***(b)*** Assembly of framework plasmid in a pUC19-derived scaffold (*ori* = bacterial origin of replication) (eDA53 validation shown in Additional file 1: Fig. S3). ***(c)*** The oligos used to construct the telomeres part (*Tel*, Additional file 1: Fig. S4). Each proto-telomere contains 16 copies of TGGATGC and the two are linked by the 18-bp *I-Sce*I-recognition site. Lower schematic: cloning of *Tel* into eDA40 (eDA131 validation shown in Additional file 1: Fig. S5).

### Construction of the telomeres part (Tel)

We used a “telomerator”-inspired approach [21] entailing insertion into the framework plasmid of a DNA sequence, which we called the telomeres part (*Tel*), comprised of proto-telomeres flanking a restriction site. In our case, two sets of 16 inverted telomere-repeats bracketed an *I-Sce*I-cleavage site (Fig. 2c and Additional file 1: Fig. S4). Thus *I-Sce*I can linearise the DNA molecule, yielding telomeres at either end. We constructed *Tel* by annealing each of two pairs of similar-length synthetic oligos, to form its 5’- and 3’-halves with complementary overhangs. The two halves were tandemly ligated into eDA40, forming eDA131 (Fig. 2c).

### An assembly line for integration and insertion arrays

We built insertion arrays (donors) and integration arrays (acceptors) to allow precise nanochromosome engineering in *K. phaffii* cells. Each array includes *Hyg^R^* or *Zeo^R^* and features genes-of-interest (*GoIs*) flanked by *LHR*s. *GoIs* are expression cassettes containing a gene plus its promoter and terminator regions. *LHR*^A^, *LHR*^B^ *etc.* are unique sequences of *∼*1000 bp from a library (*LHRs^A-Z^*), with *LHR^Z^* reserved for the terminus of each array. Thus, it should be possible to (for example) swap *Hyg^R^* within the nanochromosome-resident integration array *LHR^A^-GoI^1^-<u>LHR^E^</u>-Hyg^R^*-*LHR^Z^* for *GoI^2^-LHR^D^-Zeo^R^* by transforming cells with insertion array *<u>LHR^E^</u>-GoI^2^-LHR^D^-Zeo^R^-<u>LHR^Z^</u>* (Additional file: Fig. S6a). Subsequently, *Zeo^R^* would be available for exchange with *GoI^3^-LHR^E^-Hyg^R^*, and so forth. Theoretically, repeated cycles of this “inch-worming” process would generate a string of evenly spaced genes (and their regulatory sequences). Recycling of a single pair of selection markers (*Hyg^R^* and *Zeo^R^*) is inherent to this strategy.

The following parts were deployed in ligase-assisted directional assembly of arrays within digested pUC19 (Additional file: Fig. S6b): *LHRs^A-Z^*; synthetic codon-optimised *GoI*s; and *Hyg^R^* or *Zeo^R^* fused to *LHR^Z^*. The assembled arrays were: *(i)* PCR-amplified using primers carrying 5’-blunt-end and 3’-*Sal*I/*Xho*I restriction-enzyme sites to create integration arrays; or *(ii)* digested with *AsiS*I or *Fsp*I for insertion arrays.

### Introduction of the initial gene-landing zone into the precursor plasmid

We introduced into our *CEN3*- and *ARS*-containing framework plasmid (eDA53) an initial integration array, *LHR^A^*-*Zeo^R^*-(*P_AOX1_)I-SceI(T_AOX1_*)-*LHR^Z^*. The gene *I-SceI* encodes a restriction enzyme that recognises the cleavage site [22] located between proto-telomeres within *Tel*. Thus we anticipated that a future plasmid containing both *I-SceI* and *Tel* would self-linearise *in vivo* after *I-SceI* induction. This one-off integration array (unlike those used subsequently) was not preassembled, but was introduced stepwise by: *(i)* inserting *LHR^A^* into eDA53 (yielding eDA71); *(ii)* constructing *(P_AOX1_)I-SceI(T_AOX1_)* in pPICZα B and amplifying by PCR; *(iii)* amplifying *LHR^Z^* by PCR; and *(iv)* ligating the purified, *Bsm*BI-digested products from step *(ii)* and *(iii)* with *Bsm*BI-linearized, gel-purified, eDA71, yielding eDA83 (Fig. 3a).

**Figure 3.**
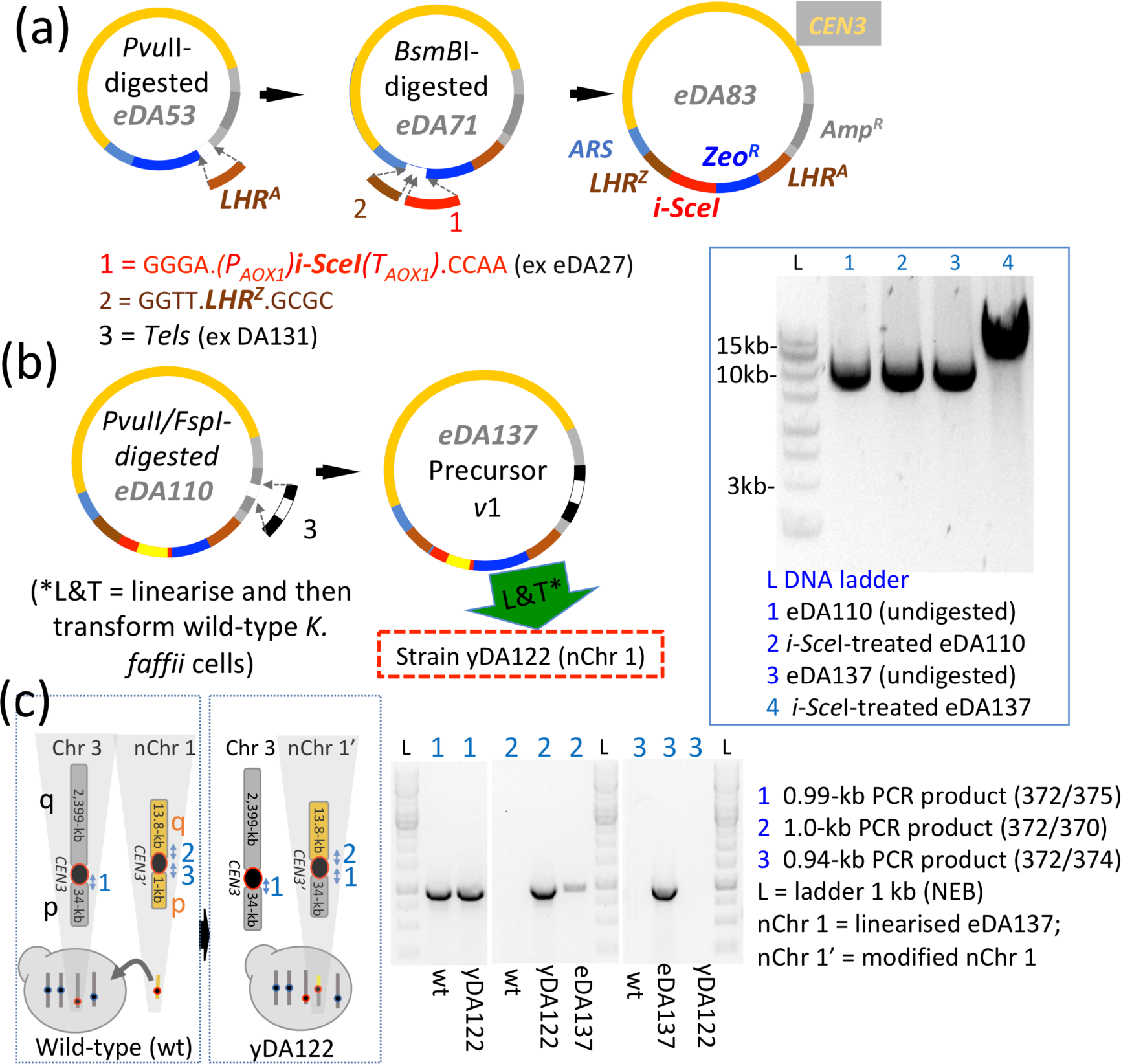
*Production and testing of precursor plasmid v1 and nChr 1*. ***(a)*** Incorporation of an initial, one-off, gene-landing pad into the precursor plasmid. Work on eDA83 was halted due to leaky *I-SceI* expression in *E. coli*. ***(b)*** Following deletion of *I-SceI* in eDA83 by insertion of *Kan^R^* (creating eDA110), *Tel* was inserted to yield eDA137, the precursor plasmid (*v*1) of nChr 1. The plasmid was linearized *in vitro* and used to transform (“L&T” in figure) CBS7435 *K. phaffii* cells creating strain yDA122. The agarose gel shows *I-Sce*I digestion of *Tel*-containing eDA137 (*vs* control, *i.e.* no-*Tel*, eDA110). ***C*** Despite nChr 1 appearing stable in chromosome-loss assays performed on yDA122, WGS (and subsequently, colony PCR, see gel) revealed major chromosomal rearrangements as indicated in the schematic. Note, in yDA122, the loss of PCR-product 3 (of nChr 1), but retention of PCR-product 2 of the suspected translocation product nChr 1* (*i.e.* nChr 1(p):Chr 3(q)).

### A prototype nanochromosome

We aimed to make a nanochromosome-precursor plasmid by excising *Tel* from eDA131 and inserting it into eDA83’s *Amp^R^* site. Disappointingly, the ligation product was unstable in *E. coli* due to leaky (*P_AOX1_*)*I-SceI* expression. We abandoned *in vivo* plasmid linearization in favour of *in vitro* linearization by *I-Sce*I, while opting to salvage our initial integration array. To this end, we replaced, in eDA83, *P_AOX1_* and part of the *I-SceI* ORF, with a bacterial *Kan^R^* expression cassette, yielding eDA110 containing the array: *LHR^A^*-*Zeo^R^*-*Kan^R^*(Δ*I-SceI*)-*LHR^Z^*. We inserted *Tel* into eDA110, deleting *Amp^R^* and yielding *I-Sce*I-cleavable eDA137 (Fig. 3b). Unlike eDA83, eDA137 (with *I-SceI* effectively deleted) could be propagated in *E. coli*. We called this nanochromosome-precursor plasmid version 1 (precursor *v*1).

We transformed wild-type *K. phaffii* cells with *I-Sce*I-linearized precursor *v*1 creating our prototype nanochromosome (nChr 1). We re-plated colonies that grew on zeocin, and based on colony-PCR, selected yDA122 for liquid culture in the presence of zeocin and used it to make a glycerol stock. We took cells from the stock and cultured them, to assay zeocin resistance after about ten generations on zeocin-free media, providing an indication of nChr 1 mitotic stability. Nine out of ten cells remained zeocin-resistant (Additional file 2: Table 5). For a more direct test of nChr 1 integrity, we submitted yDA122 cells, recovered from glycerol stock and cultured overnight, for WGS. Analysis by *de novo* assembly and visualization [23] revealed that nChr 1 had undergone a substantial, rearrangement (Fig. 3c and Additional file 1: Fig. S7a) entailing fusion of the p-arm of nChr 1 with a copy of the Chr 3 ∼34-kb p-arm. While this conflicted with the PCR-based genotyping performed shortly after transformation, it agreed with PCR-based genotyping of yDA122 after multiple generations with no antimicrobial selection (Fig. 3c). Instability of nChr 1 was likewise evident from WGS of two other colonies, similarly prepared.

### Extension of backbone failed to improve integrity

We wondered whether the telomere-to-centromere proximity within the p-arm of nChr 1 might impair maintenance, or promote chromosomal aberrations [24]. We also conjectured that a second *ARS* might expedite duplication and/or segregation [25]. To address these possibilities, we reverted to eDA110 and inserted *K. phaffii PARS-B413* [20] in tandem with some (1.3-kb) ncDNA, into the future p-arm of the precursor plasmid to yield eDA146 (Fig. 4a). The AT-rich *PARS-B413* is less efficient than the GC-rich *PARS-A76* already present in the q-arm. The resultant arrangement was designed to resemble (native) Chr 3 in which the centromere is sandwiched between (stronger) GC-rich *PARS*-*C2204* and (weaker) AT-rich *PARS*-*C2216* [26]. Into eDA146 we inserted *Tel* to create *I-Sce*I-cleavable eDA155 (precursor *v*2) (Fig. 4a,b). PCR mapping of linearized eDA155 (*i.e.* nChr 2, Fig. 4c) validated its integrity.

**Figure 4.**
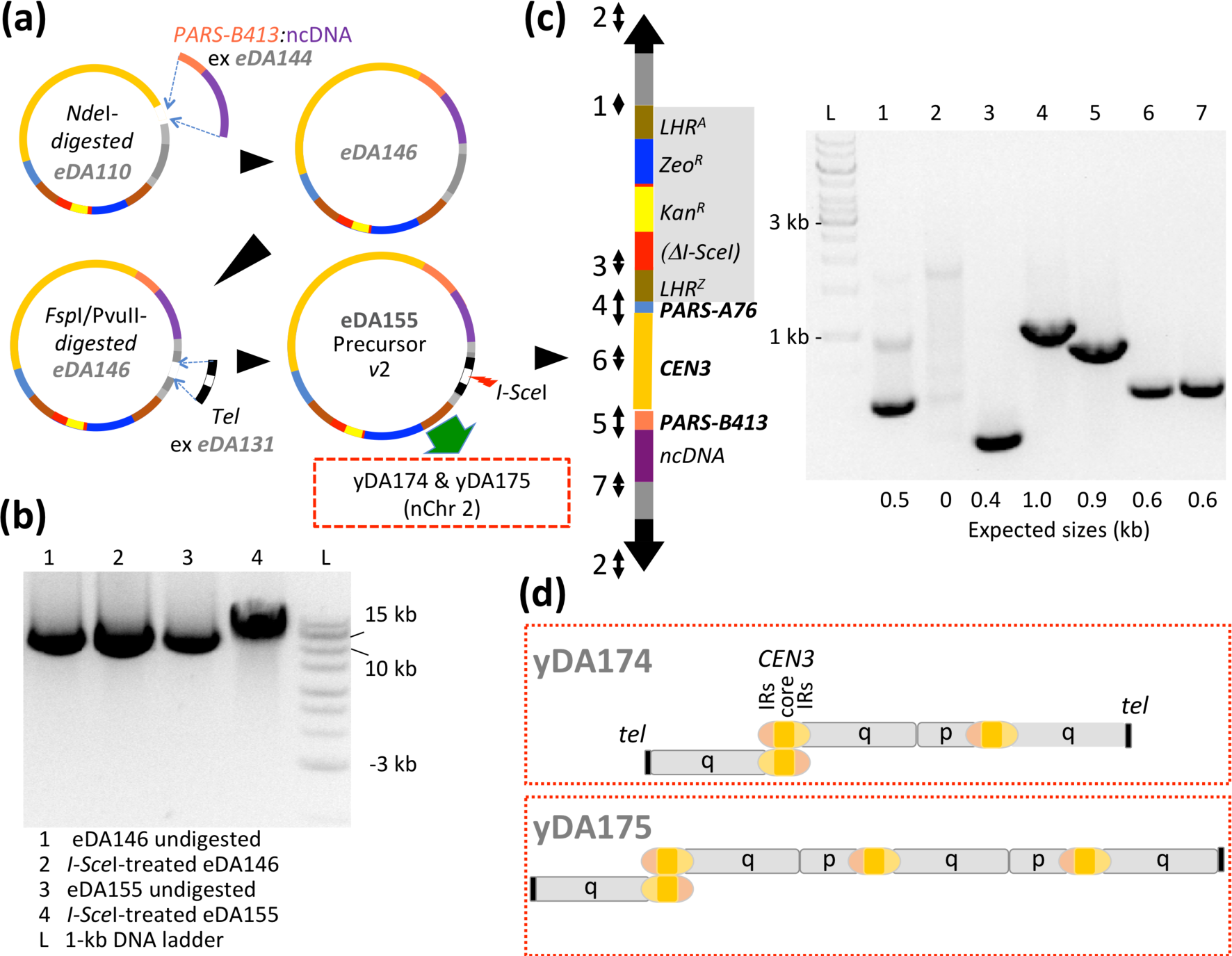
*Observed aberrations of an extended neochromosome* (nChr 2) *in wild-type cells* **(a)** The shorter (p) arm of nChr 1 was extended with non-coding (nc)DNA, and a second ARS added, by cloning components into precursor *v*1 to create *v*2. **(b)** Agarose gel showing *I-Sce*I digestion products for eDA146 (control) and eDA155**. (c)** Validation of nChr 2 by PCR before *K. phaffii* transformation (for oligonucleotides sets see Additional file 2: Table 1). **(d)** *De novo* assembly analysis of the WGS for the resultant strains (yDA174 and yDA175) revealed a preponderance of fused chromosomes indicating instability (see Fig. 5 and Additional file 1: Figs. S7 and S8). Strain yDA174 carries a triple-fusion between a nChr 2 q-arm (*i.e.,* lacking its p-arm), another nChr 2 q-arm; and a copy of nChr 2 lacking the p-arm telomere; the junction formed between q-arms involves the inverted-repeat regions (IRs, 99% sequence identity) of the centromeres but it was not possible to ascertain the nature of this fusion event. In strain yDA175 an additional copy of nChr 2, lacking its p-arm telomere, appears fused to the triple-fusion structure seen in yDA174.

Thus, we linearized precursor *v*2/eDA155 *in vitro* and used the product, nChr 2, to transform *K. phaffii* cells. From colonies growing on zeocin, we picked yDA174 and yDA175. Chromosome-loss assays (Additional file 2: Table 5) were promising and the integrity of nChr 2 in these strains was supported initially by superficial analysis of PCR-based genotyping. However, an unanticipated PCR amplicon was noted (Additional file 1: Fig. S8) while subsequent WGS (Fig. 5), *de novo* assembly (Additional file 3) and coverage analysis revealed fused end-to-end, di- or tricentric DNA molecules, consistent with this anomaly, in which telomere repeats occur exclusively at termini (Fig. 4d).

**Figure 5.**
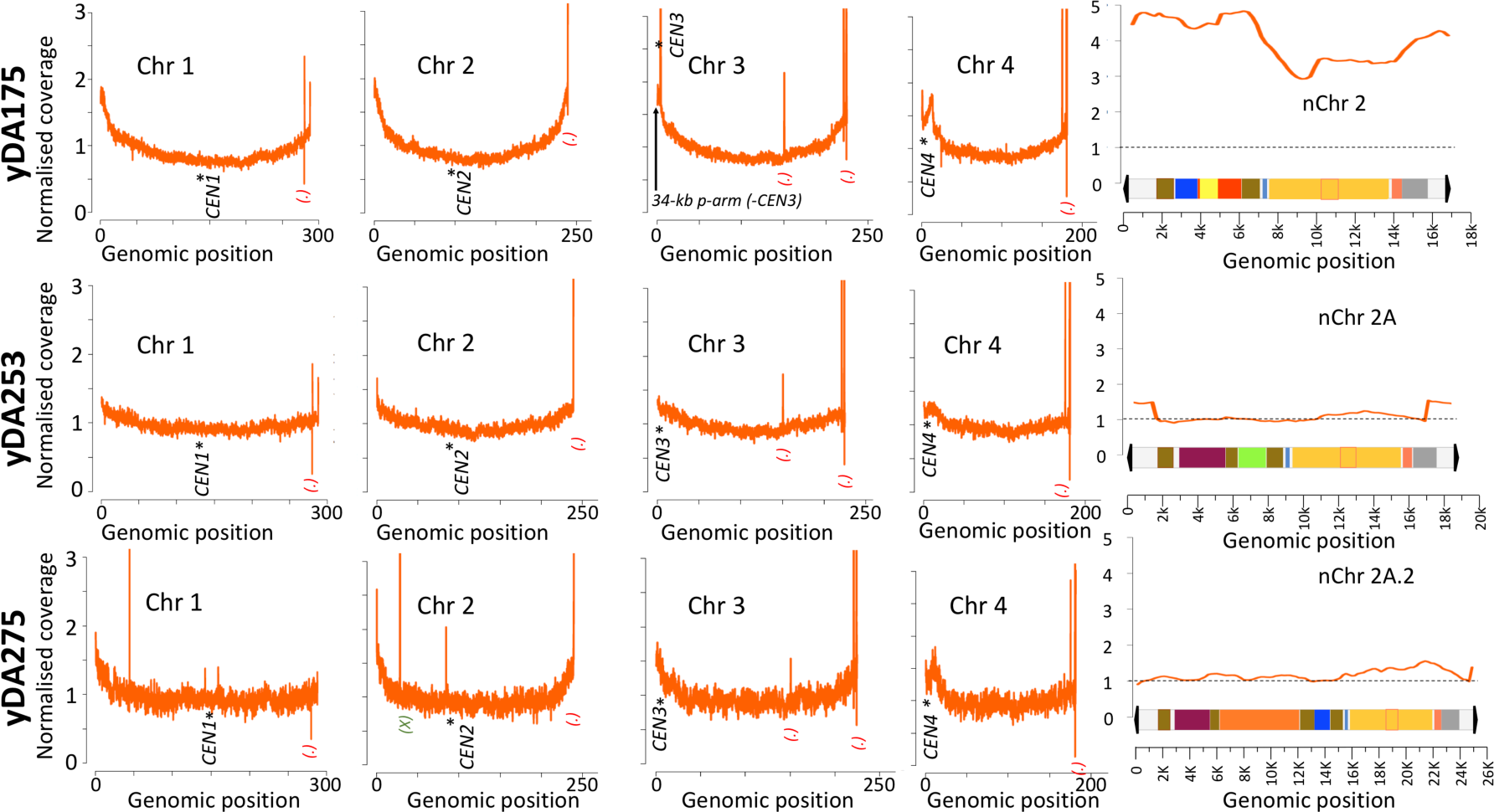
*Whole-genome sequencing coverage data for examples of wild-type and ΔKU70 K. phaffii cells containing nanochromosomes* (Also see Additional file 1: Fig. S7) The *y*-axes show “normalised coverage” values obtained by dividing the coverage of each base pair by the mean coverage of all base pairs in the respective sample. This allows comparisons, and identification of duplicated segments or deletions. The *x*-axes show the position (genomic coordinate) of each base pair in its respective chromosome (values in kilobases for native chromosomes). Each nanochromosome is shown as a schematic (colour-coding as in Fig. 1b), drawn to the same scale as the *x*-axis in each case. * Indicates a centromere. (.) Indicates anomalies attributable to native genome rearrangements (*e.g.,* duplicated loci) or artifacts arising from highly repetitive regions such as found in telomeres. (X) Indicates a ∼1-kb duplicated region of Chr 2 corresponding to an ORF encoding an unknown protein (inexplicably observed exclusively in *ΔKU70* strains transformed with insertion arrays). Red boxes, drawn on the nanochromosome schematics, indicate *CEN3* core regions characterised by the signature presence of an extra GC base pair. ***Upper panel:*** For yDA175, on wild-type background, a plot of normalized coverage indicates two-four copies of nanochromosomal sequence per cell. This elevated normalised coverage values for nChr 2 suggest multiple copies per cell of its DNA content. The high copy-number for the Chr 3 centromere correlates with a high copy-number of nChr 2 and supports the existence of chimeric multi-centric nanochromosomes (a tri-centric model is suggested in Additional file 1: Fig. S8). This observation is compatible with yDA175 and yDA177 *de novo* assembly results obtained from long-read WGS (Additional file 3). ***Middle and lower panels:*** Both these trains on a Δ*KU70* background are each consistent with a single copy per cell of its nanochromosome, specifically: yDA253 with a single copy of nChr 2A; and yDA275 with a single copy of nChr 2A.2 (*i.e.* after the inch-worming proof-of-principle experiment). [*(BioProject PRJNA971544)*]

### Switch to an NHEJ-deficient host strain

While HR events could occur between telomeres, these should lead to elongation or shortening rather than fusion, provided that telomere length exceeds a threshold value [27]. Given that HR repair is, in any case, inefficient in *K. phaffii*, we suspected that the observed end-to-end fusion of nanochromosomes in wild-type cells was a result of NHEJ. Multicentric nanochromosomes are regarded as unstable and, if not resolved, a source of further genomic abnormalities. Although, our multicentric entities appeared to persist over generations, they were deemed unsuitable as expression platforms. We therefore explored the use of a host strain in which NHEJ is impaired by *KU70*-knockout [11, 28] (Additional file 1: Fig. S9a) but which remained viable in cultures over multiple days at 30 °C (Additional file 1: Fig. S9b).

Attempts to maintain nChr 1 in *ΔKU70* cells proved no more successful than they had been in wild-type, whereas nChr 2 appeared to be stable the NHEJ-compromised cells (Additional file 1: Fig. S10). We therefore focused on nChr2. To better compare nanochromosome engineering in wild-type *vs ΔKU70* strains, we chose to create more useful, versatile versions. We therefore excised the integration array (*LHR^A^*-*Zeo^R^*-*Kan^R^*(Δ*I-SceI*)-*LHR^Z^*) of precursor *v*2 and replaced it with either: *(i)* a triple-LHR integration array expressing a methanol-inducible *PDI* gene, and *Hyg^R^*, *i.e. LHR^A^-PDI^H^-LHR^E^-Hyg^R^-LHR^Z^* (for precursor *v*2A); or *(ii)* a double-*LHR* array carrying constitutively expressed (*P_TEF1_*)GFP(T_CYC1_) paired with *Zeo^R^*, *i.e. LHR^E^-GFP-Zeo^R^-LHR^Z^* (for precursor *v*2B) (Fig. 6). The rationale for introducing the *PDI^H^* gene, encoding His-tagged protein disulfide isomerase (PDI), into the first array is explained below. We linearized precursor *v*2B (creating nChr 2B), and precursor *v*2A (creating nChr 2A), and transformed both wild-type and Δ*KU70 K. phaffii* cells with the products. Colony PCRs (on freshly transformed cells growing on agar plates) and chromosome-loss assays (Additional file 2: Table 5) suggested initial stability of nChrs 2A and 2B in both strains. Subsequently, however, WGS and PCR genotyping revealed that nChrs 2A and 2B were stable, longer term, in Δ*KU70* strains (yDA253 and yDA218) (Fig. 5 and Additional file 1: Figs. S7 and S11) but were unstable on the wild-type background. Moreover, after protracted cell culture, the wild-type background strains that initially passed PCR-based genotyping, failed (Additional file 1: Fig. S11) despite performing well in chromosome-loss assays, suggesting integration of the marker into the genome.

**Figure 6.**
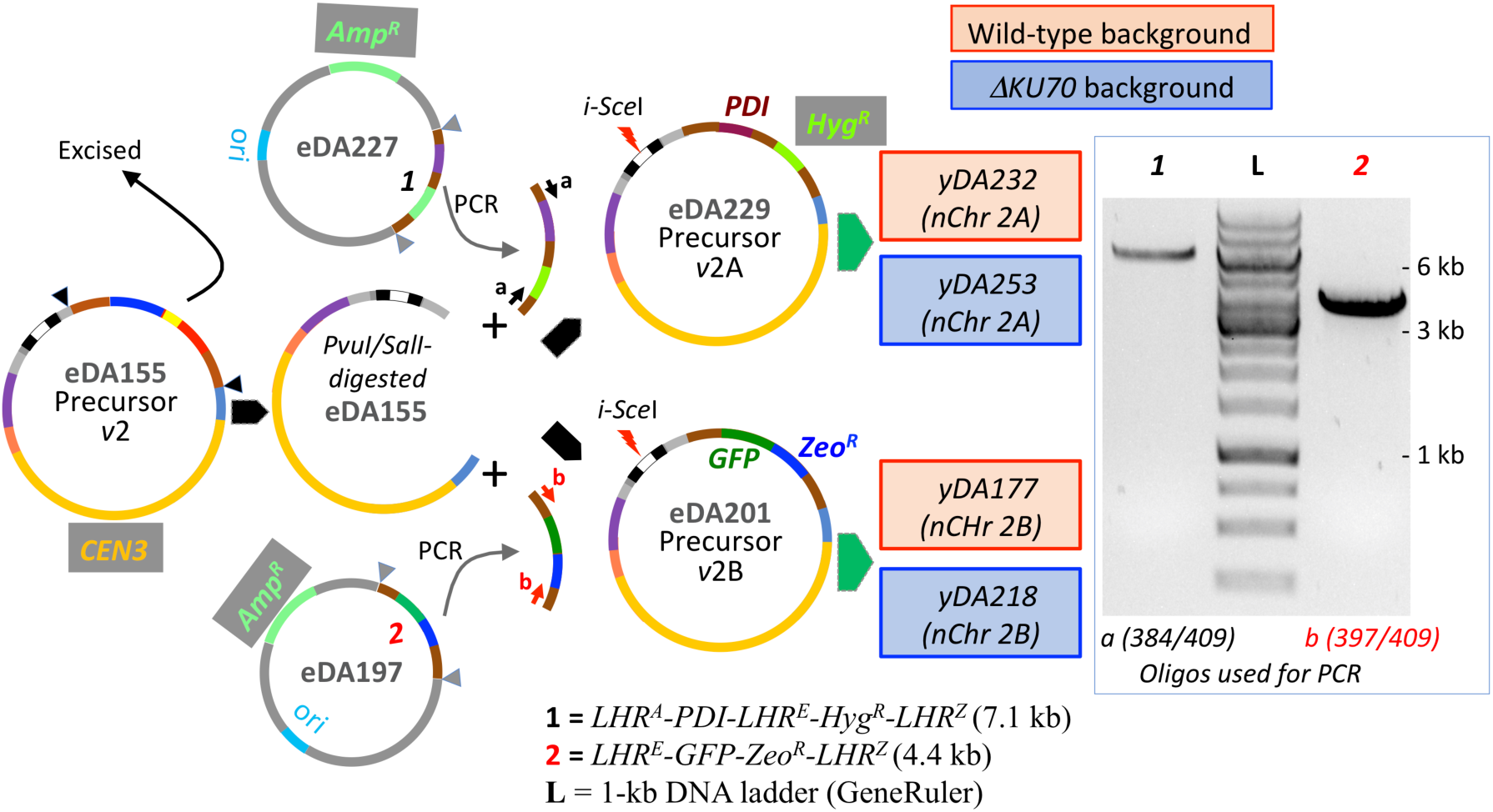
*New versions of precursor plasmids created by swapping out the original integration array.* PCR products (see agarose gel) from eDA197 or eDA227, digested with *Afe*I and *Xho*I were ligated into appropriately digested eDA155. Precursors *v*2A and *v*2B were linearized with *I-Sce*I then used to transform wild-type or Δ*KU70* cells to create a total of four strains (see also Additional file 1: Fig. S1).

### The ability to engineer nanochromosomes depended on knocking out KU70

To exemplify nanochromosome engineering, we sought to replace *GFP* and *Zeo^R^* within the integration array (*LHR^E^-GFP-Zeo^R^-LHR^Z^*) of nChr 2B, by *mCH* (codes for mCherry) and *Hyg^R^* delivered to the cell within an insertion array. We thus transformed yDA177 (despite subsequently emerging evidence for low nChr stability on the wild-type background) and yDA218 (*ΔKU70* background) with the array *LHR^E^-mCH-Hyg^R^-LHR^Z^*, aiming to create nChr 2B.1 (Additional file 1: Fig. S12a), and we selected colonies (34 from the wild-type, and 16 from *Δ*KU70 strains, in two independent experiments) on hygromycin-agar plates. We screened these for a combination of hygromycin resistance/zeocin sensitivity, and for red fluorescence/lack of green fluorescence, and then by PCR genotyping (data not shown). A total of 15 of 16 colonies on the *ΔKU70* background (including yDA226) were positively verified, but only 1 of 34 colonies on the wild-type background, passed all tests. This is consistent with the subsequently analysed PCR genotyping results and WGS analysis (data not shown) for yDA177 showing that the nChr 2B had already been compromised by aberrant fusions (Additional file 1: Figs. S11 and S12a). A qPCR analysis of yDA226 (*ΔKU70*) indicated one copy of nChr 2B.1 per cell (Additional file 2: Table 6).

In related proof-of-principle experiments we swapped *Hyg^R^* in *LHR^A^-PDI^H^-LHR^E^-Hyg^R^-LHR^Z^* of nChr 2A - in yDA232 (wild-type) and yDA253 (*ΔKU70*) - for the tandem pair *GFP-Zeo^R^* within insertion array *LHR^E^-GFP-Zeo^R^-LHR^Z^*, aiming to produce nChr 2A.1 with an extended landing zone (Additional file 1: Fig. S12b). We obtained no positively verified colonies on the wild-type background but validated (zeocin-resistant/hygromycin-sensitive, GFP-producing) eleven colonies (Additional file 1: Fig. S12b) for *ΔKU70* strains (including yDA263). Subsequent WGS of yDA263 confirmed integrity of nChr 2A.1 (Additional file 1: Fig. S7e) while qPCR was consistent with a single copy per cell (Additional file 2: Table 6).

#### Nanochromosomes support methanol-induced protein production

We investigated the potential of a nanochromosome-resident chaperonin-coding gene to facilitate protein production from a methanol-inducible heterologous gene that was integrated (using classical techniques) into the native *K. phaffii* genome (Fig. 7a). We had previously found, in *K. phaffii* cell cultures, that overexpression of *PDI* as well as a gene (*mFH*) for murine complement factor H (mFH) (both of which were genome-integrated and under control of *P_AOX1_*) was required for detectable levels of secretion of mFH, which has 40 disulfide bonds [29]. Hence, we tested if (*P_AOX1_*)*PDI^H^(T_AOX1_)* on nChr 2A could enhance mFH production in a similar manner, which would require the nanochromosome to persist throughout both growth and induction phases of the culture. We created as a positive control, nanochromosome-null strain yDA264, in which both *PDI^H^* and *mFH* are integrated into the native genome of the *ΔKU70* parental strain. For the experimental strain we took yDA250, in which only *mFH* had been integrated into the native genome, and transformed it with nChr 2A generating strain yDA260. Integrity and stability of nChr 2A in yDA260 and yDA250 was verified by PCR-based genotyping (Additional file 1: Fig. S13) and chromosome-loss assay (Additional file 2: Table 5) and WGS (Additional file 1: Fig. S7c,d). We used yDA250 (expressing mFH at a low yield, without PDI support) as a negative control.

**Figure 7.**
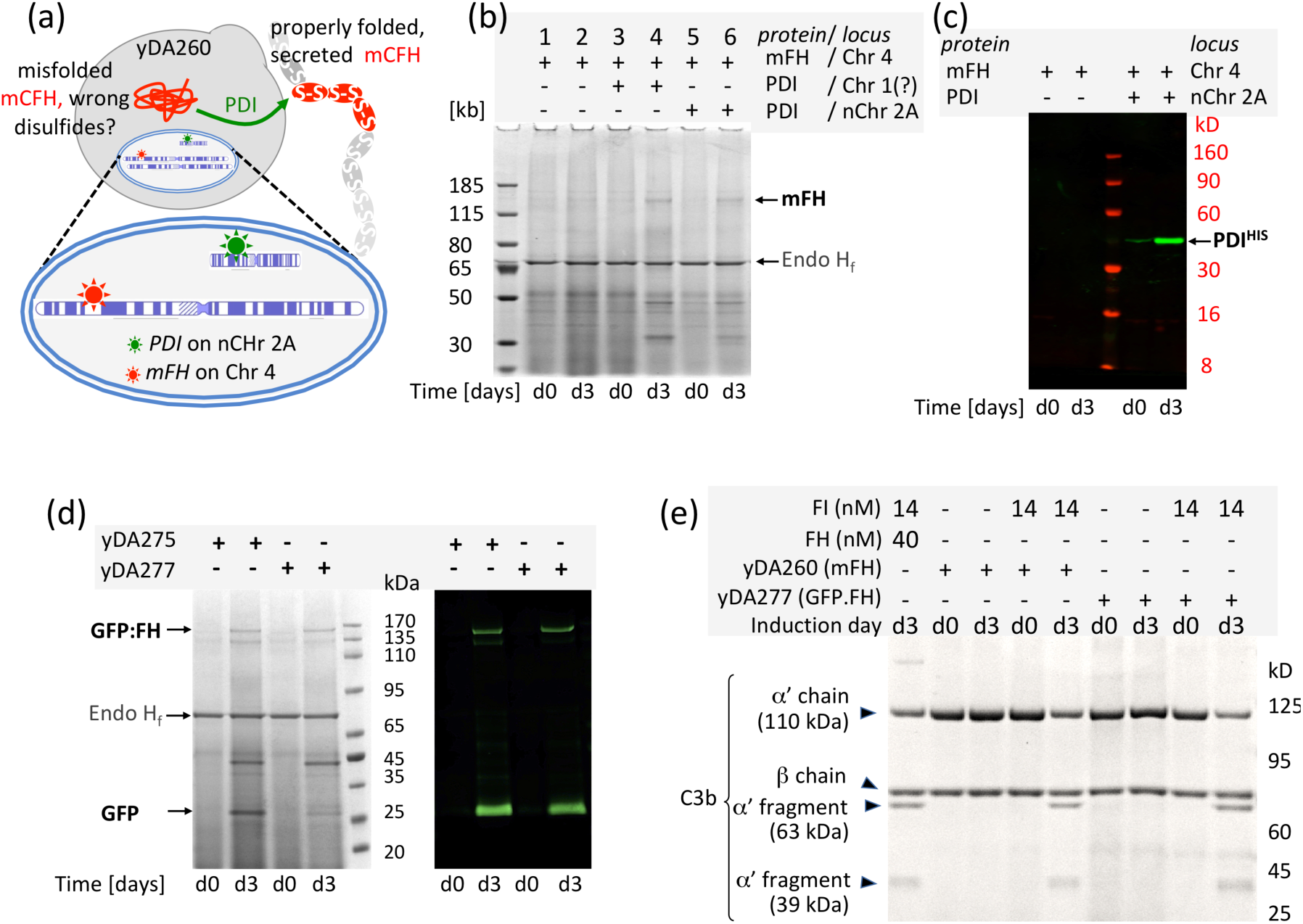
Gene expression from the nanochromosome supports production of a traditionally integrated extraneous gene, or can be used directly to produce recombinant protein. **(a)** Expression of *PDI^H^* on *nChr 2A* could assist formation of the 40 disulfide bonds of mFH encoded by DNA integrated into Chr 4. ***(b)*** SDS-PAGE (Coomassie staining) of crude cell-culture media (30 μL, reducing conditions, treated with Endo H_f_) before (d0), and three days after (d3), methanol induction. As expected, co-expression of *PDI^H^* located on either nChr 2A or (as a control) a native chromosome, is required for detectable quantities of mFH to be secreted. ***(c)*** A Western blot was used to demonstrate PDI^H^ production in cells containing nChr 2A three days post-induction. ***(d)*** A 165-kDa fusion GFP:FH fusion protein is detectable three days after induction in the cell-culture media from two strains containing nChr 2A.2 carrying both *PDI^H^* and *GFP:FH*. In each lane 40 μL of EndoH_f_–treated crude culture medium (after cell removal) was loaded under reducing conditions. Bands were detected with Coomassie blue and by Western blot using an anti-GFP primary antibody that also revealed the presence of a GFP-containing proteolytic fragment. ***(e)*** Both mFH and GFP:FH produced herein are cofactors for (30 nM) complement FI-catalysed cleavage of the 110-kDa α’-chain of complement protein C3b into 63-kD and 39-kDa fragments. In each case 12 μL of crude culture medium (at d(ay) 3 or, as a control, at d(ay) 0) was assayed. A potential contribution of non-specific protease activity is excluded by performing control reactions lacking FI.

Small-scale mFH-production tests were performed in baffled flasks, with methanol induction at day 3 (Fig. 7b). As expected, Coomassie staining of SDS-PAGE gels revealed no mFH-candidate bands for yDA250, which lacks *PDI^H^* (Fig. 7b). Conversely, a band for mFH was detected in the culture medium of *mFH*^+^ strains providing its production is supported by *PDI^H^* and regardless of whether *PDI^H^* is carried within the genome (yDA264) or on nChr 2A (yDA260) (Fig. 7b). The presence of His-tagged PDI in cultures of these strains was confirmed by Western blotting (Fig. 7c). Note that qPCR assay showed single copies of *mFH* and *PDI^H^* genes in both yDA260 and yDA264 strains (Additional file 2: Table 6). The recombinant mFH, secreted into the yDA264/yDA260 culture medium but not purified, is functionally active in an assay in which it acts as a cofactor for complement factor I (FI)-catalysed cleavage of the α’-chain of the complement protein, C3b, generating cleavage products detectable on an SDS-PAGE gel (Fig. 7e). No cleavage products were obtained in the absence of FI or when testing pre-induction cultures. Importantly, WGS validated the stability of nChr 2A (Additional file 1: Fig. S7c,d), in cells even after the induction phase, as well as the presence of a single copy of the m*FH*-expression cassette integrated into the *P_AOX1_* site on (native) Chr 4, in agreement with qPCR.

#### Demonstration of “inch-worming” for gene integration and protein production

We excised the insertion array *LHR^E^-GFP:FH-LHR^D^-Zeo^R^-LHR^Z^* (from eDA250) that includes the 4.4-kb *GoI* (*pAOX1)GFP:FH*. This *GoI* encodes a fusion protein in which green fluorescent protein (GFP) is fused to the N-terminus of human complement factor H (FH). Like mFH, this is a challenging protein to produce [29] recombinantly because it contains 40 disulfides. We used this insertion array to transform strains yDA232 (wild-type) and yDA253 (*ΔKU70*), both containing nChr 2A carrying the integration array *LHR^A^-PDI^H^-LHR^E^-Hyg^R^-LHR^Z^*. Double-crossover HR should preserve *PDI^H^* but promote insertion into nChr 2A of *GFP:FH* as well as *Zeo^R^* (replacing *Hyg^R^*), yielding nChr 2A.2 (Additional file 1: Fig. S14). It should also introduce a new landing site between *LHR^E^* and *LHR^Z^*. We selected for transformed cells with zeocin, and then screened for single colonies possessing dual zeocin resistance/hygromycin sensitivity. We then screened colonies using PCR-genotyping (Additional file 1: Fig. S14) and chromosome-loss assays (Additional file 2: Table 5). No colonies were obtained on the wild-type background but in *ΔKU70* strains, ten colonies passed screening (Additional file 1: Figs. S14 and S15). Two of these, yDA275 and yDA277 were submitted for WGS, and data analysis showed a stable, single copy of nChr 2A.2 (Fig. 5; Additional file 1: Fig. S7g, Additional file 3), also confirmed by qPCR (Additional file 2: Table 6). We cultured yDA275 and yDA277 in small-scale trials of methanol-induced GFP:FH production. Yields of protein (resolved, after treatment with EndoH_f_, on SDS-PAGE, and verified by Western blot) (Fig. 7d) were comparable with those obtained previously from strains of *K. phaffii* where *PDI^H^* and *FH* were integrated into the native genome (data not shown). Moreover, the recombinant GFP:FH secreted into the yDA275/7 culture medium is functionally active in FI-cofactor assays (Fig. 7e). Crucially, the newly created landing site in Chr 2A.2, between *LHR^E^* and *LHR^Z^*, would be available for the next of what could be multiple rounds of sequential gene insertions and hence an “inch-worming” mode of nanochromosome extension.

## DISCUSSION

We have demonstrated stable protein production from DNA on a supernumerary chromosome in a *ΔKU70* strain of the “biotechnology” yeast *K. phaffii*. This miniscule synthetic linear chromosome persists even after growing cells to high density on glucose followed by transfer to methanol-rich media for induction of heterologous gene expression. At 15-25 kb, it is smaller than even the smallest *S. cerevisiae* chromosome (250 kb), while so-called “microchromosomes” of birds and reptiles contain > 1000 kb [30]. Hence, we adopted the term “nano” chromosomes for our constructs. Telomere-capped gene-sized DNA molecules in the macronucleus of some single-celled ciliates were also called nanochromosomes, but lack centromeres and do not segregate [31]. In *S. cerevisiae*, which has highly efficient HR-based DSB-repair mechanisms, “yeast artificial chromosomes” were developed as vehicles for Mb-sized segments of foreign DNA and used early in the human genome project and for generating transgenic animals [32].

Recent work, demonstrated the feasibility of introducing centromeric and telomere-protected “neo”chromosomes into *S. cerevisiae* and then exploiting this organism’s highly efficient HR machinery to assemble multiple-gene orthogonal expression platforms [33,34] [doi: https://doi.org/10.1101/2022.10.03.510608] A study in *Y. lipolytica* achieved a similar goal despite this organism being less efficient at HR than *S. cerevisiae* and more efficient at NHEJ. In the case of *Y. lipolytica*, HR proceeded more efficiently on the synthetic neochromosome than in native chromosomes [17].

To our knowledge, ours is the first report of a stable synthetic chromosome in *K. phaffii*, and the first WGS-verified example that survives without selective pressure or the possession of essential genes. We did not establish how many copies of the nanochromosome exist per cell immediately following transformation or how many reach and enter the nucleus, but our WGS and qPCR suggest an average of one copy per cell in cultures after a few dozen generations. During mitosis, its small size will restrict the number of cohesin molecules that can encircle a nanochromosome, which might cause premature separation of sister chromatids and spindle detachment [35]. Hence it will be important to check copy numbers over time, and retain the option of transferring an essential gene onto the nanochromosome. The presence of an ARS in both arms of the nanochromosome (flanking *CEN3*) enhanced stability but we did not explore relationships between persistence and the number, spacing, sequence or positioning of ARS in a rigorous or systematic way. We have not explored nanochromosome behaviour during meiosis but this could be relevant to future prospects for using mating as a strategy for transferring it between strains. Of note is that *K. phaffii* has been suggested as a model organism for research on eukaryotic molecular cell biology. For example, its large, modular centromeres are reminiscent of those of higher organisms, unlike the 125-bp centromeres of *S. cerevisiae* [7]. Hence these and other studies of the nanochromosome (*e.g.* of chromatin structure and remodelling, and structure-function relationships of telomeres, and subtelomeric and pericentromeric regions) might have relevance for chromosome biology more widely.

On a wild-type background, analysis of WGS suggested nanochromosomes undergo translocation or fusion, creating aberrant new chromosomes. In the case of nChr 1 we observed translocation with the acrocentric *K. phaffii* Chr 3, reminiscent of centric translocation (Fig. 3). We found that nChr 1 was also unstable in *ΔKU70* strains. Compared to nChr 1, nChr 2 has a longer p-arm containing the second ARS (Fig. 1b). Interpretation of WGS results revealed that - on a wild-type background - nChr 2 formed chromosomes with more than one centromere (Fig. 4). In the breakage-fusion-bridge (BFB) cycle [36], chromosomes that lose telomeres may, following duplication and during anaphase, form end-to-end bridged dimeric chromatids [37]. Indeed, NHEJ-mediated formation of dicentric chromosomes has been reported in yeast [36, 38, 39]. Multimeric chromosomes are normally resolved, over multiple BFB cycles, because two centromeres on a mutual chromatid have a 50% chance of being pulled towards opposing spindle poles [37, 40], inducing a DSB and hence opportunities for repair via various mechanisms. In our fused nanochromosomes, the centromeres (∼10 kb apart) may be too close together for attachment to different kinetochores, so are not resolved in this way [37]. No fused versions of nChr 2 were identified in the *ΔKU70* strain. We did not yet try restoring *KU70* once the nanochromosome has become established in the nucleus.

Thus, a current limitation of our platform is the need to delete *KU70*, which encodes a DNA-binding protein, within Ku70/Ku80, that is important for NHEJ. It was reported that Δ*KU70* strains of *K. phaffii* may lack robustness and be prone to large genome deletions or depressed colony numbers when attempting genetic manipulations [41, 42]. Moreover, deletion- or insertion-promoting alternative DSB-repair mechanisms, such as microhomology-mediated end joining or single-strand annealing, might prevail when neither NHEJ (due to knockout) nor HR (due to inherent low efficiency) are adequate [43]. We encountered no such issues with our strains (Additional file 1: Fig. S9B), but did not test stability in a commercial setting, so alternatives to Ku70 depletion might be needed, and there are numerous possibilities. Recognition of DSBs by Ku70/Ku80 both recruits other NHEJ-critical proteins and blocks 5’-resection. Removing this block, by Ku70 depletion, allows resection to commence. Rapid binding of the resultant single-stranded DNA by recombinase Rad51, and its facilitator Rad52, launches HR and shuts down NHEJ [44]. Interestingly, in *K. phaffii*, genes for Ku70/Ku80 are more strongly expressed than the gene for Rad52 [42], while the inverse pertains in *S. cerevisiae* [45]. Consistently, overexpression of RAD51 and RAD52, or deletion of MPH1 (that unwinds D-loops made by Rad51) enhances HR in *K. phaffii* and improves success rates of CRISPR/Cas9-assisted gene integration [42]. A further gain in HR-mediated repair, post-Cas9 cleavage, was obtained by fusing Cas9 with endogenous Mre11 that (as part of Mre11-Rad50-Xsr2) trims back one of the strands at a DSB to create 3’-tails. Deletion of NHEJ-related protein, DNA ligase IV (Dnl4p), also improved HR efficiency in *K. phaffii* [46] and the *ΔDnl4p* strain was amenable to *in vivo* assembly of DNA fragments with concurrent integration at target sites [47]. Thus multiple alternatives, or complements, to depriving nanochromosome-possessing cells of Ku70 could be explored.

Despite the limitations of this study, our nanochromosome design has significant potential as a future robust and versatile protein-production platform. The small size of precursor plasmids required for nanochromosome construction ensures easy assembly. Insertion arrays are modular and readily constructed (Additional file 1: Fig. S6). In both cases, the process could be automated. The landing-zone design allows for effectively indefinite expansion using just two selection markers accompanied by the positioning of defined non-coding, DNA spacers between integrated genes, could facilitate metabolic engineering in *K. phaffii*. Further landing zones could, presumably, be added. There is also potential to employ, for convenience, shorter LHRs/spacers, while the prospect of *in vivo* sequential assembly, *via* transformation with overlapping insertion arrays, could be tested. The use of IRES-like sequences or bi-directional promoters to co-express multiple genes could be explored as could editing of the nanochromosome with CRISPR-Cas9.

The chemically defined nature of nanochromosomes could enhance the precision with which artificial genetic circuits and networks are assembled, facilitate progress in biocatalyst design, and improve quantitative comparisons of genetic variants. But *K. phaffii* strains containing supernumerary chromosomes will probably be most useful in biotherapeutics and vaccines applications that require expression of multiple foreign genes and where there is a mandate to cost-effectively biomanufacture large quantities of homogenous high-quality materials. Examples might include proteins with complex and extensive post-translational modifications (PTMs) such as disulfides and N-glycans, multi-protein or multi-subunit complexes [27] and virus-like particles [40]. For example, a library of selectively inducible genes for chaperonins and PTM-pathway enzymes, such as glycosyltransferases [48], could be installed on an adjunct or “supporter” nanochromosome. Subsequently, a codon-optimised gene coding for a protein of choice could be integrated into the native genome, using classical techniques. Alternatively, a bespoke array of heterologous genes could be positioned in the nanochromosome’s landing zone without engineering the native genome at all. We demonstrated minimalist versions of both strategies in the current study. Secreted yields of mFH, from a codon-optimised gene within the native genome, became detectable following *PDI* expression from a supporting nanochromosome, while human (GFP-tagged) FH and (His-tagged) PDI were co-produced from adjacent genes on a bespoke nanochromosome. To maximise yields of recombinant proteins, the following could be varied and optimised, potentially in an automated fashion, with more control and precision compared to equivalent investigations in native chromosomes: promoter selection, synonymous codon usage, gene-copy numbers, spacing between genes, and the positioning of landing zone(s) relative to centromere, ARS, and telomeres.

## CONCLUSIONS

In conclusion, we have added the biotech-friendly yeast *K. phaffii* to a small but growing list of organisms in which a synthetic linear chromosome serves as a stable source of genetic information that is orthogonal to the native genome. While highlighting the need for further development and optimisation, we have demonstrated a potential application of nanochromosome-containing strains of *K. phaffii* for the manufacture of challenging-to-make proteins.

## METHODS

### Materials

The following were sourced from New England Biolabs (NEB): type II and type IIS restriction enzymes, Q5 high-fidelity DNA polymerase, T4 and T7 DNA ligases, T4 DNA polynucleotide kinase and Antarctic phosphatase. Synthetic genes were ordered from GeneArt and plasmids purchased from Thermo Fisher Scientific. Oligonucleotides (oligos) (Additional file 2: Table 1) were ordered from Integrated DNA Technologies. *E. coli* TOP10 cells purchased from Thermo Fisher Scientific were used for sub-cloning. Wild type *K. phaffii* cells were sourced from the ATCC strains library. Non-coding DNA segments of nanochromosomes, used for HR and as fillers or spacers, were derived from a proprietary library of 8-kb synthetic DNA chunks deposited in a GeneArt plasmid (ordered from Thermo Fisher Scientific) that was custom designed for construction of yeast synthetic chromosomes (unpublished).

### Gibson Assembly

Gibson-assembly reactions were performed according to manufacturer’s (NEB) protocols. Sanger sequencing allowed verification of all DNA constructs and gel-extracted PCR products (Azenta Life Sciences). Plasmidsaurus (SNPsaurus) performed full-length sequencing of plasmids. All DNA constructs sequences are available in Additional file 3.

### Transformations

Rubidium chloride-preparation of chemically competent TOP10 *E. coli* cells (Thermo Fisher Scientific), and their heat-shock transformations with DNA, was conducted according to standard protocols (http://currentprotocols.onlinelibrary.wiley.com). Transformed *E. coli* cells were selected on LB agar medium with an appropriate antibiotic. Transformations by electroporation of *K. phaffii* cells with precursor plasmids were performed according to a modification of a published protocol [49]. Briefly, 1-10 μg of DNA in 10 μL were added to 90 μL of pretreated competent *K. phaffii* cells prior to electroporation, and the volume adjusted to 1.0 mL with yeast extract peptone dextrose (YPD) medium containing 1.0 M sorbitol (YPDS). Following recovery (four hours, 30 °C), cells were spread on YPDS agar plates with a suitable antibiotic and incubated (30 °C) for between two and (exceptionally) five days. Single colonies were then transferred to fresh YPD agar plates in preparation for PCR-based genotype screening. Finally, verified strains were grown on YPD medium ahead of the next experimental step or for deep-freeze storage in glycerol.

### Cell cultures

Cultures of *E. coli* were performed (30 °C) on LB medium supplemented with antibiotic at 50 μg/mL in the cases of zeocin and ampicillin or 100 μg/mL for hygromycin. Plasmid- and DNA-gel extractions were done with kits (Qiagen) as per manufacturer’s protocols. Yeast cells were cultivated on yeast extract, peptone with a carbon source that depended on the desired outcome (glucose for propagation; glycerol or methanol for induction). This was supplemented with 0.3 mg/mL zeocin or hygromycin.

### Yeast genomic DNA extraction

Extraction of genomic DNA (gDNA) from *K. phaffii* cells was carried out according to a protocol supplied with the Epicentre MasterPure Yeast DNA-Purification Kit from Bioresearch Technologies, but modified by inclusion of an additional RNase A treatment and a purification step using Zymo-Spin III columns (Zymo-Research). Purity of gDNA was assessed using either a Denovix DS-11 spectrophotometer or the Qubit 3 fluorometer (Thermo Fisher Scientific) and dsDNA Broad Range Assay Kit (Thermo Fisher Scientific).

### PCR-based genotyping of *K. phaffii* strains

Yeast-colony PCR was performed using green GoTaq Master Mix (Promega) as per the manufacturer’s protocol (dx.doi.org/10.17504/protocols.io.bp2l69p95lqe/v1) and employing oligos (Additional file 2: Table 1) as forward and reverse primers to target appropriate regions of nanochromosomes or their precursors. The sizes of the resulting amplicons were confirmed on agarose gels.

### qPCR for gene-copy number estimation

Estimates of gene-copy number were obtained using qPCR reactions, and according to the Luna Universal qPCR Master Mix protocol (NEB). Reaction volumes of 10 μL, in 96-well plates, were analysed using the StepOnePlus real-time PCR System (Thermo Fisher Scientific). Values for *C*_T_ were inferred from two biological repeats with three technical duplicates, and calculated using StepOnePlus software. gene-copy number values were inferred by comparisons with two “housekeeping” genes, *ACT1* and *PDH-PDA*, assumed to be single-copy.

### Chromosome-loss assays

A protocol similar to one reported previously [50] enabled quantification of nanochromosome loss over multiple mitotic events. Briefly, *K. phaffii* cells with the nanochromosome were initially inoculated into rich media containing the appropriate antimicrobial and incubated (30 °C) overnight in a shaking incubator. Cells were then transferred to a medium [1% w/v yeast extract (Sigma-Aldrich), 2% w/v peptone (Sigma-Aldrich), 0.7% w/v yeast nitrogen base without amino acids (Formedium Ltd) with 2% v/v glycerol and 1% v/v methanol], to emulate inducible recombinant protein production, with no antibiotic, and incubated (30 °C) for at least ten cell divisions. Then triplicate aliquots (500-600 cells) were spread on agar plates with or without antibiotic. Retention of AMR was equated to retention of the nanochromosome (on which the *AMR* gene had been delivered into the cell). Hence the numbers of colonies growing on the two plates were counted and compared, with the help of ImageJ64 software [51]. Note that this assay does not distinguish between scenarios in which *(i)* the *AMR* gene remains on the nanochromosome *versus (ii)* its integration into the native genome.

### Whole-genome sequencing

Illumina NextSeq (Novogene) performed WGS to generate “short” 150-nucleotide paired-end reads. Samples of gDNA were prepared as above. WGS-library preparation, and bioinformatic analysis, were performed by Novogene. The Integrative Genome Viewer 2.12.2 software (Broad Institute) was used for visualization, in Genome Browser, of mapped WGS data. Alternatively, BigWigs-formatted files were created from bedGraph files. Coverage plots were generated with custom R scripts based on data extracted from BAM files. Values were normalized to the mean in order to visualize relative coverage. To smoothen graphs a moving-window average function was deployed with a window of 2.5 kb and a step size of 200 bp. For longer reads, WGS was performed using a MinION Mk1B device (Oxford Nanopore Technologies). Briefly, gDNA was obtained using the NucleoBond HMW DNA kit (Macherey-Nagel) as per manufacturer’s guidelines, using lyticase (Sigma) for cell lysis and incubated (one hour, 37 °C) in Y1 buffer (1.0 M sorbitol, 100 mM EDTA, pH 8.0, 14 mM β-mercaptoethanol). DNA quality and concentration were assessed using a Qubit 3 Fluorimeter. Library preparation for Nanopore sequencing was performed using the Ligation Sequencing Kit SQK-LSK109 with Native Barcoding Kits EXP-NDB104 and EXP-NDB114 for multiplexing (Oxford Nanopore Technologies). Kits were used according to the manufacturer’s guidelines, except that DNA shearing was not applied and hence input DNA was increased fivefold to match the molarity expected in the protocol. Sequencing on the MinION Mk1B used MinION Flow Cells [FLO-MIN111 (R10)] or Flongle Cells [FLO-FLG001 (R9.4.1)]. Base calling was performed using Guppy [version:6.0.1+652ffd179] and reads mapped against the reference using minimap2 [version: 2.17-r941] [52]. Canu [version: 2.2] was used for *de novo* assembly [53]. All raw reads are deposited at BioProject PRJNA971544 and an overview of de novo assembly sequenced strains may be found in Additional file 2: Table 3 and Additional file 3.

### Small-scale protein production

Cell culture for small-scale tests of recombinant protein production was performed as described [29]. To assess production of GFP:FH, samples were taken from the culture medium, after spinning out cells, and Endo H_f_-treated to trim back yeast N-glycans. Then SDS-PAGE loading buffer was added and gel electrophoresis performed under reducing, denaturing conditions using 4-12% w/v Bis-Tris NuPAGE gels (Thermo Fisher Scientific) with Coomassie Blue staining. Bands of appropriate size were verified by Western blot (electrophoretic wet transfer performed on nitrocellulose membranes, in Tris-CAPS buffer using, the BioRad Mini-PROTEAN system) with anti-GFP primary antibody (Abcam, ab38689) and IRDye 800CW donkey anti-mouse IgG as a secondary (Li-COR Biosciences). Samples of secreted mFH were similarly prepared and analysed, but gel bands were identified by reference to a sample of human plasma-derived FH (Complement Technologies). To identify recombinant His-tagged PDI, cell pellets were lysed in SDS-PAGE-loading prior to gel electrophoresis. Bands were verified by Western blot, using anti-His tag monoclonal Ab, H1029 (Scientific Labs) as a primary, and IRDye 800CW as a secondary antibody. Signal was detected on the Odyssey CLx Imaging System (Li-COR Biosciences).

### Assay of FH cofactor activity

Recombinant mFH was assayed for its activity as a cofactor for cleavage of the human C3b α-chain by human FI. The 21-μL reaction volume contained 1.7 μM C3b, 14 nM FI (Complement Technology) in PBS buffer, pH 7.0, and various volumes of culture medium. The reaction mixture was incubated (one hour, 37 °C), before addition of SDS-PAGE loading buffer. After boiling (five minutes), electrophoresis was performed on NuPAGE (4-12%) gels and stained with Coomassie Brilliant Blue G250. Densitometry values inferred using Image Lab software (Bio-Rad). Plasma-derived human FH, at 10-80 nM, served as positive control and for calibration.

### The construction of parts

Note that all DNA parts used for assembly of nanochromosome precursors are free of restriction sites for *Sal*I and/or *Xho*I (needed to produce compatible cohesive ends), as well as *AsiS*I*, Bsa*I and/or *BsmB*I (used in Golden Gate-like assembly). Oligonucleotides (oligos, see Additional file 2: Table 1). Plasmids are listed in Additional file 2: Table 2. Selected plasmids were sequenced; DNA sequences are available in Additional file 3. Additional file 1: Fig. S1 includes a schematic summarizing the workflow used to generate our nanochromosomes.

To create the centromere part (*CEN3*), a DNA sequence corresponding to the centromere of *K. phaffii* Chr 3 was generated initially as two segments, in separate PCRs, using *K. phaffii* genomic DNA as a template. The 3.2-kb segment 1 was amplified (oligos 197/198) and ligated between pUC19 *Kpn*I and *EcoR*I sites, generating plasmid eDA10. The 3.1-kb segment 2 was amplified (oligos 199/200) and ligated into eDA10 between *Eco*RI and *Nde*I sites. This generated plasmid eDA24 carrying the 6.3-kb *CEN3*.

Based on previous reports [54], two *K. phaffii* ARSs[25] [26] were eventually selected to serve as origins of replication, one per nanochromosome arm. A 233-bp DNA segment corresponding to *PARS*-A76 was PCR-amplified from *K. phaffii* gDNA (oligos 217/218) and inserted into the *BamH*I site of pUC19 creating eDA26. Separately, a 530-bp molecule equating to *PARS-B413* was likewise PCR amplified (oligos 238/239) and inserted into pUC19 yielding eDA41.

Expression cassettes for AMR selection-marker were amplified from commercial vectors. *Zeo^R^*, between *P_TEF1_* and *T_CYC1_*, was PCR-amplified from pGS-GnT-I (a GlycoSwitch plasmid) (oligos 308/309), digested with *Sal*I and *BamH*I and inserted into *Sal*I/*BamH*I-linearized eDA26, creating eDA40. *Hyg^R^*, between *P_TEF1_* and *T_TEF1_*, was amplified from pGS-GnT-II (Glycoswitch) (oligos 306/307), digested with *Sal*I and *BamH1* and inserted into *SalI/BamH1*-linearised eDA26 yielding eDA37.

The telomeres part (*Tel*) comprises an 18-bp *I-Sce*I-cleavage site flanked by arrays of 16 inverted telomere repeats [26]. To construct it, two pairs of complementary oligos were annealed separately: *(i)* Oligo 246 (127-nt) with oligo 247 (120-nt); *(ii)* oligo 248 (121-nt) with oligo 249 (128-nt). In each case, oligos (100 μM) were incubated (30 minutes, 37 °C) with five units of T4 polynucleotide kinase in T4 polynucleotide kinase buffer supplemented with 2.5 mM ATP. Then, NaCl was added to 50 mM and the mixture heated (300 s, 95 °C), then cooled to room temperature. The oligos were designed so that annealing would yield two ds-DNA molecules with mutually complementary overhangs that, upon ligation, form the *Tel* with overhangs at its distal ends compatible with *Pvu*I. Thus, products of processes *(i)* and *(ii)*, together with *Pvu*I-linearized eDA40 (deleting its *Amp^R^* gene) were ligated (T4 ligase) to yield eDA131. After transformation of *E. coli* with eDA131, zeocin (50 μg/mL)-resistant, ampicillin (100 μg/mL)-sensitive cells were verified (colony PCR, not shown), and eDA131 confirmed (by Sanger sequencing) as the *Tel*-donor plasmid from which a blunt-ended *Tel* could be excised by *Fsp*I/*Pvu*II for insertion into nanochromosome precursor plasmids.

Codon-optimised for *K. phaffii* genes encoding fluorescent proteins were delivered in plasmids pM-GFP and pM-mCH (Thermo Fisher Scientific). Constitutive expression cassettes for GFP and mCherry (mCH) were PCR-amplified (oligos 302a/303a, targeting *P_TEF1_* and *T_CYC1_*), and overhangs released after *BsmB*I digestion. The products were ligated into plasmid eDA40, replacing the *Zeo^R^* gene between *P_TEF1_* and *T_CYC1_*. The resultant plasmids (eDA189 for GFP and eDA191 for mCH) were used to transform *E. coli*. Cells were selected on LB plates with ampicillin.

To prepare the *PDI* (coding for His-tagged protein disulfide isomerase) component, a previously synthesised *K. phaffii*-codon optimized *K. phaffii PDI* gene was re-amplified (oligos 447/448) from the in-house plasmid pPIC3.5K-*PDI* [29]. The primer 448 introduces six His-codons followed by the TAA stop-codon. After excision with *Not*I and *Mfe*I, the 1.64-kb DNA (*PDI^H^*) fragment was cloned into *Not*I and *Mfe*I-digested pPICZ A (between *P_AOX1_* and *T_AOX1_*) forming plasmid eDA226.

A gene cassette (*P_AOX1_)GFP:CFH(T_AOX1_*) for the GFP:FH fusion protein was prepared as follow: PCR-amplified *GFP* (from pM-GFP using oligos 276/277), digested with *Nsi*I was inserted into *Pst*I (*Nsi*I- and *Pst*I-recognition sites are compatible)-linearized pPICZα “B-cFH” plasmid prepared previously [55] forming eDA89. The DNA sequence for a flexible seven-amino acids linker (GGGGSNA) between fusion partners was delivered by oligo 277.

### Assembly of arrays

Integration (receptor) and insertion (donor) arrays contain LHRs alternating with gene-expression cassettes. Examples include *LHR^n^-GoI-AMR*-*LHR^Z^* and *LHR^n^-GoI-LHR^m^-AMR-LHR^Z^*, wherein (as described above): *LHRs* are unique sequences from a library, *LHRs^A-Z^*; *GoI* contains a gene plus promoter and terminator regions; and *AMR* is *Hyg^R^* or *Zeo^R^* (with its own promoter and terminator).

Arrays were built from the following pre-prepared components: *(i)* one or two different *LHR*s (other than *LHR^Z^*); *(ii)* the *GoI*s; and *(iii)* (common to all arrays) either *Hyg^R^*-*LHR*^Z^ or *Zeo^R^*-*LHR^Z^* tandem pairs. These components were prepared as follows. *(i) LHR*s were amplified from source plasmids *e.g.* 0.87-kb *LHR^E^* from eDA9, and amplicons digested with *BsmB*I or *Bsa*I to release overhangs. *(ii)* Synthetic *GoI*s were amplified from their delivery plasmid, then digested with restriction enzymes before insertion into pUC19. *(iiia)* For the *Zeo^R^*-*LHR^Z^* tandem pair, *LHR^Z^* was amplified by PCR from eDA8 and inserted into pUC19, to generate eDA101. The *Zeo^R^* was PCR-amplified from eDA40 for adjacent insertion into eDA101, to create eDA105. The resultant *Zeo^R^*-*LHR^Z^* construct was re-amplified and digested with *BsmB*I. *(iiib)* To make the *Hyg^R^*-*LHR^Z^* pair, *LHR^Z^* was generated by PCR and inserted into pUC19, yielding eDA99. The *Hyg^R^* cassette was PCR-amplified from eDA37 for insertion into eDA99 to create eDA103. An unwanted *AsiS*I-recognition site in *Hyg^R^* was modified by site-directed mutagenesis to generate the *AsiS*I-null plasmid eDA115. Then *Hyg^R^*-*LHR^Z^* was re-amplified and digested for subsequent assembly steps. See protocols [dx.doi.org/10.17504/protocols.io.4r3l271o3g1y/v1] and [dx.doi.org/10.17504/protocols.io.bp2l69p95lqe/v1] for details.

The integration array carrying constitutively expressed *GFP* and *Zeo^R^*, *i.e. LHR^E^- (P_TEF1_)GFP(T_CYC1_)-Zeo^R^-LHR^Z^*, was assembled by ligation of *LHR^E^*, a *GFP*-expression cassette from eDA189, and *Zeo^R^*-*LHR^Z^* pair from eDA105. Assembly was instigated by ligating *LHR^E^* with GFP. The gel-extracted (*LHR^E^*-*GFP*) product and the *Zeo^R^-LHR^Z^* pair were then ligated tandemly into pUC19 forming eDA197.

Insertion array *LHR^E^-(P_TEF_)mCH(T_CYC1_)-Hyg^R^-LHR^Z^* - carrying *mCH* paired with *Hyg^R^* - was assembled, by ligation, from *LHR^E^*, the *mCH*-expression cassettes from eDA191, and the *Hyg^R^*-*LHR^Z^* pair from eDA103. *LHR^E^* was first ligated with the *mCH*-expression cassettes forming *LHR^E^-mCH,* which was gel extracted. Then *LHR^E^-mCH* plus *Hyg^R^-LHR^Z^* were ligated into pUC19 forming eDA199.

A longer array, *LHR^A^-PDI^H^-LHR^E^-Hyg^R^-LHR^Z^* was assembled from: *LHR*^A^; *PDI^H^ (*encoding His-tagged *K. phaffii* PDI under *P_AOX1_*); *LHR^E^*; and *Hyg^R^*-*LHR^Z^*. First *LHR^A^- (P_AOX1_)PDI^H^(T_AOX1_)* and *LHR^E^*-*Hyg^R^*-*LHR^Z^* were prepared by pairwise ligations and gel purification. Then the two ligated parts were inserted, sequentially, into pUC19 forming eDA227.

Another triple-*LHR* array *LHR^E^-GFP:FH-LHR^D^-Zeo^R^-LHR^Z^* was formed from: *LHR^E^*; an expression cassette (*P_AOX1_)GFP:FH(T_AOX1_)* encoding GFP:FH (see above) amplified from eDA89; *LHR^D^* amplified from eDA9 and suitably digested; and *Zeo^R^*-*LHR^Z^* prepared as above. In two separate reactions, *LHR^E^* was ligated to *GFP:FH* and *LHR^D^* was ligated with *Zeo^R^*-*LHR^Z^*. Subsequently, gel-purified products, *i.e. LHR^E^*-*GFP:FH* and *LHR^D^*-*Zeo^R^*-*LHR^Z^*, were further ligated into pUC19 forming eDA250.

### Assembly of parts into framework and precursor plasmids

A circular, telomere-null, framework plasmid (eDA53, 10.5-kb) was constructed by Gibson assembly from: *(i)* a 1.46-kb part corresponding to *Zeo^R^* that was PCR amplified (oligos 253/254) using eDA40 as a template; *(ii)* a 0.57-kb PARS-76 part that was generated by PCR (oligos 255/256) from eDA26; and *(iii)* the 8.63-kb centromeric plasmid eDA24 linearized by *Sma*I/*PvuII*.

We initially engineered into eDA53 an integration array consisting of *LHR^A^*-*I-SceI*-*Zeo^R^*-*LHR^Z^* where *I-SceI* (encoding the meganuclease, *I-Sce*I) was under control of *P_AOX1_.* This initial integration array was not pre-assembled in the same way as the arrays described above, but was introduced in the following steps that converted eDA53 into 14.5-kb eDA83.

i. *Pvu*II-linearized eDA53 was ligated with PCR-generated 0.8-kb *LHR^A^* (from eDA8 template, oligos 264/273) carrying *Pvu*II restriction site yielding 11.3-kb eDA71, which was *BsmB*I-linearized creating -CCCT and -CGCG overhangs on antisense and sense strands, respectively.
ii. An *I-SceI*-expression cassette was made by amplifying the *I-SceI* gene from plasmid eDA22 (oligos 223/224, carrying *EcoR*I and *Sac*II sites), and inserting it into *EcoR*I/*Sac*II-linearized pPICZα B forming eDA27. Then, using eDA27 as a template, a 2.2-kb (*P_AOX1_*)*I-SceI*(*T_AOX1_*) cassette was PCR-amplified (oligos 267/268) and *BsmB*I-digested, creating –GGGA and –CCAA overhangs on sense and antisense strands, respectively.
iii. A 0.95-kb *LHR^Z^*part was PCR-generated from eDA9 (oligos 269/270) and *BsmB*I - digested, creating –GGTT and –GCGC overhangs on sense and antisense strands, respectively.
iv. Purified *BsmB*I-digested PCR products from steps *(ii)* and *(iii)* were ligated, using T7 DNA ligase, with gel-purified, linearized, eDA71 from step *(i).* After *E. coli* transformation, the product (eDA83) was verified by restriction-enzyme digestion and DNA sequencing.

The next step would have been to introduce a pair of proto-telomeres into eDA83. With this in mind, a blunt-ended 0.46-kb DNA segment containing the telomeres part (*Tel*) was gel-purified from *Pvu*II/*Fsp*I-digested eDA131. *Tel* was then ligated with *Fsp*I-linearised eDA83 (thereby deleting a *bla^R^* gene in eDA83). However, the ligation product was consistently lost when attempting propagation in *E. coli*.

### Creation of nanochromosomes

The *P_AOX1_* (in addition to ∼50-bp of the *I-SceI* open reading frame) in eDA83 was replaced with the bacterial kanamycin-resistance (*Kan^R^*) cassette to create 14.5-kb eDA110, containing the modified integration array we called *LHR^A^*-*Kan^R^*(Δ*I-SceI*)-*Zeo^R^*-*LHR^Z^*. To achieve this, the 13.5-kb *Sph*I-digested and gel-extracted eDA83 was ligated with a 980-bp PCR-amplified *Kan^R^* (using oligos 321/322 and vector pUC57-Kan (Addgene) as a template). Following extraction from kanamycin-resistant *E. coli* cells, plasmid eDA110 was verified by Sanger sequencing. A blunt-ended 0.46-kb DNA segment containing the *Tel* was gel-purified from *Pvu*II/*Fsp*I-digested eDA131, as before, but this time it was ligated with *Fsp*I-linearised eDA110 to yield 14.9-kb eDA137. Plasmid eDA137, verified by Sanger sequencing, was re-amplified in *E. coli* cells, linearized *in vitro* by *I-Sce*I and used to transform *K. phaffii* (wild type). Transformants were selected on YPDS plates with 0.3 mg/ml zeocin.

The plasmid eDA110 was extended by tandem insertion of *(i)* a second ARS - the 529-bp *K. phaffii ARS-B413* - and *(ii)* an extended (1.3-kb) non-coding spacer segment of DNA (*ncDNA*), as follows (See Suppl Fig 1). PARS-B413 was generated by PCR (oligos 238/239) from gDNA, and *BamH*I-digested then ligated into the multi-cloning site of pUC19 forming eDA41. Meanwhile, *ncDNA* was amplified (oligos 379/380) from eDA9 and then *Nde*I/*EcoR*I-digested for insertion immediately downstream of PARS-B413 in eDA41, generating intermediate plasmid eDA144. Subsequently, the combined 1.9-kb *ARS-B413-ncDNA* part was amplified from eDA144 (oligos 379/381), digested by *Nde*I, and inserted into the *Nde*I site of eDA110, yielding eDA146 (16.8 kb). The *Tel* was inserted into *Fsp*I-linearized eDA146, generating eDA155. Plasmid eDA155, verified by Sanger sequencing, was propagated in *E. coli* and then isolated and linearised with *I-Sce*I *in vitro*. The linear product was used to transform *K. phaffii* strain CBS7435 using zeocin for selection.

The modified integration array (*LHR^A^*-*Kan^R^(*Δ*I-SceI*)-*Zeo^R^*-*LHR^Z^*) of eDA155 was excised with *Pvu*I and *Sal*I, releasing 5’-blunt and 3’-*Sal*I/*Xho*I compatible ends. Subsequently, the PCR-amplified integration arrays *(i)* 4.5-kb *LHR^E^-GFP-Zeo^R^-LHR^Z^* (from template eDA197 with oligos 397/409), or *(ii)* 7.3-kb *LHR^A^-(P_AOX1_)PDI^H^(T_AOX1_)-LHR^E^-Hyg^R^-LHR^Z^* (from template eDA227 with oligos 384/409), *were* digested with *Afe*I and *Xho*I/*Sal*I, then ligated with the *Pvu*I/*Sal*I-treated eDA155. The products (*(i)* eDA201 or *(ii)* eDA229) were used to transform *E. coli* cells. Following sequence verification these nanochromosome-precursor plasmids were linearized *in vitro* by *I-Sce*I and the products used to transform *K. phaffii* cells, with selection on YPDS plates with 0.3 mg/mL hygromycin or zeocin.

### Proofs of principle

*(i)* The insertion array *LHR^E^-(P_TEF_)mCH(T_CYC1_)-Hyg^R^*-*LHR^Z^* was amplified by PCR (oligos 409/397) from eDA199, and used to transform *K. phaffii* cells (creating yDA177 on the CBS7435 background, or yDA218 on the CBS7435::Δ*KU70* background) already carrying nChr 2B (*i.e.* linearised eDA201). The aim here was to replace *GFP-Zeo^R^* (in *LHR^E^*-*GFP-Zeo^R^-LHR^Z^*) with *mCH-Hyg^R^*.
*(ii)* The insertion array *LHR^E^-(P_TEF_)GFP(T_CYC1_)-Zeo^R^-LHR^Z^* was amplified by PCR (oligos 397/409) from eDA197 and used to transform *K. phaffii* cells (strains CBS7435 (to create yDA232) and CBS7435::Δ*KU70* (to create yDA253) carrying nChr 2A (*i.e.* linearised eDA229). In this case the aim was to replace *Hyg^R^* (in *LHR^A^-(P_AOX1_)PDI^H^(T_AOX1_)-LHR^E^-Hyg^R^-LHR^Z^*) with *GFP-Zeo^R^*.

Transformed cells (strains yDA218 and yDA253 for *(i)* and *(ii),* respectively) were selected on YPDS plates containing 0.3 mg/ml zeocin or 0.3 mg/ml hygromycin. Thus, efficiency of gene replacement was estimated from the number of colonies that were fluorescent protein-producing, and resistant or sensitive to the expected antimicrobial.

To test our “inch-worming” strategy for gene insertion and landing zone extension, *LHR^E^-GFP:FH-LHR^D^-Zeo^R^-LHR^Z^* was excised from eDA250 using *AsiS*I. Following concentration by isopropanol precipitation, the array was used to transform *K. phaffii* cells containing nChr 2A so as to replace *Hyg^R^* (in *LHR^A^-(P_AOX1_)PDI^H^(T_AOX1_)-LHR^E^-Hyg^R^-LHR^Z^*) with *GFP:FH-LHR^D^-Zeo^R^*. Transformed cells were selected on YPDS plates with zeocin. Gene integration/replacement efficiency was estimated from the number of colonies that were green-fluorescent, zeocin-resistant, and hygromycin-sensitive.

## LIST of ABBREVIATIONS

Ab: antibody
*Amp^R^*: β-Lactamase (*bla*) resistance cassette
AMR: antimicrobial resistance
ARS: autonomously replicating sequence
*CEN3*: the centromere of (native) Chr 3 that doubles as the centromere part of all nanochromosomes
DSB: double-strand break
FI: complement factor I
eDA(x): plasmids created for this study
gDNA: genomic DNA
GFP: green fluorescent protein
GFP:FH: A GFP-human complement factor H fusion protein
GoI: gene of interest plus promoter and terminator
HR: homologous recombination
FH or mFH: human or murine complement factor H
*Hyg^R^*: hygromycin resistence, *hph* gene
*Kan^R^*: kanamycin resistance, *neo* gene
LHRs: long homologous recombination-promoting regions
mCH: mCherry
nCHr 1 etc: nanochromosome 1 etc
NHEJ: non-homologous end joining
*ncDNA*: non-coding DNA used as neutral spacer
nt: nucleotides
oligos: oligonucleotides
PDI: protein disulfide isomerase
PNK: polynucleotide kinase
PTMs: posttranslational modifications
*Tel*: telomeres part
v1, v2 etc: version 1, 2 etc of precursor plasmids
WGS: whole-genome screening
yDA(x): *K. phaffii* strains created in this study
YPD: yeast extract peptone dextrose
YPDS: YPD-sorbitol
*Zeo^R^*: zeocin resistance, *ble* gene

## Declarations

### Ethics approval and consent to participate

Not applicable

### Consent for publication

Not applicable

### Availability of data and materials

27th November 2023

### Authors’ contributions

DA helped to conceive of the project, performed most of the experimental work and much of the data analysis, and helped to draft the manuscript; PNB helped to conceive of the project, design experiments and interpret the results and draft the manuscript; MCSO, AARR, DS, DR performed long-read WGS and data analysis, and (DS) helped to edit the manuscript; DR performed analysis of Nanopore WGS; ALM and SM contributed to experimental strategy.

## Acknowledgements

Valentin Zulkower (University of Edinburgh, UK) for sharing the non-coding DNA (the source of LHRs). We thank J. Christopher Love (MIT, Cambridge, MA, USA) for sharing *K. phaffii* strain CBS7435 *ΔKU70*.

