## Supplementary material for "A supernumerary synthetic chromosome in *Komagataella phaffii* as a repository for extraneous genetic material": Abramczyk et al AdditiobalFile1.docx

Dariusz Abramczyk^1^*, María del Carmen Sánchez Olmos^2^, Adán Andrés Ramírez Rojas^2^, Daniel Schindler^2,3^, Daniel Robertson^4^, Stephen McColm^5^, Adele L. Marston^6^, Paul N. Barlow^1,4^ *

^1^ School of Chemistry, University of Edinburgh, United Kingdom

^2^ Max Planck Institute for Terrestrial Microbiology, Marburg, Germany

^3^ Center for Synthetic Microbiology, Philipps-Universität Marburg, Marburg, Germany

^4^ School of Biological Sciences, University of Edinburgh, United Kingdom

^5^ Ingenza Ltd Scotland, United Kingdom

^6^ The Wellcome Centre for Cell Biology, Institute of Cell Biology, School of Biological Sciences, University of Edinburgh, United Kingdom

Additional File 1 (Supplementary figures)

**Contents**

Supplementary Figure S**1** *The route to our nanochromosome-carrying strains of K. phaffii.*

Supplementary Figure S**2** *WGS confirmation of a distinctive nucleotide ‘marker’ on CEN3 on nanochromosome in comparison to chromosome 3.*

Supplementary Figure S**3** *Preparation of framework plasmid eDA53*

Supplementary Figure S**4** *Construction of the telomeres part* (*Tel*)

Supplementary Figure S**5** *Validation of Tel-carrying plasmid eDA131*

Supplementary Figure S**6** *Insertion and integration arrays*

Supplementary Figure S**7** *Sequencing-coverage plots for Chrs 1-4, and the nanochromosome (nChr 1, 2A etc).*

Supplementary Figure S**8** *Genotyping by PCR is consistent with nChr 2 instability on wild-type background*

Supplementary Figure S**9** *Validating the KU70-knockout K. phaffii strain employed in this study to host nanochromosomes*

Supplementary Figure S**10** *Verification of nanochromosomes version 1 (nChr 1) and 2 (nChr 2) stability in Pichia ΔKU70 cells growing in YPD medium over 20 generations*

Supplementary Figure S**11** *Comparison of PCR-based genotyping of nChr 2A and 2B on wild-type and KU70-compromised K. phaffii backgrounds*

Supplementary Figure S**12** *Antibiotic resistance genotype of wild-type (CBS7435) or ΔKU70 strains of P. pastoris after transformation with the insertion array*

Supplementary Figure S**13** *Assessment of nChr 2A persistence in K. phaffii strain yDA260*

Supplementary Figure S**14** *Illustration of the use of an “inch-worming” strategy for in vivo gene integration into the nanochromosome that requires a ΔKU70 background*

Supplementary Figure S**15** *Validation of inch-worming*

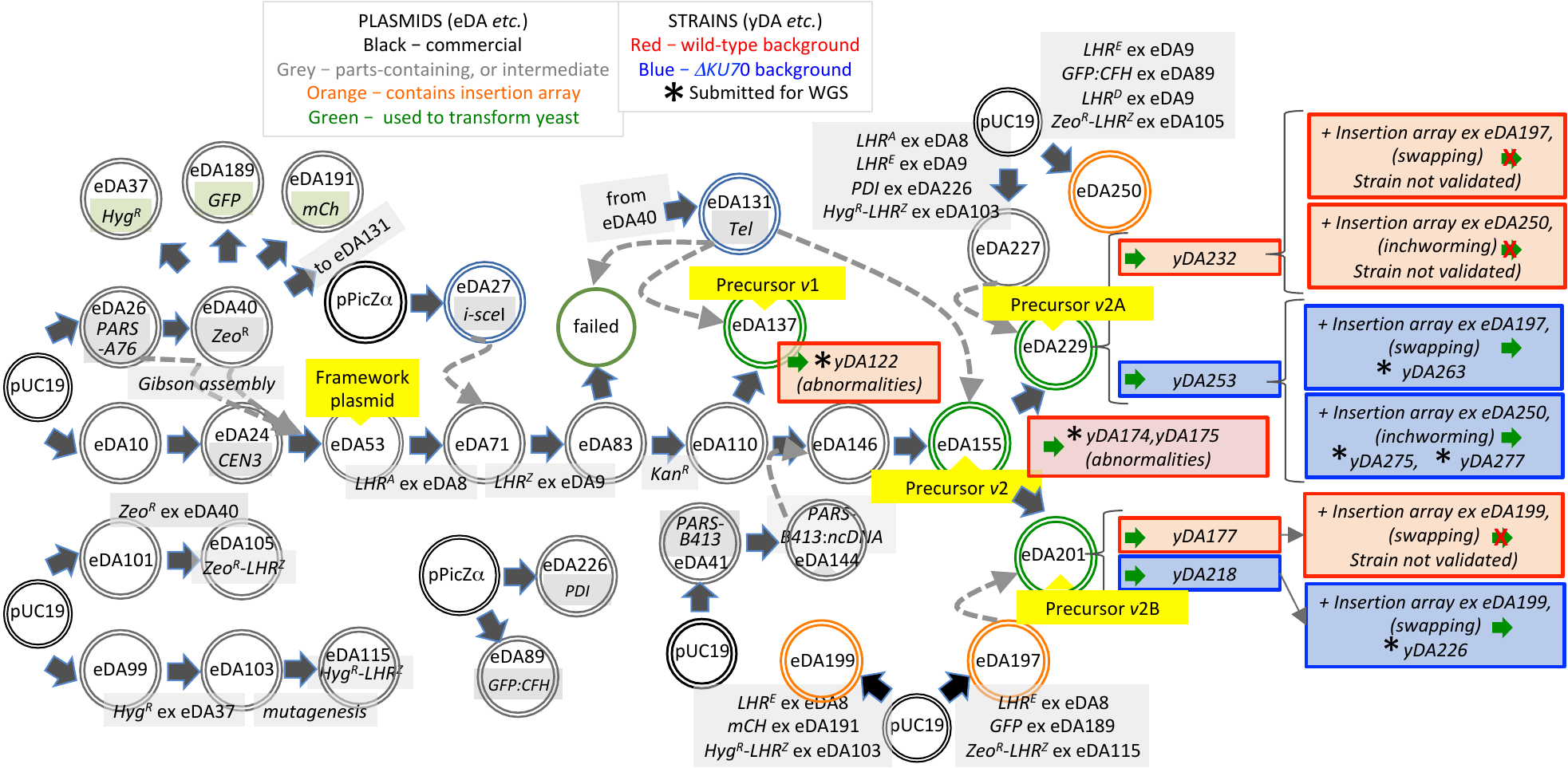

**Supplementary Figure S1**

*The route to our nanochromosome-carrying strains of K. phaffii*

This is a more detailed version of Figure 1A. Circles represents engineered in *E. coli* plasmids (eDAxxx etc). See also the list of plasmids in Additional file 2: Supplementary Table 2 and the set of nanochromosomes drawn in Figure 1B. Boxes represents *K . phaffii* strains (yDAxxx etc) that contain nanochromosomes. See also the list of strains in Additional file 2: Supplementary Table 3)

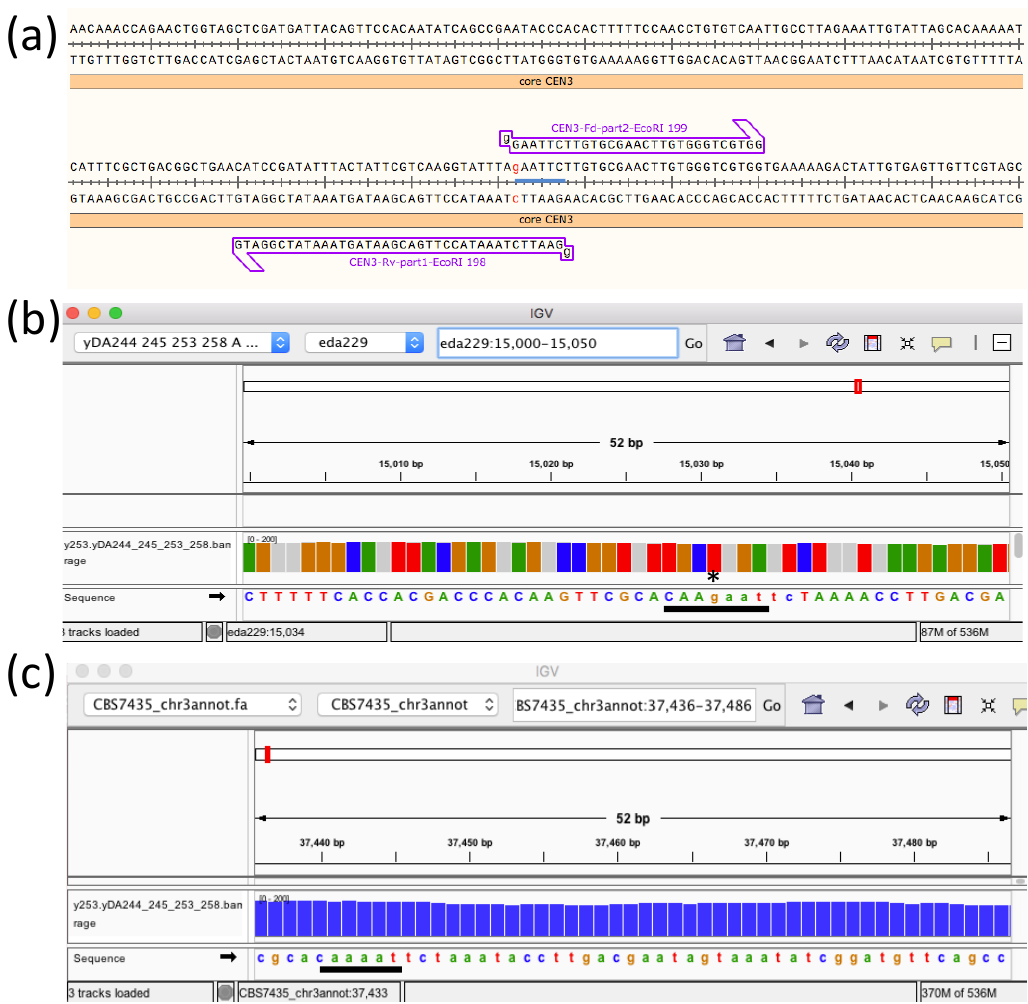

**Supplementary Figure S2**

*Confirmation of distinction between nanochromosomal CEN3 (containing additional G) and the native centromere of Chr 3*

***(a)*** A screenshot showing SnapGene representation of core region of *CEN3* from precursor plasmid *v*2A, with a G inserted to create an *EcoR*I-recoginition site (see Fig. 2) ***(b)*** Visualization (IGV) derived from WGS of nChr 2A *CEN3* core region (from strain yDA253). The black bar shows the region containing the *inserted guanosine. ***(c)*** Visulisation (IGV) derived from WGS of the core region of the native Chr 3 centromere (from strain yDA253) . The black bar indicates the sequence that distinguishes between native (CAAAAT ) and nanochromosomal (CAAGAAT) centromeres.

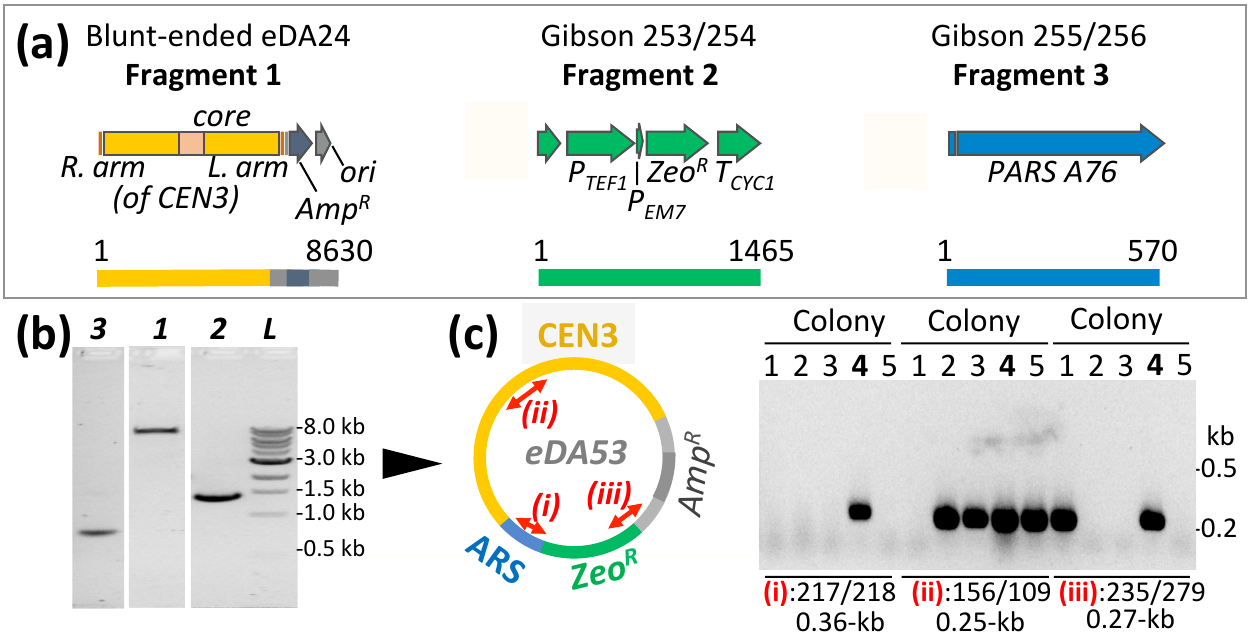

**Supplementary Figure S3**

*Preparation of framework plasmid eDA53* (see also Fig. 2)

***(a)*** The DNA parts (annotated by SnapGene Viewer) used for Gibson assembly. Fragment 1: eDA24 (source of *CEN3*) linearized with blunt-end restriction enzymes. Fragment 2: PCR-amplified Zeo^R^, ex eDA40. Fragment 3: PCR-amplified *PARS-A76*, *ex* eDA26 (see additional file 2: Suppl. Table 1 for oligo sequences). ***(b)*** Purified fragments 1-3 ran as expected on an agarose gel. ***(c)*** Map of eDA53 with location of primer-pairs used for verification by PCR. Of five *E. coli* colonies selected on plates with 50 μg/mL zeocin and 100 μg/ml ampicillin, only one colony (4, highlighted) was verified, and Sanger sequencing confirmed the expected sequence of the extracted plasmid.

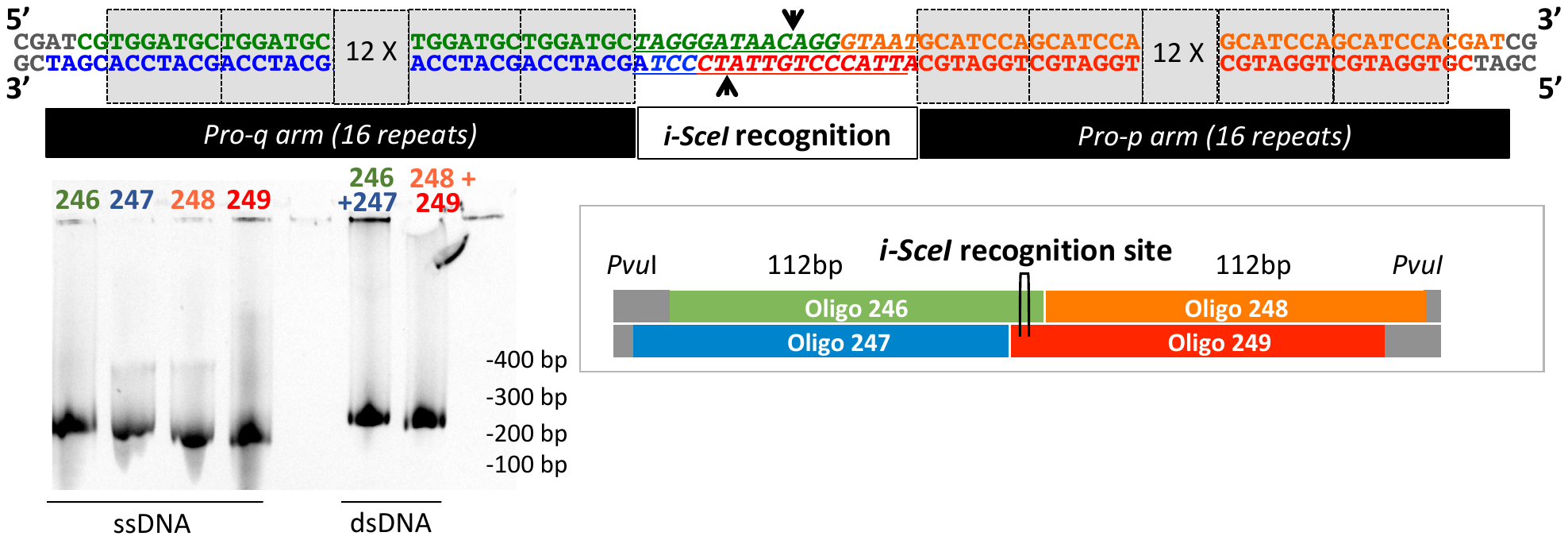

**Supplementary Figure S4**

*Construction of the telomeres part* (*Tel*) (see also Fig. 2)

The upper schematic shows the proto-telomere[*i-Sce*I-recognition site]proto-telomere structure expected after annealing and then ligation of four oligos as color-coded in the lower schematic (see additional file: Suppl. Table 1). Grey boxes represent some of the seven-base pair telomere repeats. The *Pvu*I-recognition site allows *Tel* excision from, and insertion into, plasmids. The confirmatory gel shows expected bands for the individual oligos (see schematic for numbering) and the products of annealing.

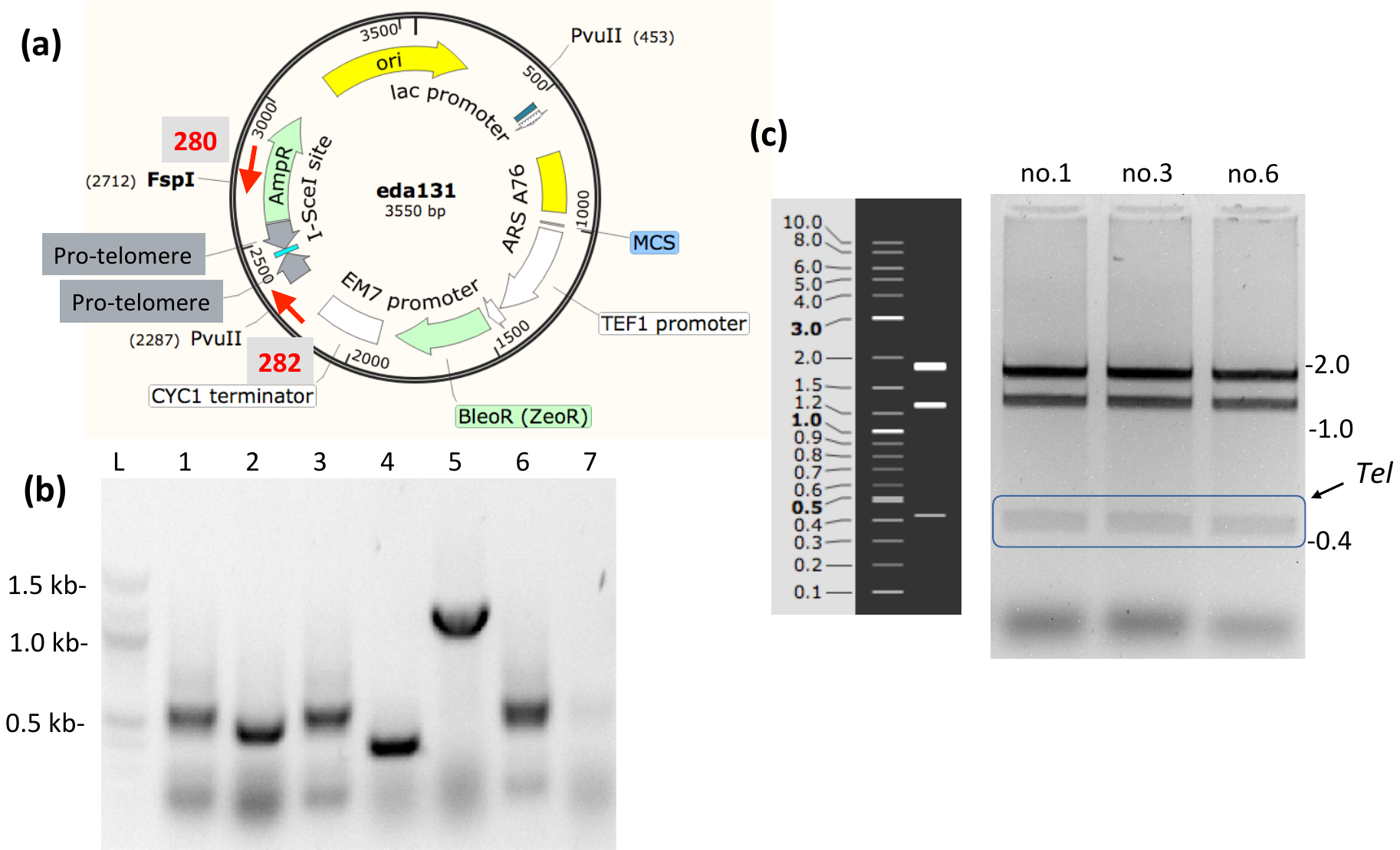

**Supplementary Figure S5**

*Validation of Tel-carrying plasmid eDA131* (see also Fig. 2)

***(a)*** Map of the eDA131 showing the locations of the various sequences targeted by oligos (red text and arrows) used for colony PCR-based screening of *E. coli* transformants, following ligation between cleaved eDA40 and *Tel*. ***(b)*** Among zeocin-resistant and ampicillin-sensitive strains, colony-PCR using oligos 280/282 generates bands corresponding to the expected (584-bp) amplicon in clones 1, 3 and 6, only. ***(c)*** Plasmids isolated from clones 1, 3 and 6, following digestion with *Fsp*I/*Pvu*II and gel electrophoresis, yielded a pattern of bands close to the one predicted (by SnapGene). The *Tel* part could thus be digested with blunt-end restriction enzymes *Pvu*II and *Fsp*I, then gel-extracted prior to ligation with framework-plasmid eDA53.

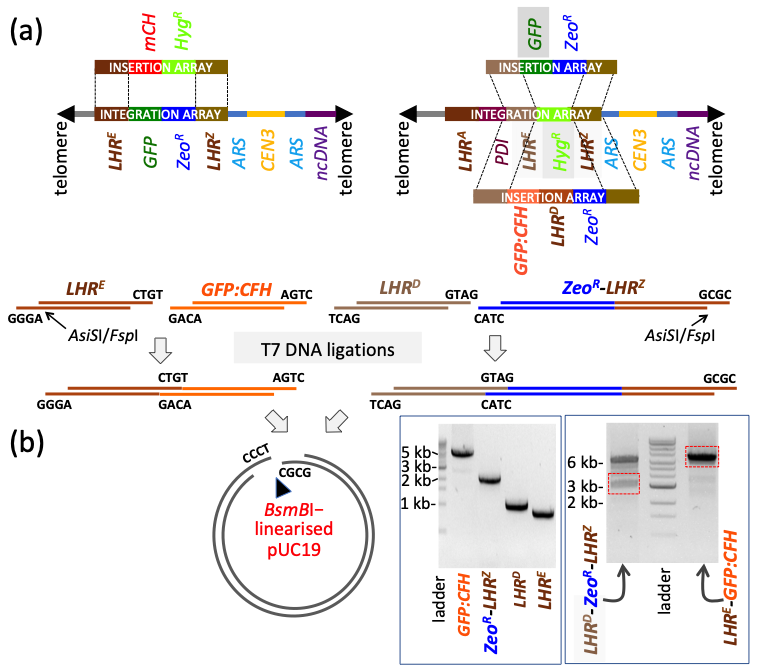

**Supplementary Figure S6**

*Insertion and integration arrays*

***(a)*** Various strategies for integration *via* double-crossover HR of genes, delivered within an *in vitro*-assembled insertion array, into a *K. phaffii* nanochromosome-resident integration array. ***(b)*** An example of array assembly. PCR-amplified parts digested with *Bsm*BI were gel-purified then ligated pairwise. Products were gel-purified and ligated into *Bsm*BI-linearised pUC19. Resultant plasmids were used to transform *E. coli* then Sanger sequenced. In preparation for deployment, arrays were subsequently excised with *AsiS*I, or amplified by PCR (dx.doi.org/10.17504/protocols.io.bp2l69p95lqe/v1). Additional file 2: Table S2 contains a list of insertion and integration arrays.

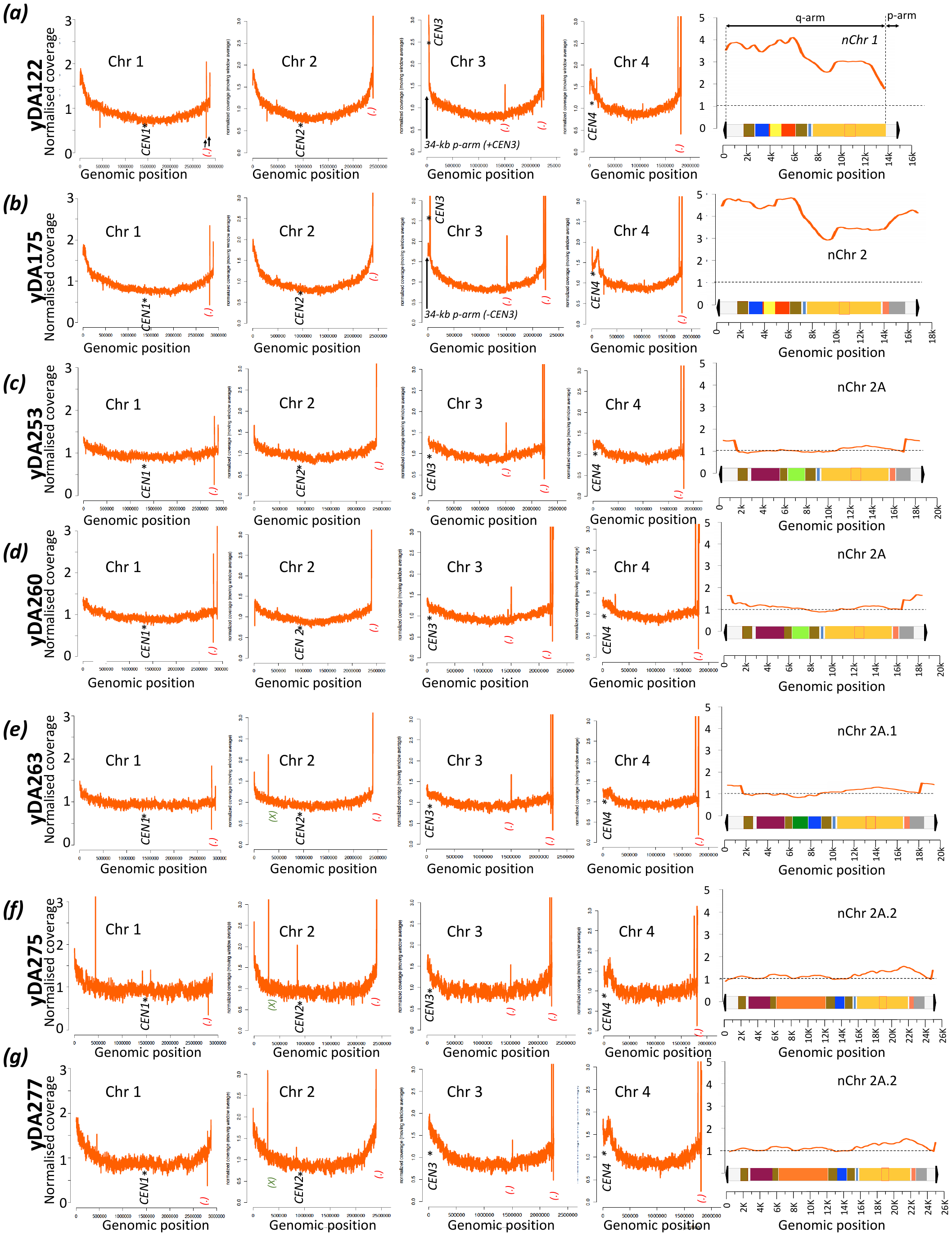

**Supplementary Figure S7**

*Whole-genome sequencing coverage data for wild-type and ΔKU70 K. phaffii cells containing nanochromosomes* (This is an extended version of Fig. 5)

See legend of Figure 5 in main text for an explanation of axes and symbols. ***(a)*** and ***(b):*** For strains on wild-type background, plots of normalized coverage indicate two-four copies of nanochromosomal sequences per cell. Note in ***(a)*** that nChr 1 of yDA122 lacks a p-arm, while high coverage suggests there are between to and four copies, per cell, of a majority of the nChr 1 sequence. This result is compatible with *de novo* assembly (details described in the Results section, sequences deposited in Additional file 3) suggesting fusion between nChr1 q-arm and the ~34-kb Chr 3 p-arm. A vertical arrow in the Chr 3 plot indicates data consistent with duplication of the Chr 3 p-arm and its centromere. In ***(b)***, the values (>2) for normalised coverage for nChr 2 suggest multiple copies per cell of its DNA content. The high copy-number for the Chr 3 centromere correlates with a high copy-number of nChr 2 and supports the existence of chimeric multi-centric nanochromosomes (a tri-centric model is suggested in Additional file, Suppl. Fig. X). This observation is compatible with yDA175 and yDA177 *de novo* assembly results obtained from long-read WGS (Suppl. file DNA.zip). ***(c)-(g):*** Strains on a Δ*KU70* background are each consistent with a single copy per cell of its nanochromosome, specifically: ***(c),*** yDA253 with a single copy of nChr 2A; ***(d),*** yDA260 with *mFH* gene integrated into (native) Chr 4 and a single copy of nChr 2A; ***(e)***, yDA263 with a single copy of nChr 2A.1 (*i.e* after replacing *Hyg*^R^ in nChr 2A with *GFP-*Zeo^R^); ***(f)*** and ***(g),*** yDA275 and yDA277 (biological replicates) with single copies of nChr 2A.2 (*i.e.* after the inch-worming proof-of-principle experiment). [*(BioProject PRJNA971544)*]

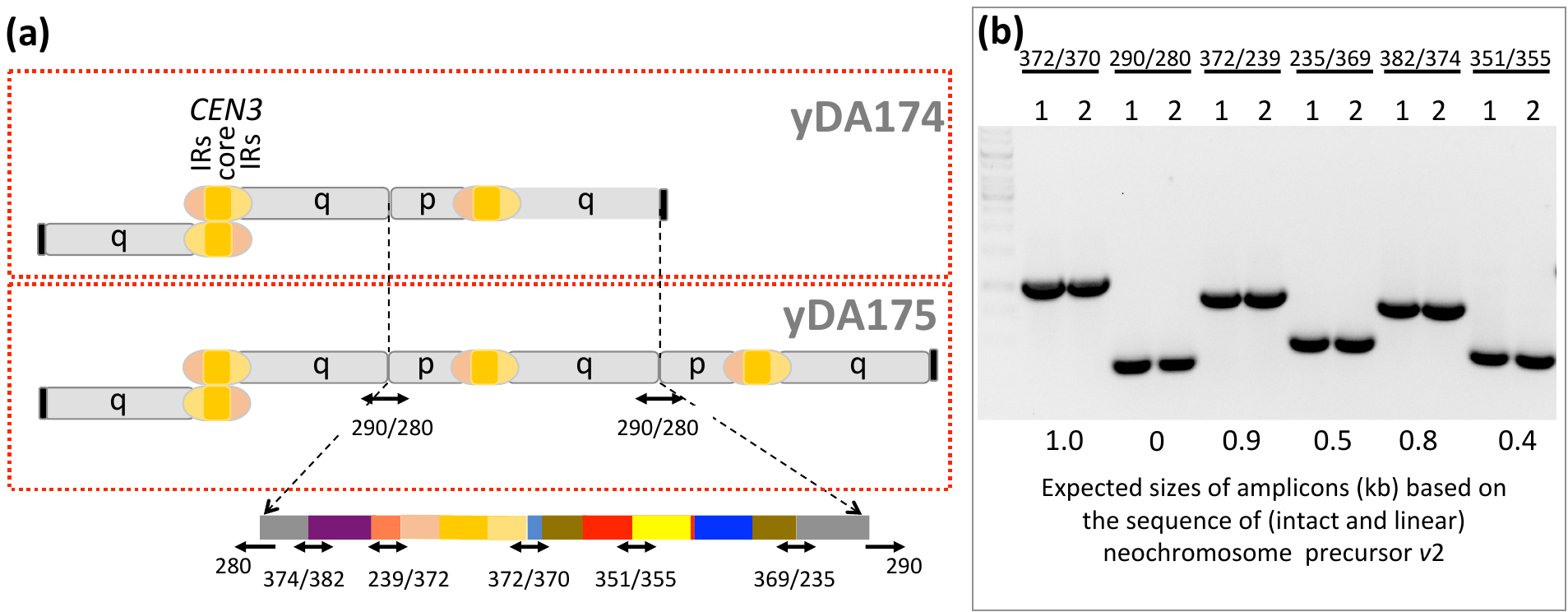

**Supplementary Figure S8**

*Genotyping by PCR is consistent with loss of nChr 2 stability on a wild-type background* (see also Fig. 4d)

***(a)*** Schematic of multi-centric chromosomes inferred from performing *de novo* assembly analysis of the WGS of two strains of *P. pastoris* created by transforming wild-type cells with nChr 2, followed by cell culture for multiple generations. A color-coded (not to scale, see Fig. 1B) map of nChr 2 (telomeres not shown) is drawn below with the sites targeted by oligo pairs (see Suppl. Table 1) for PCR-based genotyping. ***(b)*** Agarose gel showing PCR amplicons derived from the oligo pairs shown in panel A; (1) for yDA174, (2) for yDA175. The oligo pair 290/280 (lanes 3 and 4 on the gel) yields a ~0.4 kb amplicon, consistent with chromosome fusion. This would not have been detected had nChr 2 remained intact.

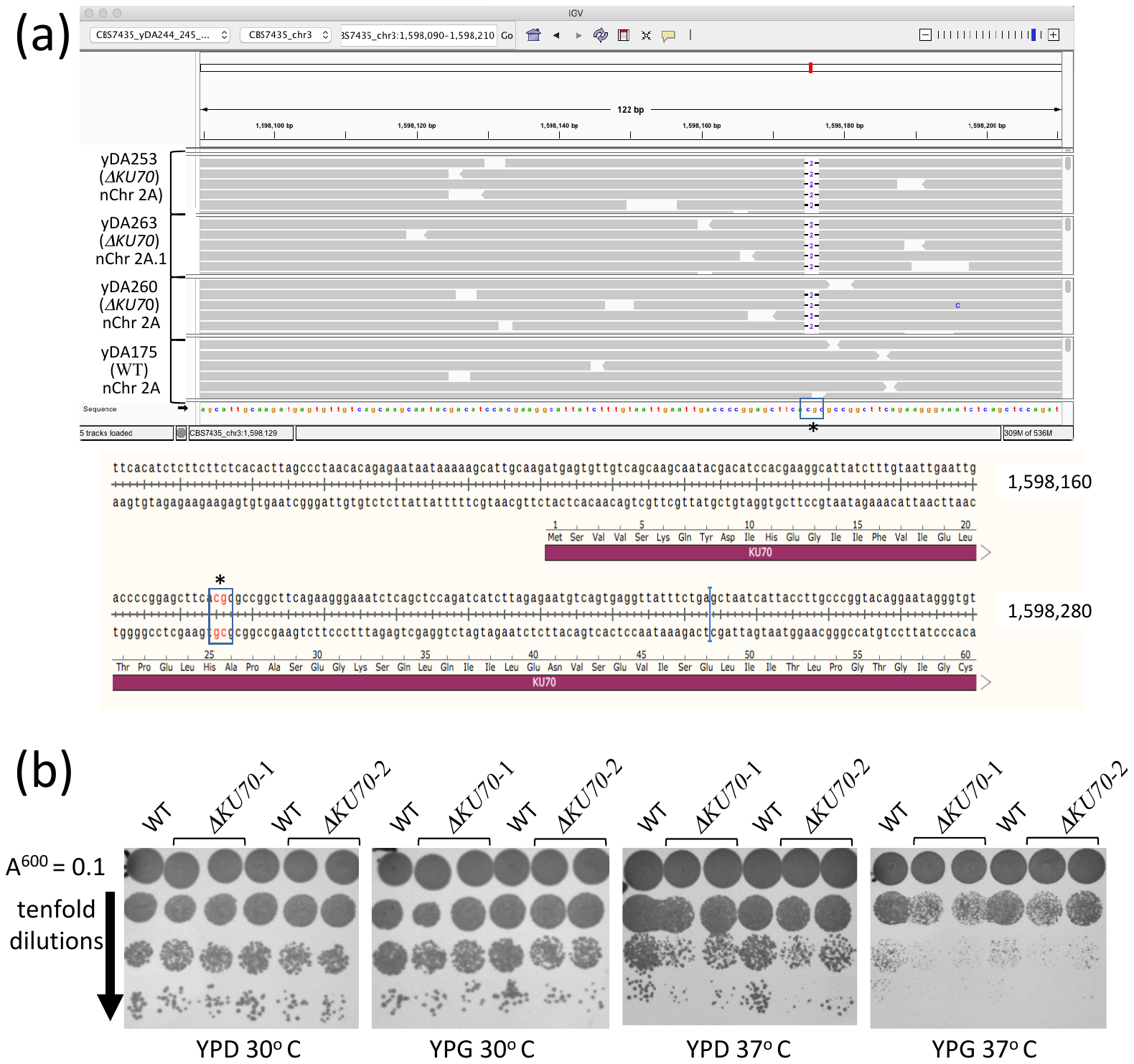

**Supplementary Figure S9**

*Validating the KU70-knockout K. phaffii strain employed in this study to host nanochromosomes*

***(a)*** Upper box: Annotated visualisation (IGV) of WGS results confirms deletion of CG dinucleotide spanning codons for *KU70* ^25^His-Ala^26^. The *KU70* start codon is indicated with a black rectangle. Lower box: Annotated SnapGene screenshot showing the sequence *of KU70* (on Chr 3) and highlighting the dinucleotide (GC) deleted in our *ΔKU70* strain. ***(b)*** Serial dilution to compare viabilities of wild-type and *ΔKU70* strains under various conditions. CBS7435 (WT) cells and two *ΔKU70* isolates, yDA208 (*ΔKU70*-1) and yDA210 (*ΔKU70*-2) were inoculated into YPD and grown for 24 hours at 30 °C. Cultures were adjusted to A^600^ = 0.1, then subjected to a series of tenfold dilutions for plating onto YPD or YPG (glycerol) agar. Plates were incubated at 30 °C for two days, or 37 °C for four days.

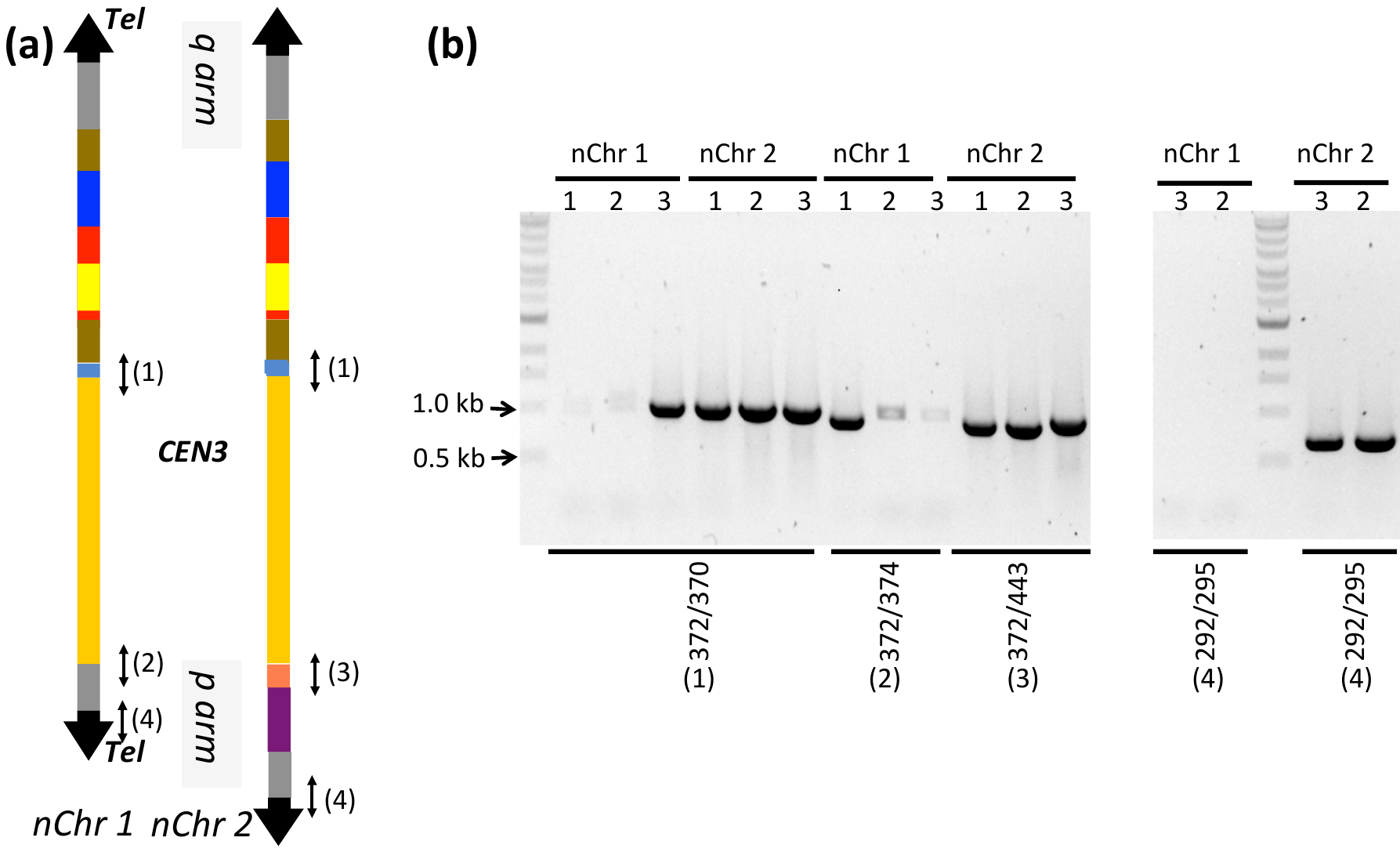

**Supplementary Figure S10**

*Verification of nChr 1 and nChr 2 stability in ΔKU70 cells growing in YPD medium for 20 generations*

PCR-based genotyping confirms integrity of extended version (nChr 2) in contrast to shorter (nChr 1) version for which one or more crucial bands are absent from each strain. ***(a)*** Primers were designed to target regions linking *CEN3* and the q-arm of both nanochromosomes (370/372; primers set 1) or the p-arm of either nChr 1 (372/374; set 2) or nChr 2 (372/443; primers set 3). Primer set 4 (292/295) validates the p-arm telomere of both nanochromsomes. ***(b)*** Note the absence of an amplicon corresponding to the p-arm telomere of nChr 1, in contrast to nChr 2. An amplicon for set 2 is missing or weak in two out of three colonies transformed with nChr 1.

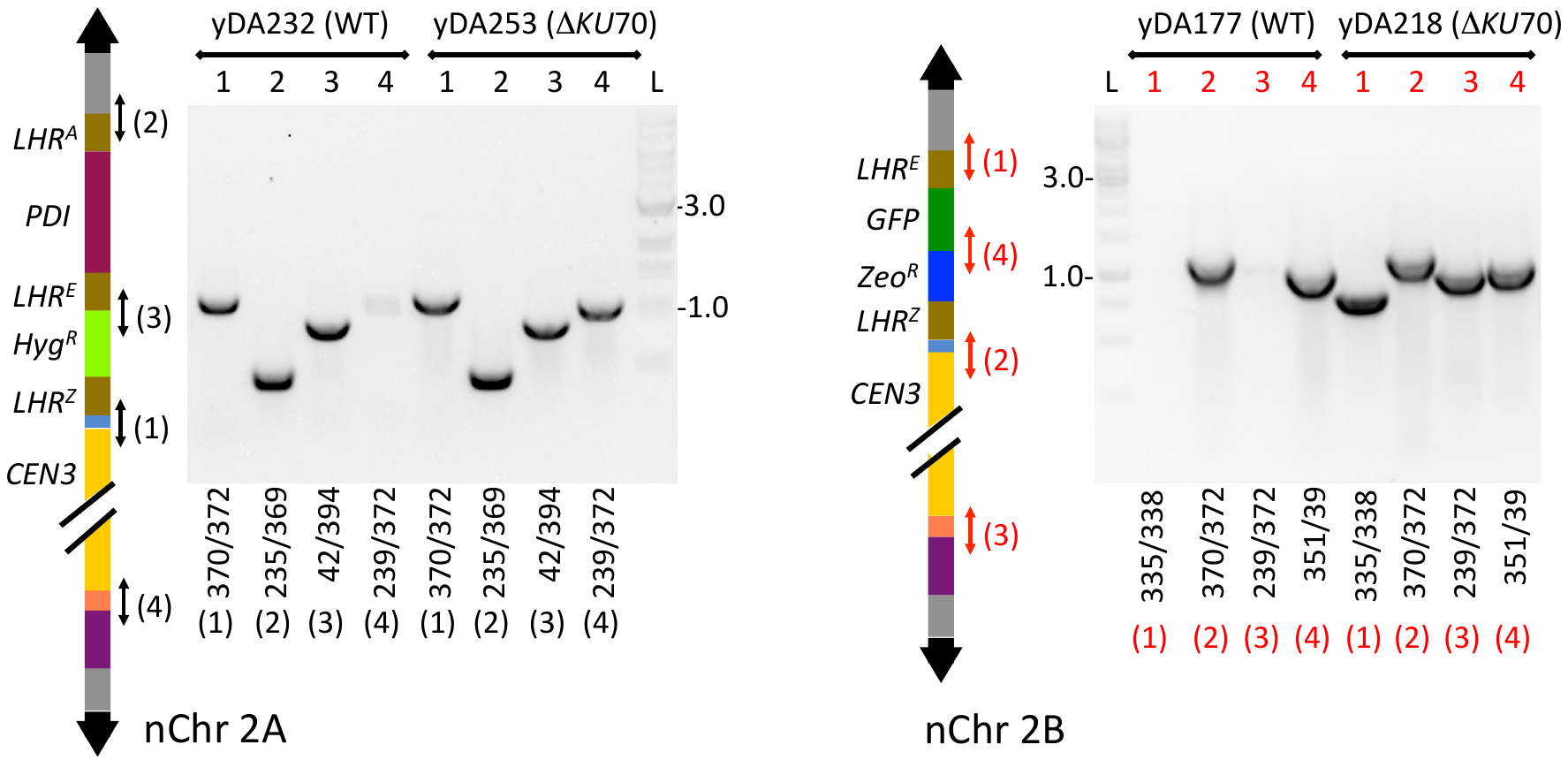

**Supplementary Figure S11**

*Comparison of PCR-based genotyping of nChr 2A and nChr 2B on wild-type and ΔKU70 K. phaffii backgrounds*

Missing amplicons in the cases of yDA232 (oligo pair 4) and yDA177 (oligo pairs 1 and 3) are consistent with the instability of both these extended nanochromosomes on wild-type backgrounds. Conversely, candidate bands are evident for all four targets in both the *ΔKU70* strains*, i.e.* yDA253 and yDA218. Analysis was performed on strains inoculated and cultivated on YPD medium, without antibiotic, for 24 hours. These data are in good agreement with WGS.

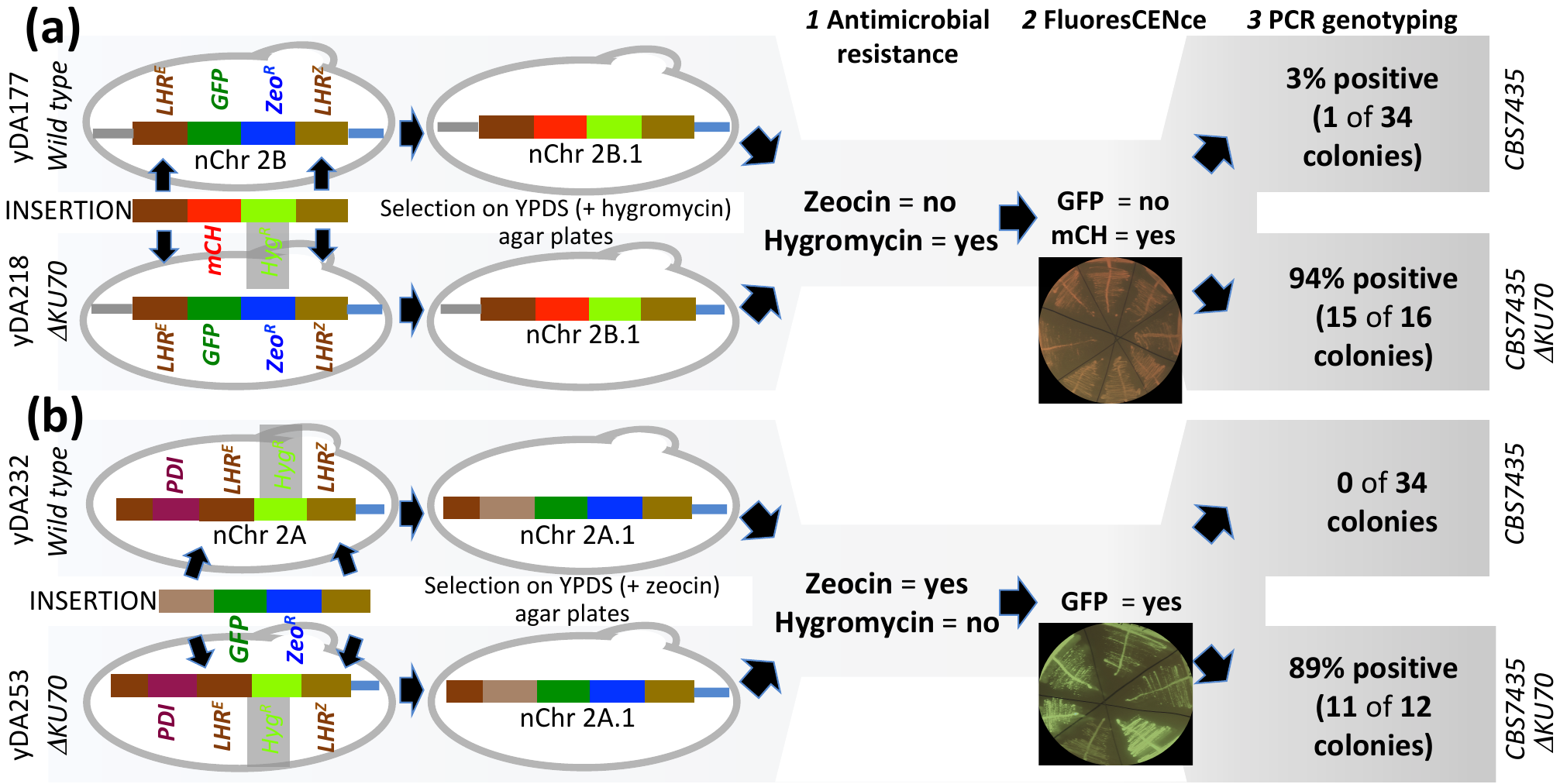

**Supplementary Figure S12**

*Engineering a nanochromosome in the context of wild-type (CBS7435) or ΔKU70 strains of K. phaffii* (See also Figure 7)

These proof-of-principle experiments were designed to assessed the feasibility of replacing genes within the landing zones of nanochromosomes with different ones, *in vivo*, by double cross-over HR. ***(a)*** Attempted replacement of *GFP*-*Zeo^R^* in the landing zone of nChr 2B with *mCH*-*Hyg^R^*. ***(b)*** Attempted replacement of *Hyg^R^* in the landing zone of nChr 2A with *GFP* and *Zeo^R^*. Note the higher success rate obtained with *ΔKU70* strains in terms of selected colonies.

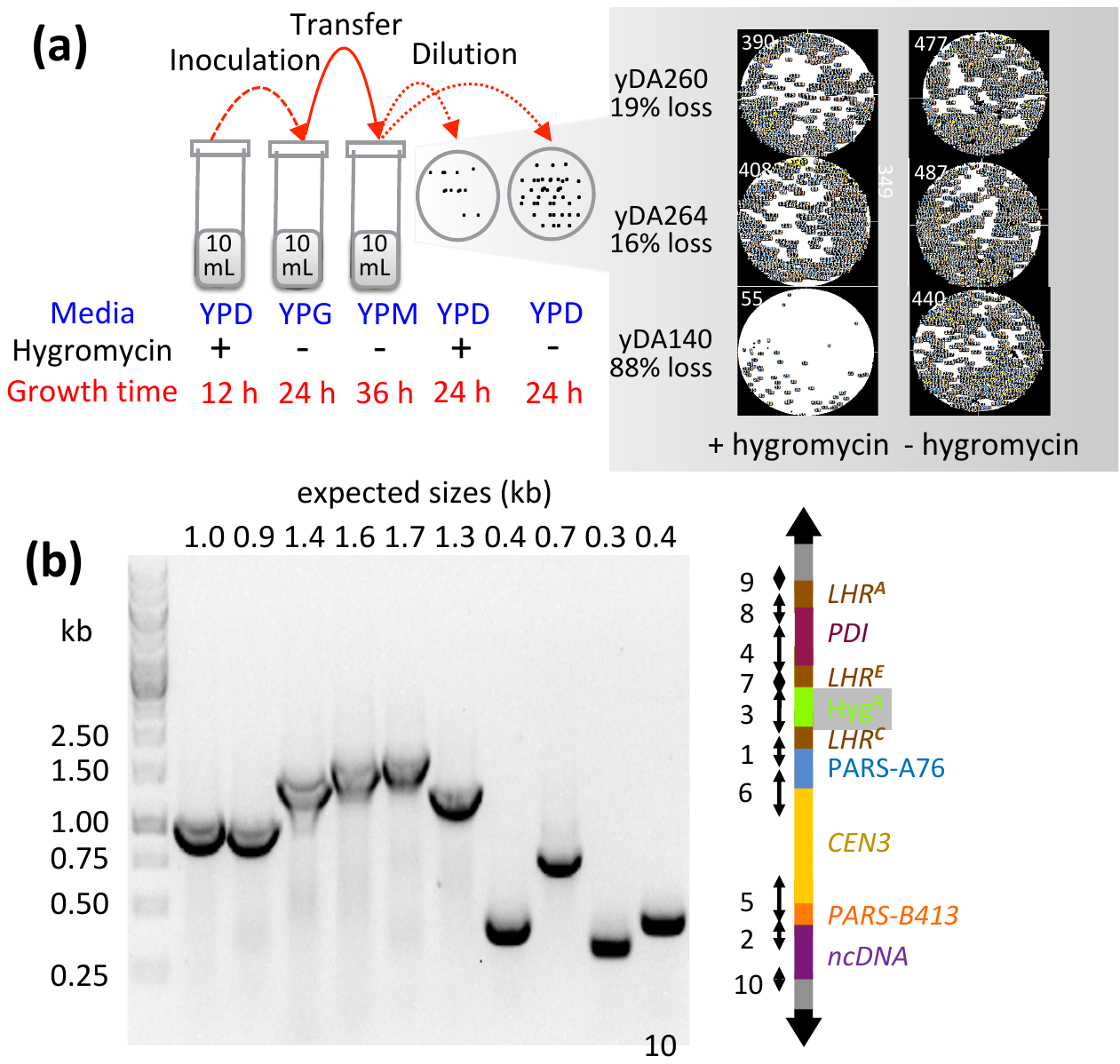

**Supplementary Figure S13**

*Assessment of nChr 2A persistence in K. phaffii strain yDA260*

***A*** Cells grown for 12 hours in YPD (+ hygromycin) were used to inoculate YPG (no antibiotic). After 24 hours of exponential growth in YPG, cells were spun down then re-suspended in YPM (no antibiotic). After 36 hours of methanol-induction, calls were streaked out on YPD plates, with or without hygromycin. An assessment of the retention of *Hyg^R^* (and, by extension, nChr 2A) was made by comparing colony counts with *vs*. without hygromycin. The results (right-hand panels, white numerals are colony counts derived from ImageJ software) were expressed as % of colonies that were no longer-hygromycin resistant (note: yDA260 carries nChr 2A; yDA264 is a control in which the *Hyg^R^* cassette is integrated into the native genome; yDA140, contains only (telomere-null/centromere-null) eDA37 (see Additional File 2: Suppl. Table 5). ***B*** Representative genotyping by colony PCR *of P. pastoris* strain yDA260 recovered from YPD plate after the chromosome-loss assay. The (numbered) sites targeted by oligo-pairs (Additional File 2: Suppl. Table 2) are indicated on on the schematic of nChr 2A.

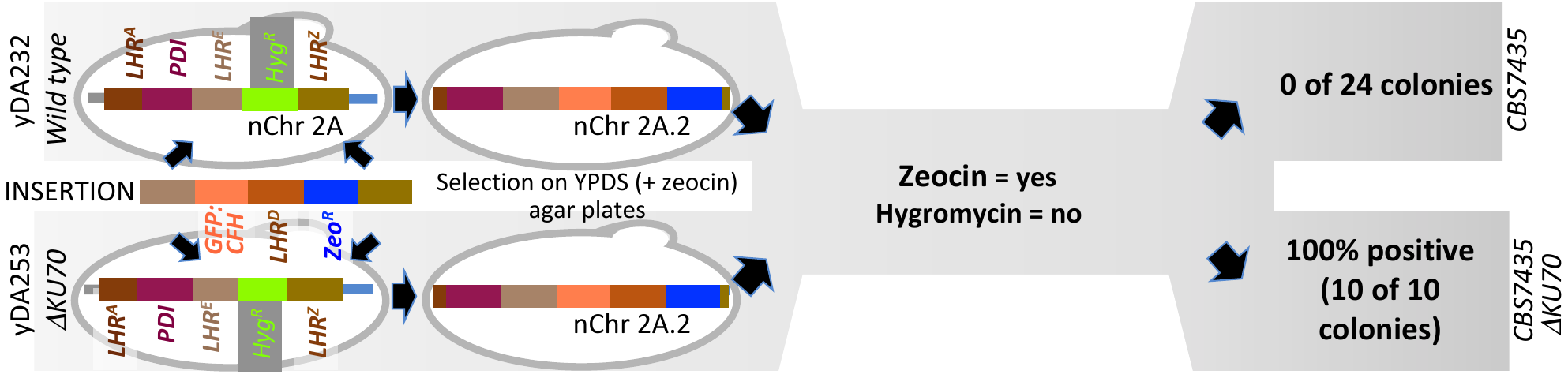

**Supplementary Figure S14**

*Illustration of the use of an “inch-worming” strategy for in vivo gene integration into the nanochromosome that requires a DKU70 background*.

The wild-type and *ΔKU70* strains shown were transformed with an integration array, schematized in the figure, which was designed to replace HygR in nChr 2A with *GFP:FH-LHR^D^-Zeo^R^*. Single colonies that grew on zeocin-containing agar plates were screened for zeocin resistance and hygromycin sensitivity following by PCR genotyping (see Additional file: Suppl. Fig. 14). Only *KU70*-compromised strains tested positive.

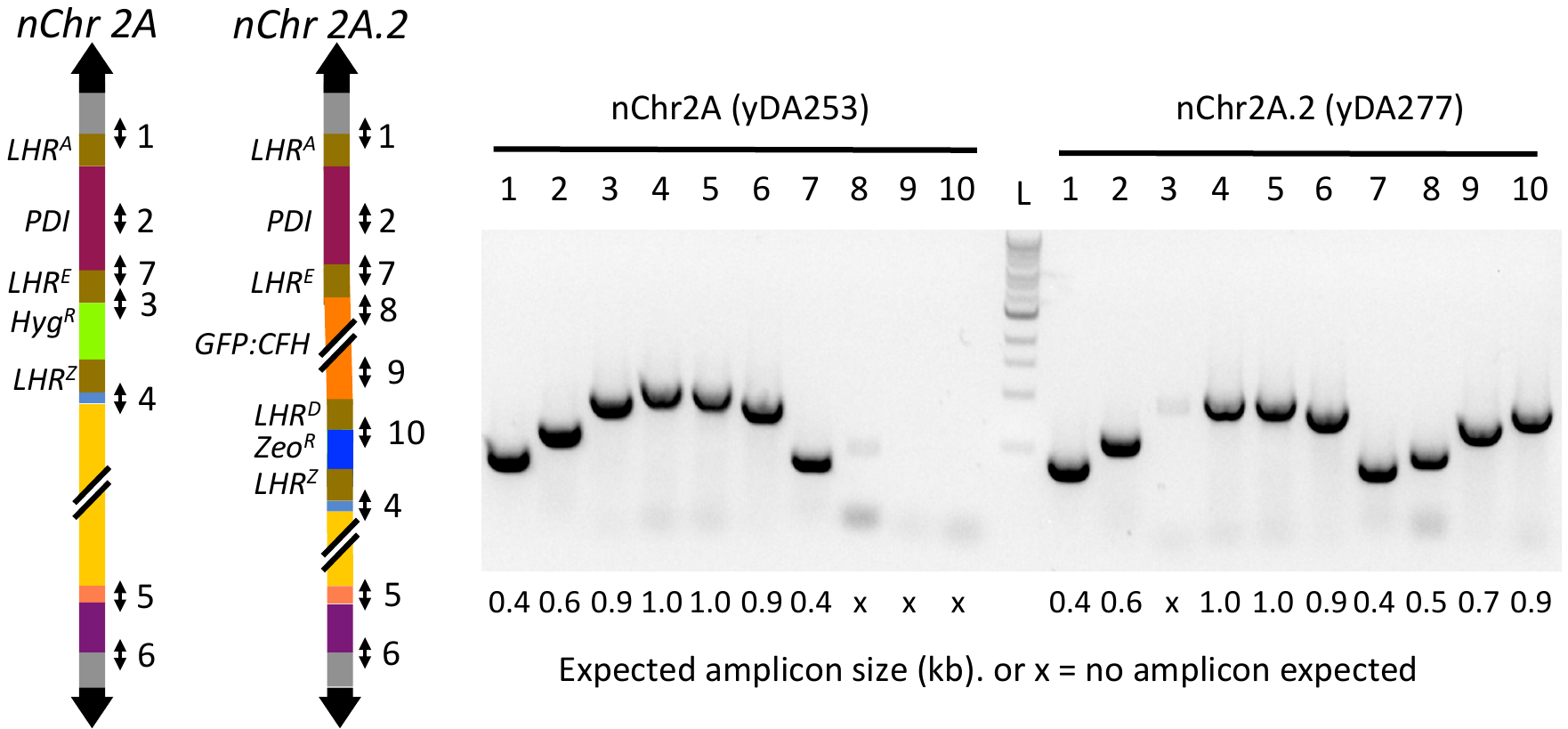

**Supplementary Figure S15**

*Validation of inch-worming*

Here, nChr 2A was engineered, *in vivo*, into nChr 2A.2, by gene integration accompanied by simultaneous extension of the landing zone (ready for a future integration) and by exchange of selection-markers. *ΔKU70* *K. phaffii* cells carrying nChr 2A were transformed with the insertion array *LHR^E^*-*GFP*:*FH*-*LHR^D^*-*Zeo^R^*-*LHR^Z^*. Transformants (growing on Zeocin) were transferred to YPD agar plates, then screened for zeocin resistance and hygromycin sensitivity. An attempt was made to validate nChr2A.2 by PCR-based genotype mapping. The amplicon numbers shown in the carton arise from the following ten oligo pairs (see Additional file: Supplementary Table 1): (1) 369/235; (2) 23/25; (3) 394/372; (4) 370/372; (5) 443/372; (6) 382/374; (7) (394/338); (8) 394/272; (9) 274/278; (10) 40/345.

Í
