## Supplementary material for "A supernumerary synthetic chromosome in *Komagataella phaffii* as a repository for extraneous genetic material": Abramczyk et al AdditionalFile2.docx

Dariusz Abramczyk^1^, María del Carmen Sánchez Olmos^2^, Adán Andrés Ramírez Rojas^2^, Daniel Schindler^2,3^, Daniel Robertson^4^, Stephen McColm^5^, Adele L. Marston^6^, Paul N. Barlow^1,4^

^1^ School of Chemistry, University of Edinburgh, United Kingdom

^2^ Max Planck Institute for Terrestrial Microbiology, Marburg, Germany

^3^ Center for Synthetic Microbiology, Philipps-Universität Marburg, Marburg, Germany

^4^ School of Biological Sciences, University of Edinburgh, United Kingdom

^5^ Ingenza Ltd Scotland, United Kingdom

^6^ The Wellcome Centre for Cell Biology, Institute of Cell Biology, School of Biological Sciences, University of Edinburgh, United Kingdom

**Additional file 2** (Tables)

**Contents**

***Table 1*** *List of oligonucleotides used in this study*

***Table 2*** *List of plasmids used in this study*

***Table 3*** *List of K. phaffii strains referred to in the current work*

***Table 4*** *List of insertion and integration arrays*

***Table 5*** *Results of (nano)chromosome-loss assays.*

***Table 6*** *Estimation of gene-copy number*

**Table 1** *List of oligonucleotides used in this study*

| **Oligo ID number**  **(F = forward;**  **R = reverse)** | **Purpose/description** | **Sequences (5’- 3’)**  Type-II restriction enzyme-recognition sites underlined; **type-IIS restriction enzyme-recognition sites in bold**; lower case letters denote overhangs generated by type-IIS restriction enzymes |
| --- | --- | --- |
| 23 (F) | Recombinant *PDI* | GCCAGAGGACGCTTCTAACTTGG |
| 25 (R) | Recombinant *PDI* | GCTGGAGTGGAGTCGGAAGTG |
| 42 (R) | For genotyping (*hph*) | GGCGCAGCTATTTACCCGC |
| 132a (R) | T*_CYC1_ (BsmBI)* | G**CGTCTC**CgtagGCCTTCGAGCGTCCC |
| 156 (F)(R) | Internal Repeats of *CEN3.* For genotyping (Fig. 4C, in set 6) | CAGGCTACCTCATATCTCCATCATC |
| 197 (F) | *CEN3* part (*Kpn*I) | GCGGTACCGTAGATCAAGTGACTTCTTGAGCTTGCC |
| 198 (R) | *CEN3* part (*EcoR*I)**;** an additional **G** is inserted into nanochromosomal *CEN3* cores (Fig. 4C, in set 6) | G**G**AATTCTAAATACCTTGACGAATAGTAAATATCGGATG |
| 199 (F) | As above | G**G**AATTCTTGTGCGAACTTGTGGGTCGTGG |
| 200 (R) | *CEN3* part (*Nde*I) | CCTTCCCCATATGTGCTCCGTCAGCTTGAATAAGCC |
| 212 (R) | Murine *FH* (for qPCR) | CCGTCGCCATTCTCGACTC |
| 215 (F) | Murine *FH* (for qPCR) | GCAGCCGAAACCGATCAGGAG |
| 217 (F) | *PARS-A76 (BamH*I*)* | GGGGATCCGGCGCTCCAACTGTATTCTAGG |
| 218 (R) | *PARS-A76 (BamH*I*)* | GGGGATCCCGAAGAGATGAGATGAGCTGATAAGATG |
| 223 (F) | *iSce*I-*ORF (EcoR*I*)* | GGAATTCATGGGATCAAGATCGCCAAAAAAGAAG |
| 224 (R) | *iSce*I*-ORF (Sac*II*)* | GCCGCGGTTATTTCAGGAAAGTTTCGGAGGAGATAGTG |
| 235 (F) | pUC region (Fig. 4C, in set 1) | AGCGAGTCAGTGAGCGAG |
| 238 (F) | *PARS-B413* (*BamH*I) | GCGGATCCCAACAATACTACGGAAACGACCCA |
| 239 (R) | *PARS-B413* (*BamH*I) (Fig. 4C, in set 5) | GGGGATCCGATAAGCTGGGGGAACATTCGC |
| 246 (F) | 5’-*Tel*, complementary strand to 247 (*in* *italic*s – *I-Sce*I-recognition site part) – includes *Pvu*I | CGTGGATGCTGGATGCTGGATGCTGGATGCTGGATGCTGGATGCTGGATGCTGGATGCTGGATGCTGGATGCTGGATGCTGGATGCTGGA­­TGCTGGATGCTGGATGCTGGATGC*TAGGGATAACAGG­* |
| 247 (R) | Complementary to strand 246  (*in* *italic*s – *I-Sce*I-recognition site part) – includes *Pvu*I | *CCTA*GCATCCAGCATCCAGCATCCAGCATCCAGCATCCAGCATCCAGCATCCAGCATCCAGCATCCAGCATCCAGCATCCAGCATCCAGCATCCAGCATCCAGCATCCAGCATCCACGAT |
| 248 (F) | 3’-*Tel*, complementary strand to 249 (*in italics* – *I-Sce*I-recognition site part) – includes *Pvu*I | *GTAAT*GCATCCAGCATCCAGCATCCAGCATCCAGCATCCAGCATCCAGCATCCAGCATCCAGCATCCAGCATCCAGCATCCAGCATCCAGCATCCAGCATCCAGCATCCAGCATCCACGAT |
| 249 (R) | 3’-*Te*l, complementary strand to 248 (*in* *italics* – *I-Sce*I-recognition site part) – includes *Pvu*I | CGTGGATGCTGGATGCTGGATGCTGGATGCTGGATGCTGGATGCTGGATGCTGGATGCTGGATGCTGGATGCTGGATGCTGGATGCTGGATGCTGGATGCTGGATGCTGGATGC*ATTACCCTGTTATC* |
| 253 (R) | 1.4-kb Gibson assembly fragment 1 for framework plasmid | GAAATTGTTATCCGCTCACAATTCGCGGCCTTTTTACGGTTCCT |
| 254 (F) | 1.4-kb Gibson assembly fragment 1 for framework plasmid | CCCGCGCGTTGGCCGATTCATTAATGCAGTTGTCATTTTGTTTCTTCTCGTACGAGCTTG |
| 255 (F) | 0.51-kb Gibson assembly fragment 2 for framework plasmid | AGGAACCGTAAAAAGGCCGCGAATTGTGAGCGGATAACAATTTCACACAG |
| 256 (R) | 0.51-kb Gibson assembly fragment 2 for framework plasmid | TGCTTGGCAAGCTCAAGAAGTCACTTGATCTACGGTACCCGGGATGTGCTGCAAGGCG |
| 264 (F) | *LHR^A^* (*Pvu*II) | AACAGCTGTGTTAAACCGTCTTTAAG |
| 267 (F) | *i-Sce*I cassette part (*BsmB*I) | C**CGTCTCA**gggaATCTAACATCCAAAGACGAAAGGTTG |
| 268 (R) | *i-Sce*I cassette part (*BsmB*I) | C**CGTCTCA**ttggCTCACTTAATCTTCTGTACTCTGAAGAGG |
| 269 (F) | *LHR^Z^* part (*BsmB*I) | C**CGTCTCCccaa**CCGTGTAGGTCTACAAACTGG |
| 270 (R) | *LHR^Z^* part (*BsmB*I) | C**CGTCTCAcgcg**CCTGCAGGTTGAGTTGGCGAAGGTGCG |
| 272 (R) | *P_AOX1_* (for genotyping) | CCTCCCCTATGTGATCGTCG |
| 273 (R) | *LHR^A^* (*Pvu*II) | AACAGCTGCAAACTAAGATTGGACTATTGCTATCC |
| 274 (F) | *GFP* (for genotyping) | AAGTTGCCCGTCCCCTG |
| 276 (F) | *GFP*-linker-fusion (*Nsi*I) | CCCATGCATCCATGGTATCTAAAGGTGAAGAACTATTCACAG |
| 277 (R) | *GFP*-linker-fusion (*Nsi*I) | CCCATGCATGACGACCTTCAATTTTGTAAAGCTCATCCATTCCCAAG |
| 278 (R) | *Human FH* (for genotyping) | CCCACTCACCCTTTCTACAAACC |
| 280 (R) | *Amp^R^* (Fig. 4C, in set 2) | TTATCCGCCTCCATCCAG |
| 282 (R) | *M13* (genotyping) | CTGGCCGTCGTTTTACAACG |
| 290 (F) | *Amp^R^* ( Fig. 4C, in set 2) | CTATGTGGCGCGGTATTATC |
| 300 (F) | *LHR^A^ (BsmB*I*)* | C**CGTCTCC**gggaCTGTGTTAAACCGTCTTTAAGTCAAC |
| 301 (R) | *LHR^A^ (BsmB*I*)* | G**CGTCTCC**TgtccAAACTAAGATTGGACTATTGCTATC |
| 302 (F) | *P_AOX1_* (*BsmB*I) | C**CGTCTC**CgacaAACATCCAAAGACGAAAGGTTG |
| 302a (F) | *P_TEF1_* (for *GFP* and *mCH* cassettes) (*BsmB*I) | C**CGTCTC**CgacaGTTTCTACTCCTTTTTTACTCTTCC |
| 303 (R) | *T_AOX1_* (*BsmB*I) | G**CGTCTC**CctgaTCTCACTTAATCTTCTGTACTCTG |
| 303a (R) | *T_CYC1_* (for *GFP* and *mCH* cassettes) (*BsmB*I) | G**CGTCTC**C**ctga**GCCTTCGAGCGTCCC |
| 304 (F) | *LHR^D^ (BsmB*I) | C**CGTCTC**CtcagGACAGCAACCTAACCGAC |
| 305 (R) | *LHR^D^ (BsmB*I) | G**CGTCTC**CctacCAGTCCTCGTGAAAGACGAG |
| 306 (F) | *Hyg^R^* (*Sal*I-*Bsa*I) | CCTGTCGAC**GGTCTC**AgtagGACATGGAGGCCCAGAATACCC |
| 307 (R) | *Hyg^R^* (*BamH*I) | GCGGATCCCAGTATAGCGACCAGCATTCACATAC |
| 308 (F) | *Zeo^R^* cassette (*Sal*I-*BsmB*I) overhang GTAG | CTGTCGAC**CGTCTC**AgtagCCCACACACCATAGCTTCAAAATG |
| 309 (R) | *Zeo^R^* cassette (*BamH*I) | GGGATCCGCAAATTAAAGCCTTCGAGCG |
| 315 (F) | *LHR^A^ (BsmB*I*-AsiS*I*)* | CCGTCTCCgggaGCGATCGCCTGTGTTAAACCGTCTTTAAGTCAACCC |
| 317 (R) | *LHR^Z^* (*Kpn*I-*Bsa*I-*AsiS*I) | GGGGTACC**GGTCTC**AcgcgGCGATCGCCCTGCAGGTTGAGTTGGCGAAGGTGCG |
| 318 (R) | *LHR^Z^* (*Kpn*I-*BsmBI*-*AsiS*I) | GCGGTACC**CGTCTC**A**cgcg**GCGATCGCCCTGCAGGTTGAGTTGGCGAAGGTGCG |
| 321 (F) | Bacterial *Kan^R^* cassette (*Sph*I) | CGCGCATGCTGATCGGCACGTAAGAGG |
| 322 (R) | Bacterial *Kan^R^* cassette (*Sph*I) | GCGGCATGCCGCAGAAAGGCCCACCCG |
| 335 (F) | *LHR^E^* (*Bsa*I-*AsiS*I) | C**GGTCTC**CgggaGCGATCGCGACTTACCTCGTTTTAACTTAGTCGG |
| 336 (R) | *LHR^E^* (*Bsa*I) | C**GGTCTC**C**Ttcag**GACTTACCTCGTTTTAACTTAGTCGG |
| 338 (R) | *LHR^E^* (*Bsa*I) | G**GGTCTCCtgtc**ACAAGTTTAACTAAGCGTCAGCC |
| 345 (F) | *LHR^D^* (genotyping) | CGGCAATCCTGTACTCTGC |
| 351 (R) | *LHR^Z^* verification ( Fig. 4C, in set 3) | GCTTCTGCTCTTAAACTAGCTGC |
| 355 (F) | *T_AOX1_* ( Fig. 4C, in set 3) | TCGTACGAGCTTGCTCCTG |
| 369 (R) | *LHR^A^* | CTGTTGAGGCACAAACCGTCAG |
| 370 (F) | *LHR^C^* (qPCR, genotyping) ( Fig. 4C, in set 4) | CGTTACACGGCAGTCCTGTTCC |
| 372 (F)(R) | Internal repeats /CEN*3* (genotyping) ( Fig. 4C, in sets 4 and 5) | CCGTTGGACCAATGTGGAATC |
| 374 (R) | 3’-p arm (genotyping) | ATAATACCGCGCCACATAGC |
| 375 (R) | 3’-p arm of Chr 3 (genotyping) | CATGTGTATTGGAGAGCAGAG |
| 379 (R) | 1.3-kb ncDNA (*Nde*I) | CCTTCCCCATATGCTAGTGCAGAGTGAGCCTG |
| 380 (F) | 1.3-kb ncDNA (*EcoR*I) | GGAATTCGCTAGCCAGATCGGCTTAAG |
| 381 (F) | 1.9-kb *ncDNA*-*PARS-B413* (*Nde*I) | CCTTCCCCATATGCAACAATACTACGGAAACGACCC |
| 382 (F) | 1.3-kb *ncDNA* verification ( Fig. 4C, in set 7) | GATTCGCCCTGCAGATTC |
| 384 (F) | Amplification of insertion array (*Afe*I), blunt end (*LHR^A^*-) | CCCAGCGCTGTGTTAAACCGTCTTTAAG |
| 394 (F) | *LHR^E^* verification | GCCAACCAAAGGTGGAGG |
| 397 (F) | Amplification of insertion array (*Afe*I) blunt end (*LHR^E^*-) | TTATAGCGCTGACTTACCTCGTTTTAACTTAGTCGG |
| 409 (R) | Amplification of insertion array (*LHR^Z^*-) (*XhoI*) (*SalI*I compatible end) | TTATCTCGAGTGAGTTGGCGAAGGTGCG |
| 417 (R) | *LHR^Z^* verification | TTGAGTTGGCGAAGGTGCG |
| 443 (R) | Verification of 1.9-kb *ncDNA*-*PARS-B413* | AAAGTCAACTGTCCTTTCTGGC |
| 448 (R) | *PDI* part (*Not*I) introducing HIS-tag-TAA stop codon (*italics*) | TTTTTGCGGCCGC*TTAATGATGATGGTGGTGATG*CAACTCATCGTGAGCATCAGCTTC |

**Table 2** *List of plasmids used in this study*

| **Plasmid ID** | **Size (Kb)** | **Key contents (notes)** | **Purpose** | **Source** |
| --- | --- | --- | --- | --- |
| pUC19 | 2.68 |  | Cloning | Thermo Fisher Scientific |
| pPICZα B | 3.6 |  | Cloning and expression | Thermo Fisher Scientific |
| pPICZ A | 3.3 |  | Cloning and expression | Thermo Fisher Scientific |
| pGS-GnT-II | 5.8 |  | Cloning | Glycoswitch |
| pGS-GnT-I | 4.2 |  | Cloning | Glycoswitch |
| eDA8 | 10.5 | Contains 8 kb of DNA designed to be non-coding, have no restriction sites *etc.* | Source of *LHRs^A-D^*, and *ncDNA* extension in p-arm of nanochromosome | In house (V. Zulkower & A. Marston, unpublished) |
| eDA9 | 10.5 | Contains 8 kb of DNA designed to be non-coding, have no restriction sites *etc.* | Source of *LHR^E^*, *LHR^D^*, *LHR^Z^* | In house (V. Zulkower & A. Marston, unpublished) |
| eDA10 | 5.92 | Chr 3 centromere (*CEN3)* (5’-part) in pUC19 | En route to eDA24 | This work |
| eDA22 | 6.98 | pRS413*-(P_GAL_)I-SceI* | Source of meganuclease gene | (Shen, Wang et al. 2017) |
| eDA24 | 8.85 | *CEN3* in pUC19 | DNA-parts assembly (centromere) | This work |
| eDA26 | 3.0 | *pARS-A76* in pUC19 | DNA-parts assembly (ARS) | This work |
| eDA27 | 4.12 | pPICZ A-(*P_AOX1_)I-SceI(T_AOX1_)* | Meganuclease gene cassette, intended for *in vivo* linearisation | This work |
| eDA37 | 4.2 | *PARS-A76-Hyg^R^* in pUC19 | Source of *Hyg^R^* and negative control for chromosome-loss assay | This work |
| eDA40 | 4.2 | *PARS-A76-Zeo^R^* in pUC19 | Source of *Zeo^R^* and negative control for chromosome-loss assay | This work |
| eDA41 | 3.2 | *PARS-B413* in pUC19 | DNA-parts preparation (ARS) | This work |
| eDA53 (framework plasmid) | 10.5 | pUC-*PARS-A76*-*CEN3*-*Zeo^R^* | Framework plasmid (see Fig. 1A) | This work |
| eDA71 | 11.3 | pUC-*PARS-A76-CEN3-LHR^A^* | Intermediate plasmid en route to eDA83 | This work |
| eDA83 | 14.5 | pUC-*PARS-A76/CEN3-LHR^A^- Zeo^R^-(P_AOX1_)I-SceI-LHR^Z^* | Abandoned strategy for self-cleaving precursor plasmid; intermediate en route to eDA110 | This work |
| eDA89 | 7.9 | pPICZα B-*GFP:FH* | DNA-parts prep. (GFP-FH fusion) | unpublished |
| eDA99 | 3.66 | 0.95-kb *LHR^Z^* in pUC19 (*Bsa*I version) | DNA-parts prep. | This work |
| eDA101 | 3.66 | 0.95-kb *LHR^Z^* in pUC19 (*BsmB*I version) | DNA-parts prep. | This work |
| eDA103 | 5.24 | *Hyg^R^*-*LHR^Z^* part in pUC19 (*Bsa*I version) (with *AsiS*I in *Hyg^R^*) | DNA-parts prep. | This work |
| eDA105 | 4.83 | *Zeo^R^-LHR^Z^* part in pUC19 (*BsmB*I version) | DNA-parts prep. | This work |
| eDA110 | 14.5 | pUC*-PARS-A76-Cen3-LHR^A^- Zeo^R^-Kan^R^(ΔI-SceI)-LHR^Z^* | Intermediate plasmid en route to precursor plasmid *v*1 | This work |
| eDA115 | 5.24 | *Hyg^R^*-*LHR^Z^* in pUC19 (*Bsa*I version) (no *AsiS*I) | DNA-parts prep. | This work |
| eDA131 | 3.55 | *telomere-I-SceI-telomere* in pUC19 | DNA-parts prep. (Proto-telomeres*, Tel*) | This work |
| eDA137 | 14.9 | pUC-*PARS-A76-CEN3-LHR^A^- Zeo^R^-Kan^R^::(P_AOX1_)I-SceI-LHR^Z^ AmpR::telomere-I-SceI-telomere* | Precursor plasmid *v*1 | This work |
| eDA143 | 10.5 | *PDI* in pPIC3.5K | Source of *PDI* | (Kerr, Herbert et al. 2021) |
| eDA144 | 4.6 | 1.9-kb *PARS-B413-ncDNA* part in pUC19 | DNA-parts prep. (p-arm extension) | This work |
| eDA146 | 16.8 | pUC-*PARS-A76-CEN3-PARS-B413-ncDNA-LHR^A^- Zeo^R^-Kan^R^(ΔI-SceI)-LHR^Z^* | Intermediate plasmid, en route to eDA155 | This work |
| eDA155 | 17.3 | pUC-*PARS-A76-CEN3-PARS-B413-ncDNA-LHR^A^- Zeo^R^- Kan^R^::(P_AOX1_)I-SceI-LHR^Z^-AmpR::telomere-I-SceI-telomere* | Precursor plasmid *v*2 | This work |
| eDA189 | 4.5 | pUC-(*PARS-A76)-(P_TEF1_)GFP(T_CYC1_)* | DNA-parts prep. (*GFP* cassette) | This work |
| _eDA191_ | 4.5 | pUC-(*PARS-A76)-(P_TEF1_)mCH(T_CYC1_)* | DNA-parts prep. (*mCH* cassette) | This work |
| eDA197 | 7.14 | pUC-*LHR^E^-(P_TEF1_)GFP(T_CYC1_)-HygR-LHR^Z^* | Assembly/repository of integration array | This work |
| eDA199 | 7.5 | pUC-*LHR^E^-(P_TEF1_)mCH(T_CYC1_)-ZeoR-LHR^Z^* | Assembly/repository of insertion array | This work |
| eDA201 | 15.8 | pUC-*PARS-A76-CEN3-PARS-B413-ncDNA-LHR^E^-GFP-HygR-LHR^Z^-AmpR::telomere-I-SceI-telomere* | Precursor plasmid *v*2b | This work |
| eDA226 | 4.8 | pPICZ A-(*P_AOX1_)PDI_H_(T_AOX1_)* | DNA-parts prep. (His-tagged *PDI*) | This work |
| eDA227 | 10.0 | pUC-*LHR^A^-(P_AOX1_)PDI(T_AOX1_)-LHR^E^-HygR-LHR^Z^* | Assembly/repository of insertion array | This work |
| eDA229 | 18.7 | pUC-*PARS-A76-CEN3-PARS-B413-ncDNA-LHR^A^-(P_AOX1_)PDI(T_AOX1_)-LHR^E^-HygR-LHR^Z^ -AmpR::telomere-I-SceI-telomere* | Precursor plasmid *v*2a | This work |
| eDA250 | 9.0 | pUC-*LHR^E^-(P_TEF1_)GFP(T_CYC1_)-LHR^D^-ZeoR-LHR^Z^* | Assembly/repository of integration array | This work |

**Table 3** *List of K. phaffii strains referred to in the current work*

| ***K. phaffii* strain** | **Genotype** | **Description (notes)** | **Landing Zone** | **Source** |
| --- | --- | --- | --- | --- |
| Background | CBS7435 (note: *Mut^S^*) | Wild type (WT) | n/a | TFS |
| yDA34 | CBS7435;eDA53 | WT, transformed with framework plasmid | n/a | This work |
| yDA39 | CBS743;eDA40 | WT, transformed with episomal *Zeo^R^*-carrying plasmid, as negative control for chromosome-loss assays | n/a | This work |
| yDA122 | CBS743;linear eDA137 | WT, transformed with linearised precursor plasmid *v*1 – [isolate no.1] | *LHR^A^*- *Zeo^R^-Kan^R^*(Δ*I-SceI*)-*LHR^Z^* | This work |
| yDA140 | CBS7435;eDA37 | WT, transformed with episomal, *Hyg^R^*-containing plasmid as negative control for chromosome-loss assays | n/a | This work |
| yDA149 | CBS7435;linear eDA137 | WT, transformed with linearised precursor plasmid *v*1 – [isolate no.2] | *LHR^A^*- *Zeo^R^-Kan^R^*(Δ*I-SceI*)-*LHR^Z^* | This work |
| yDA174 | CBS7435;linear eDA155 | WT, transformed with linearised precursor plasmid *v*2 – [isolate no.1] | *LHR^A^*- *Zeo^R^-Kan^R^*(Δ*I-SceI*)-*LHR^Z^* | This work |
| yDA175 | CBS7435;linear eDA155 | WT, transformed with linearised precursor plasmid *v*2 – [isolate no.2] | *LHR^A^*- *Zeo^R^-Kan^R^*(Δ*I-SceI*)-*LHR^Z^* | This work |
| yDA177 | CBS7435;linear eDA201 | WT, transformed with precursor plasmid *v*2B – [isolate no.2] | *LHR^E^-P_TEF_GFP-Zeo^R^-LHR^Z^* | This work |
| yDA208 | CBS7435 Δ*KU70* | WT with KU70-encoding gene knocked out (KO) | n/a | (Dalvie, Leal et al. 2020) |
| yDA218 | CBS7435 Δ*KU70;*linear eDA201 (nChr 2B) | *KU70*-KO, transformed with linearised precursor *v*2B – [isolate no.1] | *LHR^E^-P_TEF_GFP-Zeo^R^-LHR^Z^* | This work |
| yDA221 | CBS7435 P*_AOX1_::mFH(Zeo*^R^) | WT with genome-integrated murine mFH-expression cassette | n/a | This work |
| yDA226 | CBS7435 Δ*KU70;*nChr 2B.1 | *KU70*-KO, with nChr 2B.1 (after double-crossover HR performed on the landing zone of nChr 2B in yDA218) | *LHR^E^-(P_TEF_)mCH-Hyg^R^-LHR^Z^* | This work |
| yDA232 | CBS7435;linear eDA229 (nChr 2A) | WT, transformed with linearised precursor plasmid *v*2A | *LHR^A^-(P_AOX1_)PDI_H_(T_AOX1_)-LHR^E^-Hyg^R^-LHR^Z^* | This work |
| yDA245 | CBS7435 P*_AOX1_::*P*_AOX_ mFH(Zeo*^R^)*;*linear eDA229 (nChr 2A) | WT, with genome-integrated *mFH*-expression cassette*,* transformed with linearised precursor plasmid *v*2A | *LHR^A^-(P_AOX1_)PDI_H_(T_AOX1_)-LHR^E^-Hyg^R^-LHR^Z^* | This work |
| yDA250 | CBS7435 Δ*KU70* P*_AOX1_::*P*_AOX1_mFH(Zeo*^R^) | *KU70*-KO, with genome-integrated *mFH*-expression cassette | n/a | This work |
| yDA253 | CBS7435 Δ*KU70;*linear eDA229 (nChr 2A) | *KU70*-KO, transformed with linearised precursor plasmid *v*2A | *LHR^A^-(P_AOX1_)PDI_H_(T_AOX1_)-LHR^E^-Hyg^R^-LHR^Z^* | This work |
| yDA260 | CBS7435 Δ*KU70* P*_AOX1_::mFH(Zeo*^R^*);*linear eDA229 (nChr 2A) | *KU70*-KO, with genome-integrated *mFH*-expression cassette*,* transformed with linearised precursor plasmid *v*2A | *LHR^A^-(P_AOX1_)PDI_H_(T_AOX1_)-LHR^E^-Hyg^R^-LHR^Z^* | This work |
| yDA263 | CBS7435 Δ*KU70* with nChr 2A.1 | *KU70*-KO, with nChr 2A.1 ( after double-crossover HR attempted on the landing zone of nChr 2A in yDA253) | *LHR^A^-(P_AOX1_)PDI_H_(T_AOX1_)-LHR^E^-(P_TEF_)GFP(T_TEF_)-Zeo^R^-LHR^Z^* | This work |
| yDA264 | CBS7435 Δ*KU70* P*_AOX1_::mFH (Zeo*^R^) P*_AOX1_::PDIHIS (Hyg*^R^) | *KU70*-KO with genome-integrated *mCFH* and *PDI_H_* expression cassettes | n/a | This work |
| yDA275 | CBS7435 Δ*KU70* nChr 2A.2 | *KU70*-KO with nChr 2A.2 [clone 1] (after double-crossover HR attempted on the landing zone of nChr 2A in yDA253) | *LHR^A^-(P_AOX1_)PDI_H_(T_AOX1_)-LHR^E^-(P_AOX1_)GFP:FH(T_AOX1_)-LHR^D^-Zeo^R^-LHR^Z^* | This work |
| yDA277 | CBS7435 Δ*KU70* nChr 2A.2 | *KU70*-KO with nChr 2A.2 [clone 2] (after double-crossover HR attempted on the landing zone of nChr 2A in yDA253) | *LHR^A^-(P_AOX1_)PDI_H_(T_AOX1_)-LHR^E^-(P_AOX1_)GFP:CFH-LHR^D^-Zeo^R^* | This work |

**Table 4** *List of insertion and integration arrays*

| **Array** | **Int/**  **Ins*** | **Assembled from parts (size in kb)** | **Prepared from (oligos used for PCR)** | **First ligation product (size in kb)** | **Array size (kb)** |
| --- | --- | --- | --- | --- | --- |
| *LHR^A^*-(*Kan*^R^)-  *I-SceI*-*Zeo^R^*-*LHR^Z^* | Int | *LHR^A^* (0.85) | eDA8 (264/273) | n/a | ~6.2 |
|  |  | *Kan*^R^ (0.98) | pGS-Man-II (GlycoSwitch) |  |  |
|  |  | *I-SceI* (2.2) | eDA27 (267/268) |  |  |
|  |  | *Zeo^R^-LHR^Z^* (2.12) | pGS-GnT-I (GlycoSwitch) |  |  |
| *LHR^E^-GFP-Zeo^R^-LHR^Z^* | Int | *LHR^E^* (0.87) | eDA9 (335/336) | *LHR^E^-*  *(P_TEF1_)GFP*  *(T_CYC1_)* (2.31) | ~4.4 |
|  |  | *(P_TEF1_)GFP(T_CYC1_)* (1.44) | eDA189 (302a/132a) |  |  |
|  |  | *Zeo^R^-LHR^Z^* (2.12) | eDA105 (308/318) |  |  |
| *LHR^E^-mCH-Hyg^R^-LHR^Z^* | Int/  Ins | *LHR^E^* (0.87) | eDA9 (335/336) | *LHR^E^-*(*P_TEF1_*)*mCH*  *(T_CYC1_)* (2.28) | ~4.8 |
|  |  | *(P_TEF1_)mCH(T_CYC1_)* (1.41) | eDA191 (302a/132a) |  |  |
|  |  | *Hyg^R^-LHR^Z^* (2.54) | eDA115 (306/317) |  |  |
| *LHR^A^-PDI^H^-LHR^E^-Hyg^R^-LHR^Z^* | Int | *LHR^A^* (0.85) | eDA8 (300/301) | *LHR^A^*-*(P_AOX1_)PDI_HIS_*  *(T_AOX1_)* (3.7) | ~7.11 |
|  |  | *(P_AOX1_)PDI_HIS_*  *(T_AOX1_)*  (2.85) | eDA227 (302/303) |  |  |
|  |  | *LHR^E^* (0.87) | eDA9 (335/336) | *LHR^E^*-*Hyg^R^-LHR^Z^* (3.41) |  |
|  |  | *Hyg^R^-LHR^Z^* (2.54) | eDA115 (306/317) |  |  |
| *LHR^E^-GFP:FH- LHR^D^ -Zeo^R^-LHR^Z^* | Ins | *LHR^E^* (0.9) | eDA9 (335/338) | *LHR^E^*-*(P_AOX1_)GFP:FH*  *(T_AOX1_)* (6.8) | ~10.1 |
|  |  | *(P_AOX1_)GFP:FH(T_AOX1_)* (5.9) | eDA89 (302/303) |  |  |
|  |  | *LHR^D^* (1.1) | eDA9 (304/305) | *LHR^D^-Zeo^R^-LHR^Z^* (3.2) |  |
|  |  | *Zeo^R^-LHR^Z^* (2.1) | eDA105 (308/318) |  |  |

**Table 5** *Results of nanochromosome-loss assays performed for various strains*

| **Strain ID** | Background strain (host) | Transformation with linearised versions of (and hence the **nanochromosome** that is potentially introduced) | % of Zeo- or Hyg-resistant isolates lost (red font) after growth in non-selective media* |
| --- | --- | --- | --- |
| yDA122 | GS115 | eDA137 (**nChr 1**) | 10% loss 90% retain |
| yDA140 | CBS7435 | eDA37 (episomal plasmid) | 88% loss 12% retain |
| yDA174 | CBS7435 | eDA155 (**nChr 2**) | 17% loss 83% retain |
| yDA175 | CBS7435 | eDA155 (**nChr 2**) | 16% loss 84% retain |
| yDA177 | CBS7435 | eDA201 (**nChr 2B**) | 20% loss 80% retain |
| yDA218 | CBS7435 *ΔKU70* | eDA201 (**nChr 2B**) | 12% loss 88% retain |
| yDA226 | CBS7435 *ΔKU70* | eDA201 (**nChr 2B.1**) | 13% loss 87% retain |
| yDA232 | CBS7435 | eDA229 (**nChr 2A** ) | 17% loss 83% retain |
| yDA250 | CBS7435 *ΔKU70* + genome-integrated mFH | none | 98% loss 2% retain |
| yDA253 | CBS7435 *ΔKU70* | eDA229 (**nChr 2A**) | 19% loss 81% retain |
| yDA260 | CBS7435 Δ*KU70* + genome-integrated mFH | eDA229 (**nChr 2A**) | 19% loss 81% retain |
| yDA263 | CBS7435 *ΔKU70* transformed w. linearised precursor plasmid *v*2A | eDA229 *i.e.* nChr 2 subsequently engineered to create **nChr 2A.1** | 12% loss 88% retain |
| yDA264 | CBS7435 *ΔKU70* with genome-integrated *Hyg^R^*) | none | 16% loss 84% retain |
| yDA275 | CBS7435 *ΔKU70 transformed w. linearised precursor plasmid* v*2A*) | eDA229 *i.e.* nChr 2 subsequently engineered to create **nChr2 A.2** | 12% loss 88% retain |
| yDA277 | (CBS7435 *ΔKU70 transformed with linearised precursor* v*2A*) | eDA229 *i.e.* nChr 2 subsequently engineered to create **nChr2 A.2** | 11% loss 89% retain |

* Transformed cells were cultured for at least ten generations in the absence of any selection, then an assay based on counting colonies on agar plates was performed to assess retention of antibiotic-resistance. The values listed here were calculated from the total number of colonies counted growing on antibiotic-resistance selection medium, divided by the number of colonies counted growing on antibiotic-resistance *non-*selection medium. At least three biological replicates were performed for each strain. Standard deviations are illustrated by bars.

**Table 6 A** *Validation of primers sets for qPCR assays*. Slope and efficiency (%) values calculated from the sample dilution using StepOnePlus software (Thermo Fisher)

| **Gene** | **Target locus**  **(amplicon length, bp)** | **Oligonucleotide primer pair** | **Slope** | **Efficiency**  **range (%)** |
| --- | --- | --- | --- | --- |
| ***ACT1*** | Gene reference  (137 bp) | 35(F): TTCGTCGGTGACGAGGCTC | -3.299 | 100.9 |
|  |  | 36(R): GGGGCCAGACGCAACTCG |  |  |
| ***PDHPDA*** | Gene reference  (132 bp) | 258(F): GCGAAGCCGTTCTTCTCGAAG | -3.350 | 98.84 |
|  |  | 259(R): CTGAACTCGCTCCTCCCCC |  |  |
| ***rPDI^1^*** | Chaperon gene  (135 bp) | 24(F): GCCAGAGGACGCTTCTAACTTGG | -3.319 | 100.1 |
|  |  | 25(R): GCTGGAGTGGAGTCGGAAGTG |  |  |
| ***HPH*** | Hygromycin resistance gene (135 bp) | 4  1(F): CGACAGCGTCTCCGACCTG | -3.245 | 103.3 |
|  |  | 42(R): GGCGCAGCTATTTACCCGC |  |  |
| ***mCFH^1^*** | Murine CFH  (179 bp) | 212(F): CCGTCGCCATTCTCGACTC | -3.32 | 100.1 |
|  |  | 215(R): GCAGCCGAAACCGATCAGGAG |  |  |
| ***CEN3 core*** | Centromere (129 bp) | 364(F): CACGCGAGATGCACTCAGTGC | -3.213 | 104.7 |
|  |  | 365(R): CGTAGCTGCTTCATCCGGC |  |  |
| ***LHR^C^*** | synthetic DNA  (120 bp) | 370(F): CGTTACACGGCAGTCCTGTTCC | -3.223 | 104.3 |
|  |  | 371(R): CTTGTAGCTCAGCTTACCCACC |  |  |

^1^ – gene codon optimized

**Table 6 B** *Estimated GCN (normalised to GCN for ACT1 = 1)*

|  |  | **Analysed strains (all on background of CBS7435 *∆KU70)*** | | | | | | |
| --- | --- | --- | --- | --- | --- | --- | --- | --- |
| **GENE** | **Copy no. /Ave. *C_T_*** | yDA218  nChr 2B | yDA226  nChr 2B | yDA253  nChr 2A | yDA263  nChr 2A | yDA277  nChr 2A | yDA260  *P_AOX1_::mFH;* nChr 2A | yDA264  *P_AOX1_::mFH- P_AOX1_::PDI^H^* |
| *ACT1*  (Ref) | **Copy no.** | **1** | **1** | **1** | **1** | **1** | **1** | **1** |
|  | Average *C_T_* | 21.15±0.03 | 21.63±0.02 | 22.57±0.04 | 19.08±0.02 | 20.46±0.04 | 18.82±0.08 | 16.26±0.3 |
| *PDI^H^* | **Copy no**. | n/a | n/a | 1.1 | 1.0 | 1.2 | 1.1 | 1.2 |
|  | Average *C_T_* | n/a | n/a | 22.37±0.19 | 19.28±0.01 | 20.21±0.1 | 18.65±0.23 | 15.98±0.03 |
| *HPH* | **Copy no.** | n/a | **1.1** | **0.8** | **0.2** | **0.2** | **0.8** | **0.9** |
|  | Average *C_T_* | n/a | 21.52±0.12 | 22.88±0.26 | 21.17±0.22 | 22.75±0.13 | 19.14±0.25 | 16.44±0.13 |
| *CEN3* | **Copy no.** | **2.1** | **2.2** | **2.1** | **2.1** | **2.3** | **2.3** | **1.8** |
|  | Average *C_T_* | 20.06±0.02 | 20.50±0.03 | 22.53±0.13 | 18.01±0.01 | 19.25±0.11 | 17.60±0.08 | 15.43±0.14 |
| *LHR^C^* | **Copy no.** | **0.9** | **1.0** | **0.9** | **0.9** | **0.8** | **1.0** | **0.1** |
|  | Average *C_T_* | 21.25±0.07 | 21.64±0.06 | 22.67±0.03 | 19.28±0.01 | 20.8±0.05 | 18.81±0.22 | 19.95±0.03 |
| *mFH* | **Copy no.** | n/a | n/a | n/a | n/a | n/a | **1.1** | **1.2** |
|  | Average *C_T_* | n/a | n/a | n/a | n/a | n/a | 18.66±0.22 | 15.95±0.08 |
| *PDHPDA*  (2^nd^ Ref) | **Copy no.** | **0.9** | **0.9** | **0.9** | **1.1** | **1.0** | **1.0** | **0.9** |
|  | Average *C_T_* | 21.31±0.21 | 21.73±0.09 | 22.67±0.01 | 18.91±0.05 | 20.49±0.07 | 18.79±0.14 | 16.45±0.2 |

Average *C_T_* (cycle threshold) values were assayed from two biological sample repeats in three technical replicates. GCNs were calculated by normalizing *C_T_* values separately with respect to the housekeeping gene *ACT1*. *PDHPDA* is an additional control; both *ACT1* and PDHPDA are assumed to be present as single copies. The *HPH* gene encodes hygromycin B phosphotransferase.

Dalvie, N. C., J. Leal, C. A. Whittaker, Y. Yang, J. R. Brady, K. R. Love and J. C. Love (2020). "Host-Informed Expression of CRISPR Guide RNA for Genomic Engineering in Komagataella phaffii." ACS Synth Biol **9**(1): 26-35.

Kerr, H., A. P. Herbert, E. Makou, D. Abramczyk, T. H. Malik, H. Lomax-Browne, Y. Yang, I. Y. Pappworth, H. Denton, A. Richards, K. J. Marchbank, M. C. Pickering and P. N. Barlow (2021). "Murine Factor H Co-Produced in Yeast With Protein Disulfide Isomerase Ameliorated C3 Dysregulation in Factor H-Deficient Mice." Front Immunol **12**: 681098.

Shen, Y., Y. Wang, T. Chen, F. Gao, J. Gong, D. Abramczyk, R. Walker, H. Zhao, S. Chen, W. Liu, Y. Luo, C. A. Müller, A. Paul-Dubois-Taine, B. Alver, G. Stracquadanio, L. A. Mitchell, Z. Luo, Y. Fan, B. Zhou, B. Wen, F. Tan, J. Zi, Z. Xie, B. Li, K. Yang, S. M. Richardson, H. Jiang, C. E. French, C. A. Nieduszynski, R. Koszul, A. L. Marston, Y. Yuan, J. Wang, J. S. Bader, J. Dai, J. D. Boeke, X. Xu, Y. Cai and H. Yang (2017). "Deep functional analysis of synII, a 770-kilobase synthetic yeast chromosome." Science **355**(6329).

**REFERENCES**
