## Supplementary material for "A supernumerary synthetic chromosome in *Komagataella phaffii* as a repository for extraneous genetic material": Abramczyk et al AdditionalFille3.docx

Dariusz Abramczyk^1^, María del Carmen Sánchez Olmos^2^, Adán Andrés Ramírez Rojas^2^, Daniel Schindler^2,3^, Daniel Robertson^4^, Stephen McColm^5^, Adele L. Marston^6^, Paul N. Barlow^1,4^

^1^ School of Chemistry, University of Edinburgh, United Kingdom

^2^ Max Planck Institute for Terrestrial Microbiology, Marburg, Germany

^3^ Center for Synthetic Microbiology, Philipps-Universität Marburg, Marburg, Germany

^4^ School of Biological Sciences, University of Edinburgh, United Kingdom

^5^ Ingenza Ltd Scotland, United Kingdom

^6^ The Wellcome Centre for Cell Biology, Institute of Cell Biology, School of Biological Sciences, University of Edinburgh, United Kingdom

**Additional file 3** (DNA sequences)

**Contents** (also refer to Additional file 1: Fig. S1)

The validated sequences of the following plasmids and contigs are shown below.

**eDA40**

**eDA53** (framework plasmid (see Fig. 1a))

**eDA83**

**eDA110**

linearized **eDA137** (nanochromosome-precursor plasmid *v*1)

linearized **eDA155** (nanochromosome-precursor plasmid *v*2)

**eDA197** (contains integration array *LHR^E^- (P_TEF_)GFP(T_CYC1_)-Hyg^R^-LHR^Z^*

**eDA199** (contains insertion array *LHR^E^- (P_TEF_)mCH(T_CYC1_)-Zeo^R^-LHR^Z^*

linearized **eDA201** (nanochromosome-precursor plasmid *v*2B)

linearized **eDA229** (nanochromosome-precursor plasmid *v*2A)

**eDA250** (contains insertion array *LHR^E^-(P_TEF_)GFP(T_CYC1_)-LHR^D^-Zeo^R^-LHR^Z^*

Contig (length = 48123 bp): *de novo* assembly for strain **yDA122** (see Fig. 4)

Contig (length 37878 bp): *de novo* assembly for strain **yDA174**

Contig (length 54145 bp): *de novo* assembly for strain **yDA175**

**eDA40:**

gagatacctacagcgtgagctatgagaaagcgccacgcttcccgaagggagaaaggcggacaggtatccggtaagcggcagggtcggaacaggagagcgcacgagggagcttccagggggaaacgcctggtatctttatagtcctgtcgggtttcgccacctctgacttgagcgtcgatttttgtgatgctcgtcaggggggcggagcctatggaaaaacgccagcaacgcggcctttttacggttcctggccttttgctggccttttgctcacatgttctttcctgcgttatcccctgattctgtggataaccgtattaccgcctttgagtgagctgataccgctcgccgcagccgaacgaccgagcgcagcgagtcagtgagcgaggaagcggaagagcgcccaatacgcaaaccgcctctccccgcgcgttggccgattcattaatgcagctggcacgacaggtttcccgactggaaagcgggcagtgagcgcaacgcaattaatgtgagttagctcactcattaggcaccccaggctttacactttatgcttccggctcgtatgttgtgtggaattgtgagcggataacaatttcacacaggaaacagctatgaccatgattacgccaagcttgcatgcctgcaggtcgactctagaggggattcggcgctccaactgtattctaggcctttccacaatttaaaaaaagacctccgcgacggggaattgaaccccggtctaccgcgcgacaagcggtggttctaccactaaactatcacggatttcgatgctgctttcccacttccttactttacattctctaggcgtttagcctgtagaataaaaccttttttcaagactacaatactgctgctagtaataccgaattattacatgttttaacaacttgagagcgggtcgtatcctgtcttccgattgtacaatctcatcttatcagctcatctcatctcttcggaatcccccccgggtacccccccacacaccatagcttcaaaatgtttctactccttttttactcttccagattttctcggactccgcgcatcgccgtaccacttcaaaacacccaagcacagcatactaaattttccctctttcttcctctagggtgtcgttaattacccgtactaaaggtttggaaaagaaaaaagagaccgcctcgtttctttttcttcgtcgaaaaaggcaataaaaatttttatcacgtttctttttcttgaaatttttttttttagtttttttctctttcagtgacctccattgatatttaagttaataaacggtcttcaatttctcaagtttcagtttcatttttcttgttctattacaactttttttacttcttgttcattagaaagaaagcatagcaatctaatctaaggggcggtgttgacaattaatcatcggcatagtatatcggcatagtataatacgacaaggtgaggaactaaaccatggccaagttgaccagtgccgttccggtgctcaccgcgcgcgacgtcgccggagcggtcgagttctggaccgaccggctcgggttctcccgggacttcgtggaggacgacttcgccggtgtggtccgggacgacgtgaccctgttcatcagcgcggtccaggaccaggtggtgccggacaacaccctggcctgggtgtgggtgcgcggcctggacgagctgtacgccgagtggtcggaggtcgtgtccacgaacttccgggacgcctccgggccggccatgaccgagatcggcgagcagccgtgggggcgggagttcgccctgcgcgacccggccggcaactgcgtgcacttcgtggccgaggagcaggactgacacgtccgacggcggcccacgggtcccaggcctcggagatccgtcccccttttcctttgtcgatatcatgtaattagttatgtcacgcttacattcacgccctccccccacatccgctctaaccgaaaaggaaggagttagacaacctgaagtctaggtccctatttatttttttatagttatgttagtattaagaacgttatttatatttcaaatttttcttttttttctgtacagacgcgtgtacgcatgtaacattatactgaaaaccttgcttgagaaggttttgggacgctcgaaggctttaatttgcaagctggagaccaacggtaccgagctcgaattcactggccgtcgttttacaacgtcatgactgggaaaaccctggcgttacccaacttaatcgccttgcaacacatccccctttcgccagctggcgtaatagcgaagaggcccgcaccgatcgcccttcccaacagttgcgcagcctgaatggcgaatggcgcctgatgcggtattttctccttacgcatctgtgcggtatttcacaccgcatatggtgcactctcagtacaatctgctctgatgccgcatagttaagccagccccgacacccgccaacacccgctgacgcgccctgacgggcttgtctgctcccggcatccgcttacagacaagctgtgaccgtctccgggagctgcatgtgtcagaggttttcaccgtcatcaccgaaacgcgcgagacgaaagggcctcgtgatacgcctatttttataggttaatgtcatgataataatggtttcttagacgtcaggtggcacttttcggggaaatgtgcgcggaacccctatttgtttatttttctaaatacattcaaatatgtatccgctcatgagacaataaccctgataaatgcttcaataatattgaaaaaggaagagtatgagtattcaacatttccgtgtcgcccttattcccttttttgcggcattttgccttcctgtttttgctcacccagaaacgctggtgaaagtaaaagatgctgaagatcagttgggtgcacgagtgggttacatcgaactggatctcaacagcggtaagatccttgagagttttcgccccgaagaacgttttccaatgatgagcacttttaaagttctgctatgtggcgcggtattatcccgtattgacgccgggcaagagcaactcggtcgccgcatacactattctcagaatgacttggttgagtactcaccagtcacagaaaagcatcttacggatggcatgacagtaagagaattatgcagtgctgccataaccatgagtgataacactgcggccaacttacttctgacaacgatcggaggaccgaaggagctaaccgcttttttgcacaacatgggggatcatgtaactcgccttgatcgttgggaaccggagctgaatgaagccataccaaacgacgagcgtgacaccacgatgcctgtagcaatggcaacaacgttgcgcaaactattaactggcgaactacttactctagcttcccggcaacaattaatagactggatggaggcggataaagttgcaggaccacttctgcgctcggcccttccggctggctggtttattgctgataaatctggagccggtgagcgtgggtctcgcggtatcattgcagcactggggccagatggtaagccctcccgtatcgtagttatctacacgacggggagtcaggcaactatggatgaacgaaatagacagatcgctgagataggtgcctcactgattaagcattggtaactgtcagaccaagtttactcatatatactttagattgatttaaaacttcatttttaatttaaaaggatctaggtgaagatcctttttgataatctcatgaccaaaatcccttaacgtgagttttcgttccactgagcgtcagaccccgtagaaaagatcaaaggatcttcttgagatcctttttttctgcgcgtaatctgctgcttgcaaacaaaaaaaccaccgctaccagcggtggtttgtttgccggatcaagagctaccaactctttttccgaaggtaactggcttcagcagagcgcagataccaaatactgttcttctagtgtagccgtagttaggccaccacttcaagaactctgtagcaccgcctacatacctcgctctgctaatcctgttaccagtggctgctgccagtggcgataagtcgtgtcttaccgggttggactcaagacgatagttaccggataaggcgcagcggtcgggctgaacggggggttcgtgcacacagcccagcttggagcgaacgacctacaccgaact

**eDA53:**

agttcggtgtaggtcgttcgctccaagctgggctgtgtgcacgaaccccccgttcagcccgaccgctgcgccttatccggtaactatcgtcttgagtccaacccggtaagacacgacttatcgccactggcagcagccactggtaacaggattagcagagcgaggtatgtaggcggtgctacagagttcttgaagtggtggcctaactacggctacactagaagaacagtatttggtatctgcgctctgctgaagccagttaccttcggaaaaagagttggtagctcttgatccggcaaacaaaccaccgctggtagcggtggtttttttgtttgcaagcagcagattacgcgcagaaaaaaaggatctcaagaagatcctttgatcttttctacggggtctgacgctcagtggaacgaaaactcacgttaagggattttggtcatgagattatcaaaaaggatcttcacctagatccttttaaattaaaaatgaagttttaaatcaatctaaagtatatatgagtaaacttggtctgacagttaccaatgcttaatcagtgaggcacctatctcagcgatctgtctatttcgttcatccatagttgcctgactccccgtcgtgtagataactacgatacgggagggcttaccatctggccccagtgctgcaatgataccgcgagacccacgctcaccggctccagatttatcagcaataaaccagccagccggaagggccgagcgcagaagtggtcctgcaactttatccgcctccatccagtctattaattgttgccgggaagctagagtaagtagttcgccagttaatagtttgcgcaacgttgttgccattgctacaggcatcgtggtgtcacgctcgtcgtttggtatggcttcattcagctccggttcccaacgatcaaggcgagttacatgatcccccatgttgtgcaaaaaagcggttagctccttcggtcctccgatcgttgtcagaagtaagttggccgcagtgttatcactcatggttatggcagcactgcataattctcttactgtcatgccatccgtaagatgcttttctgtgactggtgagtactcaaccaagtcattctgagaatagtgtatgcggcgaccgagttgctcttgcccggcgtcaatacgggataataccgcgccacatagcagaactttaaaagtgctcatcattggaaaacgttcttcggggcgaaaactctcaaggatcttaccgctgttgagatccagttcgatgtaacccactcgtgcacccaactgatcttcagcatcttttactttcaccagcgtttctgggtgagcaaaaacaggaaggcaaaatgccgcaaaaaagggaataagggcgacacggaaatgttgaatactcatactcttcctttttcaatattattgaagcatttatcagggttattgtctcatgagcggatacatatttgaatgtatttagaaaaataaacaaataggggttccgcgcacatttccccgaaaagtgccacctgacgtctaagaaaccattattatcatgacattaacctataaaaataggcgtatcacgaggccctttcgtctcgcgcgtttcggtgatgacggtgaaaacctctgacacatgcagctcccggagacggtcacagcttgtctgtaagcggatgccgggagcagacaagcccgtcagggcgcgtcagcgggtgttggcgggtgtcggggctggcttaactatgcggcatcagagcagattgtactgagagtgcaccatatgttgctccgtcagcttgaataagcccctgaaatagaacgtagtcttgaatgctcagttagcaagtctgttactatctattttataacgcaactcaatgtttttgtttctctcgacgaaacgtttttgcttagcatttcggtatcttaaaccgtgtgattttaagtctcataaccttttacatacaggatcaataataatattattgttgatacaatataggattagaatacatagattctgatgttcgaatcaatcaatgtcttaatattcacgtacttgtataccctctgattccacattggtccaacggaatcttctagcaacagccagttgcactagatgtggttctccaccagaaattatagtactgacagattttttggtttacggtgagaggctccaatttccaaagctaaaccaatttcttcataagcaacctccacgcgcgcttttagtatgatgtagaaactggacctaaagatcgaccaatacttgcttacacggcagcgcgaaataaaagacattgataatgtagagtaagtactatttcttcaaaagatagaaccactctcagaggctaactagaagatctaaaacggatgtacagctaatatgtttcgaggcaatcatgttcaggtgcgcctcaagtggctcttaatctattttaaagcataatccacgaaaatttcgatacagagagatacgtcataaactgagaacgatagttagaggcatagcttcggcaatgttcgataaacaagtcaacgagtcagaatttcatgtttcttttttttcttgtagtaatatgtaataaccgaatcagataacttgccagtaatgacctgagctctataaattgaaaccgctacaaaaagtaagggatcgtgttaagggcacagagaagaaagtgtaaagaagtaagctagcgttgtacaacaactcaaaagcatggagcccggtgggtactagataataagaacaaggtttcagtgagtttacaagttcataaaacacggaacctggcaacatagcataccagtaaatcgatttgaagaactctattaggctggcagcctctctcaatcaatagtctaaaacggttaagagacacaatcatctctaggtgtcgttgtggtctaccatcgaaacactggaaatcaatccaatgaacttcaagtgatatatttttcaagagtcttcaaagaccaagcaatgttcttattgattgataattttaaagttgaagtctaaatttgtagcagtgaagtctgagcagggggaatgttaagccatatctaatagtacctattagacgcttgattgccgtttgcaaatatgccatgtttctttgaaaccaaaaccacgtaagggggcttaagttttcggcttaggattgttacggagctcaaaccaaataaggcagagagcatacaaaacgtttattaaaaagaacaactacttctggatagctcaattatcttttgtttcttttgaggcgtgcccttcatgacatgacattgcatccatacaattaatagtagataggggaggataagtttgctagtgtgagcatttagagcaaggggtcagtttcctccatctgcctactctgctccatttaaaagcgagccgattgccataagcttgctcacgtagataggacttcaaaaagacgttaagaggctgctctctagaatacgatggaataaacaaatctcgttagtttcttgaaacggaaagtacatgtaggataattgctggatcattcggaagctaaaattggtcctttccccaaagacaatcgactatagttcctgaagcttcctgaggtgagctggatagaaccagaagatagtacctataacgtcaacaacaactgaaactacagatggaagagctaggcatatccataggggatagccatagaaaactttgattgacgcagcaaatcgagacgttaacctgaagctgccataacttcaggaacttgaacacaagagtggcattaaaattcctatgttgcatttcagaaataccacaagtaaactgagcaattaacttgttcataccgacactacaagttaactacaaaccgagacccttatatgcagactgataatacagaatgatacgtatcactcctaagactaggaacatacagagcttttgagcttttggtattttcaagttatttgaaaaattaaatctttatactagggacgaggttcgtgacaaaagaagacaatcatgagactcaccgtcttgtcatctacaagtttcaataagcttccaacttgagtaacctagccatgtgattagtaactttcgaagcatgattcagaacgttctgctctgccgcgtcaaaaagggcagctacttgaagttaaaagaaagatcaaaaattcctgagcattgttcaaacagctataaatcaaaacagaataagaaataaggccatcacttaccaaggaaaaacaaaaaagttcgagagatcaacaatgcacttcagggacgcacctgaacctttacattagtgtaaaaataaattattagattgttcggttcgagcgttagccatatacaaaatgcggccttcaaagtcattggaaaagagctgctttaggacctctccaaaagctaaattgacaaacaggctacctcatatctccatcatctttgccaaccttttcagtaacaaaaaacaaaagcaatatcgtttatttgtccattggacgatgaagaaaagatcaaacttagtctaggtgaggctaatcaataaagtttgttgatcctactaattacgatcctatcctagccttagcatttacatctaaaggtctggtttcacattctaattgtgaaatgatctagcaagctctccctagcataattattcacctccaaatttgtgtacttggttcatcgttcactgaatgagaatgagaaagattcaaaattgagaagtgtgttcgaacacttcaactgtcgcactgttgctttctacagatatacgcataacaaatacattagttgcttaaagggcagcttcttttaatgtgtatcagtatagtcaaggactcagctacgaacaactcacaatagtctttttcaccacgacccacaagttcgcacaagaattctaaataccttgacgaatagtaaatatcggatgttcagccgtcagcgaaatgatttttgtgctaatacaatttctaaggcaattgacacaggttggaaaaagtgtgggtattcggctgatattgtggaactgtaatcatcgagctaccagttctggtttgttacttgttcgtctcattgtcgtggataagaacaaataatgtaccggtttattcgaccaccaatcagcgcgtaaagcacgcagcagaccaagcaagtagccggatgaagcagctacgttacccaaccgtttatcctacccaagtaggcgagttcaatttgtctcacgcgagatgcactcagtgctacaagattgaatgggtaattagaagcatgttagtttctacaatgatcgattcatagtatccggcgacaattgatcttactgaagaaaggaggcagacacgcttaccgaaaaaagtgcctacctagcaatatttgatgatcgtccaatggacaaataaacgatattgcttttgttttttgttactgaaaaggttggcaaagatgatggagatatgaggtagcctgtttgtcaatttagcttttggagaggtcctaaagcagctcttttccaatgactttgaaggccgcattttgtatatggctaacgctcgaaccgaacaatctaataatttatttttacactaatgtaaaggttcaggtgcgtccctgaagtgcattgttgatctctcgaacttttttgtttttccttggtaagtgatggccttatttcttattctgttttgatttatagctgtttgaacaatgctcaggaatttttgatctttcttttaacttcaagtagctgccctttttgacgcggcagagcagaacgttctgaatcatgcttcgaaagttactaatcacatggctaggttactcaagttggaagcttattgaaacttgtagatgacaagacggtgagtctcatgattgtcttcttttgtcacgaacctcgtccctagtataaagatttaatttttcaaataacttgaaaataccaaaagctcaaaagctctgtatgttcctagtcttaggagtgatacgtatcattctgtattatcagtctgcatataagggtctcggtttgtagttaacttgtagtgtcggtatgaacaagttaattgctcagtttacttgtggtatttctgaaatgcaacataggaattttaatgccactcttgtgttcaagttcctgaagttatggcagcttcaggttaacgtctcgatttgctgcgtcaatcaaagttttctatggctatcccctatggatatgcctagctcttccatctgtagtttcagttgttgttgacgttataggtactatcttctggttctatccagctcacctcaggaagcttcaggaactatagtcgattgtctttggggaaaggaccaattttagcttccgaatgatccagcaattatcctacatgtactttccgtttcaagaaactaacgagatttgtttattccatcgtattctagagagcagcctcttaacgtctttttgaagtcctatctacgtgagcaagcttatggcaatcggctcgcttttaaatggagcagagtaggcagatggaggaaactgaccccttgctctaaatgctcacactagcaaacttatcctcccctatctactattaattgtatggatgcaatgtcatgtcatgaagggcacgcctcaaaagaaacaaaagataattgagctatccagaagtagttgttctttttaataaacgttttgtatgctctctgccttatttggtttgagctccgtaacaatcctaagccgaaaacttaagcccccttacgtggttttggtttcaaagaaacatggcatatttgcaaacggcaatcaagcgtctaataggtactattagatatggcttaacattccccctgctcagacttcactgctacaaatttagacttcaactttaaaattatcaatcaataagaacattgcttggtctttgaagactcttgaaaaatatatcacttgaagttcattggattgatttccagtgtttcgatggtagaccacaacgacacctagagatgattgtgtctcttaaccgttttagactattgattgagagaggctgccagcctaatagagttcttcaaatcgatttactggtatgctatgttgccaggttccgtgttttatgaacttgtaaactcactgaaaccttgttcttattatctagtacccaccgggctccatgcttttgagttgttgtacaacgctagcttacttctttacactttcttctctgtgcccttaacacgatcccttactttttgtagcggtttcaatttatagagctcaggtcattactggcaagttatctgattcggttattacatattactacaagaaaaaaagaaacatgaaattctgactcgttgacttgtttatcgaacattgccgaagctatgcctctaactatcgttctcagtttatgacgtatctctctgtatcgaaattttcgtggattatgctttaaaatagattaagagccacttgaggcgcacctgaacatgattgcctcgaaacatattagctgtacatccgttttagatcttctagttagcctctgagagtggttctatcttttgaagaaatagtacttactctacattatcaatgtcttttatttcgcgctgccgtgtaagcaagtattggtcgatctttaggtccagtttctacatcatactaaaagcgcgcgtggaggttgcttatgaagaaattggtttagctttggaaattggagcctctcaccgtaaaccaaaaaatctgtcagtactataatttctggtggagaaccacatctagtgcaactggctgttgctagaagattccgttggaccaatgtggaatcagagggtatacaagtacgtgaatattaagacattgattgattcgaacatcagaatctatgtattctaatcctatattgtatcaacaataatattattattgatcctgtatgtagaaggttatgagacttaaaatcacacggtttaagataccgaaatgctaagcaaaaacgtttcgtcgagagaaacaaaaacattgagttgtgaatgggacaatttgaattaaagaatgtatggaaagataacgctgactggtgtaccacttgtcggtgggacttgatgcttggcaagctcaagaagtcacttgatctacggtacccgggatgtgctgcaaggcgattaagttgggtaacgccagggttttcccagtcatgacgttgtaaaacgacggccagtgaattcgagctcggtacccgggggggattccgaagagatgagatgagctgataagatgagattgtacaatcggaagacaggatacgacccgctctcaagttgttaaaacatgtaataattcggtattactagcagcagtattgtagtcttgaaaaaaggttttattctacaggctaaacgcctagagaatgtaaagtaaggaagtgggaaagcagcatcgaaatccgtgatagtttagtggtagaaccaccgcttgtcgcgcggtagaccggggttcaattccccgtcgcggaggtctttttttaaattgtggaaaggcctagaatacagttggagcgccgaatcccctctagagtcgacctgcaggcatgcaagcttggcgtaatcatggtcatagctgtttcctgtgtgaaattgttatccgctcacaattcgcggcctttttacggttcctggccttttgctggccttttgctcacatgttggtctccagcttgcaaattaaagccttcgagcgtcccaaaaccttctcaagcaaggttttcagtataatgttacatgcgtacacgcgtctgtacagaaaaaaaagaaaaatttgaaatataaataacgttcttaatactaacataactataaaaaaataaatagggacctagacttcaggttgtctaactccttccttttcggttagagcggatgtggggggagggcgtgaatgtaagcgtgacataactaattacatgatatcgacaaaggaaaagggggacggatctccgaggcctcggacccgtcgggccgcgtcggacgtgtcagtcctgctcctcggccacgaagtgcacgcagttgccggccgggtcgcgcagggcgaactcccgcccccacggctgctcgccgatctcggtcatggccggcccggaggcgtcccggaagttcgtggacacgacctccgaccactcggcgtacagctcgtccaggccgcgcacccacacccaggccagggtgttgtccggcaccacctggtcctggaccgcgctgatgaacagggtcacgtcgtcccggaccacaccggcgaagtcgtcctccacgaagtcccgggagaacccgagccggtcggtccagaactcgaccgctccggcgacgtcgcgcgcggtgagcaccggaacggcactggtcaacttggccatggtttagttcctcaccttgtcgtattatactatgccgatatactatgccgatgattaattgtcaacaccgcccttagattagattgctatgctttctttctaatgagcaagaagtaaaaaaagttgtaatagaacaagaaaaatgaaactgaaacttgagaaattgaagaccgtttattaacttaaatatcaatgggaggtcatcgaaagagaaaaaaatcaaaaaaaaaaattttcaagaaaaagaaacgtgataaaaatttttattgcctttttcgacgaagaaaaagaaacgaggcggtctcttttttcttttccaaacctttagtacgggtaattaacgacaccctagaggaagaaagaggggaaatttagtatgctgtgcttgggtgttttgaagtggtacggcgatgcgcggagtccgagaaaatctggaagagtaaaaaaggagtagaaacattttgaagctatggtgtgtgggggatccgcacaaacgaaggtctcacttaatcttctgtactctgaagaggagtgggaaataccaagaaaaacatcaaactcgaatgattttcccaaacccctaccacaagatattcatcagctgcgagataggctgatcaggagcaagctcgtacgagaagaaacaaaatgacaactgcattaatgaatcggccaacgcgcggggagaggcggtttgcgtattgggcgctcttccgcttcctcgctcactgactcgctgcgctcggtcgttcggctgcggcgagcggtatcagctcactcaaaggcggtaatacggttatccacagaatcaggggataacgcaggaaagaacatgtgagcaaaaggccagcaaaaggccaggaaccgtaaaaaggccgcgttgctggcgtttttccataggctccgcccccctgacgagcatcacaaaaatcgacgctcaagtcagaggtggcgaaacccgacaggactataaagataccaggcgtttccccctggaagctccctcgtgcgctctcctgttccgaccctgccgcttaccggatacctgtccgcctttctcccttcgggaagcgtggcgctttctcatagctcacgctgtaggtatctc

**eDA83:**

gagatacctacagcgtgagctatgagaaagcgccacgcttcccgaagggagaaaggcggacaggtatccggtaagcggcagggtcggaacaggagagcgcacgagggagcttccagggggaaacgcctggtatctttatagtcctgtcgggtttcgccacctctgacttgagcgtcgatttttgtgatgctcgtcaggggggcggagcctatggaaaaacgccagcaacgcggcctttttacggttcctggccttttgctggccttttgctcacatgttctttcctgcgttatcccctgattctgtggataaccgtattaccgcctttgagtgagctgataccgctcgccgcagccgaacgaccgagcgcagcgagtcagtgagcgaggaagcggaagagcgcccaatacgcaaaccgcctctccccgcgcgttggccgattcattaatgcagttgtcattttgtttcttctcgtacgagcttgctcctgatcagcctatctcgcagctgtgttaaaccgtctttaagtcaacccacagtctactgcaatcgtattcagaactagccactagactgttacaagtcgaaacctcagttaaccaactagaaactctagttaccaagttagaactgtgagtacgaaaagtctgaaaagcagaaagattcaatagatttgtctgtgttacacaagagttcaacaagtagctgcgttcgtgctagttgtctaggtttaaacgtttaaaaagacaactagtagtttacttcacctctgtattactgacggtttgtgcctcaacagtttacgttaacaaactagtaagcgtctacttcgtggacttgtacttttagtagaaagattgagtgtctggcagttttaaacctaagtcagttagaatctcaatcctctgcctttggtttagaaacagtgtagctgttttgacctagactgagtacaactgttcttgctgcttattcgaaacttgcctaactcactgcaacagacaagtctagtttagaataaagtaggctcagttattcaagtctaacttcagacagtgtttctcagatttgacttaagtagacagtagtgtgtgaaaaaccagactagtttgaatcttagcctgttcagaaaaaactaactgacttgtgtcaggcctcccaaaaaaactaacttcaatcctctgtactattccacaactattagttgttgcaagtctgaagcctagtcttgctttgaccttacaggatagaggttggctcacaactgagtctgtgtgaaaacaaaagtaaagattgccacaagaagtctattctagttagtccttttacgttggatagcaatagtccaatcttagtttgcagctgatgaatatcttgtggtaggggtttgggaaaatcattcgagtttgatgtttttcttggtatttcccactcctcttcagagtacagaagattaagtgagaccttcgtttgtgcggatcccccacacaccatagcttcaaaatgtttctactccttttttactcttccagattttctcggactccgcgcatcgccgtaccacttcaaaacacccaagcacagcatactaaatttcccctctttcttcctctagggtgtcgttaattacccgtactaaaggtttggaaaagaaaaaagagaccgcctcgtttctttttcttcgtcgaaaaaggcaataaaaatttttatcacgtttctttttcttgaaaatttttttttttgatttttttctctttcgatgacctcccattgatatttaagttaataaacggtcttcaatttctcaagtttcagtttcatttttcttgttctattacaactttttttacttcttgctcattagaaagaaagcatagcaatctaatctaagggcggtgttgacaattaatcatcggcatagtatatcggcatagtataatacgacaaggtgaggaactaaaccatggccaagttgaccagtgccgttccggtgctcaccgcgcgcgacgtcgccggagcggtcgagttctggaccgaccggctcgggttctcccgggacttcgtggaggacgacttcgccggtgtggtccgggacgacgtgaccctgttcatcagcgcggtccaggaccaggtggtgccggacaacaccctggcctgggtgtgggtgcgcggcctggacgagctgtacgccgagtggtcggaggtcgtgtccacgaacttccgggacgcctccgggccggccatgaccgagatcggcgagcagccgtgggggcgggagttcgccctgcgcgacccggccggcaactgcgtgcacttcgtggccgaggagcaggactgacacgtccgacgcggcccgacgggtccgaggcctcggagatccgtcccccttttcctttgtcgatatcatgtaattagttatgtcacgcttacattcacgccctccccccacatccgctctaaccgaaaaggaaggagttagacaacctgaagtctaggtccctatttatttttttatagttatgttagtattaagaacgttatttatatttcaaatttttcttttttttctgtacagacgcgtgtacgcatgtaacattatactgaaaaccttgcttgagaaggttttgggacgctcgaaggctttaatttgcaagctggagaccaacatgtgagcaaaaggccagcaaaaggccaggaaccgtaaaaaggccgcgaattgtgagcggataacaatttcacacaggaaacagctatgaccatgattacgccaagcttgcatgctgtatctaacatccaaagacgaaaggttgaatgaaacctttttgccatccgacatccacaggtccattctcacacataagtgccaaacgcaacaggaggggatacactagcagcagaccgttgcaaacgcaggacctccactcctcttctcctcaacacccacttttgccatcgaaaaaccagcccagttattgggcttgattggagctcgctcattccaattccttctattaggctactaacaccatgactttattagcctgtctatcctggcccccctggcgaggttcatgtttgtttatttccgaatgcaacaagctccgcattacacccgaacatcactccagatgagggctttctgagtgtggggtcaaatagtttcatgttccccaaatggcccaaaactgacagtttaaacgctgtcttggaacctaatatgacaaaagcgtgatctcatccaagatgaactaagtttggttcgttgaaatgctaacggccagttggtcaaaaagaaacttccaaaagtcggcataccgtttgtcttgtttggtattgattgacgaatgctcaaaaataatctcattaatgcttagcgcagtctctctatcgcttctgaaccccggtgcacctgtgccgaaacgcaaatggggaaacacccgctttttggatgattatgcattgtctccacattgtatgcttccaagattctggtgggaatactgctgatagcctaacgttcatgatcaaaatttaactgttctaacccctacttgacagcaatatataaacagaaggaagctgccctgtcttaaacctttttttttatcatcattattagcttactttcataattgcgactggttccaattgacaagcttttgattttaacgacttttaacgacaacttgagaagatcaaaaaacaactaattattcgaaacgaggaattcatgggatcaagatcgccaaaaaagaagagaaaggtgccgaagaagcatgcagcaccaccaaaaaaaaaacgaaaagtagaagacccacgatttatgtacccatacgatgttcctgactatgcgggtatgaaaaacatcaaaaaaaaccaggtaatgaacctgggtccgaactctaaactgctgaaagaatacaaatcccagctgatcgaactgaacatcgaacagttcgaagcaggtatcggtctgatcctgggtgatgcttacatccgttctcgtgatgaaggtaaaacctactgtatgcagttcgagtggaaaaacaaagcatacatggaccacgtatgtctgctgtacgatcagtgggtactgtccccgccgcacaaaaaagaacgtgttaaccacctgggtaacctggtaatcacctggggcgcccagactttcaaacaccaagctttcaacaaactggctaacctgttcatcgttaacaacaaaaaaaccatcccgaacaacctggttgaaaactacctgaccccgatgtctctggcatactggttcatggatgatggtggtaaatgggattacaacaaaaactctaccaacaaatcgatcgtactgaacacccagtctttcactttcgaagaagtagaatacctggttaagggtctgcgtaacaaattccaactgaactgttacgtaaaaatcaacaaaaacaaaccgatcatctacatcgattctatgtcttacctgatcttctacaacctgatcaaaccgtacctgatcccgcagatgatgtacaaactgccgaacactatctcctccgaaactttcctgaaataaccgcggcggccgccagcttgggcccgaacaaaaactcatctcagaagaggatctgaatagcgccgtcgaccatcatcatcatcatcattgagttttagccttagacatgactgttcctcagttcaagttgggcacttacgagaagaccggtcttgctagattctaatcaagaggatgtcagaatgccatttgcctgagagatgcaggcttcatttttgatacttttttatttgtaacctatatagtataggattttttttgtcattttgtttcttctcgtacgagcttgctcctgatcagcctatctcgcagctgatgaatatcttgtggtaggggtttgggaaaatcattcgagtttgatgtttttcttggtatttcccactcctcttcagagtacagaagattaagtgagccaaccgtgtaggtctacaaactggtattctcgtttgttgtgaatcttgtcttttctgcttactttctaatagattgagtgagttaggtctgtatttgacggctaaagttaaacttaactgttgtgtgggatagttgttagctgtcaagaaaccgttaaacgtagaaagagtaacttttctaaaccgtagtgtctactcaatattgagacaagcagctagtttaagagcagaagctgaagtctgtgagtttggaaactaaaaggtctgtgaacagctaactagaactaggcttgcttttagtattagaataacaacaaacgaggtctattagttgtgggatctagctagttgcctaagaacaaaggactaagtctgcagaaaaacgagtttcaataggtctgtgtgtaacagtgttaaggctactagcagaactgccacctaagtttcgtaagttccacctcagccttcaatattccagacttgaccaactttcagtttttaaactctaccgtactagtagtttctgcacagtattagacctacttacactgtctgtgcttgcaccacaaaccaaatagtagaatctcttttaagtagaacagtcttttcttgttagactaaaacaacgagagacaactgagtgtagggttacagttaaaaaactgtacagtctgctgtacagacgttacacggcagtcctgttccaaacttcacttaagggttagattggctaagttcaacttgtccttttaagttagttcagacctagttagcagacttaggtgggtaagctgagctacaagagttaaaaacaactgctaagactaaagtaagtgtacgtaagcttcaaactagtaagttaaacttagaaacagctaacgtctaaagctagaagtaagctcaatcttagtactgttaacctaagttgctcacgcaccttcgccaactcaacctgcaggtcgactctagaggggattcggcgctccaactgtattctaggcctttccacaatttaaaaaaagacctccgcgacggggaattgaaccccggtctaccgcgcgacaagcggtggttctaccactaaactatcacggatttcgatgctgctttcccacttccttactttacattctctaggcgtttagcctgtagaataaaaccttttttcaagactacaatactgctgctagtaataccgaattattacatgttttaacaacttgagagcgggtcgtatcctgtcttccgattgtacaatctcatcttatcagctcatctcatctcttcggaatcccccccgggtaccgagctcgaattcactggccgtcgttttacaacgtcatgactgggaaaaccctggcgttacccaacttaatcgccttgcagcacatcccgggtaccgtagatcaagtgacttcttgagcttgccaagcatcaagtcccaccgacaagtggtacaccagtcagcgttatctttccatacattctttaattcaaattgtcccattcacaactcaatgtttttgtttctctcgacgaaacgtttttgcttagcatttcggtatcttaaaccgtgtgattttaagtctcataaccttctacatacaggatcaataataatattattgttgatacaatataggattagaatacatagattctgatgttcgaatcaatcaatgtcttaatattcacgtacttgtataccctctgattccacattggtccaacggaatcttctagcaacagccagttgcactagatgtggttctccaccagaaattatagtactgacagattttttggtttacggtgagaggctccaatttccaaagctaaaccaatttcttcataagcaacctccacgcgcgcttttagtatgatgtagaaactggacctaaagatcgaccaatacttgcttacacggcagcgcgaaataaaagacattgataatgtagagtaagtactatttcttcaaaagatagaaccactctcagaggctaactagaagatctaaaacggatgtacagctaatatgtttcgaggcaatcatgttcaggtgcgcctcaagtggctcttaatctattttaaagcataatccacgaaaatttcgatacagagagatacgtcataaactgagaacgatagttagaggcatagcttcggcaatgttcgataaacaagtcaacgagtcagaatttcatgtttctttttttcttgtagtaatatgtaataaccgaatcagataacttgccagtaatgacctgagctctataaattgaaaccgctacaaaaagtaagggatcgtgttaagggcacagagaagaaagtgtaaagaagtaagctagcgttgtacaacaactcaaaagcatggagcccggtgggtactagataataagaacaaggtttcagtgagtttacaagttcataaaacacggaacctggcaacatagcataccagtaaatcgatttgaagaactctattaggctggcagcctctctcaatcaatagtctaaaacggttaagagacacaatcatctctaggtgtcgttgtggtctaccatcgaaacactggaaatcaatccaatgaacttcaagtgatatatttttcaagagtcttcaaagaccaagcaatgttcttattgattgataattttaaagttgaagtctaaatttgtagcagtgaagtctgagcagggggaatgttaagccatatctaatagtacctattagacgcttgattgccgtttgcaaatatgccatgtttctttgaaaccaaaaccacgtaagggggcttaagttttcggcttaggattgttacggagctcaaaccaaataaggcagagagcatacaaaacgtttattaaaaagaacaactacttctggatagctcaattatcttttgtttcttttgaggcgtgcccttcatgacatgacattgcatccatacaattaatagtagataggggaggataagtttgctagtgtgagcatttagagcaaggggtcagtttcctccatctgcctactctgctccatttaaaagcgagccgattgccataagcttgctcacgtagataggacttcaaaaagacgttaagaggctgctctctagaatacgatggaataaacaaatctcgttagtttcttgaaacggaaagtacatgtaggataattgctggatcattcggaagctaaaattggtcctttccccaaagacaatcgactatagttcctgaagcttcctgaggtgagctggatagaaccagaagatagtacctataacgtcaacaacaactgaaactacagatggaagagctaggcatatccataggggatagccatagaaaactttgattgacgcagcaaatcgagacgttaacctgaagctgccataacttcaggaacttgaacacaagagtggcattaaaattcctatgttgcatttcagaaataccacaagtaaactgagcaattaacttgttcataccgacactacaagttaactacaaaccgagacccttatatgcagactgataatacagaatgatacgtatcactcctaagactaggaacatacagagcttttgagcttttggtattttcaagttatttgaaaaattaaatctttatactagggacgaggttcgtgacaaaagaagacaatcatgagactcaccgtcttgtcatctacaagtttcaataagcttccaacttgagtaacctagccatgtgattagtaactttcgaagcatgattcagaacgttctgctctgccgcgtcaaaaagggcagctacttgaagttaaaagaaagatcaaaaattcctgagcattgttcaaacagctataaatcaaaacagaataagaaataaggccatcacttaccaaggaaaaacaaaaaagttcgagagatcaacaatgcacttcagggacgcacctgaacctttacattagtgtaaaaataaattattagattgttcggttcgagcgttagccatatacaaaatgcggccttcaaagtcattggaaaagagctgctttaggacctctccaaaagctaaattgacaaacaggctacctcatatctccatcatctttgccaaccttttcagtaacaaaaaacaaaagcaatatcgtttatttgtccattggacgatcatcaaatattgctaggtaggcacttttttcggtaagcgtgtctgcctcctttcttcagtaagatcaattgtcgccggatactatgaatcgatcattgtagaaactaacatgcttctaattacccattcaatcttgtagcactgagtgcatctcgcgtgagacaaattgaactcgcctacttgggtaggataaacggttgggtaacgtagctgcttcatccggctacttgcttggtctgctgcgtgctttacgcgctgattggtggtcgaataaaccggtacattatttgttcttatccacgacaatgagacgaacaagtaacaaaccagaactggtagctcgatgattacagttccacaatatcagccgaatacccacactttttccaacctgtgtcaattgccttagaaattgtattagcacaaaaatcatttcgctgacggctgaacatccgatatttactattcgtcaaggtatttagaattcttgtgcgaacttgtgggtcgtggtgaaaaagactattgtgagttgttcgtagctgagtccttgactatactgatacacattaaaagaagctgccctttaagcaactaatgtatttgttatgcgtatatctgtagaaagcaacagtgcgacagttgaagtgttcgaacacacttctcaattttgaatctttctcattctcattcagtgaacgatgaaccaagtacacaaatttggaggtgaataattatgctagggagagcttgctagatcatttcacaattagaatgtgaaaccagacctttagatgtaaatgctaaggctaggataggatcgtaattagtaggatcaacaaactttattgattagcctcacctagactaagtttgatcttttcttcatcgtccaatggacaaataaacgatattgcttttgttttttgttactgaaaaggttggcaaagatgatggagatatgaggtagcctgtttgtcaatttagcttttggagaggtcctaaagcagctcttttccaatgactttgaaggccgcattttgtatatggctaacgctcgaaccgaacaatctaataatttatttttacactaatgtaaaggttcaggtgcgtccctgaagtgcattgttgatctctcgaacttttttgtttttccttggtaagtgatggccttatttcttattctgttttgatttatagctgtttgaacaatgctcaggaatttttgatctttcttttaacttcaagtagctgccctttttgacgcggcagagcagaacgttctgaatcatgcttcgaaagttactaatcacatggctaggttactcaagttggaagcttattgaaacttgtagatgacaagacggtgagtctcatgattgtcttcttttgtcacgaacctcgtccctagtataaagatttaatttttcaaataacttgaaaataccaaaagctcaaaagctctgtatgttcctagtcttaggagtgatacgtatcattctgtattatcagtctgcatataagggtctcggtttgtagttaacttgtagtgtcggtatgaacaagttaattgctcagtttacttgtggtatttctgaaatgcaacataggaattttaatgccactcttgtgttcaagttcctgaagttatggcagcttcaggttaacgtctcgatttgctgcgtcaatcaaagttttctatggctatcccctatggatatgcctagctcttccatctgtagtttcagttgttgttgacgttataggtactatcttctggttctatccagctcacctcaggaagcttcaggaactatagtcgattgtctttggggaaaggaccaattttagcttccgaatgatccagcaattatcctacatgtactttccgtttcaagaaactaacgagatttgtttattccatcgtattctagagagcagcctcttaacgtctttttgaagtcctatctacgtgagcaagcttatggcaatcggctcgcttttaaatggagcagagtaggcagatggaggaaactgaccccttgctctaaatgctcacactagcaaacttatcctcccctatctactattaattgtatggatgcaatgtcatgtcatgaagggcacgcctcaaaagaaacaaaagataattgagctatccagaagtagttgttctttttaataaacgttttgtatgctctctgccttatttggtttgagctccgtaacaatcctaagccgaaaacttaagcccccttacgtggttttggtttcaaagaaacatggcatatttgcaaacggcaatcaagcgtctaataggtactattagatatggcttaacattccccctgctcagacttcactgctacaaatttagacttcaactttaaaattatcaatcaataagaacattgcttggtctttgaagactcttgaaaaatatatcacttgaagttcattggattgatttccagtgtttcgatggtagaccacaacgacacctagagatgattgtgtctcttaaccgttttagactattgattgagagaggctgccagcctaatagagttcttcaaatcgatttactggtatgctatgttgccaggttccgtgttttatgaacttgtaaactcactgaaaccttgttcttattatctagtacccaccgggctccatgcttttgagttgttgtacaacgctagcttacttctttacactttcttctctgtgcccttaacacgatcccttactttttgtagcggtttcaatttatagagctcaggtcattactggcaagttatctgattcggttattacatattactacaagaaaaaaaagaaacatgaaattctgactcgttgacttgtttatcgaacattgccgaagctatgcctctaactatcgttctcagtttatgacgtatctctctgtatcgaaattttcgtggattatgctttaaaatagattaagagccacttgaggcgcacctgaacatgattgcctcgaaacatattagctgtacatccgttttagatcttctagttagcctctgagagtggttctatcttttgaagaaatagtacttactctacattatcaatgtcttttatttcgcgctgccgtgtaagcaagtattggtcgatctttaggtccagtttctacatcatactaaaagcgcgcgtggaggttgcttatgaagaaattggtttagctttggaaattggagcctctcaccgtaaaccaaaaaatctgtcagtactataatttctggtggagaaccacatctagtgcaactggctgttgctagaagattccgttggaccaatgtggaatcagagggtatacaagtacgtgaatattaagacattgattgattcgaacatcagaatctatgtattctaatcctatattgtatcaacaataatattattattgatcctgtatgtaaaaggttatgagacttaaaatcacacggtttaagataccgaaatgctaagcaaaaacgtttcgtcgagagaaacaaaaacattgagttgcgttataaaatagatagtaacagacttgctaactgagcattcaagactacgttctatttcaggggcttattcaagctgacggagcaacatatggtgcactctcagtacaatctgctctgatgccgcatagttaagccagccccgacacccgccaacacccgctgacgcgccctgacgggcttgtctgctcccggcatccgcttacagacaagctgtgaccgtctccgggagctgcatgtgtcagaggttttcaccgtcatcaccgaaacgcgcgagacgaaagggcctcgtgatacgcctatttttataggttaatgtcatgataataatggtttcttagacgtcaggtggcacttttcggggaaatgtgcgcggaacccctatttgtttatttttctaaatacattcaaatatgtatccgctcatgagacaataaccctgataaatgcttcaataatattgaaaaaggaagagtatgagtattcaacatttccgtgtcgcccttattcccttttttgcggcattttgccttcctgtttttgctcacccagaaacgctggtgaaagtaaaagatgctgaagatcagttgggtgcacgagtgggttacatcgaactggatctcaacagcggtaagatccttgagagttttcgccccgaagaacgttttccaatgatgagcacttttaaagttctgctatgtggcgcggtattatcccgtattgacgccgggcaagagcaactcggtcgccgcatacactattctcagaatgacttggttgagtactcaccagtcacagaaaagcatcttacggatggcatgacagtaagagaattatgcagtgctgccataaccatgagtgataacactgcggccaacttacttctgacaacgatcggaggaccgaaggagctaaccgcttttttgcacaacatgggggatcatgtaactcgccttgatcgttgggaaccggagctgaatgaagccataccaaacgacgagcgtgacaccacgatgcctgtagcaatggcaacaacgttgcgcaaactattaactggcgaactacttactctagcttcccggcaacaattaatagactggatggaggcggataaagttgcaggaccacttctgcgctcggcccttccggctggctggtttattgctgataaatctggagccggtgagcgtgggtctcgcggtatcattgcagcactggggccagatggtaagccctcccgtatcgtagttatctacacgacggggagtcaggcaactatggatgaacgaaatagacagatcgctgagataggtgcctcactgattaagcattggtaactgtcagaccaagtttactcatatatactttagattgatttaaaacttcatttttaatttaaaaggatctaggtgaagatcctttttgataatctcatgaccaaaatcccttaacgtgagttttcgttccactgagcgtcagaccccgtagaaaagatcaaaggatcttcttgagatcctttttttctgcgcgtaatctgctgcttgcaaacaaaaaaaccaccgctaccagcggtggtttgtttgccggatcaagagctaccaactctttttccgaaggtaactggcttcagcagagcgcagataccaaatactgttcttctagtgtagccgtagttaggccaccacttcaagaactctgtagcaccgcctacatacctcgctctgctaatcctgttaccagtggctgctgccagtggcgataagtcgtgtcttaccgggttggactcaagacgatagttaccggataaggcgcagcggtcgggctgaacggggggttcgtgcacacagcccagcttggagcgaacgacctacaccgaact

**eDA110:**

gagatacctacagcgtgagctatgagaaagcgccacgcttcccgaagggagaaaggcggacaggtatccggtaagcggcagggtcggaacaggagagcgcacgagggagcttccagggggaaacgcctggtatctttatagtcctgtcgggtttcgccacctctgacttgagcgtcgatttttgtgatgctcgtcaggggggcggagcctatggaaaaacgccagcaacgcggcctttttacggttcctggccttttgctggccttttgctcacatgttctttcctgcgttatcccctgattctgtggataaccgtattaccgcctttgagtgagctgataccgctcgccgcagccgaacgaccgagcgcagcgagtcagtgagcgaggaagcggaagagcgcccaatacgcaaaccgcctctccccgcgcgttggccgattcattaatgcagttgtcattttgtttcttctcgtacgagcttgctcctgatcagcctatctcgcagctgtgttaaaccgtctttaagtcaacccacagtctactgcaatcgtattcagaactagccactagactgttacaagtcgaaacctcagttaaccaactagaaactctagttaccaagttagaactgtgagtacgaaaagtctgaaaagcagaaagattcaatagatttgtctgtgttacacaagagttcaacaagtagctgcgttcgtgctagttgtctaggtttaaacgtttaaaaagacaactagtagtttacttcacctctgtattactgacggtttgtgcctcaacagtttacgttaacaaactagtaagcgtctacttcgtggacttgtacttttagtagaaagattgagtgtctggcagttttaaacctaagtcagttagaatctcaatcctctgcctttggtttagaaacagtgtagctgttttgacctagactgagtacaactgttcttgctgcttattcgaaacttgcctaactcactgcaacagacaagtctagtttagaataaagtaggctcagttattcaagtctaacttcagacagtgtttctcagatttgacttaagtagacagtagtgtgtgaaaaaccagactagtttgaatcttagcctgttcagaaaaaactaactgacttgtgtcaggcctcccaaaaaaactaacttcaatcctctgtactattccacaactattagttgttgcaagtctgaagcctagtcttgctttgaccttacaggatagaggttggctcacaactgagtctgtgtgaaaacaaaagtaaagattgccacaagaagtctattctagttagtccttttacgttggatagcaatagtccaatcttagtttgcagctgatgaatatcttgtggtaggggtttgggaaaatcattcgagtttgatgtttttcttggtatttcccactcctcttcagagtacagaagattaagtgagaccttcgtttgtgcggatcccccacacaccatagcttcaaaatgtttctactccttttttactcttccagattttctcggactccgcgcatcgccgtaccacttcaaaacacccaagcacagcatactaaatttcccctctttcttcctctagggtgtcgttaattacccgtactaaaggtttggaaaagaaaaaagagaccgcctcgtttctttttcttcgtcgaaaaaggcaataaaaatttttatcacgtttctttttcttgaaaatttttttttttgatttttttctctttcgatgacctcccattgatatttaagttaataaacggtcttcaatttctcaagtttcagtttcatttttcttgttctattacaactttttttacttcttgctcattagaaagaaagcatagcaatctaatctaagggcggtgttgacaattaatcatcggcatagtatatcggcatagtataatacgacaaggtgaggaactaaaccatggccaagttgaccagtgccgttccggtgctcaccgcgcgcgacgtcgccggagcggtcgagttctggaccgaccggctcgggttctcccgggacttcgtggaggacgacttcgccggtgtggtccgggacgacgtgaccctgttcatcagcgcggtccaggaccaggtggtgccggacaacaccctggcctgggtgtgggtgcgcggcctggacgagctgtacgccgagtggtcggaggtcgtgtccacgaacttccgggacgcctccgggccggccatgaccgagatcggcgagcagccgtgggggcgggagttcgccctgcgcgacccggccggcaactgcgtgcacttcgtggccgaggagcaggactgacacgtccgacgcggcccgacgggtccgaggcctcggagatccgtcccccttttcctttgtcgatatcatgtaattagttatgtcacgcttacattcacgccctccccccacatccgctctaaccgaaaaggaaggagttagacaacctgaagtctaggtccctatttatttttttatagttatgttagtattaagaacgttatttatatttcaaatttttcttttttttctgtacagacgcgtgtacgcatgtaacattatactgaaaaccttgcttgagaaggttttgggacgctcgaaggctttaatttgcaagctggagaccaacatgtgagcaaaaggccagcaaaaggccaggaaccgtaaaaaggccgcgaattgtgagcggataacaatttcacacaggaaacagctatgaccatgattacgccaagcttgcatgctgatcggcacgtaagaggttccaactttcaccataatgaaataagatcactaccgggcgtattttttgagttatcgagattttcaggagctaaggaagctaaaatgagccatattcaacgggaaacgtcttgctcgaggccgcgattaaattccaacatggatgctgatttatatgggtataaatgggctcgcgataatgtcgggcaatcaggtgcgacaatctatcgattgtatgggaagcccgatgcgccagagttgtttctgaaacatggcaaaggtagcgttgccaatgatgttacagatgagatggtcaggctaaactggctgacggaatttatgcctcttccgaccatcaagcattttatccgtactcctgatgatgcatggttactcaccactgcgatcccagggaaaacagcattccaggtattagaagaatatcctgattcaggtgaaaatattgttgatgcgctggcagtgttcctgcgccggttgcattcgattcctgtttgtaattgtccttttaacggcgatcgcgtatttcgtctcgctcaggcgcaatcacgaatgaataacggtttggttggtgcgagtgattttgatgacgagcgtaatggctggcctgttgaacaagtctggaaagaaatgcataagcttttgccattctcaccggattcagtcgtcactcatggtgatttctcacttgataaccttatttttgacgaggggaaattaataggttgtattgatgttggacgagtcggaatcgcagaccgataccaggatcttgccatcctatggaactgcctcggtgagttttctccttcattacagaaacggctttttcaaaaatatggtattgataatcctgatatgaataaattgcagtttcacttgatgctcgatgagtttttctaatgagggcccaaatgtaatcacctggctcaccttcgggtgggcctttctgcggcatgcagcaccaccaaaaaaaaaacgaaaagtagaagacccacgatttatgtacccatacgatgttcctgactatgcgggtatgaaaaacatcaaaaaaaaccaggtaatgaacctgggtccgaactctaaactgctgaaagaatacaaatcccagctgatcgaactgaacatcgaacagttcgaagcaggtatcggtctgatcctgggtgatgcttacatccgttctcgtgatgaaggtaaaacctactgtatgcagttcgagtggaaaaacaaagcatacatggaccacgtatgtctgctgtacgatcagtgggtactgtccccgccgcacaaaaaagaacgtgttaaccacctgggtaacctggtaatcacctggggcgcccagactttcaaacaccaagctttcaacaaactggctaacctgttcatcgttaacaacaaaaaaaccatcccgaacaacctggttgaaaactacctgaccccgatgtctctggcatactggttcatggatgatggtggtaaatgggattacaacaaaaactctaccaacaaatcgatcgtactgaacacccagtctttcactttcgaagaagtagaatacctggttaagggtctgcgtaacaaattccaactgaactgttacgtaaaaatcaacaaaaacaaaccgatcatctacatcgattctatgtcttacctgatcttctacaacctgatcaaaccgtacctgatcccgcagatgatgtacaaactgccgaacactatctcctccgaaactttcctgaaataacggcggccgccagcttgggcccgaacaaaaactcatctcagaagaggatctgaatagcgccgtcgaccatcatcatcatcatcattgagttttagccttagacatgactgttcctcagttcaagttgggcacttacgagaagaccggtcttgctagattctaatcaagaggatgtcagaatgccatttgcctgagagatgcaggcttcatttttgatacttttttatttgtaacctatatagtataggattttttttgtcattttgtttcttctcgtacgagcttgctcctgatcagcctatctcgcagctgatgaatatcttgtggtaggggtttgggaaaatcattcgagtttgatgtttttcttggtatttcccactcctcttcagagtacagaagattaagtgagccaaccgtgtaggtctacaaactggtattctcgtttgttgtgaatcttgtcttttctgcttactttctaatagattgagtgagttaggtctgtatttgacggctaaagttaaacttaactgttgtgtgggatagttgttagctgtcaagaaaccgttaaacgtagaaagagtaacttttctaaaccgtagtgtctactcaatattgagacaagcagctagtttaagagcagaagctgaagtctgtgagtttggaaactaaaaggtctgtgaacagctaactagaactaggcttgcttttagtattagaataacaacaaacgaggtctattagttgtgggatctagctagttgcctaagaacaaaggactaagtctgcagaaaaacgagtttcaataggtctgtgtgtaacagtgttaaggctactagcagaactgccacctaagtttcgtaagttccacctcagccttcaatattccagacttgaccaactttcagtttttaaactctaccgtactagtagtttctgcacagtattagacctacttacactgtctgtgcttgcaccacaaaccaaatagtagaatctcttttaagtagaacagtcttttcttgttagactaaaacaacgagagacaactgagtgtagggttacagttaaaaaactgtacagtctgctgtacagacgttacacggcagtcctgttccaaacttcacttaagggttagattggctaagttcaacttgtccttttaagttagttcagacctagttagcagacttaggtgggtaagctgagctacaagagttaaaaacaactgctaagactaaagtaagtgtacgtaagcttcaaactagtaagttaaacttagaaacagctaacgtctaaagctagaagtaagctcaatcttagtactgttaacctaagttgctcacgcaccttcgccaactcaacctgcaggtcgactctagaggggattcggcgctccaactgtattctaggcctttccacaatttaaaaaaagacctccgcgacggggaattgaaccccggtctaccgcgcgacaagcggtggttctaccactaaactatcacggatttcgatgctgctttcccacttccttactttacattctctaggcgtttagcctgtagaataaaaccttttttcaagactacaatactgctgctagtaataccgaattattacatgttttaacaacttgagagcgggtcgtatcctgtcttccgattgtacaatctcatcttatcagctcatctcatctcttcggaatcccccccgggtaccgagctcgaattcactggccgtcgttttacaacgtcatgactgggaaaaccctggcgttacccaacttaatcgccttgcagcacatcccgggtaccgtagatcaagtgacttcttgagcttgccaagcatcaagtcccaccgacaagtggtacaccagtcagcgttatctttccatacattctttaattcaaattgtcccattcacaactcaatgtttttgtttctctcgacgaaacgtttttgcttagcatttcggtatcttaaaccgtgtgattttaagtctcataaccttctacatacaggatcaataataatattattgttgatacaatataggattagaatacatagattctgatgttcgaatcaatcaatgtcttaatattcacgtacttgtataccctctgattccacattggtccaacggaatcttctagcaacagccagttgcactagatgtggttctccaccagaaattatagtactgacagattttttggtttacggtgagaggctccaatttccaaagctaaaccaatttcttcataagcaacctccacgcgcgcttttagtatgatgtagaaactggacctaaagatcgaccaatacttgcttacacggcagcgcgaaataaaagacattgataatgtagagtaagtactatttcttcaaaagatagaaccactctcagaggctaactagaagatctaaaacggatgtacagctaatatgtttcgaggcaatcatgttcaggtgcgcctcaagtggctcttaatctattttaaagcataatccacgaaaatttcgatacagagagatacgtcataaactgagaacgatagttagaggcatagcttcggcaatgttcgataaacaagtcaacgagtcagaatttcatgtttctttttttcttgtagtaatatgtaataaccgaatcagataacttgccagtaatgacctgagctctataaattgaaaccgctacaaaaagtaagggatcgtgttaagggcacagagaagaaagtgtaaagaagtaagctagcgttgtacaacaactcaaaagcatggagcccggtgggtactagataataagaacaaggtttcagtgagtttacaagttcataaaacacggaacctggcaacatagcataccagtaaatcgatttgaagaactctattaggctggcagcctctctcaatcaatagtctaaaacggttaagagacacaatcatctctaggtgtcgttgtggtctaccatcgaaacactggaaatcaatccaatgaacttcaagtgatatatttttcaagagtcttcaaagaccaagcaatgttcttattgattgataattttaaagttgaagtctaaatttgtagcagtgaagtctgagcagggggaatgttaagccatatctaatagtacctattagacgcttgattgccgtttgcaaatatgccatgtttctttgaaaccaaaaccacgtaagggggcttaagttttcggcttaggattgttacggagctcaaaccaaataaggcagagagcatacaaaacgtttattaaaaagaacaactacttctggatagctcaattatcttttgtttcttttgaggcgtgcccttcatgacatgacattgcatccatacaattaatagtagataggggaggataagtttgctagtgtgagcatttagagcaaggggtcagtttcctccatctgcctactctgctccatttaaaagcgagccgattgccataagcttgctcacgtagataggacttcaaaaagacgttaagaggctgctctctagaatacgatggaataaacaaatctcgttagtttcttgaaacggaaagtacatgtaggataattgctggatcattcggaagctaaaattggtcctttccccaaagacaatcgactatagttcctgaagcttcctgaggtgagctggatagaaccagaagatagtacctataacgtcaacaacaactgaaactacagatggaagagctaggcatatccataggggatagccatagaaaactttgattgacgcagcaaatcgagacgttaacctgaagctgccataacttcaggaacttgaacacaagagtggcattaaaattcctatgttgcatttcagaaataccacaagtaaactgagcaattaacttgttcataccgacactacaagttaactacaaaccgagacccttatatgcagactgataatacagaatgatacgtatcactcctaagactaggaacatacagagcttttgagcttttggtattttcaagttatttgaaaaattaaatctttatactagggacgaggttcgtgacaaaagaagacaatcatgagactcaccgtcttgtcatctacaagtttcaataagcttccaacttgagtaacctagccatgtgattagtaactttcgaagcatgattcagaacgttctgctctgccgcgtcaaaaagggcagctacttgaagttaaaagaaagatcaaaaattcctgagcattgttcaaacagctataaatcaaaacagaataagaaataaggccatcacttaccaaggaaaaacaaaaaagttcgagagatcaacaatgcacttcagggacgcacctgaacctttacattagtgtaaaaataaattattagattgttcggttcgagcgttagccatatacaaaatgcggccttcaaagtcattggaaaagagctgctttaggacctctccaaaagctaaattgacaaacaggctacctcatatctccatcatctttgccaaccttttcagtaacaaaaaacaaaagcaatatcgtttatttgtccattggacgatcatcaaatattgctaggtaggcacttttttcggtaagcgtgtctgcctcctttcttcagtaagatcaattgtcgccggatactatgaatcgatcattgtagaaactaacatgcttctaattacccattcaatcttgtagcactgagtgcatctcgcgtgagacaaattgaactcgcctacttgggtaggataaacggttgggtaacgtagctgcttcatccggctacttgcttggtctgctgcgtgctttacgcgctgattggtggtcgaataaaccggtacattatttgttcttatccacgacaatgagacgaacaagtaacaaaccagaactggtagctcgatgattacagttccacaatatcagccgaatacccacactttttccaacctgtgtcaattgccttagaaattgtattagcacaaaaatcatttcgctgacggctgaacatccgatatttactattcgtcaaggtatttagaattcttgtgcgaacttgtgggtcgtggtgaaaaagactattgtgagttgttcgtagctgagtccttgactatactgatacacattaaaagaagctgccctttaagcaactaatgtatttgttatgcgtatatctgtagaaagcaacagtgcgacagttgaagtgttcgaacacacttctcaattttgaatctttctcattctcattcagtgaacgatgaaccaagtacacaaatttggaggtgaataattatgctagggagagcttgctagatcatttcacaattagaatgtgaaaccagacctttagatgtaaatgctaaggctaggataggatcgtaattagtaggatcaacaaactttattgattagcctcacctagactaagtttgatcttttcttcatcgtccaatggacaaataaacgatattgcttttgttttttgttactgaaaaggttggcaaagatgatggagatatgaggtagcctgtttgtcaatttagcttttggagaggtcctaaagcagctcttttccaatgactttgaaggccgcattttgtatatggctaacgctcgaaccgaacaatctaataatttatttttacactaatgtaaaggttcaggtgcgtccctgaagtgcattgttgatctctcgaacttttttgtttttccttggtaagtgatggccttatttcttattctgttttgatttatagctgtttgaacaatgctcaggaatttttgatctttcttttaacttcaagtagctgccctttttgacgcggcagagcagaacgttctgaatcatgcttcgaaagttactaatcacatggctaggttactcaagttggaagcttattgaaacttgtagatgacaagacggtgagtctcatgattgtcttcttttgtcacgaacctcgtccctagtataaagatttaatttttcaaataacttgaaaataccaaaagctcaaaagctctgtatgttcctagtcttaggagtgatacgtatcattctgtattatcagtctgcatataagggtctcggtttgtagttaacttgtagtgtcggtatgaacaagttaattgctcagtttacttgtggtatttctgaaatgcaacataggaattttaatgccactcttgtgttcaagttcctgaagttatggcagcttcaggttaacgtctcgatttgctgcgtcaatcaaagttttctatggctatcccctatggatatgcctagctcttccatctgtagtttcagttgttgttgacgttataggtactatcttctggttctatccagctcacctcaggaagcttcaggaactatagtcgattgtctttggggaaaggaccaattttagcttccgaatgatccagcaattatcctacatgtactttccgtttcaagaaactaacgagatttgtttattccatcgtattctagagagcagcctcttaacgtctttttgaagtcctatctacgtgagcaagcttatggcaatcggctcgcttttaaatggagcagagtaggcagatggaggaaactgaccccttgctctaaatgctcacactagcaaacttatcctcccctatctactattaattgtatggatgcaatgtcatgtcatgaagggcacgcctcaaaagaaacaaaagataattgagctatccagaagtagttgttctttttaataaacgttttgtatgctctctgccttatttggtttgagctccgtaacaatcctaagccgaaaacttaagcccccttacgtggttttggtttcaaagaaacatggcatatttgcaaacggcaatcaagcgtctaataggtactattagatatggcttaacattccccctgctcagacttcactgctacaaatttagacttcaactttaaaattatcaatcaataagaacattgcttggtctttgaagactcttgaaaaatatatcacttgaagttcattggattgatttccagtgtttcgatggtagaccacaacgacacctagagatgattgtgtctcttaaccgttttagactattgattgagagaggctgccagcctaatagagttcttcaaatcgatttactggtatgctatgttgccaggttccgtgttttatgaacttgtaaactcactgaaaccttgttcttattatctagtacccaccgggctccatgcttttgagttgttgtacaacgctagcttacttctttacactttcttctctgtgcccttaacacgatcccttactttttgtagcggtttcaatttatagagctcaggtcattactggcaagttatctgattcggttattacatattactacaagaaaaaaaagaaacatgaaattctgactcgttgacttgtttatcgaacattgccgaagctatgcctctaactatcgttctcagtttatgacgtatctctctgtatcgaaattttcgtggattatgctttaaaatagattaagagccacttgaggcgcacctgaacatgattgcctcgaaacatattagctgtacatccgttttagatcttctagttagcctctgagagtggttctatcttttgaagaaatagtacttactctacattatcaatgtcttttatttcgcgctgccgtgtaagcaagtattggtcgatctttaggtccagtttctacatcatactaaaagcgcgcgtggaggttgcttatgaagaaattggtttagctttggaaattggagcctctcaccgtaaaccaaaaaatctgtcagtactataatttctggtggagaaccacatctagtgcaactggctgttgctagaagattccgttggaccaatgtggaatcagagggtatacaagtacgtgaatattaagacattgattgattcgaacatcagaatctatgtattctaatcctatattgtatcaacaataatattattattgatcctgtatgtaaaaggttatgagacttaaaatcacacggtttaagataccgaaatgctaagcaaaaacgtttcgtcgagagaaacaaaaacattgagttgcgttataaaatagatagtaacagacttgctaactgagcattcaagactacgttctatttcaggggcttattcaagctgacggagcaacatatggtgcactctcagtacaatctgctctgatgccgcatagttaagccagccccgacacccgccaacacccgctgacgcgccctgacgggcttgtctgctcccggcatccgcttacagacaagctgtgaccgtctccgggagctgcatgtgtcagaggttttcaccgtcatcaccgaaacgcgcgagacgaaagggcctcgtgatacgcctatttttataggttaatgtcatgataataatggtttcttagacgtcaggtggcacttttcggggaaatgtgcgcggaacccctatttgtttatttttctaaatacattcaaatatgtatccgctcatgagacaataaccctgataaatgcttcaataatattgaaaaaggaagagtatgagtattcaacatttccgtgtcgcccttattcccttttttgcggcattttgccttcctgtttttgctcacccagaaacgctggtgaaagtaaaagatgctgaagatcagttgggtgcacgagtgggttacatcgaactggatctcaacagcggtaagatccttgagagttttcgccccgaagaacgttttccaatgatgagcacttttaaagttctgctatgtggcgcggtattatcccgtattgacgccgggcaagagcaactcggtcgccgcatacactattctcagaatgacttggttgagtactcaccagtcacagaaaagcatcttacggatggcatgacagtaagagaattatgcagtgctgccataaccatgagtgataacactgcggccaacttacttctgacaacgatcggaggaccgaaggagctaaccgcttttttgcacaacatgggggatcatgtaactcgccttgatcgttgggaaccggagctgaatgaagccataccaaacgacgagcgtgacaccacgatgcctgtagcaatggcaacaacgttgcgcaaactattaactggcgaactacttactctagcttcccggcaacaattaatagactggatggaggcggataaagttgcaggaccacttctgcgctcggcccttccggctggctggtttattgctgataaatctggagccggtgagcgtgggtctcgcggtatcattgcagcactggggccagatggtaagccctcccgtatcgtagttatctacacgacggggagtcaggcaactatggatgaacgaaatagacagatcgctgagataggtgcctcactgattaagcattggtaactgtcagaccaagtttactcatatatactttagattgatttaaaacttcatttttaatttaaaaggatctaggtgaagatcctttttgataatctcatgaccaaaatcccttaacgtgagttttcgttccactgagcgtcagaccccgtagaaaagatcaaaggatcttcttgagatcctttttttctgcgcgtaatctgctgcttgcaaacaaaaaaaccaccgctaccagcggtggtttgtttgccggatcaagagctaccaactctttttccgaaggtaactggcttcagcagagcgcagataccaaatactgttcttctagtgtagccgtagttaggccaccacttcaagaactctgtagcaccgcctacatacctcgctctgctaatcctgttaccagtggctgctgccagtggcgataagtcgtgtcttaccgggttggactcaagacgatagttaccggataaggcgcagcggtcgggctgaacggggggttcgtgcacacagcccagcttggagcgaacgacctacaccgaact

**eDA137:**

cagggtaatgcatccagcatccagcatccagcatccagcatccagcatccgcatccagcatccagcatccagcatccagcatccagcatccagcatccagcatccagcatccagcatccagcatccagcatccagcatccagcatccagcatccagcatccacgatcggaggaccgaaggagctaaccgcttttttgcacaacatgggggatcatgtaactcgccttgatcgttgggaaccggagctgaatgaagccataccaaacgacgagcgtgacaccacgatgcctgtagcaatggcaacaacgttgcgcaaactattaactggcgaactacttactctagcttcccggcaacaattaatagactggatggaggcggataaagttgcaggaccacttctgcgctcggcccttccggctggctggtttattgctgataaatctggagccggtgagcgtgggtctcgcggtatcattgcagcactggggccagatggtaagccctcccgtatcgtagttatctacacgacggggagtcaggcaactatggatgaacgaaatagacagatcgctgagataggtgcctcactgattaagcattggtaactgtcagaccaagtttactcatatatactttagattgatttaaaacttcatttttaatttaaaaggatctaggtgaagatcctttttgataatctcatgaccaaaatcccttaacgtgagttttcgttccactgagcgtcagaccccgtagaaaagatcaaaggatcttcttgagatcctttttttctgcgcgtaatctgctgcttgcaaacaaaaaaaccaccgctaccagcggtggtttgtttgccggatcaagagctaccaactctttttccgaaggtaactggcttcagcagagcgcagataccaaatactgttcttctagtgtagccgtagttaggccaccacttcaagaactctgtagcaccgcctacatacctcgctctgctaatcctgttaccagtggctgctgccagtggcgataagtcgtgtcttaccgggttggactcaagacgatagttaccggataaggcgcagcggtcgggctgaacggggggttcgtgcacacagcccagcttggagcgaacgacctacaccgaactgagatacctacagcgtgagctatgagaaagcgccacgcttcccgaagggagaaaggcggacaggtatccggtaagcggcagggtcggaacaggagagcgcacgagggagcttccagggggaaacgcctggtatctttatagtcctgtcgggtttcgccacctctgacttgagcgtcgatttttgtgatgctcgtcaggggggcggagcctatggaaaaacgccagcaacgcggcctttttacggttcctggccttttgctggccttttgctcacatgttctttcctgcgttatcccctgattctgtggataaccgtattaccgcctttgagtgagctgataccgctcgccgcagccgaacgaccgagcgcagcgagtcagtgagcgaggaagcggaagagcgcccaatacgcaaaccgcctctccccgcgcgttggccgattcattaatgcagttgtcattttgtttcttctcgtacgagcttgctcctgatcagcctatctcgcagctgtgttaaaccgtctttaagtcaacccacagtctactgcaatcgtattcagaactagccactagactgttacaagtcgaaacctcagttaaccaactagaaactctagttaccaagttagaactgtgagtacgaaaagtctgaaaagcagaaagattcaatagatttgtctgtgttacacaagagttcaacaagtagctgcgttcgtgctagttgtctaggtttaaacgtttaaaaagacaactagtagtttacttcacctctgtattactgacggtttgtgcctcaacagtttacgttaacaaactagtaagcgtctacttcgtggacttgtacttttagtagaaagattgagtgtctggcagttttaaacctaagtcagttagaatctcaatcctctgcctttggtttagaaacagtgtagctgttttgacctagactgagtacaactgttcttgctgcttattcgaaacttgcctaactcactgcaacagacaagtctagtttagaataaagtaggctcagttattcaagtctaacttcagacagtgtttctcagatttgacttaagtagacagtagtgtgtgaaaaaccagactagtttgaatcttagcctgttcagaaaaaactaactgacttgtgtcaggcctcccaaaaaaactaacttcaatcctctgtactattccacaactattagttgttgcaagtctgaagcctagtcttgctttgaccttacaggatagaggttggctcacaactgagtctgtgtgaaaacaaaagtaaagattgccacaagaagtctattctagttagtccttttacgttggatagcaatagtccaatcttagtttgcagctgatgaatatcttgtggtaggggtttgggaaaatcattcgagtttgatgtttttcttggtatttcccactcctcttcagagtacagaagattaagtgagaccttcgtttgtgcggatcccccacacaccatagcttcaaaatgtttctactccttttttactcttccagattttctcggactccgcgcatcgccgtaccacttcaaaacacccaagcacagcatactaaatttcccctctttcttcctctagggtgtcgttaattacccgtactaaaggtttggaaaagaaaaaagagaccgcctcgtttctttttcttcgtcgaaaaaggcaataaaaatttttatcacgtttctttttcttgaaaatttttttttttgatttttttctctttcgatgacctcccattgatatttaagttaataaacggtcttcaatttctcaagtttcagtttcatttttcttgttctattacaactttttttacttcttgctcattagaaagaaagcatagcaatctaatctaagggcggtgttgacaattaatcatcggcatagtatatcggcatagtataatacgacaaggtgaggaactaaaccatggccaagttgaccagtgccgttccggtgctcaccgcgcgcgacgtcgccggagcggtcgagttctggaccgaccggctcgggttctcccgggacttcgtggaggacgacttcgccggtgtggtccgggacgacgtgaccctgttcatcagcgcggtccaggaccaggtggtgccggacaacaccctggcctgggtgtgggtgcgcggcctggacgagctgtacgccgagtggtcggaggtcgtgtccacgaacttccgggacgcctccgggccggccatgaccgagatcggcgagcagccgtgggggcgggagttcgccctgcgcgacccggccggcaactgcgtgcacttcgtggccgaggagcaggactgacacgtccgacgcggcccgacgggtccgaggcctcggagatccgtcccccttttcctttgtcgatatcatgtaattagttatgtcacgcttacattcacgccctccccccacatccgctctaaccgaaaaggaaggagttagacaacctgaagtctaggtccctatttatttttttatagttatgttagtattaagaacgttatttatatttcaaatttttcttttttttctgtacagacgcgtgtacgcatgtaacattatactgaaaaccttgcttgagaaggttttgggacgctcgaaggctttaatttgcaagctggagaccaacatgtgagcaaaaggccagcaaaaggccaggaaccgtaaaaaggccgcgaattgtgagcggataacaatttcacacaggaaacagctatgaccatgattacgccaagcttgcatgccgcagaaaggcccacccgaaggtgagccaggtgattacatttgggccctcattagaaaaactcatcgagcatcaagtgaaactgcaatttattcatatcaggattatcaataccatatttttgaaaaagccgtttctgtaatgaaggagaaaactcaccgaggcagttccataggatggcaagatcctggtatcggtctgcgattccgactcgtccaacatcaatacaacctattaatttcccctcgtcaaaaataaggttatcaagtgagaaatcaccatgagtgacgactgaatccggtgagaatggcaaaagcttatgcatttctttccagacttgttcaacaggccagccattacgctcgtcatcaaaatcactcgcaccaaccaaaccgttattcattcgtgattgcgcctgagcgagacgaaatacgcgatcgccgttaaaaggacaattacaaacaggaatcgaatgcaaccggcgcaggaacactgccagcgcatcaacaatattttcacctgaatcaggatattcttctaatacctggaatgctgttttccctgggatcgcagtggtgagtaaccatgcatcatcaggagtacggataaaatgcttgatggtcggaagaggcataaattccgtcagccagtttagcctgaccatctcatctgtaacatcattggcaacgctacctttgccatgtttcagaaacaactctggcgcatcgggcttcccatacaatcgatagattgtcgcacctgattgcccgacattatcgcgagcccatttatacccatataaatcagcatccatgttggaatttaatcgcggcctcgagcaagacgtttcccgttgaatatggctcattttagcttccttagctcctgaaaatctcgataactcaaaaaatacgcccggtagtgatcttatttcattatggtgaaagttggaacctcttacgtgccgatcagcatgcagcaccaccaaaaaaaaaacgaaaagtagaagacccacgatttatgtacccatacgatgttcctgactatgcgggtatgaaaaacatcaaaaaaaaccaggtaatgaacctgggtccgaactctaaactgctgaaagaatacaaatcccagctgatcgaactgaacatcgaacagttcgaagcaggtatcggtctgatcctgggtgatgcttacatccgttctcgtgatgaaggtaaaacctactgtatgcagttcgagtggaaaaacaaagcatacatggaccacgtatgtctgctgtacgatcagtgggtactgtccccgccgcacaaaaaagaacgtgttaaccacctgggtaacctggtaatcacctggggcgcccagactttcaaacaccaagctttcaacaaactggctaacctgttcatcgttaacaacaaaaaaaccatcccgaacaacctggttgaaaactacctgaccccgatgtctctggcatactggttcatggatgatggtggtaaatgggattacaacaaaaactctaccaacaaatcgatcgtactgaacacccagtctttcactttcgaagaagtagaatacctggttaagggtctgcgtaacaaattccaactgaactgttacgtaaaaatcaacaaaaacaaaccgatcatctacatcgattctatgtcttacctgatcttctacaacctgatcaaaccgtacctgatcccgcagatgatgtacaaactgccgaacactatctcctccgaaactttcctgaaataacggcggccgccagcttgggcccgaacaaaaactcatctcagaagaggatctgaatagcgccgtcgaccatcatcatcatcatcattgagttttagccttagacatgactgttcctcagttcaagttgggcacttacgagaagaccggtcttgctagattctaatcaagaggatgtcagaatgccatttgcctgagagatgcaggcttcatttttgatacttttttatttgtaacctatatagtataggattttttttgtcattttgtttcttctcgtacgagcttgctcctgatcagcctatctcgcagctgatgaatatcttgtggtaggggtttgggaaaatcattcgagtttgatgtttttcttggtatttcccactcctcttcagagtacagaagattaagtgagccaaccgtgtaggtctacaaactggtattctcgtttgttgtgaatcttgtcttttctgcttactttctaatagattgagtgagttaggtctgtatttgacggctaaagttaaacttaactgttgtgtgggatagttgttagctgtcaagaaaccgttaaacgtagaaagagtaacttttctaaaccgtagtgtctactcaatattgagacaagcagctagtttaagagcagaagctgaagtctgtgagtttggaaactaaaaggtctgtgaacagctaactagaactaggcttgcttttagtattagaataacaacaaacgaggtctattagttgtgggatctagctagttgcctaagaacaaaggactaagtctgcagaaaaacgagtttcaataggtctgtgtgtaacagtgttaaggctactagcagaactgccacctaagtttcgtaagttccacctcagccttcaatattccagacttgaccaactttcagtttttaaactctaccgtactagtagtttctgcacagtattagacctacttacactgtctgtgcttgcaccacaaaccaaatagtagaatctcttttaagtagaacagtcttttcttgttagactaaaacaacgagagacaactgagtgtagggttacagttaaaaaactgtacagtctgctgtacagacgttacacggcagtcctgttccaaacttcacttaagggttagattggctaagttcaacttgtccttttaagttagttcagacctagttagcagacttaggtgggtaagctgagctacaagagttaaaaacaactgctaagactaaagtaagtgtacgtaagcttcaaactagtaagttaaacttagaaacagctaacgtctaaagctagaagtaagctcaatcttagtactgttaacctaagttgctcacgcaccttcgccaactcaacctgcaggtcgactctagaggggattcggcgctccaactgtattctaggcctttccacaatttaaaaaaagacctccgcgacggggaattgaaccccggtctaccgcgcgacaagcggtggttctaccactaaactatcacggatttcgatgctgctttcccacttccttactttacattctctaggcgtttagcctgtagaataaaaccttttttcaagactacaatactgctgctagtaataccgaattattacatgttttaacaacttgagagcgggtcgtatcctgtcttccgattgtacaatctcatcttatcagctcatctcatctcttcggaatcccccccgggtaccgagctcgaattcactggccgtcgttttacaacgtcatgactgggaaaaccctggcgttacccaacttaatcgccttgcagcacatcccgggtaccgtagatcaagtgacttcttgagcttgccaagcatcaagtcccaccgacaagtggtacaccagtcagcgttatctttccatacattctttaattcaaattgtcccattcacaactcaatgtttttgtttctctcgacgaaacgtttttgcttagcatttcggtatcttaaaccgtgtgattttaagtctcataaccttctacatacaggatcaataataatattattgttgatacaatataggattagaatacatagattctgatgttcgaatcaatcaatgtcttaatattcacgtacttgtataccctctgattccacattggtccaacggaatcttctagcaacagccagttgcactagatgtggttctccaccagaaattatagtactgacagattttttggtttacggtgagaggctccaatttccaaagctaaaccaatttcttcataagcaacctccacgcgcgcttttagtatgatgtagaaactggacctaaagatcgaccaatacttgcttacacggcagcgcgaaataaaagacattgataatgtagagtaagtactatttcttcaaaagatagaaccactctcagaggctaactagaagatctaaaacggatgtacagctaatatgtttcgaggcaatcatgttcaggtgcgcctcaagtggctcttaatctattttaaagcataatccacgaaaatttcgatacagagagatacgtcataaactgagaacgatagttagaggcatagcttcggcaatgttcgataaacaagtcaacgagtcagaatttcatgtttctttttttcttgtagtaatatgtaataaccgaatcagataacttgccagtaatgacctgagctctataaattgaaaccgctacaaaaagtaagggatcgtgttaagggcacagagaagaaagtgtaaagaagtaagctagcgttgtacaacaactcaaaagcatggagcccggtgggtactagataataagaacaaggtttcagtgagtttacaagttcataaaacacggaacctggcaacatagcataccagtaaatcgatttgaagaactctattaggctggcagcctctctcaatcaatagtctaaaacggttaagagacacaatcatctctaggtgtcgttgtggtctaccatcgaaacactggaaatcaatccaatgaacttcaagtgatatatttttcaagagtcttcaaagaccaagcaatgttcttattgattgataattttaaagttgaagtctaaatttgtagcagtgaagtctgagcagggggaatgttaagccatatctaatagtacctattagacgcttgattgccgtttgcaaatatgccatgtttctttgaaaccaaaaccacgtaagggggcttaagttttcggcttaggattgttacggagctcaaaccaaataaggcagagagcatacaaaacgtttattaaaaagaacaactacttctggatagctcaattatcttttgtttcttttgaggcgtgcccttcatgacatgacattgcatccatacaattaatagtagataggggaggataagtttgctagtgtgagcatttagagcaaggggtcagtttcctccatctgcctactctgctccatttaaaagcgagccgattgccataagcttgctcacgtagataggacttcaaaaagacgttaagaggctgctctctagaatacgatggaataaacaaatctcgttagtttcttgaaacggaaagtacatgtaggataattgctggatcattcggaagctaaaattggtcctttccccaaagacaatcgactatagttcctgaagcttcctgaggtgagctggatagaaccagaagatagtacctataacgtcaacaacaactgaaactacagatggaagagctaggcatatccataggggatagccatagaaaactttgattgacgcagcaaatcgagacgttaacctgaagctgccataacttcaggaacttgaacacaagagtggcattaaaattcctatgttgcatttcagaaataccacaagtaaactgagcaattaacttgttcataccgacactacaagttaactacaaaccgagacccttatatgcagactgataatacagaatgatacgtatcactcctaagactaggaacatacagagcttttgagcttttggtattttcaagttatttgaaaaattaaatctttatactagggacgaggttcgtgacaaaagaagacaatcatgagactcaccgtcttgtcatctacaagtttcaataagcttccaacttgagtaacctagccatgtgattagtaactttcgaagcatgattcagaacgttctgctctgccgcgtcaaaaagggcagctacttgaagttaaaagaaagatcaaaaattcctgagcattgttcaaacagctataaatcaaaacagaataagaaataaggccatcacttaccaaggaaaaacaaaaaagttcgagagatcaacaatgcacttcagggacgcacctgaacctttacattagtgtaaaaataaattattagattgttcggttcgagcgttagccatatacaaaatgcggccttcaaagtcattggaaaagagctgctttaggacctctccaaaagctaaattgacaaacaggctacctcatatctccatcatctttgccaaccttttcagtaacaaaaaacaaaagcaatatcgtttatttgtccattggacgatcatcaaatattgctaggtaggcacttttttcggtaagcgtgtctgcctcctttcttcagtaagatcaattgtcgccggatactatgaatcgatcattgtagaaactaacatgcttctaattacccattcaatcttgtagcactgagtgcatctcgcgtgagacaaattgaactcgcctacttgggtaggataaacggttgggtaacgtagctgcttcatccggctacttgcttggtctgctgcgtgctttacgcgctgattggtggtcgaataaaccggtacattatttgttcttatccacgacaatgagacgaacaagtaacaaaccagaactggtagctcgatgattacagttccacaatatcagccgaatacccacactttttccaacctgtgtcaattgccttagaaattgtattagcacaaaaatcatttcgctgacggctgaacatccgatatttactattcgtcaaggtatttagaattcttgtgcgaacttgtgggtcgtggtgaaaaagactattgtgagttgttcgtagctgagtccttgactatactgatacacattaaaagaagctgccctttaagcaactaatgtatttgttatgcgtatatctgtagaaagcaacagtgcgacagttgaagtgttcgaacacacttctcaattttgaatctttctcattctcattcagtgaacgatgaaccaagtacacaaatttggaggtgaataattatgctagggagagcttgctagatcatttcacaattagaatgtgaaaccagacctttagatgtaaatgctaaggctaggataggatcgtaattagtaggatcaacaaactttattgattagcctcacctagactaagtttgatcttttcttcatcgtccaatggacaaataaacgatattgcttttgttttttgttactgaaaaggttggcaaagatgatggagatatgaggtagcctgtttgtcaatttagcttttggagaggtcctaaagcagctcttttccaatgactttgaaggccgcattttgtatatggctaacgctcgaaccgaacaatctaataatttatttttacactaatgtaaaggttcaggtgcgtccctgaagtgcattgttgatctctcgaacttttttgtttttccttggtaagtgatggccttatttcttattctgttttgatttatagctgtttgaacaatgctcaggaatttttgatctttcttttaacttcaagtagctgccctttttgacgcggcagagcagaacgttctgaatcatgcttcgaaagttactaatcacatggctaggttactcaagttggaagcttattgaaacttgtagatgacaagacggtgagtctcatgattgtcttcttttgtcacgaacctcgtccctagtataaagatttaatttttcaaataacttgaaaataccaaaagctcaaaagctctgtatgttcctagtcttaggagtgatacgtatcattctgtattatcagtctgcatataagggtctcggtttgtagttaacttgtagtgtcggtatgaacaagttaattgctcagtttacttgtggtatttctgaaatgcaacataggaattttaatgccactcttgtgttcaagttcctgaagttatggcagcttcaggttaacgtctcgatttgctgcgtcaatcaaagttttctatggctatcccctatggatatgcctagctcttccatctgtagtttcagttgttgttgacgttataggtactatcttctggttctatccagctcacctcaggaagcttcaggaactatagtcgattgtctttggggaaaggaccaattttagcttccgaatgatccagcaattatcctacatgtactttccgtttcaagaaactaacgagatttgtttattccatcgtattctagagagcagcctcttaacgtctttttgaagtcctatctacgtgagcaagcttatggcaatcggctcgcttttaaatggagcagagtaggcagatggaggaaactgaccccttgctctaaatgctcacactagcaaacttatcctcccctatctactattaattgtatggatgcaatgtcatgtcatgaagggcacgcctcaaaagaaacaaaagataattgagctatccagaagtagttgttctttttaataaacgttttgtatgctctctgccttatttggtttgagctccgtaacaatcctaagccgaaaacttaagcccccttacgtggttttggtttcaaagaaacatggcatatttgcaaacggcaatcaagcgtctaataggtactattagatatggcttaacattccccctgctcagacttcactgctacaaatttagacttcaactttaaaattatcaatcaataagaacattgcttggtctttgaagactcttgaaaaatatatcacttgaagttcattggattgatttccagtgtttcgatggtagaccacaacgacacctagagatgattgtgtctcttaaccgttttagactattgattgagagaggctgccagcctaatagagttcttcaaatcgatttactggtatgctatgttgccaggttccgtgttttatgaacttgtaaactcactgaaaccttgttcttattatctagtacccaccgggctccatgcttttgagttgttgtacaacgctagcttacttctttacactttcttctctgtgcccttaacacgatcccttactttttgtagcggtttcaatttatagagctcaggtcattactggcaagttatctgattcggttattacatattactacaagaaaaaaaagaaacatgaaattctgactcgttgacttgtttatcgaacattgccgaagctatgcctctaactatcgttctcagtttatgacgtatctctctgtatcgaaattttcgtggattatgctttaaaatagattaagagccacttgaggcgcacctgaacatgattgcctcgaaacatattagctgtacatccgttttagatcttctagttagcctctgagagtggttctatcttttgaagaaatagtacttactctacattatcaatgtcttttatttcgcgctgccgtgtaagcaagtattggtcgatctttaggtccagtttctacatcatactaaaagcgcgcgtggaggttgcttatgaagaaattggtttagctttggaaattggagcctctcaccgtaaaccaaaaaatctgtcagtactataatttctggtggagaaccacatctagtgcaactggctgttgctagaagattccgttggaccaatgtggaatcagagggtatacaagtacgtgaatattaagacattgattgattcgaacatcagaatctatgtattctaatcctatattgtatcaacaataatattattattgatcctgtatgtaaaaggttatgagacttaaaatcacacggtttaagataccgaaatgctaagcaaaaacgtttcgtcgagagaaacaaaaacattgagttgcgttataaaatagatagtaacagacttgctaactgagcattcaagactacgttctatttcaggggcttattcaagctgacggagcaacatatggtgcactctcagtacaatctgctctgatgccgcatagttaagccagccccgacacccgccaacacccgctgacgcgccctgacgggcttgtctgctcccggcatccgcttacagacaagctgtgaccgtctccgggagctgcatgtgtcagaggttttcaccgtcatcaccgaaacgcgcgagacgaaagggcctcgtgatacgcctatttttataggttaatgtcatgataataatggtttcttagacgtcaggtggcacttttcggggaaatgtgcgcggaacccctatttgtttatttttctaaatacattcaaatatgtatccgctcatgagacaataaccctgataaatgcttcaataatattgaaaaaggaagagtatgagtattcaacatttccgtgtcgcccttattcccttttttgcggcattttgccttcctgtttttgctcacccagaaacgctggtgaaagtaaaagatgctgaagatcagttgggtgcacgagtgggttacatcgaactggatctcaacagcggtaagatccttgagagttttcgccccgaagaacgttttccaatgatgagcacttttaaagttctgctatgtggcgcggtattatcccgtattgacgccgggcaagagcaactcggtcgccgcatacactattctcagaatgacttggttgagtactcaccagtcacagaaaagcatcttacggatggcatgacagtaagagaattatgcagtgctgccataaccatgagtgataacactgcggccaacttacttctgacaacgatcggaggaccgaaggagctaaccgcttttttgcacaacatgggggatcatgtaactcgccttgatcgttgggaaccggagctgaatgaagccataccaaacgacgagcgtgacaccacgatgcctgtagcaatggcaacaacgttgcctggcgtaatagcgaagaggcccgcaccgatcgtggatgctggatgctggatgctggatgctggatgctggatgctggatgctggatgctggatgctggatgctggatgctggatgctggatgctggatgctggatgctggagcatccagcatccagcatccagcatccagcatccagcatcctgctagggataa

**eDA155:**

ataacagggtaatgcatccagcatccagcatccagcatccagcatccagcatccgcatccagcatccagcatccagcatccagcatccagcatccgcatccagcatccagcatccagcatccagcatccagcatccacgatcggaggaccgaaggagctaaccgcttttttgcacaacatgggggatcatgtaactcgccttgatcgttgggaaccggagctgaatgaagccataccaaacgacgagcgtgacaccacgatgcctgtagcaatggcaacaacgttgcgcaaactattaactggcgaactacttactctagcttcccggcaacaattaatagactggatggaggcggataaagttgcaggaccacttctgcgctcggcccttccggctggctggtttattgctgataaatctggagccggtgagcgtgggtctcgcggtatcattgcagcactggggccagatggtaagccctcccgtatcgtagttatctacacgacggggagtcaggcaactatggatgaacgaaatagacagatcgctgagataggtgcctcactgattaagcattggtaactgtcagaccaagtttactcatatatactttagattgatttaaaacttcatttttaatttaaaaggatctaggtgaagatcctttttgataatctcatgaccaaaatcccttaacgtgagttttcgttccactgagcgtcagaccccgtagaaaagatcaaaggatcttcttgagatcctttttttctgcgcgtaatctgctgcttgcaaacaaaaaaaccaccgctaccagcggtggtttgtttgccggatcaagagctaccaactctttttccgaaggtaactggcttcagcagagcgcagataccaaatactgttcttctagtgtagccgtagttaggccaccacttcaagaactctgtagcaccgcctacatacctcgctctgctaatcctgttaccagtggctgctgccagtggcgataagtcgtgtcttaccgggttggactcaagacgatagttaccggataaggcgcagcggtcgggctgaacggggggttcgtgcacacagcccagcttggagcgaacgacctacaccgaactgagatacctacagcgtgagctatgagaaagcgccacgcttcccgaagggagaaaggcggacaggtatccggtaagcggcagggtcggaacaggagagcgcacgagggagcttccagggggaaacgcctggtatctttatagtcctgtcgggtttcgccacctctgacttgagcgtcgatttttgtgatgctcgtcaggggggcggagcctatggaaaaacgccagcaacgcggcctttttacggttcctggccttttgctggccttttgctcacatgttctttcctgcgttatcccctgattctgtggataaccgtattaccgcctttgagtgagctgataccgctcgccgcagccgaacgaccgagcgcagcgagtcagtgagcgaggaagcggaagagcgcccaatacgcaaaccgcctctccccgcgcgttggccgattcattaatgcagttgtcattttgtttcttctcgtacgagcttgctcctgatcagcctatctcgcagctgtgttaaaccgtctttaagtcaacccacagtctactgcaatcgtattcagaactagccactagactgttacaagtcgaaacctcagttaaccaactagaaactctagttaccaagttagaactgtgagtacgaaaagtctgaaaagcagaaagattcaatagatttgtctgtgttacacaagagttcaacaagtagctgcgttcgtgctagttgtctaggtttaaacgtttaaaaagacaactagtagtttacttcacctctgtattactgacggtttgtgcctcaacagtttacgttaacaaactagtaagcgtctacttcgtggacttgtacttttagtagaaagattgagtgtctggcagttttaaacctaagtcagttagaatctcaatcctctgcctttggtttagaaacagtgtagctgttttgacctagactgagtacaactgttcttgctgcttattcgaaacttgcctaactcactgcaacagacaagtctagtttagaataaagtaggctcagttattcaagtctaacttcagacagtgtttctcagatttgacttaagtagacagtagtgtgtgaaaaaccagactagtttgaatcttagcctgttcagaaaaaactaactgacttgtgtcaggcctcccaaaaaaactaacttcaatcctctgtactattccacaactattagttgttgcaagtctgaagcctagtcttgctttgaccttacaggatagaggttggctcacaactgagtctgtgtgaaaacaaaagtaaagattgccacaagaagtctattctagttagtccttttacgttggatagcaatagtccaatcttagtttgcagctgatgaatatcttgtggtaggggtttgggaaaatcattcgagtttgatgtttttcttggtatttcccactcctcttcagagtacagaagattaagtgagaccttcgtttgtgcggatcccccacacaccatagcttcaaaatgtttctactccttttttactcttccagattttctcggactccgcgcatcgccgtaccacttcaaaacacccaagcacagcatactaaatttcccctctttcttcctctagggtgtcgttaattacccgtactaaaggtttggaaaagaaaaaagagaccgcctcgtttctttttcttcgtcgaaaaaggcaataaaaatttttatcacgtttctttttcttgaaaatttttttttttgatttttttctctttcgatgacctcccattgatatttaagttaataaacggtcttcaatttctcaagtttcagtttcatttttcttgttctattacaactttttttacttcttgctcattagaaagaaagcatagcaatctaatctaagggcggtgttgacaattaatcatcggcatagtatatcggcatagtataatacgacaaggtgaggaactaaaccatggccaagttgaccagtgccgttccggtgctcaccgcgcgcgacgtcgccggagcggtcgagttctggaccgaccggctcgggttctcccgggacttcgtggaggacgacttcgccggtgtggtccgggacgacgtgaccctgttcatcagcgcggtccaggaccaggtggtgccggacaacaccctggcctgggtgtgggtgcgcggcctggacgagctgtacgccgagtggtcggaggtcgtgtccacgaacttccgggacgcctccgggccggccatgaccgagatcggcgagcagccgtgggggcgggagttcgccctgcgcgacccggccggcaactgcgtgcacttcgtggccgaggagcaggactgacacgtccgacgcggcccgacgggtccgaggcctcggagatccgtcccccttttcctttgtcgatatcatgtaattagttatgtcacgcttacattcacgccctccccccacatccgctctaaccgaaaaggaaggagttagacaacctgaagtctaggtccctatttatttttttatagttatgttagtattaagaacgttatttatatttcaaatttttcttttttttctgtacagacgcgtgtacgcatgtaacattatactgaaaaccttgcttgagaaggttttgggacgctcgaaggctttaatttgcaagctggagaccaacatgtgagcaaaaggccagcaaaaggccaggaaccgtaaaaaggccgcgaattgtgagcggataacaatttcacacaggaaacagctatgaccatgattacgccaagcttgcatgccgcagaaaggcccacccgaaggtgagccaggtgattacatttgggccctcattagaaaaactcatcgagcatcaagtgaaactgcaatttattcatatcaggattatcaataccatatttttgaaaaagccgtttctgtaatgaaggagaaaactcaccgaggcagttccataggatggcaagatcctggtatcggtctgcgattccgactcgtccaacatcaatacaacctattaatttcccctcgtcaaaaataaggttatcaagtgagaaatcaccatgagtgacgactgaatccggtgagaatggcaaaagcttatgcatttctttccagacttgttcaacaggccagccattacgctcgtcatcaaaatcactcgcaccaaccaaaccgttattcattcgtgattgcgcctgagcgagacgaaatacgcgatcgccgttaaaaggacaattacaaacaggaatcgaatgcaaccggcgcaggaacactgccagcgcatcaacaatattttcacctgaatcaggatattcttctaatacctggaatgctgttttccctgggatcgcagtggtgagtaaccatgcatcatcaggagtacggataaaatgcttgatggtcggaagaggcataaattccgtcagccagtttagcctgaccatctcatctgtaacatcattggcaacgctacctttgccatgtttcagaaacaactctggcgcatcgggcttcccatacaatcgatagattgtcgcacctgattgcccgacattatcgcgagcccatttatacccatataaatcagcatccatgttggaatttaatcgcggcctcgagcaagacgtttcccgttgaatatggctcattttagcttccttagctcctgaaaatctcgataactcaaaaaatacgcccggtagtgatcttatttcattatggtgaaagttggaacctcttacgtgccgatcagcatgcagcaccaccaaaaaaaaaacgaaaagtagaagacccacgatttatgtacccatacgatgttcctgactatgcgggtatgaaaaacatcaaaaaaaaccaggtaatgaacctgggtccgaactctaaactgctgaaagaatacaaatcccagctgatcgaactgaacatcgaacagttcgaagcaggtatcggtctgatcctgggtgatgcttacatccgttctcgtgatgaaggtaaaacctactgtatgcagttcgagtggaaaaacaaagcatacatggaccacgtatgtctgctgtacgatcagtgggtactgtccccgccgcacaaaaaagaacgtgttaaccacctgggtaacctggtaatcacctggggcgcccagactttcaaacaccaagctttcaacaaactggctaacctgttcatcgttaacaacaaaaaaaccatcccgaacaacctggttgaaaactacctgaccccgatgtctctggcatactggttcatggatgatggtggtaaatgggattacaacaaaaactctaccaacaaatcgatcgtactgaacacccagtctttcactttcgaagaagtagaatacctggttaagggtctgcgtaacaaattccaactgaactgttacgtaaaaatcaacaaaaacaaaccgatcatctacatcgattctatgtcttacctgatcttctacaacctgatcaaaccgtacctgatcccgcagatgatgtacaaactgccgaacactatctcctccgaaactttcctgaaataacggcggccgccagcttgggcccgaacaaaaactcatctcagaagaggatctgaatagcgccgtcgaccatcatcatcatcatcattgagttttagccttagacatgactgttcctcagttcaagttgggcacttacgagaagaccggtcttgctagattctaatcaagaggatgtcagaatgccatttgcctgagagatgcaggcttcatttttgatacttttttatttgtaacctatatagtataggattttttttgtcattttgtttcttctcgtacgagcttgctcctgatcagcctatctcgcagctgatgaatatcttgtggtaggggtttgggaaaatcattcgagtttgatgtttttcttggtatttcccactcctcttcagagtacagaagattaagtgagccaaccgtgtaggtctacaaactggtattctcgtttgttgtgaatcttgtcttttctgcttactttctaatagattgagtgagttaggtctgtatttgacggctaaagttaaacttaactgttgtgtgggatagttgttagctgtcaagaaaccgttaaacgtagaaagagtaacttttctaaaccgtagtgtctactcaatattgagacaagcagctagtttaagagcagaagctgaagtctgtgagtttggaaactaaaaggtctgtgaacagctaactagaactaggcttgcttttagtattagaataacaacaaacgaggtctattagttgtgggatctagctagttgcctaagaacaaaggactaagtctgcagaaaaacgagtttcaataggtctgtgtgtaacagtgttaaggctactagcagaactgccacctaagtttcgtaagttccacctcagccttcaatattccagacttgaccaactttcagtttttaaactctaccgtactagtagtttctgcacagtattagacctacttacactgtctgtgcttgcaccacaaaccaaatagtagaatctcttttaagtagaacagtcttttcttgttagactaaaacaacgagagacaactgagtgtagggttacagttaaaaaactgtacagtctgctgtacagacgttacacggcagtcctgttccaaacttcacttaagggttagattggctaagttcaacttgtccttttaagttagttcagacctagttagcagacttaggtgggtaagctgagctacaagagttaaaaacaactgctaagactaaagtaagtgtacgtaagcttcaaactagtaagttaaacttagaaacagctaacgtctaaagctagaagtaagctcaatcttagtactgttaacctaagttgctcacgcaccttcgccaactcactcgactctagaggggattcggcgctccaactgtattctaggcctttccacaatttaaaaaaagacctccgcgacggggaattgaaccccggtctaccgcgcgacaagcggtggttctaccactaaactatcacggatttcgatgctgctttcccacttccttactttacattctctaggcgtttagcctgtagaataaaaccttttttcaagactacaatactgctgctagtaataccgaattattacatgttttaacaacttgagagcgggtcgtatcctgtcttccgattgtacaatctcatcttatcagctcatctcatctcttcggaatcccccccgggtaccgagctcgaattcactggccgtcgttttacaacgtcgtgactgggaaaaccctggcgttacccaacttaatcgccttgcagcacatcccgggtaccgtagatcaagtgacttcttgagcttgccaagcatcaagtcccaccgacaagtggtacaccagtcagcgttatctttccatacattctttaattcaaattgtcccattcacaactcaatgtttttgtttctctcgacgaaacgtttttgcttagcatttcggtatcttaaaccgtgtgattttaagtctcataaccttctacatacaggatcaataataatattattgttgatacaatataggattagaatacatagattctgatgttcgaatcaatcaatgtcttaatattcacgtacttgtataccctctgattccacattggtccaacggaatcttctagcaacagccagttgcactagatgtggttctccaccagaaattatagtactgacagattttttggtttacggtgagaggctccaatttccaaagctaaaccaatttcttcataagcaacctccacgcgcgcttttagtatgatgtagaaactggacctaaagatcgaccaatacttgcttacacggcagcgcgaaataaaagacattgataatgtagagtaagtactatttcttcaaaagatagaaccactctcagaggctaactagaagatctaaaacggatgtacagctaatatgtttcgaggcaatcatgttcaggtgcgcctcaagtggctcttaatctattttaaagcataatccacgaaaatttcgatacagagagatacgtcataaactgagaacgatagttagaggcatagcttcggcaatgttcgataaacaagtcaacgagtcagaatttcatgtttcttttttttcttgtagtaatatgtaataaccgaatcagataacttgccagtaatgacctgagctctataaattgaaaccgctacaaaaagtaagggatcgtgttaagggcacagagaagaaagtgtaaagaagtaagctagcgttgtacaacaactcaaaagcatggagcccggtgggtactagataataagaacaaggtttcagtgagtttacaagttcataaaacacggaacctggcaacatagcataccagtaaatcgatttgaagaactctattaggctggcagcctctctcaatcaatagtctaaaacggttaagagacacaatcatctctaggtgtcgttgtggtctaccatcgaaacactggaaatcaatccaatgaacttcaagtgatatatttttcaagagtcttcaaagaccaagcaatgttcttattgattgataattttaaagttgaagtctaaatttgtagcagtgaagtctgagcagggggaatgttaagccatatctaatagtacctattagacgcttgattgccgtttgcaaatatgccatgtttctttgaaaccaaaaccacgtaagggggcttaagttttcggcttaggattgttacggagctcaaaccaaataaggcagagagcatacaaaacgtttattaaaaagaacaactacttctggatagctcaattatcttttgtttcttttgaggcgtgcccttcatgacatgacattgcatccatacaattaatagtagataggggaggataagtttgctagtgtgagcatttagagcaaggggtcagtttcctccatctgcctactctgctccatttaaaagcgagccgattgccataagcttgctcacgtagataggacttcaaaaagacgttaagaggctgctctctagaatacgatggaataaacaaatctcgttagtttcttgaaacggaaagtacatgtaggataattgctggatcattcggaagctaaaattggtcctttccccaaagacaatcgactatagttcctgaagcttcctgaggtgagctggatagaaccagaagatagtacctataacgtcaacaacaactgaaactacagatggaagagctaggcatatccataggggatagccatagaaaactttgattgacgcagcaaatcgagacgttaacctgaagctgccataacttcaggaacttgaacacaagagtggcattaaaattcctatgttgcatttcagaaataccacaagtaaactgagcaattaacttgttcataccgacactacaagttaactacaaaccgagacccttatatgcagactgataatacagaatgatacgtatcactcctaagactaggaacatacagagcttttgagcttttggtattttcaagttatttgaaaaattaaatctttatactagggacgaggttcgtgacaaaagaagacaatcatgagactcaccgtcttgtcatctacaagtttcaataagcttccaacttgagtaacctagccatgtgattagtaactttcgaagcatgattcagaacgttctgctctgccgcgtcaaaaagggcagctacttgaagttaaaagaaagatcaaaaattcctgagcattgttcaaacagctataaatcaaaacagaataagaaataaggccatcacttaccaaggaaaaacaaaaaagttcgagagatcaacaatgcacttcagggacgcacctgaacctttacattagtgtaaaaataaattattagattgttcggttcgagcgttagccatatacaaaatgcggccttcaaagtcattggaaaagagctgctttaggacctctccaaaagctaaattgacaaacaggctacctcatatctccatcatctttgccaaccttttcagtaacaaaaaacaaaagcaatatcgtttatttgtccattggacgatcatcaaatattgctaggtaggcacttttttcggtaagcgtgtctgcctcctttcttcagtaagatcaattgtcgccggatactatgaatcgatcattgtagaaactaacatgcttctaattacccattcaatcttgtagcactgagtgcatctcgcgtgagacaaattgaactcgcctacttgggtaggataaacggttgggtaacgtagctgcttcatccggctacttgcttggtctgctgcgtgctttacgcgctgattggtggtcgaataaaccggtacattatttgttcttatccacgacaatgagacgaacaagtaacaaaccagaactggtagctcgatgattacagttccacaatatcagccgaatacccacactttttccaacctgtgtcaattgccttagaaattgtattagcacaaaaatcatttcgctgacggctgaacatccgatatttactattcgtcaaggtatttagaattcttgtgcgaacttgtgggtcgtggtgaaaaagactattgtgagttgttcgtagctgagtccttgactatactgatacacattaaaagaagctgccctttaagcaactaatgtatttgttatgcgtatatctgtagaaagcaacagtgcgacagttgaagtgttcgaacacacttctcaattttgaatctttctcattctcattcagtgaacgatgaaccaagtacacaaatttggaggtgaataattatgctagggagagcttgctagatcatttcacaattagaatgtgaaaccagacctttagatgtaaatgctaaggctaggataggatcgtaattagtaggatcaacaaactttattgattagcctcacctagactaagtttgatcttttcttcatcgtccaatggacaaataaacgatattgcttttgttttttgttactgaaaaggttggcaaagatgatggagatatgaggtagcctgtttgtcaatttagcttttggagaggtcctaaagcagctcttttccaatgactttgaaggccgcattttgtatatggctaacgctcgaaccgaacaatctaataatttatttttacactaatgtaaaggttcaggtgcgtccctgaagtgcattgttgatctctcgaacttttttgtttttccttggtaagtgatggccttatttcttattctgttttgatttatagctgtttgaacaatgctcaggaatttttgatctttcttttaacttcaagtagctgccctttttgacgcggcagagcagaacgttctgaatcatgcttcgaaagttactaatcacatggctaggttactcaagttggaagcttattgaaacttgtagatgacaagacggtgagtctcatgattgtcttcttttgtcacgaacctcgtccctagtataaagatttaatttttcaaataacttgaaaataccaaaagctcaaaagctctgtatgttcctagtcttaggagtgatacgtatcattctgtattatcagtctgcatataagggtctcggtttgtagttaacttgtagtgtcggtatgaacaagttaattgctcagtttacttgtggtatttctgaaatgcaacataggaattttaatgccactcttgtgttcaagttcctgaagttatggcagcttcaggttaacgtctcgatttgctgcgtcaatcaaagttttctatggctatcccctatggatatgcctagctcttccatctgtagtttcagttgttgttgacgttataggtactatcttctggttctatccagctcacctcaggaagcttcaggaactatagtcgattgtctttggggaaaggaccaattttagcttccgaatgatccagcaattatcctacatgtactttccgtttcaagaaactaacgagatttgtttattccatcgtattctagagagcagcctcttaacgtctttttgaagtcctatctacgtgagcaagcttatggcaatcggctcgcttttaaatggagcagagtaggcagatggaggaaactgaccccttgctctaaatgctcacactagcaaacttatcctcccctatctactattaattgtatggatgcaatgtcatgtcatgaagggcacgcctcaaaagaaacaaaagataattgagctatccagaagtagttgttctttttaataaacgttttgtatgctctctgccttatttggtttgagctccgtaacaatcctaagccgaaaacttaagcccccttacgtggttttggtttcaaagaaacatggcatatttgcaaacggcaatcaagcgtctaataggtactattagatatggcttaacattccccctgctcagacttcactgctacaaatttagacttcaactttaaaattatcaatcaataagaacattgcttggtctttgaagactcttgaaaaatatatcacttgaagttcattggattgatttccagtgtttcgatggtagaccacaacgacacctagagatgattgtgtctcttaaccgttttagactattgattgagagaggctgccagcctaatagagttcttcaaatcgatttactggtatgctatgttgccaggttccgtgttttatgaacttgtaaactcactgaaaccttgttcttattatctagtacccaccgggctccatgcttttgagttgttgtacaacgctagcttacttctttacactttcttctctgtgcccttaacacgatcccttactttttgtagcggtttcaatttatagagctcaggtcattactggcaagttatctgattcggttattacatattactacaagaaaaaaaagaaacatgaaattctgactcgttgacttgtttatcgaacattgccgaagctatgcctctaactatcgttctcagtttatgacgtatctctctgtatcgaaattttcgtggattatgctttaaaatagattaagagccacttgaggcgcacctgaacatgattgcctcgaaacatattagctgtacatccgttttagatcttctagttagcctctgagagtggttctatcttttgaagaaatagtacttactctacattatcaatgtcttttatttcgcgctgccgtgtaagcaagtattggtcgatctttaggtccagtttctacatcatactaaaagcgcgcgtggaggttgcttatgaagaaattggtttagctttggaaattggagcctctcaccgtaaaccaaaaaatctgtcagtactataatttctggtggagaaccacatctagtgcaactggctgttgctagaagattccgttggaccaatgtggaatcagagggtatacaagtacgtgaatattaagacattgattgattcgaacatcagaatctatgtattctaatcctatattgtatcaacaataatattattattgatcctgtatgtaaaaggttatgagacttaaaatcacacggtttaagataccgaaatgctaagcaaaaacgtttcgtcgagagaaacaaaaacattgagttgcgttataaaatagatagtaacagacttgctaactgagcattcaagactacgttctatttcaggggcttattcaagctgacggagcacatatgcaacaatactacggaaacgacccataacgctgaaacagccgtggaactcgatggaactttcttgccgatttgttgcccaatgtgctccaaaacggagggaagtctttccgcggacagtctggcttgaacgggctagagatggcagagatcggtttgacttagcactcttaaattcatgctttagggttatgtcgcgagacattaatattgggcaagccagggcagatttggctatattcaggctggtttaggcgatccaaaaaggattatgtaagggattttgactgggatggataaaatagttctgattggggcttaatggggctcaagtggctgtgttttctgattgatattggaacctgctgtcatttcgacaattaatatttacttattttggtcaaccccaaataggttgatttcatacttggttcattcaaaaataagtagtcttttgagatctttcaatattataataaatatactataacagccgacttgtttcattttcgcgaatgttcccccagcttatcggatcccgagctcgaattcgctagccagatcggcttaagacaaccaaacccagaactagtcgaatagtctgtgctaaaaaaccaatcggttaaaactagaatcgatctttaagtttaagattcaggcttgttgttaaacctattgccagaaaggacagttgactttggaacaaactcagctaacttgaaactcaaacaacagttttgttacttgccggcttacctcacttgctcagctagtctagcaagaaaacagtgtttcttgtcaacttagaactaatagcagagttaagcagaatcttgaaggctactctgtatttgaaaaccaattgacttgctttttacttagccttacttgctagtcaatcttcagggcagaagtaacttgaaaagttagaaaccaagtttaaaaagacaacaagaaacttgcacacaggactgagaaagtaactaagatttgaaactgagtagactaactctaccagtggttcacaagggttccaccctactttaacttttgacaagagttcaaagtcgaactttaaactgagcttacctctgtggactgtaaactcttgagttaccaaacttttgagctagcctctcagactgttactggttagctctacacagtcttttactattctaacgtaagcttagactagttcttggaataagtgaatctaagaagttgaggctaaactctcaaatagtgtagactaggcacgaactagaagttagtttcttacgttggtagaaaaaaacagttgagtaggctgaaaaggtcaactaagccaaacaagtttcttgtcagaagtctaactgagtgaatacagttgtattcagaaactgtagctagttcgtcttctcaaaccaacttaccagacttttttaaacttgcagtttttactgtacgagacagtttgttactcttcacagtcagttagacgtaaagtaacttgaataagctgaatactaacgtccacgaggtaggcagactatcctgttactgtcttttttagtctgcagtacaaaaagttaaactgttagcaatcctcggttaggcaactttgagttaacttagaagttaccactgcaagctcaatagattcgccctgcagattctaaaaacaaaagctgagctaaagctgctaactgcagttaggtttagatttgagaatagacttggtaaactgcagtaacttacgagaaagtaactaaaactacagttttcaacttcagttttcgaactgtagttgtcttagtctgagttgaaagactgagttaagtaagttaactcactaggttacgctaaagtgtagctactttggattcaggctcactctgcactagcatatggtgcactctcagtacaatctgctctgatgccgcatagttaagccagccccgacacccgccaacacccgctgacgcgccctgacgggcttgtctgctcccggcatccgcttacagacaagctgtgaccgtctccgggagctgcatgtgtcagaggttttcaccgtcatcaccgaaacgcgcgagacgaaagggcctcgtgatacgcctatttttataggttaatgtcatgataataatggtttcttagacgtcaggtggcacttttcggggaaatgtgcgcggaacccctatttgtttatttttctaaatacattcaaatatgtatccgctcatgagacaataaccctgataaatgcttcaataatattgaaaaaggaagagtatgagtattcaacatttccgtgtcgcccttattcccttttttgcggcattttgccttcctgtttttgctcacccagaaacgctggtgaaagtaaaagatgctgaagatcagttgggtgcacgagtgggttacatcgaactggatctcaacagcggtaagatccttgagagttttcgccccgaagaacgttttccaatgatgagcacttttaaagttctgctatgtggcgcggtattatcccgtattgacgccgggcaagagcaactcggtcgccgcatacactattctcagaatgacttggttgagtactcaccagtcacagaaaagcatcttacggatggcatgacagtaagagaattatgcagtgctgccataaccatgagtgataacactgcggccaacttacttctgacaacgatcggaggaccgaaggagctaaccgcttttttgcacaacatgggggatcatgtaactcgccttgatcgttgggaaccggagctgaatgaagccataccaaacgacgagcgtgacaccacgatgcctgtagcaatggcaacaacgttgcctggcgtaatagcgaagaggcccgcaccgatcgtggatgctggatgctggatgctggatgctggatgctggatgctggatgctggatgctggatgctggatgctggatgctggatgctggatgctggatgctggatgctggatgctggatgctggatgctggatgctggatgctaggg

**eDA197:**

gagatacctacagcgtgagctatgagaaagcgccacgcttcccgaagggagaaaggcggacaggtatccggtaagcggcagggtcggaacaggagagcgcacgagggagcttccagggggaaacgcctggtatctttatagtcctgtcgggtttcgccacctctgacttgagcgtcgatttttgtgatgctcgtcaggggggcggagcctatggaaaaacgccagcaacgcggcctttttacggttcctggccttttgctggccttttgctcacatgttctttcctgcgttatcccctgattctgtggataaccgtattaccgcctttgagtgagctgataccgctcgccgcagccgaacgaccgagcgcagcgagtcagtgagcgaggaagcggaagagcgcccaatacgcaaaccgcctctccccgcgcgttggccgattcattaatgcagctggcacgacaggtttcccgactggaaagcgggcagtgagcgcaacgcaattaatgtgagttagctcactcattaggcaccccaggctttacactttatgcttccggctcgtatgttgtgtggaattgtgagcggataacaatttcacacaggaaacagctatgaccatgattacgccaagcttgcatgcgacttacctcgttttaacttagtcggttagtgaagtttaagtctaagttagttcagacctttattgcagattcgtcttactgtctacttagccctgcgtttgacttgcctattagctactcaacttaaaacagttaaactcacagactttacgattgaaaacaagttgacttgaaccccagtggacgaactccacaagtctagtttggcttagagtttctactttacctctattctaccgcagtaactttcaacctgaagtaagttcaagtgaaacttagagtgaagcttgtgatcgactacaagctcaatacaaacttctagaccttagaactagcttagttgcttaacctagtgaatccaacaacaaagttaccaagaatagacagtacgacaactggttacgagtgtaactaagaatcttacactctagttctgccttgctgcttattcaccttttagcttttgttgtttttagttttagccaaccaaaggtggaggtttaaacccttgaactaagcttaggttcaactgaaagtctgtctgccttacagacttgtacagacaaaagttaaacctctaaagcaaccagtctactagttttcgtctttctacgtaactattaccacgagagtaacaagtttctaacacaagtctgttgagccaaatccaaatctttaaacccttgagttagttgagtttcgacgttagaactagctttttgtagtaagcttattcacaaacctacttaagtttagactgtagttaccgtactgaaaagattgctgacttttctacttttacgtttacaacagctgtgtagcgtgcaacagacgtgagattcaaggctgacgcttagttaaacttgtcgaccacacaccatagcttcaaaatgtttctactccttttttactcttccagattttctcggactccgcgcatcgccgtaccacttcaaaacacccaagcacagcatactaaattttccctctttcttcctctagggtgtcgttaattacccgtactaaaggtttggaaaagaaaaaagagaccgcctcgtttctttttcttcgtcgaaaaaggcaataaaaatttttatcacgtttctttttcttgaaatttttttttttagtttttttctctttcagtgacctccattgatatttaagttaataaacggtcttcaatttctcaagtttcagtttcatttttcttgttctattacaactttttttacttcttgttcattagaaagaaagcatagcaatctaatctaaggggcggtgttgacaattaatcatcggcatagtatatcggcatagtataatacgacaaggtgaggaactaaaccatggtatctaaaggtgaagaactattcacaggtgtggtgcccattttagttgaattggatggtgatgtaaacggccacaagttcagtgtcagtggagaaggtgagggtgatgccacctatggaaaactaacattaaaatttatatgcactactggaaagttgcccgtcccctggccaactctagtcactaccctgacatatggagtacagtgtttctcaagatatcctgaccatatgaagcagcatgacttctttaagtccgctatgcctgaaggatacgtccaggaacgtaccatattcttcaaagatgacggtaactataagactagagcagaggtaaaattcgagggagatacgcttgtcaacagaattgagcttaagggaatcgacttcaaggaggatggtaatattttaggccacaaattagagtacaattataattcccacaacgtgtacatcatggccgacaagcagaaaaacggcatcaaagtcaacttcaagatccgtcacaatattgaagatggcagtgtgcaacttgcagaccattatcaacaaaacacaccaataggcgatggacctgtcttgttgcccgacaatcattatttgtcaactcagtccgctctttccaaggacccaaacgagaaaagggaccatatggtactactagagttcgttactgcagctggcataaccttgggaatggatgagctttacaaataagcatcatgtaattagttatgtcacgcttacattcacgccctccccccacatccgctctaaccgaaaaggaaggagttagacaacctgaagtctaggtccctatttatttttttatagttatgttagtattaagaacgttatttatatttcaaatttttcttttttttctgtacagacgcgtgtacgcatgtaacattatactgaaaaccttgcttgagaaggttttgggacgctcgaaggctgtcgaccgtctcagtagcccacacaccatagcttcaaaatgtttctactccttttttactcttccagattttctcggactccgcgcatcgccgtaccacttcaaaacacccaagcacagcatactaaatttcccctctttcttcctctagggtgtcgttaattacccgtactaaaggtttggaaaagaaaaaagagaccgcctcgtttctttttcttcgtcgaaaaaggcaataaaaatttttatcacgtttctttttcttgaaaatttttttttttgatttttttctctttcgatgacctcccattgatatttaagttaataaacggtcttcaatttctcaagtttcagtttcatttttcttgttctattacaactttttttacttcttgctcattagaaagaaagcatagcaatctaatctaagggcggtgttgacaattaatcatcggcatagtatatcggcatagtataatacgacaaggtgaggaactaaaccatggccaagttgaccagtgccgttccggtgctcaccgcgcgcgacgtcgccggagcggtcgagttctggaccgaccggctcgggttctcccgggacttcgtggaggacgacttcgccggtgtggtccgggacgacgtgaccctgttcatcagcgcggtccaggaccaggtggtgccggacaacaccctggcctgggtgtgggtgcgcggcctggacgagctgtacgccgagtggtcggaggtcgtgtccacgaacttccgggacgcctccgggccggccatgaccgagatcggcgagcagccgtgggggcgggagttcgccctgcgcgacccggccggcaactgcgtgcacttcgtggccgaggagcaggactgacacgtccgacgcggcccgacgggtccgaggcctcggagatccgtcccccttttcctttgtcgatatcatgtaattagttatgtcacgcttacattcacgccctccccccacatccgctctaaccgaaaaggaaggagttagacaacctgaagtctaggtccctatttatttttttatagttatgttagtattaagaacgttatttatatttcaaatttttcttttttttctgtacagacgcgtgtacgcatgtaacattatactgaaaaccttgcttgagaaggttttgggacgctcgaaggctttaatttgcggatccccgtgtaggtctacaaactggtattctcgtttgttgtgaatcttgtcttttctgcttactttctaatagattgagtgagttaggtctgtatttgacggctaaagttaaacttaactgttgtgtgggatagttgttagctgtcaagaaaccgttaaacgtagaaagagtaacttttctaaaccgtagtgtctactcaatattgagacaagcagctagtttaagagcagaagctgaagtctgtgagtttggaaactaaaaggtctgtgaacagctaactagaactaggcttgcttttagtattagaataacaacaaacgaggtctattagttgtgggatctagctagttgcctaagaacaaaggactaagtctgcagaaaaacgagtttcaataggtctgtgtgtaacagtgttaaggctactagcagaactgccacctaagtttcgtaagttccacctcagccttcaatattccagacttgaccaactttcagtttttaaactctaccgtactagtagtttctgcacagtattagacctacttacactgtctgtgcttgcaccacaaaccaaatagtagaatctcttttaagtagaacagtcttttcttgttagactaaaacaacgagagacaactgagtgtagggttacagttaaaaaactgtacagtctgctgtacagacgttacacggcagtcctgttccaaacttcacttaagggttagattggctaagttcaacttgtccttttaagttagttcagacctagttagcagacttaggtgggtaagctgagctacaagagttaaaaacaactgctaagactaaagtaagtgtacgtaagcttcaaactagtaagttaaacttagaaacagctaacgtctaaagctagaagtaagctcaatcttagtactgttaacctaagttgctcacgcaccttcgccaactcactcgagtcctgcagggcgatcgccgcgtgagacgggtaccgagctcgaattcactggccgtcgttttacaacgtcgtgactgggaaaaccctggcgttacccaacttaatcgccttgcagcacatccccctttcgccagctggcgtaatagcgaagaggcccgcaccgatcgcccttcccaacagttgcgcagcctgaatggcgaatggcgcctgatgcggtattttctccttacgcatctgtgcggtatttcacaccgcatatggtgcactctcagtacaatctgctctgatgccgcatagttaagccagccccgacacccgccaacacccgctgacgcgccctgacgggcttgtctgctcccggcatccgcttacagacaagctgtgaccgtctccgggagctgcatgtgtcagaggttttcaccgtcatcaccgaaacgcgcgagacgaaagggcctcgtgatacgcctatttttataggttaatgtcatgataataatggtttcttagacgtcaggtggcacttttcggggaaatgtgcgcggaacccctatttgtttatttttctaaatacattcaaatatgtatccgctcatgagacaataaccctgataaatgcttcaataatattgaaaaaggaagagtatgagtattcaacatttccgtgtcgcccttattcccttttttgcggcattttgccttcctgtttttgctcacccagaaacgctggtgaaagtaaaagatgctgaagatcagttgggtgcacgagtgggttacatcgaactggatctcaacagcggtaagatccttgagagttttcgccccgaagaacgttttccaatgatgagcacttttaaagttctgctatgtggcgcggtattatcccgtattgacgccgggcaagagcaactcggtcgccgcatacactattctcagaatgacttggttgagtactcaccagtcacagaaaagcatcttacggatggcatgacagtaagagaattatgcagtgctgccataaccatgagtgataacactgcggccaacttacttctgacaacgatcggaggaccgaaggagctaaccgcttttttgcacaacatgggggatcatgtaactcgccttgatcgttgggaaccggagctgaatgaagccataccaaacgacgagcgtgacaccacgatgcctgtagcaatggcaacaacgttgcgcaaactattaactggcgaactacttactctagcttcccggcaacaattaatagactggatggaggcggataaagttgcaggaccacttctgcgctcggcccttccggctggctggtttattgctgataaatctggagccggtgagcgtgggtctcgcggtatcattgcagcactggggccagatggtaagccctcccgtatcgtagttatctacacgacggggagtcaggcaactatggatgaacgaaatagacagatcgctgagataggtgcctcactgattaagcattggtaactgtcagaccaagtttactcatatatactttagattgatttaaaacttcatttttaatttaaaaggatctaggtgaagatcctttttgataatctcatgaccaaaatcccttaacgtgagttttcgttccactgagcgtcagaccccgtagaaaagatcaaaggatcttcttgagatcctttttttctgcgcgtaatctgctgcttgcaaacaaaaaaaccaccgctaccagcggtggtttgtttgccggatcaagagctaccaactctttttccgaaggtaactggcttcagcagagcgcagataccaaatactgttcttctagtgtagccgtagttaggccaccacttcaagaactctgtagcaccgcctacatacctcgctctgctaatcctgttaccagtggctgctgccagtggcgataagtcgtgtcttaccgggttggactcaagacgatagttaccggataaggcgcagcggtcgggctgaacggggggttcgtgcacacagcccagcttggagcgaacgacctacaccgaact

**eDA199:**

gagatacctacagcgtgagctatgagaaagcgccacgcttcccgaagggagaaaggcggacaggtatccggtaagcggcagggtcggaacaggagagcgcacgagggagcttccagggggaaacgcctggtatctttatagtcctgtcgggtttcgccacctctgacttgagcgtcgatttttgtgatgctcgtcaggggggcggagcctatggaaaaacgccagcaacgcggcctttttacggttcctggccttttgctggccttttgctcacatgttctttcctgcgttatcccctgattctgtggataaccgtattaccgcctttgagtgagctgataccgctcgccgcagccgaacgaccgagcgcagcgagtcagtgagcgaggaagcggaagagcgcccaatacgcaaaccgcctctccccgcgcgttggccgattcattaatgcagctggcacgacaggtttcccgactggaaagcgggcagtgagcgcaacgcaattaatgtgagttagctcactcattaggcaccccaggctttacactttatgcttccggctcgtatgttgtgtggaattgtgagcggataacaatttcacacaggaaacagctatgaccatgattacgccaagcttgcatgcgacttacctcgttttaacttagtcggttagtgaagtttaagtctaagttagttcagacctttattgcagattcgtcttactgtctacttagccctgcgtttgacttgcctattagctactcaacttaaaacagttaaactcacagactttacgattgaaaacaagttgacttgaaccccagtggacgaactccacaagtctagtttggcttagagtttctactttacctctattctaccgcagtaactttcaacctgaagtaagttcaagtgaaacttagagtgaagcttgtgatcgactacaagctcaatacaaacttctagaccttagaactagcttagttgcttaacctagtgaatccaacaacaaagttaccaagaatagacagtacgacaactggttacgagtgtaactaagaatcttacactctagttctgccttgctgcttattcaccttttagcttttgttgtttttagttttagccaaccaaaggtggaggtttaaacccttgaactaagcttaggttcaactgaaagtctgtctgccttacagacttgtacagacaaaagttaaacctctaaagcaaccagtctactagttttcgtctttctacgtaactattaccacgagagtaacaagtttctaacacaagtctgttgagccaaatccaaatctttaaacccttgagttagttgagtttcgacgttagaactagctttttgtagtaagcttattcacaaacctacttaagtttagactgtagttaccgtactgaaaagattgctgacttttctacttttacgtttacaacagctgtgtagcgtgcaacagacgtgagattcaaggctgacgcttagttaaacttgtcgaccacacaccatagcttcaaaatgtttctactccttttttactcttccagattttctcggactccgcgcatcgccgtaccacttcaaaacacccaagcacagcatactaaattttccctctttcttcctctagggtgtcgttaattacccgtactaaaggtttggaaaagaaaaaagagaccgcctcgtttctttttcttcgtcgaaaaaggcaataaaaatttttatcacgtttctttttcttgaaatttttttttttagtttttttctctttcagtgacctccattgatatttaagttaataaacggtcttcaatttctcaagtttcagtttcatttttcttgttctattacaactttttttacttcttgttcattagaaagaaagcatagcaatctaatctaaggggcggtgttgacaattaatcatcggcatagtatatcggcatagtataatacgacaaggtgaggaactaaaccatggtttcaaagggagaagaagacaacatggctatcataaaggaatttatgagattcaaagtgcacatggagggatcagtaaacggccatgaatttgaaattgagggagagggtgagggtagaccttatgagggaacgcagacggctaaactaaaagtgactaaaggaggtcccctgccctttgcatgggacattctgtctcctcaattcatgtacggaagtaaagcctacgtcaagcacccagccgatattcctgattatttgaagttgtccttccctgagggctttaagtgggaaagggtgatgaacttcgaggacggtggcgtggttaccgtaacacaagacagtagtctacaagacggtgaatttatctacaaagtgaaactacgtggaaccaactttccctctgacggacctgtaatgcagaaaaagacgatgggttgggaggcatcctctgagcgtatgtatcccgaggatggcgctttaaagggtgagattaaacaaagactgaagttgaaggatggtggtcactatgacgcagaggttaaaacgacctacaaggccaagaagccagtacaactgcctggagcatataatgtgaatataaagttagacattacatcccataatgaagactacacaattgttgaacaatatgaacgtgctgagggccgtcatagtacaggcggaatggatgaactatacaaatagcatcatgtaattagttatgtcacgcttacattcacgccctccccccacatccgctctaaccgaaaaggaaggagttagacaacctgaagtctaggtccctatttatttttttatagttatgttagtattaagaacgttatttatatttcaaatttttcttttttttctgtacagacgcgtgtacgcatgtaacattatactgaaaaccttgcttgagaaggttttgggacgctcgaaggctgtcgacggtctcagtaggacatggaggcccagaataccctccttgacagtcttgacgtgcgcagctcaggggcatgatgtgactgtcgcccgtacatttagcccatacatccccatgtataatcatttgcatccatacattttgatggccgcacggcgcgaagcaaaaattacggctcctcgctgcggacctgcgagcagggaaacgctcccctcacagacgcgttgaattgtccccacgccgcgcccctgtagagaaatataaaaggttaggatttgccactgaggttcttctttcatatacttccttttaaaatcttgctaggatacagttctcacatcacatccgaacataaacaaccatgggtaaaaagcctgaactcaccgcgacgtctgtcgagaagtttctgatcgaaaagttcgacagcgtctccgacctgatgcagctctcggagggcgaagaatctcgtgctttcagcttcgatgtaggagggcgtggatatgtcctgcgggtaaatagctgcgccgatggtttctacaaagatcgttatgtttatcggcactttgcatcggccgcgctcccgattccggaagtgcttgacattggggaattcagcgagagcctgacctattgcatctcccgccgtgcacagggtgtcacgttgcaagacctgcctgaaaccgaactgcccgctgttctgcagccggtcgcggaggccatggatgcgattgctgcggccgatcttagccagacgagcgggttcggcccattcggaccgcaaggaatcggtcaatacactacatggcgtgatttcatatgcgcgattgctgatccccatgtgtatcactggcaaactgtgatggacgacaccgtcagtgcgtccgtcgcgcaggctctcgatgagctgatgctttgggccgaggactgccccgaagtccggcacctcgtgcacgcggatttcggctccaacaatgtcctgacggacaatggccgcataacagcggtcattgactggagcgaggcgatgttcggggattcccaatacgaggtcgccaacatcttcttctggaggccgtggttggcttgtatggagcagcagacgcgctacttcgagcggaggcatccggagcttgcaggatcgccgcggctccgggcgtatatgctccgcattggtcttgaccaactctatcagagcttggttgacggcaatttcgatgatgcagcttgggcgcagggtcgatgcgacgcaatcgtccgatccggagccgggactgtcgggcgtacacaaatcgcccgcagaagcgcggccgtctggaccgatggctgtgtagaagtactcgccgatagtggaaaccgacgccccagcactcgtccgagggcaaaggaataatcagtactgacaataaaaagattcttgttttcaagaacttgtcatttgtatagtttttttatattgtagttgttctattttaatcaaatgttagcgtgatttatattttttttcgcctcgacatcatctgcccagatgcgaagttaagtgcgcagaaagtaatatcatgcgtcaatcgtatgtgaatgctggtcgctatactgggatccccgtgtaggtctacaaactggtattctcgtttgttgtgaatcttgtcttttctgcttactttctaatagattgagtgagttaggtctgtatttgacggctaaagttaaacttaactgttgtgtgggatagttgttagctgtcaagaaaccgttaaacgtagaaagagtaacttttctaaaccgtagtgtctactcaatattgagacaagcagctagtttaagagcagaagctgaagtctgtgagtttggaaactaaaaggtctgtgaacagctaactagaactaggcttgcttttagtattagaataacaacaaacgaggtctattagttgtgggatctagctagttgcctaagaacaaaggactaagtctgcagaaaaacgagtttcaataggtctgtgtgtaacagtgttaaggctactagcagaactgccacctaagtttcgtaagttccacctcagccttcaatattccagacttgaccaactttcagtttttaaactctaccgtactagtagtttctgcacagtattagacctacttacactgtctgtgcttgcaccacaaaccaaatagtagaatctcttttaagtagaacagtcttttcttgttagactaaaacaacgagagacaactgagtgtagggttacagttaaaaaactgtacagtctgctgtacagacgttacacggcagtcctgttccaaacttcacttaagggttagattggctaagttcaacttgtccttttaagttagttcagacctagttagcagacttaggtgggtaagctgagctacaagagttaaaaacaactgctaagactaaagtaagtgtacgtaagcttcaaactagtaagttaaacttagaaacagctaacgtctaaagctagaagtaagctcaatcttagtactgttaacctaagttgctcacgcaccttcgccaactcaacctgcagggcgatcgccgcgtgagaccggtaccgagctcgaattcactggccgtcgttttacaacgtcgtgactgggaaaaccctggcgttacccaacttaatcgccttgcagcacatccccctttcgccagctggcgtaatagcgaagaggcccgcaccgatcgcccttcccaacagttgcgcagcctgaatggcgaatggcgcctgatgcggtattttctccttacgcatctgtgcggtatttcacaccgcatatggtgcactctcagtacaatctgctctgatgccgcatagttaagccagccccgacacccgccaacacccgctgacgcgccctgacgggcttgtctgctcccggcatccgcttacagacaagctgtgaccgtctccgggagctgcatgtgtcagaggttttcaccgtcatcaccgaaacgcgcgagacgaaagggcctcgtgatacgcctatttttataggttaatgtcatgataataatggtttcttagacgtcaggtggcacttttcggggaaatgtgcgcggaacccctatttgtttatttttctaaatacattcaaatatgtatccgctcatgagacaataaccctgataaatgcttcaataatattgaaaaaggaagagtatgagtattcaacatttccgtgtcgcccttattcccttttttgcggcattttgccttcctgtttttgctcacccagaaacgctggtgaaagtaaaagatgctgaagatcagttgggtgcacgagtgggttacatcgaactggatctcaacagcggtaagatccttgagagttttcgccccgaagaacgttttccaatgatgagcacttttaaagttctgctatgtggcgcggtattatcccgtattgacgccgggcaagagcaactcggtcgccgcatacactattctcagaatgacttggttgagtactcaccagtcacagaaaagcatcttacggatggcatgacagtaagagaattatgcagtgctgccataaccatgagtgataacactgcggccaacttacttctgacaacgatcggaggaccgaaggagctaaccgcttttttgcacaacatgggggatcatgtaactcgccttgatcgttgggaaccggagctgaatgaagccataccaaacgacgagcgtgacaccacgatgcctgtagcaatggcaacaacgttgcgcaaactattaactggcgaactacttactctagcttcccggcaacaattaatagactggatggaggcggataaagttgcaggaccacttctgcgctcggcccttccggctggctggtttattgctgataaatctggagccggtgagcgtgggtctcgcggtatcattgcagcactggggccagatggtaagccctcccgtatcgtagttatctacacgacggggagtcaggcaactatggatgaacgaaatagacagatcgctgagataggtgcctcactgattaagcattggtaactgtcagaccaagtttactcatatatactttagattgatttaaaacttcatttttaatttaaaaggatctaggtgaagatcctttttgataatctcatgaccaaaatcccttaacgtgagttttcgttccactgagcgtcagaccccgtagaaaagatcaaaggatcttcttgagatcctttttttctgcgcgtaatctgctgcttgcaaacaaaaaaaccaccgctaccagcggtggtttgtttgccggatcaagagctaccaactctttttccgaaggtaactggcttcagcagagcgcagataccaaatactgttcttctagtgtagccgtagttaggccaccacttcaagaactctgtagcaccgcctacatacctcgctctgctaatcctgttaccagtggctgctgccagtggcgataagtcgtgtcttaccgggttggactcaagacgatagttaccggataaggcgcagcggtcgggctgaacggggggttcgtgcacacagcccagcttggagcgaacgacctacaccgaact

**eDA201**

cagggtaatgcatccagcatccagcatccagcatccagcatccagcatccgcatccagcatccagcatccagcatccagcatccagcatccacgatcggaggaccgaaggagctaaccgcttttttgcacaacatgggggatcatgtaactcgccttgatcgttgggaaccggagctgaatgaagccataccaaacgacgagcgtgacaccacgatgcctgtagcaatggcaacaacgttgcgcaaactattaactggcgaactacttactctagcttcccggcaacaattaatagactggatggaggcggataaagttgcaggaccacttctgcgctcggcccttccggctggctggtttattgctgataaatctggagccggtgagcgtgggtctcgcggtatcattgcagcactggggccagatggtaagccctcccgtatcgtagttatctacacgacggggagtcaggcaactatggatgaacgaaatagacagatcgctgagataggtgcctcactgattaagcattggtaactgtcagaccaagtttactcatatatactttagattgatttaaaacttcatttttaatttaaaaggatctaggtgaagatcctttttgataatctcatgaccaaaatcccttaacgtgagttttcgttccactgagcgtcagaccccgtagaaaagatcaaaggatcttcttgagatcctttttttctgcgcgtaatctgctgcttgcaaacaaaaaaaccaccgctaccagcggtggtttgtttgccggatcaagagctaccaactctttttccgaaggtaactggcttcagcagagcgcagataccaaatactgttcttctagtgtagccgtagttaggccaccacttcaagaactctgtagcaccgcctacatacctcgctctgctaatcctgttaccagtggctgctgccagtggcgataagtcgtgtcttaccgggttggactcaagacgatagttaccggataaggcgcagcggtcgggctgaacggggggttcgtgcacacagcccagcttggagcgaacgacctacaccgaactgagatacctacagcgtgagctatgagaaagcgccacgcttcccgaagggagaaaggcggacaggtatccggtaagcggcagggtcggaacaggagagcgcacgagggagcttccagggggaaacgcctggtatctttatagtcctgtcgggtttcgccacctctgacttgagcgtcgatttttgtgatgctcgtcaggggggcggagcctatggaaaaacgccagcaacgcggcctttttacggttcctggccttttgctggccttttgctggccttttgctcacatgttctttcctgcgttatcccctgattctgtggataaccgtattaccgcctttgagtgagctgataccgctcgccgcagccgaacgaccgagcgcagcgagtcagtgagcgaggaagcggaagagcgcccaatacgcaaaccgcctctccccgcgcgttggccgattcattaatgcagttgtcattttgtttcttctcgtacgagcttgctcctgatcagcctatctcgcaggctgacttacctcgttttaacttagtcggttagtgaagtttaagtctaagttagttcagacctttattgcagattcgtcttactgtctacttagccctgcgtttgacttgcctattagctactcaacttaaaacagttaaactcacagactttacgattgaaaacaagttgacttgaaccccagtggacgaactccacaagtctagtttggcttagagtttctactttacctctattctaccgcagtaactttcaacctgaagtaagttcaagtgaaacttagagtgaagcttgtgatcgactacaagctcaatacaaacttctagaccttagaactagcttagttgcttaacctagtgaatccaacaacaaagttaccaagaatagacagtacgacaactggttacgagtgtaactaagaatcttacactctagttctgccttgctgcttattcaccttttagcttttgttgtttttagttttagccaaccaaaggtggaggtttaaacccttgaactaagcttaggttcaactgaaagtctgtctgccttacagacttgtacagacaaaagttaaacctctaaagcaaccagtctactagttttcgtctttctacgtaactattaccacgagagtaacaagtttctaacacaagtctgttgagccaaatccaaatctttaaacccttgagttagttgagtttcgacgttagaactagctttttgtagtaagcttattcacaaacctacttaagtttagactgtagttaccgtactgaaaagattgctgacttttctacttttacgtttacaacagctgtgtagcgtgcaacagacgtgagattcaaggctgacgcttagttaaacttgtcgaccacacaccatagcttcaaaatgtttctactccttttttactcttccagattttctcggactccgcgcatcgccgtaccacttcaaaacacccaagcacagcatactaaattttccctctttcttcctctagggtgtcgttaattacccgtactaaaggtttggaaaagaaaaaagagaccgcctcgtttctttttcttcgtcgaaaaaggcaataaaaatttttatcacgtttctttttcttgaaatttttttttttagtttttttctctttcagtgacctccattgatatttaagttaataaacggtcttcaatttctcaagtttcagtttcatttttcttgttctattacaactttttttacttcttgttcattagaaagaaagcatagcaatctaatctaaggggcggtgttgacaattaatcatcggcatagtatatcggcatagtataatacgacaaggtgaggaactaaaccatggtatctaaaggtgaagaactattcacaggtgtggtgcccattttagttgaattggatggtgatgtaaacggccacaagttcagtgtcagtggagaaggtgagggtgatgccacctatggaaaactaacattaaaatttatatgcactactggaaagttgcccgtcccctggccaactctagtcactaccctgacatatggagtacagtgtttctcaagatatcctgaccatatgaagcagcatgacttctttaagtccgctatgcctgaaggatacgtccaggaacgtaccatattcttcaaagatgacggtaactataagactagagcagaggtaaaattcgagggagatacgcttgtcaacagaattgagcttaagggaatcgacttcaaggaggatggtaatattttaggccacaaattagagtacaattataattcccacaacgtgtacatcatggccgacaagcagaaaaacggcatcaaagtcaacttcaagatccgtcacaatattgaagatggcagtgtgcaacttgcagaccattatcaacaaaacacaccaataggcgatggacctgtcttgttgcccgacaatcattatttgtcaactcagtccgctctttccaaggacccaaacgagaaaagggaccatatggtactactagagttcgttactgcagctggcataaccttgggaatggatgagctttacaaataagcatcatgtaattagttatgtcacgcttacattcacgccctccccccacatccgctctaaccgaaaaggaaggagttagacaacctgaagtctaggtccctatttatttttttatagttatgttagtattaagaacgttatttatatttcaaatttttcttttttttctgtacagacgcgtgtacgcatgtaacattatactgaaaaccttgcttgagaaggttttgggacgctcgaaggctgtcgaccgtctcagtagcccacacaccatagcttcaaaatgtttctactccttttttactcttccagattttctcggactccgcgcatcgccgtaccacttcaaaacacccaagcacagcatactaaatttcccctctttcttcctctagggtgtcgttaattacccgtactaaaggtttggaaaagaaaaaagagaccgcctcgtttctttttcttcgtcgaaaaaggcaataaaaatttttatcacgtttctttttcttgaaaatttttttttttgatttttttctctttcgatgacctcccattgatatttaagttaataaacggtcttcaatttctcaagtttcagtttcatttttcttgttctattacaactttttttacttcttgctcattagaaagaaagcatagcaatctaatctaagggcggtgttgacaattaatcatcggcatagtatatcggcatagtataatacgacaaggtgaggaactaaaccatggccaagttgaccagtgccgttccggtgctcaccgcgcgcgacgtcgccggagcggtcgagttctggaccgaccggctcgggttctcccgggacttcgtggaggacgacttcgccggtgtggtccgggacgacgtgaccctgttcatcagcgcggtccaggaccaggtggtgccggacaacaccctggcctgggtgtgggtgcgcggcctggacgagctgtacgccgagtggtcggaggtcgtgtccacgaacttccgggacgcctccgggccggccatgaccgagatcggcgagcagccgtgggggcgggagttcgccctgcgcgacccggccggcaactgcgtgcacttcgtggccgaggagcaggactgacacgtccgacgcggcccgacgggtccgaggcctcggagatccgtcccccttttcctttgtcgatatcatgtaattagttatgtcacgcttacattcacgccctccccccacatccgctctaaccgaaaaggaaggagttagacaacctgaagtctaggtccctatttatttttttatagttatgttagtattaagaacgttatttatatttcaaatttttcttttttttctgtacagacgcgtgtacgcatgtaacattatactgaaaaccttgcttgagaaggttttgggacgctcgaaggctttaatttgcggatccccgtgtaggtctacaaactggtattctcgtttgttgtgaatcttgtcttttctgcttactttctaatagattgagtgagttaggtctgtatttgacggctaaagttaaacttaactgttgtgtgggatagttgttagctgtcaagaaaccgttaaacgtagaaagagtaacttttctaaaccgtagtgtctactcaatattgagacaagcagctagtttaagagcagaagctgaagtctgtgagtttggaaactaaaaggtctgtgaacagctaactagaactaggcttgcttttagtattagaataacaacaaacgaggtctattagttgtgggatctagctagttgcctaagaacaaaggactaagtctgcagaaaaacgagtttcaataggtctgtgtgtaacagtgttaaggctactagcagaactgccacctaagtttcgtaagttccacctcagccttcaatattccagacttgaccaactttcagtttttaaactctaccgtactagtagtttctgcacagtattagacctacttacactgtctgtgcttgcaccacaaaccaaatagtagaatctcttttaagtagaacagtcttttcttgttagactaaaacaacgagagacaactgagtgtagggttacagttaaaaaactgtacagtctgctgtacagacgttacacggcagtcctgttccaaacttcacttaagggttagattggctaagttcaacttgtccttttaagttagttcagacctagttagcagacttaggtgggtaagctgagctacaagagttaaaaacaactgctaagactaaagtaagtgtacgtaagcttcaaactagtaagttaaacttagaaacagctaacgtctaaagctagaagtaagctcaatcttagtactgttaacctaagttgctcacgcaccttcgccaactcactcgactctagaggggattcggcgctccaactgtattctaggcctttccacaatttaaaaaaagacctccgcgacggggaattgaaccccggtctaccgcgcgacaagcggtggttctaccactaaactatcacggatttcgatgctgctttcccacttccttactttacattctctaggcgtttagcctgtagaataaaaccttttttcaagactacaatactgctgctagtaataccgaattattacatgttttaacaacttgagagcgggtcgtatcctgtcttccgattgtacaatctcatcttatcagctcatctcatctcttcggaatcccccccgggtaccgagctcgaattcactggccgtcgttttacaacgtcgtgactgggaaaaccctggcgttacccaacttaatcgccttgcagcacatcccgggtaccgtagatcaagtgacttcttgagcttgccaagcatcaagtcccaccgacaagtggtacaccagtcagcgttatctttccatacattctttaattcaaattgtcccattcacaactcaatgtttttgtttctctcgacgaaacgtttttgcttagcatttcggtatcttaaaccgtgtgattttaagtctcataaccttctacatacaggatcaataataatattattgttgatacaatataggattagaatacatagattctgatgttcgaatcaatcaatgtcttaatattcacgtacttgtataccctctgattccacattggtccaacggaatcttctagcaacagccagttgcactagatgtggttctccaccagaaattatagtactgacagattttttggtttacggtgagaggctccaatttccaaagctaaaccaatttcttcataagcaacctccacgcgcgcttttagtatgatgtagaaactggacctaaagatcgaccaatacttgcttacacggcagcgcgaaataaaagacattgataatgtagagtaagtactatttcttcaaaagatagaaccactctcagaggctaactagaagatctaaaacggatgtacagctaatatgtttcgaggcaatcatgttcaggtgcgcctcaagtggctcttaatctattttaaagcataatccacgaaaatttcgatacagagagatacgtcataaactgagaacgatagttagaggcatagcttcggcaatgttcgataaacaagtcaacgagtcagaatttcatgtttctttttttcttgtagtaatatgtaataaccgaatcagataacttgccagtaatgacctgagctctataaattgaaaccgctacaaaaagtaagggatcgtgttaagggcacagagaagaaagtgtaaagaagtaagctagcgttgtacaacaactcaaaagcatggagcccggtgggtactagataataagaacaaggtttcagtgagtttacaagttcataaaacacggaacctggcaacatagcataccagtaaatcgatttgaagaactctattaggctggcagcctctctcaatcaatagtctaaaacggttaagagacacaatcatctctaggtgtcgttgtggtctaccatcgaaacactggaaatcaatccaatgaacttcaagtgatatatttttcaagagtcttcaaagaccaagcaatgttcttattgattgataattttaaagttgaagtctaaatttgtagcagtgaagtctgagcagggggaatgttaagccatatctaatagtacctattagacgcttgattgccgtttgcaaatatgccatgtttctttgaaaccaaaaccacgtaagggggcttaagttttcggcttaggattgttacggagctcaaaccaaataaggcagagagcatacaaaacgtttattaaaaagaacaactacttctggatagctcaattatcttttgtttcttttgaggcgtgcccttcatgacatgacattgcatccatacaattaatagtagataggggaggataagtttgctagtgtgagcatttagagcaaggggtcagtttcctccatctgcctactctgctccatttaaaagcgagccgattgccataagcttgctcacgtagataggacttcaaaaagacgttaagaggctgctctctagaatacgatggaataaacaaatctcgttagtttcttgaaacggaaagtacatgtaggataattgctggatcattcggaagctaaaattggtcctttccccaaagacaatcgactatagttcctgaagcttcctgaggtgagctggatagaaccagaagatagtacctataacgtcaacaacaactgaaactacagatggaagagctaggcatatccataggggatagccatagaaaactttgattgacgcagcaaatcgagacgttaacctgaagctgccataacttcaggaacttgaacacaagagtggcattaaaattcctatgttgcatttcagaaataccacaagtaaactgagcaattaacttgttcataccgacactacaagttaactacaaaccgagacccttatatgcagactgataatacagaatgatacgtatcactcctaagactaggaacatacagagcttttgagcttttggtattttcaagttatttgaaaaattaaatctttatactagggacgaggttcgtgacaaaagaagacaatcatgagactcaccgtcttgtcatctacaagtttcaataagcttccaacttgagtaacctagccatgtgattagtaactttcgaagcatgattcagaacgttctgctctgccgcgtcaaaaagggcagctacttgaagttaaaagaaagatcaaaaattcctgagcattgttcaaacagctataaatcaaaacagaataagaaataaggccatcacttaccaaggaaaaacaaaaaagttcgagagatcaacaatgcacttcagggacgcacctgaacctttacattagtgtaaaaataaattattagattgttcggttcgagcgttagccatatacaaaatgcggccttcaaagtcattggaaaagagctgctttaggacctctccaaaagctaaattgacaaacaggctacctcatatctccatcatctttgccaaccttttcagtaacaaaaaacaaaagcaatatcgtttatttgtccattggacgatcatcaaatattgctaggtaggcacttttttcggtaagcgtgtctgcctcctttcttcagtaagatcaattgtcgccggatactatgaatcgatcattgtagaaactaacatgcttctaattacccattcaatcttgtagcactgagtgcatctcgcgtgagacaaattgaactcgcctacttgggtaggataaacggttgggtaacgtagctgcttcatccggctacttgcttggtctgctgcgtgctttacgcgctgattggtggtcgaataaaccggtacattatttgttcttatccacgacaatgagacgaacaagtaacaaaccagaactggtagctcgatgattacagttccacaatatcagccgaatacccacactttttccaacctgtgtcaattgccttagaaattgtattagcacaaaaatcatttcgctgacggctgaacatccgatatttactattcgtcaaggtatttagaattcttgtgcgaacttgtgggtcgtggtgaaaaagactattgtgagttgttcgtagctgagtccttgactatactgatacacattaaaagaagctgccctttaagcaactaatgtatttgttatgcgtatatctgtagaaagcaacagtgcgacagttgaagtgttcgaacacacttctcaattttgaatctttctcattctcattcagtgaacgatgaaccaagtacacaaatttggaggtgaataattatgctagggagagcttgctagatcatttcacaattagaatgtgaaaccagacctttagatgtaaatgctaaggctaggataggatcgtaattagtaggatcaacaaactttattgattagcctcacctagactaagtttgatcttttcttcatcgtccaatggacaaataaacgatattgcttttgttttttgttactgaaaaggttggcaaagatgatggagatatgaggtagcctgtttgtcaatttagcttttggagaggtcctaaagcagctcttttccaatgactttgaaggccgcattttgtatatggctaacgctcgaaccgaacaatctaataatttatttttacactaatgtaaaggttcaggtgcgtccctgaagtgcattgttgatctctcgaacttttttgtttttccttggtaagtgatggccttatttcttattctgttttgatttatagctgtttgaacaatgctcaggaatttttgatctttcttttaacttcaagtagctgccctttttgacgcggcagagcagaacgttctgaatcatgcttcgaaagttactaatcacatggctaggttactcaagttggaagcttattgaaacttgtagatgacaagacggtgagtctcatgattgtcttcttttgtcacgaacctcgtccctagtataaagatttaatttttcaaataacttgaaaataccaaaagctcaaaagctctgtatgttcctagtcttaggagtgatacgtatcattctgtattatcagtctgcatataagggtctcggtttgtagttaacttgtagtgtcggtatgaacaagttaattgctcagtttacttgtggtatttctgaaatgcaacataggaattttaatgccactcttgtgttcaagttcctgaagttatggcagcttcaggttaacgtctcgatttgctgcgtcaatcaaagttttctatggctatcccctatggatatgcctagctcttccatctgtagtttcagttgttgttgacgttataggtactatcttctggttctatccagctcacctcaggaagcttcaggaactatagtcgattgtctttggggaaaggaccaattttagcttccgaatgatccagcaattatcctacatgtactttccgtttcaagaaactaacgagatttgtttattccatcgtattctagagagcagcctcttaacgtctttttgaagtcctatctacgtgagcaagcttatggcaatcggctcgcttttaaatggagcagagtaggcagatggaggaaactgaccccttgctctaaatgctcacactagcaaacttatcctcccctatctactattaattgtatggatgcaatgtcatgtcatgaagggcacgcctcaaaagaaacaaaagataattgagctatccagaagtagttgttctttttaataaacgttttgtatgctctctgccttatttggtttgagctccgtaacaatcctaagccgaaaacttaagcccccttacgtggttttggtttcaaagaaacatggcatatttgcaaacggcaatcaagcgtctaataggtactattagatatggcttaacattccccctgctcagacttcactgctacaaatttagacttcaactttaaaattatcaatcaataagaacattgcttggtctttgaagactcttgaaaaatatatcacttgaagttcattggattgatttccagtgtttcgatggtagaccacaacgacacctagagatgattgtgtctcttaaccgttttagactattgattgagagaggctgccagcctaatagagttcttcaaatcgatttactggtatgctatgttgccaggttccgtgttttatgaacttgtaaactcactgaaaccttgttcttattatctagtacccaccgggctccatgcttttgagttgttgtacaacgctagcttacttctttacactttcttctctgtgcccttaacacgatcccttactttttgtagcggtttcaatttatagagctcaggtcattactggcaagttatctgattcggttattacatattactacaagaaaaaaaagaaacatgaaattctgactcgttgacttgtttatcgaacattgccgaagctatgcctctaactatcgttctcagtttatgacgtatctctctgtatcgaaattttcgtggattatgctttaaaatagattaagagccacttgaggcgcacctgaacatgattgcctcgaaacatattagctgtacatccgttttagatcttctagttagcctctgagagtggttctatcttttgaagaaatagtacttactctacattatcaatgtcttttatttcgcgctgccgtgtaagcaagtattggtcgatctttaggtccagtttctacatcatactaaaagcgcgcgtggaggttgcttatgaagaaattggtttagctttggaaattggagcctctcaccgtaaaccaaaaaatctgtcagtactataatttctggtggagaaccacatctagtgcaactggctgttgctagaagattccgttggaccaatgtggaatcagagggtatacaagtacgtgaatattaagacattgattgattcgaacatcagaatctatgtattctaatcctatattgtatcaacaataatattattattgatcctgtatgtaaaaggttatgagacttaaaatcacacggtttaagataccgaaatgctaagcaaaaacgtttcgtcgagagaaacaaaaacattgagttgcgttataaaatagatagtaacagacttgctaactgagcattcaagactacgttctatttcaggggcttattcaagctgacggagcacatatgcaacaatactacggaaacgacccataacgctgaaacagccgtggaactcgatggaactttcttgccgatttgttgcccaatgtgctccaaaacggagggaagtctttccgcggacagtctggcttgaacgggctagagatggcagagatcggtttgacttagcactcttaaattcatgctttagggttatgtcgcgagacattaatattgggcaagccagggcagatttggctatattcaggctggtttaggcgatccaaaaaggattatgtaagggattttgactgggatggataaaatagttctgattggggcttaatggggctcaagtggctgtgttttctgattgatattggaacctgctgtcatttcgacaattaatatttacttattttggtcaaccccaaataggttgatttcatacttggttcattcaaaaataagtagtcttttgagatctttcaatattataataaatatactataacagccgacttgtttcattttcgcgaatgttcccccagcttatcggatcccgagctcgaattcgctagccagatcggcttaagacaaccaaacccagaactagtcgaatagtctgtgctaaaaaaccaatcggttaaaactagaatcgatctttaagtttaagattcaggcttgttgttaaacctattgccagaaaggacagttgactttggaacaaactcagctaacttgaaactcaaacaacagttttgttacttgccggcttacctcacttgctcagctagtctagcaagaaaacagtgtttcttgtcaacttagaactaatagcagagttaagcagaatcttgaaggctactctgtatttgaaaaccaattgacttgctttttacttagccttacttgctagtcaatcttcagggcagaagtaacttgaaaagttagaaaccaagtttaaaaagacaacaagaaacttgcacacaggactgagaaagtaactaagatttgaaactgagtagactaactctaccagtggttcacaagggttccaccctactttaacttttgacaagagttcaaagtcgaactttaaactgagcttacctctgtggactgtaaactcttgagttaccaaacttttgagctagcctctcagactgttactggttagctctacacagtcttttactattctaacgtaagcttagactagttcttggaataagtgaatctaagaagttgaggctaaactctcaaatagtgtagactaggcacgaactagaagttagtttcttacgttggtagaaaaaaacagttgagtaggctgaaaaggtcaactaagccaaacaagtttcttgtcagaagtctaactgagtgaatacagttgtattcagaaactgtagctagttcgtcttctcaaaccaacttaccagacttttttaaacttgcagtttttactgtacgagacagtttgttactcttcacagtcagttagacgtaaagtaacttgaataagctgaatactaacgtccacgaggtaggcagactatcctgttactgtcttttttagtctgcagtacaaaaagttaaactgttagcaatcctcggttaggcaactttgagttaacttagaagttaccactgcaagctcaatagattcgccctgcagattctaaaaacaaaagctgagctaaagctgctaactgcagttaggtttagatttgagaatagacttggtaaactgcagtaacttacgagaaagtaactaaaactacagttttcaacttcagttttcgaactgtagttgtcttagtctgagttgaaagactgagttaagtaagttaactcactaggttacgctaaagtgtagctactttggattcaggctcactctgcactagcatatggtgcactctcagtacaatctgctctgatgccgcatagttaagccagccccgacacccgccaacacccgctgacgcgccctgacgggcttgtctgctcccggcatccgcttacagacaagctgtgaccgtctccgggagctgcatgtgtcagaggttttcaccgtcatcaccgaaacgcgcgagacgaaagggcctcgtgatacgcctatttttataggttaatgtcatgataataatggtttcttagacgtcaggtggcacttttcggggaaatgtgcgcggaacccctatttgtttatttttctaaatacattcaaatatgtatccgctcatgagacaataaccctgataaatgcttcaataatattgaaaaaggaagagtatgagtattcaacatttccgtgtcgcccttattcccttttttgcggcattttgccttcctgtttttgctcacccagaaacgctggtgaaagtaaaagatgctgaagatcagttgggtgcacgagtgggttacatcgaactggatctcaacagcggtaagatccttgagagttttcgccccgaagaacgttttccaatgatgagcacttttaaagttctgctatgtggcgcggtattatcccgtattgacgccgggcaagagcaactcggtcgccgcatacactattctcagaatgacttggttgagtactcaccagtcacagaaaagcatcttacggatggcatgacagtaagagaattatgcagtgctgccataaccatgagtgataacactgcggccaacttacttctgacaacgatcggaggaccgaaggagctaaccgcttttttgcacaacatgggggatcatgtaactcgccttgatcgttgggaaccggagctgaatgaagccataccaaacgacgagcgtgacaccacgatgcctgtagcaatggcaacaacgttgcctggcgtaatagcgaagaggcccgcaccgatcgtggatgctggatgctggatgctggatgctggatgctggatgcgcatccagcatccagcatccagcatccagcatccagcatcctggatgctggatgctggatgctggatgctggatgctggatgctggatgctggatgctggatgctggatgctggatgctggatgctggatgctggatgcta

s

***eDA229:***

cagggtaatgcatccagcatccagcatccagcatccagcatccagcatccagcatccagcatccagcatccagcatccagcatccagcatccacgatcggaggaccgaaggagctaaccgcttttttgcacaacatgggggatcatgtaactcgccttgatcgttgggaaccggagctgaatgaagccataccaaacgacgagcgtgacaccacgatgcctgtagcaatggcaacaacgttgcgcaaactattaactggcgaactacttactctagcttcccggcaacaattaatagactggatggaggcggataaagttgcaggaccacttctgcgctcggcccttccggctggctggtttattgctgataaatctggagccggtgagcgtgggtctcgcggtatcattgcagcactggggccagatggtaagccctcccgtatcgtagttatctacacgacggggagtcaggcaactatggatgaacgaaatagacagatcgctgagataggtgcctcactgattaagcattggtaactgtcagaccaagtttactcatatatactttagattgatttaaaacttcatttttaatttaaaaggatctaggtgaagatcctttttgataatctcatgaccaaaatcccttaacgtgagttttcgttccactgagcgtcagaccccgtagaaaagatcaaaggatcttcttgagatcctttttttctgcgcgtaatctgctgcttgcaaacaaaaaaaccaccgctaccagcggtggtttgtttgccggatcaagagctaccaactctttttccgaaggtaactggcttcagcagagcgcagataccaaatactgttcttctagtgtagccgtagttaggccaccacttcaagaactctgtagcaccgcctacatacctcgctctgctaatcctgttaccagtggctgctgccagtggcgataagtcgtgtcttaccgggttggactcaagacgatagttaccggataaggcgcagcggtcgggctgaacggggggttcgtgcacacagcccagcttggagcgaacgacctacaccgaactgagatacctacagcgtgagctatgagaaagcgccacgcttcccgaagggagaaaggcggacaggtatccggtaagcggcagggtcggaacaggagagcgcacgagggagcttccagggggaaacgcctggtatctttatagtcctgtcgggtttcgccacctctgacttgagcgtcgatttttgtgatgctcgtcaggggggcggagcctatggaaaaacgccagcaacgcggcctttttacggttcctggccttttgctggccttttgctggccttttgctcacatgttctttcctgcgttatcccctgattctgtggataaccgtattaccgcctttgagtgagctgataccgctcgccgcagccgaacgaccgagcgcagcgagtcagtgagcgaggaagcggaagagcgcccaatacgcaaaccgcctctccccgcgcgttggccgattcattaatgcagttgtcattttgtttcttctcgtacgagcttgctcctgatcagcctatctcgcaggctgtgttaaaccgtctttaagtcaacccacagtctactgcaatcgtattcagaactagccactagactgttacaagtcgaaacctcagttaaccaactagaaactctagttaccaagttagaactgtgagtacgaaaagtctgaaaagcagaaagattcaatagatttgtctgtgttacacaagagttcaacaagtagctgcgttcgtgctagttgtctaggtttaaacgtttaaaaagacaactagtagtttacttcacctctgtattactgacggtttgtgcctcaacagtttacgttaacaaactagtaagcgtctacttcgtggacttgtacttttagtagaaagattgagtgtctggcagttttaaacctaagtcagttagaatctcaatcctctgcctttggtttagaaacagtgtagctgttttgacctagactgagtacaactgttcttgctgcttattcgaaacttgcctaactcactgcaacagacaagtctagtttagaataaagtaggctcagttattcaagtctaacttcagacagtgtttctcagatttgacttaagtagacagtagtgtgtgaaaaaccagactagtttgaatcttagcctgttcagaaaaaactaactgacttgtgtcaggcctcccaaaaaaactaacttcaatcctctgtactattccacaactattagttgttgcaagtctgaagcctagtcttgctttgaccttacaggatagaggttggctcacaactgagtctgtgtgaaaacaaaagtaaagattgccacaagaagtctattctagttagtccttttacgttggatagcaatagtccaatcttagtttgacgtctaagaaaccattattatcatgacattaacctataaaaataggcgtatcacgaggccctttcgtcttcaagaattaattctcatgtttgacagcttatcatcgataagctgactcatgttggtattgtgaaatagacgcagatcgggaacactgaaaaataacagttattattcgagatctaacatccaaagacgaaaggttgaatgaaacctttttgccatccgacatccacaggtccattctcacacataagtgccaaacgcaacaggaggggatacactagcagcagaccgttgcaaacgcaggacctccactcctcttctcctcaacacccacttttgccatcgaaaaaccagcccagttattgggcttgattggagctcgctcattccaattccttctattaggctactaacaccatgactttattagcctgtctatcctggcccccctggcgaggttcatgtttgtttatttccgaatgcaacaagctccgcattacacccgaacatcactccagatgagggctttctgagtgtggggtcaaatagtttcatgttccccaaatggcccaaaactgacagtttaaacgctgtcttggaacctaatatgacaaaagcgtgatctcatccaagatgaactaagtttggttcgttgaaatgctaacggccagttggtcaaaaagaaacttccaaaagtcgccataccgtttgtcttgtttggtattgattgacgaatgctcaaaaataatctcattaatgcttagcgcagtctctctatcgcttctgaaccccggtgcacctgtgccgaaacgcaaatggggaaacacccgctttttggatgattatgcattgtctccacattgtatgcttccaagattctggtgggaatactgctgatagcctaacgttcatgatcaaaatttaactgttctaacccctacttgacagcaatatataaacagaaggaagctgccctgtcttaaacctttttttttatcatcattattagcttactttcataattgcgactggttccaattgacaagttttgattttaacgacttttaacgacaacttgagaagatcaaaaaacaactaattattcgaaggatccaaaacgatgcagttcaactgggacatcaagactgttgcttccatcttgtccgctttgactttggctcaagcttctgaccaagaggctattgctccagaagattcccacgttgttaagttgactgaggctactttcgagtccttcatcacttccaacccacacgttttggctgagtttttcgctccatggtgtggtcactgtaagaagttgggtccagaattggtttccgctgctgagattttgaaggacaacgagcaggttaagatcgctcagatcgactgtactgaagagaaagagttgtgtcagggttacgagatcaagggttacccaactttgaaggttttccacggtgaggttgaagttccatccgactaccaaggtcaaagacaatcccaatccatcgtttcctacatgttgaagcagtccttgccaccagtttccgagatcaacgctactaaggatttggacgacactatcgctgaggctaaagagccagttatcgttcaggttttgccagaggacgcttctaacttggagtccaacactactttctacggtgttgctggtactttgagagagaagttcactttcgtttccactaagtccactgactacgctaagaagtacacttccgactccactccagcttacttgttggttagaccaggtgaggaaccatccgtttactctggtgaagaattggacgagactcatttggttcactggatcgacattgagtccaagcctttgttcggtgacattgacggttccactttcaagtcctacgctgaagctaacatcccattggcttactacttctacgagaacgaggaacagagagccgctgctgctgacattattaagccattcgctaaagaacaaagaggtaagatcaacttcgttggtttggacgctgttaagttcggtaagcacgctaaaaacttgaacatggacgaagagaagttgcctttgtttgttatccacgacttggtttccaacaagaagttcggagttccacaggaccaagagttgactaacaaggacgttactgagttgatcgagaagtttatcgctggtgaggctgagccaatcgttaagtctgaacctatcccagagatccaagaggaaaaggttttcaagttggttggtaaggctcacgacgaggttgttttcgacgaatctaaggacgttttggttaagtactacgctccttggtgtggacactgtaaaagaatggctccagcttatgaagagttggctactttgtacgctaacgacgaagatgcttcctccaaggttgttatcgctaagttggaccacactttgaacgacgttgataacgttgacatccagggataccctacattgatcttgtacccagctggtgacaagtccaaccctcagttgtacgatggttctagagacttggaatccttggctgaattcgttaaggaaagaggtactcacaaggttgacgctttggctttgagacctgttgaggaagaaaaagaggctgaagaggaagctgaatctgaagctgatgctcacgatgagttgcatcaccaccatcatcattaagcggccgcgaattaattcgccttagacatgactgttcctcagttcaagttgggcacttacgagaagaccggtcttgctagattctaatcaagaggatgtcagaatgccatttgcctgagagatgcaggcttcatttttgatacttttttatttgtaacctatatagtataggattttttttgtcattttgtttcttctcgtacgagcttgctcctgatcagcctatctcgcagctgatgaatatcttgtggtaggggtttgggaaaatcattcgagtttgatgtttttcttggtatttcccactcctcttcagagtacagaagattaagtgaggcatgcgacttacctcgttttaacttagtcggttagtgaagtttaagtctaagttagttcagacctttattgcagattcgtcttactgtctacttagccctgcgtttgacttgcctattagctactcaacttaaaacagttaaactcacagactttacgattgaaaacaagttgacttgaaccccagtggacgaactccacaagtctagtttggcttagagtttctactttacctctattctaccgcagtaactttcaacctgaagtaagttcaagtgaaacttagagtgaagcttgtgatcgactacaagctcaatacaaacttctagaccttagaactagcttagttgcttaacctagtgaatccaacaacaaagttaccaagaatagacagtacgacaactggttacgagtgtaactaagaatcttacactctagttctgccttgctgcttattcaccttttagcttttgttgtttttagttttagccaaccaaaggtggaggtttaaacccttgaactaagcttaggttcaactgaaagtctgtctgccttacagacttgtacagacaaaagttaaacctctaaagcaaccagtctactagttttcgtctttctacgtaactattaccacgagagtaacaagtttctaacacaagtctgttgagccaaatccaaatctttaaacccttgagttagttgagtttcgacgttagaactagctttttgtagtaagcttattcacaaacctacttaagtttagactgtagttaccgtactgaaaagattgctgacttttctacttttacgtttacaacagctgtgtagcgtgcaacagacgtgagattcaaggctgacgcttagttaaacttgtcgacggtctcagtaggacatggaggcccagaataccctccttgacagtcttgacgtgcgcagctcaggggcatgatgtgactgtcgcccgtacatttagcccatacatccccatgtataatcatttgcatccatacattttgatggccgcacggcgcgaagcaaaaattacggctcctcgctgcggacctgcgagcagggaaacgctcccctcacagacgcgttgaattgtccccacgccgcgcccctgtagagaaatataaaaggttaggatttgccactgaggttcttctttcatatacttccttttaaaatcttgctaggatacagttctcacatcacatccgaacataaacaaccatgggtaaaaagcctgaactcaccgcgacgtctgtcgagaagtttctgatcgaaaagttcgacagcgtctccgacctgatgcagctctcggagggcgaagaatctcgtgctttcagcttcgatgtaggagggcgtggatatgtcctgcgggtaaatagctgcgccgatggtttctacaaagatcgttatgtttatcggcactttgcatcggccgcgctcccgattccggaagtgcttgacattggggaattcagcgagagcctgacctattgcatctcccgccgtgcacagggtgtcacgttgcaagacctgcctgaaaccgaactgcccgctgttctgcagccggtcgcggaggccatggatgcgattgctgcggccgatcttagccagacgagcgggttcggcccattcggaccgcaaggaatcggtcaatacactacatggcgtgatttcatatgcgcgattgctgatccccatgtgtatcactggcaaactgtgatggacgacaccgtcagtgcgtccgtcgcgcaggctctcgatgagctgatgctttgggccgaggactgccccgaagtccggcacctcgtgcacgcggatttcggctccaacaatgtcctgacggacaatggccgcataacagcggtcattgactggagcgaggcgatgttcggggattcccaatacgaggtcgccaacatcttcttctggaggccgtggttggcttgtatggagcagcagacgcgctacttcgagcggaggcatccggagcttgcaggatcgccgcggctccgggcgtatatgctccgcattggtcttgaccaactctatcagagcttggttgacggcaatttcgatgatgcagcttgggcgcagggtcgatgcgacgcaatcgtccgatccggagccgggactgtcgggcgtacacaaatcgcccgcagaagcgcggccgtctggaccgatggctgtgtagaagtactcgccgatagtggaaaccgacgccccagcactcgtccgagggcaaaggaataatcagtactgacaataaaaagattcttgttttcaagaacttgtcatttgtatagtttttttatattgtagttgttctattttaatcaaatgttagcgtgatttatattttttttcgcctcgacatcatctgcccagatgcgaagttaagtgcgcagaaagtaatatcatgcgtcaatcgtatgtgaatgctggtcgctatactgggatccccgtgtaggtctacaaactggtattctcgtttgttgtgaatcttgtcttttctgcttactttctaatagattgagtgagttaggtctgtatttgacggctaaagttaaacttaactgttgtgtgggatagttgttagctgtcaagaaaccgttaaacgtagaaagagtaacttttctaaaccgtagtgtctactcaatattgagacaagcagctagtttaagagcagaagctgaagtctgtgagtttggaaactaaaaggtctgtgaacagctaactagaactaggcttgcttttagtattagaataacaacaaacgaggtctattagttgtgggatctagctagttgcctaagaacaaaggactaagtctgcagaaaaacgagtttcaataggtctgtgtgtaacagtgttaaggctactagcagaactgccacctaagtttcgtaagttccacctcagccttcaatattccagacttgaccaactttcagtttttaaactctaccgtactagtagtttctgcacagtattagacctacttacactgtctgtgcttgcaccacaaaccaaatagtagaatctcttttaagtagaacagtcttttcttgttagactaaaacaacgagagacaactgagtgtagggttacagttaaaaaactgtacagtctgctgtacagacgttacacggcagtcctgttccaaacttcacttaagggttagattggctaagttcaacttgtccttttaagttagttcagacctagttagcagacttaggtgggtaagctgagctacaagagttaaaaacaactgctaagactaaagtaagtgtacgtaagcttcaaactagtaagttaaacttagaaacagctaacgtctaaagctagaagtaagctcaatcttagtactgttaacctaagttgctcacgcaccttcgccaactcactcgactctagaggggattcggcgctccaactgtattctaggcctttccacaatttaaaaaaagacctccgcgacggggaattgaaccccggtctaccgcgcgacaagcggtggttctaccactaaactatcacggatttcgatgctgctttcccacttccttactttacattctctaggcgtttagcctgtagaataaaaccttttttcaagactacaatactgctgctagtaataccgaattattacatgttttaacaacttgagagcgggtcgtatcctgtcttccgattgtacaatctcatcttatcagctcatctcatctcttcggaatcccccccgggtaccgagctcgaattcactggccgtcgttttacaacgtcgtgactgggaaaaccctggcgttacccaacttaatcgccttgcagcacatcccgggtaccgtagatcaagtgacttcttgagcttgccaagcatcaagtcccaccgacaagtggtacaccagtcagcgttatctttccatacattctttaattcaaattgtcccattcacaactcaatgtttttgtttctctcgacgaaacgtttttgcttagcatttcggtatcttaaaccgtgtgattttaagtctcataaccttctacatacaggatcaataataatattattgttgatacaatataggattagaatacatagattctgatgttcgaatcaatcaatgtcttaatattcacgtacttgtataccctctgattccacattggtccaacggaatcttctagcaacagccagttgcactagatgtggttctccaccagaaattatagtactgacagattttttggtttacggtgagaggctccaatttccaaagctaaaccaatttcttcataagcaacctccacgcgcgcttttagtatgatgtagaaactggacctaaagatcgaccaatacttgcttacacggcagcgcgaaataaaagacattgataatgtagagtaagtactatttcttcaaaagatagaaccactctcagaggctaactagaagatctaaaacggatgtacagctaatatgtttcgaggcaatcatgttcaggtgcgcctcaagtggctcttaatctattttaaagcataatccacgaaaatttcgatacagagagatacgtcataaactgagaacgatagttagaggcatagcttcggcaatgttcgataaacaagtcaacgagtcagaatttcatgtttctttttttcttgtagtaatatgtaataaccgaatcagataacttgccagtaatgacctgagctctataaattgaaaccgctacaaaaagtaagggatcgtgttaagggcacagagaagaaagtgtaaagaagtaagctagcgttgtacaacaactcaaaagcatggagcccggtgggtactagataataagaacaaggtttcagtgagtttacaagttcataaaacacggaacctggcaacatagcataccagtaaatcgatttgaagaactctattaggctggcagcctctctcaatcaatagtctaaaacggttaagagacacaatcatctctaggtgtcgttgtggtctaccatcgaaacactggaaatcaatccaatgaacttcaagtgatatatttttcaagagtcttcaaagaccaagcaatgttcttattgattgataattttaaagttgaagtctaaatttgtagcagtgaagtctgagcagggggaatgttaagccatatctaatagtacctattagacgcttgattgccgtttgcaaatatgccatgtttctttgaaaccaaaaccacgtaagggggcttaagttttcggcttaggattgttacggagctcaaaccaaataaggcagagagcatacaaaacgtttattaaaaagaacaactacttctggatagctcaattatcttttgtttcttttgaggcgtgcccttcatgacatgacattgcatccatacaattaatagtagataggggaggataagtttgctagtgtgagcatttagagcaaggggtcagtttcctccatctgcctactctgctccatttaaaagcgagccgattgccataagcttgctcacgtagataggacttcaaaaagacgttaagaggctgctctctagaatacgatggaataaacaaatctcgttagtttcttgaaacggaaagtacatgtaggataattgctggatcattcggaagctaaaattggtcctttccccaaagacaatcgactatagttcctgaagcttcctgaggtgagctggatagaaccagaagatagtacctataacgtcaacaacaactgaaactacagatggaagagctaggcatatccataggggatagccatagaaaactttgattgacgcagcaaatcgagacgttaacctgaagctgccataacttcaggaacttgaacacaagagtggcattaaaattcctatgttgcatttcagaaataccacaagtaaactgagcaattaacttgttcataccgacactacaagttaactacaaaccgagacccttatatgcagactgataatacagaatgatacgtatcactcctaagactaggaacatacagagcttttgagcttttggtattttcaagttatttgaaaaattaaatctttatactagggacgaggttcgtgacaaaagaagacaatcatgagactcaccgtcttgtcatctacaagtttcaataagcttccaacttgagtaacctagccatgtgattagtaactttcgaagcatgattcagaacgttctgctctgccgcgtcaaaaagggcagctacttgaagttaaaagaaagatcaaaaattcctgagcattgttcaaacagctataaatcaaaacagaataagaaataaggccatcacttaccaaggaaaaacaaaaaagttcgagagatcaacaatgcacttcagggacgcacctgaacctttacattagtgtaaaaataaattattagattgttcggttcgagcgttagccatatacaaaatgcggccttcaaagtcattggaaaagagctgctttaggacctctccaaaagctaaattgacaaacaggctacctcatatctccatcatctttgccaaccttttcagtaacaaaaaacaaaagcaatatcgtttatttgtccattggacgatcatcaaatattgctaggtaggcacttttttcggtaagcgtgtctgcctcctttcttcagtaagatcaattgtcgccggatactatgaatcgatcattgtagaaactaacatgcttctaattacccattcaatcttgtagcactgagtgcatctcgcgtgagacaaattgaactcgcctacttgggtaggataaacggttgggtaacgtagctgcttcatccggctacttgcttggtctgctgcgtgctttacgcgctgattggtggtcgaataaaccggtacattatttgttcttatccacgacaatgagacgaacaagtaacaaaccagaactggtagctcgatgattacagttccacaatatcagccgaatacccacactttttccaacctgtgtcaattgccttagaaattgtattagcacaaaaatcatttcgctgacggctgaacatccgatatttactattcgtcaaggtatttagaattcttgtgcgaacttgtgggtcgtggtgaaaaagactattgtgagttgttcgtagctgagtccttgactatactgatacacattaaaagaagctgccctttaagcaactaatgtatttgttatgcgtatatctgtagaaagcaacagtgcgacagttgaagtgttcgaacacacttctcaattttgaatctttctcattctcattcagtgaacgatgaaccaagtacacaaatttggaggtgaataattatgctagggagagcttgctagatcatttcacaattagaatgtgaaaccagacctttagatgtaaatgctaaggctaggataggatcgtaattagtaggatcaacaaactttattgattagcctcacctagactaagtttgatcttttcttcatcgtccaatggacaaataaacgatattgcttttgttttttgttactgaaaaggttggcaaagatgatggagatatgaggtagcctgtttgtcaatttagcttttggagaggtcctaaagcagctcttttccaatgactttgaaggccgcattttgtatatggctaacgctcgaaccgaacaatctaataatttatttttacactaatgtaaaggttcaggtgcgtccctgaagtgcattgttgatctctcgaacttttttgtttttccttggtaagtgatggccttatttcttattctgttttgatttatagctgtttgaacaatgctcaggaatttttgatctttcttttaacttcaagtagctgccctttttgacgcggcagagcagaacgttctgaatcatgcttcgaaagttactaatcacatggctaggttactcaagttggaagcttattgaaacttgtagatgacaagacggtgagtctcatgattgtcttcttttgtcacgaacctcgtccctagtataaagatttaatttttcaaataacttgaaaataccaaaagctcaaaagctctgtatgttcctagtcttaggagtgatacgtatcattctgtattatcagtctgcatataagggtctcggtttgtagttaacttgtagtgtcggtatgaacaagttaattgctcagtttacttgtggtatttctgaaatgcaacataggaattttaatgccactcttgtgttcaagttcctgaagttatggcagcttcaggttaacgtctcgatttgctgcgtcaatcaaagttttctatggctatcccctatggatatgcctagctcttccatctgtagtttcagttgttgttgacgttataggtactatcttctggttctatccagctcacctcaggaagcttcaggaactatagtcgattgtctttggggaaaggaccaattttagcttccgaatgatccagcaattatcctacatgtactttccgtttcaagaaactaacgagatttgtttattccatcgtattctagagagcagcctcttaacgtctttttgaagtcctatctacgtgagcaagcttatggcaatcggctcgcttttaaatggagcagagtaggcagatggaggaaactgaccccttgctctaaatgctcacactagcaaacttatcctcccctatctactattaattgtatggatgcaatgtcatgtcatgaagggcacgcctcaaaagaaacaaaagataattgagctatccagaagtagttgttctttttaataaacgttttgtatgctctctgccttatttggtttgagctccgtaacaatcctaagccgaaaacttaagcccccttacgtggttttggtttcaaagaaacatggcatatttgcaaacggcaatcaagcgtctaataggtactattagatatggcttaacattccccctgctcagacttcactgctacaaatttagacttcaactttaaaattatcaatcaataagaacattgcttggtctttgaagactcttgaaaaatatatcacttgaagttcattggattgatttccagtgtttcgatggtagaccacaacgacacctagagatgattgtgtctcttaaccgttttagactattgattgagagaggctgccagcctaatagagttcttcaaatcgatttactggtatgctatgttgccaggttccgtgttttatgaacttgtaaactcactgaaaccttgttcttattatctagtacccaccgggctccatgcttttgagttgttgtacaacgctagcttacttctttacactttcttctctgtgcccttaacacgatcccttactttttgtagcggtttcaatttatagagctcaggtcattactggcaagttatctgattcggttattacatattactacaagaaaaaaaagaaacatgaaattctgactcgttgacttgtttatcgaacattgccgaagctatgcctctaactatcgttctcagtttatgacgtatctctctgtatcgaaattttcgtggattatgctttaaaatagattaagagccacttgaggcgcacctgaacatgattgcctcgaaacatattagctgtacatccgttttagatcttctagttagcctctgagagtggttctatcttttgaagaaatagtacttactctacattatcaatgtcttttatttcgcgctgccgtgtaagcaagtattggtcgatctttaggtccagtttctacatcatactaaaagcgcgcgtggaggttgcttatgaagaaattggtttagctttggaaattggagcctctcaccgtaaaccaaaaaatctgtcagtactataatttctggtggagaaccacatctagtgcaactggctgttgctagaagattccgttggaccaatgtggaatcagagggtatacaagtacgtgaatattaagacattgattgattcgaacatcagaatctatgtattctaatcctatattgtatcaacaataatattattattgatcctgtatgtaaaaggttatgagacttaaaatcacacggtttaagataccgaaatgctaagcaaaaacgtttcgtcgagagaaacaaaaacattgagttgcgttataaaatagatagtaacagacttgctaactgagcattcaagactacgttctatttcaggggcttattcaagctgacggagcacatatgcaacaatactacggaaacgacccataacgctgaaacagccgtggaactcgatggaactttcttgccgatttgttgcccaatgtgctccaaaacggagggaagtctttccgcggacagtctggcttgaacgggctagagatggcagagatcggtttgacttagcactcttaaattcatgctttagggttatgtcgcgagacattaatattgggcaagccagggcagatttggctatattcaggctggtttaggcgatccaaaaaggattatgtaagggattttgactgggatggataaaatagttctgattggggcttaatggggctcaagtggctgtgttttctgattgatattggaacctgctgtcatttcgacaattaatatttacttattttggtcaaccccaaataggttgatttcatacttggttcattcaaaaataagtagtcttttgagatctttcaatattataataaatatactataacagccgacttgtttcattttcgcgaatgttcccccagcttatcggatcccgagctcgaattcgctagccagatcggcttaagacaaccaaacccagaactagtcgaatagtctgtgctaaaaaaccaatcggttaaaactagaatcgatctttaagtttaagattcaggcttgttgttaaacctattgccagaaaggacagttgactttggaacaaactcagctaacttgaaactcaaacaacagttttgttacttgccggcttacctcacttgctcagctagtctagcaagaaaacagtgtttcttgtcaacttagaactaatagcagagttaagcagaatcttgaaggctactctgtatttgaaaaccaattgacttgctttttacttagccttacttgctagtcaatcttcagggcagaagtaacttgaaaagttagaaaccaagtttaaaaagacaacaagaaacttgcacacaggactgagaaagtaactaagatttgaaactgagtagactaactctaccagtggttcacaagggttccaccctactttaacttttgacaagagttcaaagtcgaactttaaactgagcttacctctgtggactgtaaactcttgagttaccaaacttttgagctagcctctcagactgttactggttagctctacacagtcttttactattctaacgtaagcttagactagttcttggaataagtgaatctaagaagttgaggctaaactctcaaatagtgtagactaggcacgaactagaagttagtttcttacgttggtagaaaaaaacagttgagtaggctgaaaaggtcaactaagccaaacaagtttcttgtcagaagtctaactgagtgaatacagttgtattcagaaactgtagctagttcgtcttctcaaaccaacttaccagacttttttaaacttgcagtttttactgtacgagacagtttgttactcttcacagtcagttagacgtaaagtaacttgaataagctgaatactaacgtccacgaggtaggcagactatcctgttactgtcttttttagtctgcagtacaaaaagttaaactgttagcaatcctcggttaggcaactttgagttaacttagaagttaccactgcaagctcaatagattcgccctgcagattctaaaaacaaaagctgagctaaagctgctaactgcagttaggtttagatttgagaatagacttggtaaactgcagtaacttacgagaaagtaactaaaactacagttttcaacttcagttttcgaactgtagttgtcttagtctgagttgaaagactgagttaagtaagttaactcactaggttacgctaaagtgtagctactttggattcaggctcactctgcactagcatatggtgcactctcagtacaatctgctctgatgccgcatagttaagccagccccgacacccgccaacacccgctgacgcgccctgacgggcttgtctgctcccggcatccgcttacagacaagctgtgaccgtctccgggagctgcatgtgtcagaggttttcaccgtcatcaccgaaacgcgcgagacgaaagggcctcgtgatacgcctatttttataggttaatgtcatgataataatggtttcttagacgtcaggtggcacttttcggggaaatgtgcgcggaacccctatttgtttatttttctaaatacattcaaatatgtatccgctcatgagacaataaccctgataaatgcttcaataatattgaaaaaggaagagtatgagtattcaacatttccgtgtcgcccttattcccttttttgcggcattttgccttcctgtttttgctcacccagaaacgctggtgaaagtaaaagatgctgaagatcagttgggtgcacgagtgggttacatcgaactggatctcaacagcggtaagatccttgagagttttcgccccgaagaacgttttccaatgatgagcacttttaaagttctgctatgtggcgcggtattatcccgtattgacgccgggcaagagcaactcggtcgccgcatacactattctcagaatgacttggttgagtactcaccagtcacagaaaagcatcttacggatggcatgacagtaagagaattatgcagtgctgccataaccatgagtgataacactgcggccaacttacttctgacaacgatcggaggaccgaaggagctaaccgcttttttgcacaacatgggggatcatgtaactcgccttgatcgttgggaaccggagctgaatgaagccataccaaacgacgagcgtgacaccacgatgcctgtagcaatggcaacaacgttgcctggcgtaatagcgaagaggcccgcaccgatcgtggatgctggatgctggatgctggatgctggatgctggatgctggatgctggatgctggatgctggatgctggatgctggatgctggatgctggatgctggatgctggatgctggatgctggatgctggatgctggatgctagggataa

***eDA250:***

gagatacctacagcgtgagctatgagaaagcgccacgcttcccgaagggagaaaggcggacaggtatccggtaagcggcagggtcggaacaggagagcgcacgagggagcttccagggggaaacgcctggtatctttatagtcctgtcgggtttcgccacctctgacttgagcgtcgatttttgtgatgctcgtcaggggggcggagcctatggaaaaacgccagcaacgcggcctttttacggttcctggccttttgctggccttttgctcacatgttctttcctgcgttatcccctgattctgtggataaccgtattaccgcctttgagtgagctgataccgctcgccgcagccgaacgaccgagcgcagcgagtcagtgagcgaggaagcggaagagcgcccaatacgcaaaccgcctctccccgcgcgttggccgattcattaatgcagctggcacgacaggtttcccgactggaaagcgggcagtgagcgcaacgcaattaatgtgagttagctcactcattaggcaccccaggctttacactttatgcttccggctcgtatgttgtgtggaattgtgagcggataacaatttcacacaggaaacagctatgaccatgattacgccaagcttgcatgcctgcaggtcgactctagaggatccccgggtaccgagctcgaattcactggccgtcgttttacaacgtcgtgactgggaaaaccctggcgttacccaacttaatcgccttgcagcacatccccctttcgccagctggcgtaatagcgaagaggcccgcaccgatcgcccttcccaacagttgcgcagcctgaatggcgaatggcgcctgatgcggtattttctccttacgcatctgtgcggtatttcacaccgcatatggtgcactctcagtacaatctgctctgatgccgcatagttaagccagccccgacacccgccaacacccgctgacgcgccctgacgggcttgtctgctcccggcatccgcttacagacaagctgtgaccgtctccgggagcgatcgcgacttacctcgttttaacttagtcggttagtgaagtttaagtctaagttagttcagacctttattgcagattcgtcttactgtctacttagccctgcgtttgacttgcctattagctactcaacttaaaacagttaaactcacagactttacgattgaaaacaagttgacttgaaccccagtggacgaactccacaagtctagtttggcttagagtttctactttacctctattctaccgcagtaactttcaacctgaagtaagttcaagtgaaacttagagtgaagcttgtgatcgactacaagctcaatacaaacttctagaccttagaactagcttagttgcttaacctagtgaatccaacaacaaagttaccaagaatagacagtacgacaactggttacgagtgtaactaagaatcttacactctagttctgccttgctgcttattcaccttttagcttttgttgtttttagttttagccaaccaaaggtggaggtttaaacccttgaactaagcttaggttcaactgaaagtctgtctgccttacagacttgtacagacaaaagttaaacctctaaagcaaccagtctactagttttcgtctttctacgtaactattaccacgagagtaacaagtttctaacacaagtctgttgagccaaatccaaatctttaaacccttgagttagttgagtttcgacgttagaactagctttttgtagtaagcttattcacaaacctacttaagtttagactgtagttaccgtactgaaaagattgctgacttttctacttttacgtttacaacagctgtgtagcgtgcaacagacgtgagattcaaggctgacgcttagttaaacttgtgacaaacatccaaagacgaaaggttgaatgaaacctttttgccatccgacatccacaggtccattctcacacataagtgccaaacgcaacaggaggggatacactagcagcagaccgttgcaaacgcaggacctccactcctcttctcctcaacacccacttttgccatcgaaaaaccagcccagttattgggcttgattggagctcgctcattccaattccttctattaggctactaacaccatgactttattagcctgtctatcctggcccccctggcgaggttcatgtttgtttatttccgaatgcaacaagctccgcattacacccgaacatcactccagatgagggctttctgagtgtggggtcaaatagtttcatgttccccaaatggcccaaaactgacagtttaaacgctgtcttggaacctaatatgacaaaagcgtgatctcatccaagatgaactaagtttggttcgttgaaatgctaacggccagttggtcaaaaagaaacttccaaaagtcggcataccgtttgtcttgtttggtattgattgacgaatgctcaaaaataatctcattaatgcttagcgcagtctctctatcgcttctgaaccccggtgcacctgtgccgaaacgcaaatggggaaacacccgctttttggatgattatgcattgtctccacattgtatgcttccaagattctggtgggaatactgctgatagcctaacgttcatgatcaaaatttaactgttctaacccctacttgacagcaatatataaacagaaggaagctgccctgtcttaaacctttttttttatcatcattattagcttactttcataattgcgactggttccaattgacaagcttttgattttaacgacttttaacgacaacttgagaagatcaaaaaacaactaattattcgaaacgatgagatttccttcaatttttactgctgttttattcgcagcatcctccgcattagctgctccagtcaacactacaacagaagatgaaacggcacaaattccggctgaagctgtcatcggttactcagatttagaaggggatttcgatgttgctgttttgccattttccaacagcacaaataacgggttattgtttataaatactactattgccagcattgctgctaaagaagaaggggtatctctcgagaaaagagaggctgaagctgcatccatggtatctaaaggtgaagaactattcacaggtgtggtgcccattttagttgaattggatggtgatgtaaacggccacaagttcagtgtcagtggagaaggtgagggtgatgccacctatggaaaactaacattaaaatttatatgcactactggaaagttgcccgtcccctggccaactctagtcactaccctgacatatggagtacagtgtttctcaagatatcctgaccatatgaagcagcatgacttctttaagtccgctatgcctgaaggatacgtccaggaacgtaccatattcttcaaagatgacggtaactataagactagagcagaggtaaaattcgagggagatacgcttgtcaacagaattgagcttaagggaatcgacttcaaggaggatggtaatattttaggccacaaattagagtacaattataattcccacaacgtgtacatcatggccgacaagcagaaaaacggcatcaaagtcaacttcaagatccgtcacaatattgaagatggcagtgtgcaacttgcagaccattatcaacaaaacacaccaataggcgatggacctgtcttgttgcccgacaatcattatttgtcaactcagtccgctctttccaaggacccaaacgagaaaagggaccatatggtactactagagttcgttactgcagctggcataaccttgggaatggatgagctttacaaaggaggcggtggctctaatgcagcagaggattgtaacgagttgccaccaagaagaaacactgagatcttgactggttcttggagtgatcaaacttacccagagggtactcaggctatctacaagtgtagaccaggttacagatccttgggtaacgttatcatggtttgtagaaagggtgagtgggttgcattgaacccattgagaaagtgtcagaaaagaccatgtggtcacccaggtgatactccattcggtactttcactttgactggtggtaacgttttcgagtacggtgttaaggctgtttacacttgtaacgagggttaccagttgttgggagagatcaactacagagagtgtgatactgacggatggactaacgacattccaatctgtgaagttgttaagtgtttgccagttactgctccagagaacggaaagattgtttcctccgctatggaaccagatagagagtaccacttcggacaggctgttagattcgtttgtaactccggttacaagattgaaggtgacgaagagatgcactgttctgatgacggtttctggtccaaagaaaagccaaagtgtgttgagatctcctgtaagtccccagacgttattaacggttccccaatctcccaaaagatcatctacaaagagaacgagagattccagtacaagtgtaacatgggttacgagtactctgaaagaggtgacgctgtttgtactgaatctggatggagaccattgccatcctgtgaagagaagtcctgtgacaacccatacattccaaacggtgactactccccattgagaatcaagcacagaactggtgacgagatcacttaccagtgtagaaatggtttctacccagctactagaggtaacactgctaagtgtacttccactggatggattccagctccaagatgtactttgaagccatgtgactacccagatatcaagcacggtggtttgtaccacgagaacatgagaaggccatacttcccagttgctgttggaaagtactactcctactactgtgacgaacacttcgaaactccatctggttcttactgggaccacatccactgtactcaagatggttggtccccagctgttccatgtttgagaaaatgttacttcccatacttggagaacggttacaaccagaactacggtagaaagttcgttcagggaaagtccattgacgttgcttgtcatccaggttacgctttgccaaaggctcagactactgttacttgtatggaaaacggttggtcccctactcctagatgtatcagagttaagacttgttccaagtcctccatcgacattgagaacggtttcatttccgagtcccagtacacttacgctttgaaagagaaggctaagtaccagtgtaaattgggatacgttactgctgacggtgaaacttccggatcaatcacatgtggaaaagacggatggagtgctcaaccaacttgtatcaagtcttgtgacatcccagttttcatgaacgctagaactaagaacgacttcacatggttcaagttgaacgacactttggactacgaatgtcacgacggttacgaatctaacactggttccactactggttccatcgtttgtggttacaatggatggagtgacttgccaatctgttacgagagagagtgcgagttgccaaagatcgacgttcatttggttccagacagaaagaaggaccagtacaaagttggagaggttttgaagttctcctgtaagccaggtttcactatcgttggtccaaactccgttcagtgttaccacttcggtttgtctccagacttgcctatctgtaaagagcaggttcaatcctgcggaccaccaccagaattgttgaacggtaacgttaaagaaaagactaaagaagagtacggtcactccgaagttgttgagtactactgtaacccaagattcttgatgaagggtccaaacaagatccaatgtgttgacggtgagtggactactttgccagtttgtatcgttgaagagtccacttgtggtgacattccagaattggaacacggatgggctcaattgtcatccccaccatactactacggtgactccgttgaattcaactgttccgagtccttcactatgattggtcacagatccatcacatgtatccacggtgtttggactcaattgccacagtgtgttgctatcgacaagttgaagaagtgtaaatcatccaaccttatcatcttggaggaacacttgaagaacaagaaagagttcgaccacaactccaacatcagatacagatgtagaggtaaagagggatggatccacactgtttgtatcaacggtagatgggaccctgaagttaactgttccatggctcagattcagttgtgtccaccaccaccacaaattccaaactcccacaacatgactactactttgaactacagagatggtgaaaaggtttccgttttgtgtcaagagaactacttgatccaagagggtgaagagatcacatgtaaggacggtagatggcagtccatccctttgtgtgttgagaagatcccatgttcccaaccacctcaaattgagcacggtactatcaactcttccagatcctctcaagagtcttacgctcacggtactaagttgtcctacacttgtgagggaggtttcagaatctctgaggaaaacgagactacttgttacatgggaaagtggtcatctccaccacaatgtgaaggattgccttgtaagtctccaccagagatttctcacggtgttgttgctcacatgtccgactcttaccaatacggagaagaggttacctacaagtgtttcgagggtttcggtattgatggtccagctatcgctaagtgtttgggagaaaagtggtcccatcctccatcctgtatcaagactgattgtttgtccttgccatccttcgaaaacgctatcccaatgggagaaaagaaggacgtttacaaggctggtgaacaagttacttatacttgtgctacttactacaagatggacggtgcttccaacgttacttgtatcaactccagatggactggtagaccaacttgtagagacacttcctgtgttaacccaccaactgttcagaacgcttacatcgtttccagacagatgtctaagtacccatccggagaacgtgttagataccaatgtagatccccatacgagatgttcggtgacgaagaggttatgtgtttgaacggtaattggactgaaccaccacagtgtaaggactccactggtaagtgtggtccacctccaccaattgacaacggtgacatcacttctttccctttgtccgtttacgctccagcttcttccgttgagtaccagtgtcagaacttgtaccagttggagggtaacaagagaatcacttgtagaaacggacaatggagtgagccaccaaagtgtttgcacccatgtgttatctccagagaaatcatggaaaactacaacattgctttgagatggactgctaaacagaagttgtactccagaactggtgaatccgttgagttcgtttgtaagagaggttacagattgtcctccagatcccacactttgagaactacatgttgggacggaaaattggagtacccaacttgtgctaagagatagtagtctagaacaaaaactcatctcagaagaggatctgaatagcgccgtcgaccatcatcatcatcatcattgagtttgtagccttagacatgactgttcctcagttcaagttgggcacttacgagaagaccggtcttgctagattctaatcaagaggatgtcagaatgccatttgcctgagagatgcaggcttcatttttgatacttttttatttgtaacctatatagtataggattttttttgtcattttgtttcttctcgtacgagcttgctcctgatcagcctatctcgcagctgatgaatatcttgtggtaggggtttgggaaaatcattcgagtttgatgtttttcttggtatttcccactcctcttcagagtacagaagattaagtgagatcaggacagcaacctaaccgacttttacagtgttaacggttgctgttagagttagtccaccagtattcgcagagactttgaatagcagctatccacttacagacaactgtagcagttttagctctttaactctggtccacccgtttgtctagctagcaagtttcttaacttttgaatactggcaagaatagagctaactgtggtttagctcttattcaactaatacaagcctactcagttgtcttgtagaaacaaaaccagttacttcaaacttaacaacagtttcgtcaatctgtcttaagcaacagaggcagcttcaaatacggcaataagtttaactttctaaactagtagtgtagccaatcttagccacgcttctaacttctagcttcttaacccactagtcgaatctgctgtaatacaaagcagacgaactagactcaatccagttgttttttgaactattctaaaacagacttgacgtgcaacgttcaatccaatattcagaatatccacctgactttcaaagacagtaaccgtggactttttgtacactgttttgacttaacccaaccaaagtaacagagctagctaggtttagaaggtgtctgtaaaactaaaactaaaagctaagttagttttcaaaagtttcctgcgttgcacactagctgagtacaagctcaagcacggttttgttgtattgaacttactagtaagtttgaatacagttcttgcgctcacacttgaactctgcaatagttctacttagtagttaacttaagtctgtgcaagtacggcaatcctgtactctgcaatctgtctaaaacaactatttggaactcttagttgacttaactcttggctaaagtggagtttgtacagtcaggttaagtagaactattctaacagtttaaccacaaacctcgttgagttgcccttaaaactaggtttgtttttgggagattcccagacgtaggactactgttgagattggatagagctagactaaagttgaaactcaagtaacctgagagtaacttagccacccaccctactgaacttaaactcgtctttcacgaggactggtagcccacacaccatagcttcaaaatgtttctactccttttttactcttccagattttctcggactccgcgcatcgccgtaccacttcaaaacacccaagcacagcatactaaatttcccctctttcttcctctagggtgtcgttaattacccgtactaaaggtttggaaaagaaaaaagagaccgcctcgtttctttttcttcgtcgaaaaaggcaataaaaatttttatcacgtttctttttcttgaaaatttttttttttgatttttttctctttcgatgacctcccattgatatttaagttaataaacggtcttcaatttctcaagtttcagtttcatttttcttgttctattacaactttttttacttcttgctcattagaaagaaagcatagcaatctaatctaagggcggtgttgacaattaatcatcggcatagtatatcggcatagtataatacgacaaggtgaggaactaaaccatggccaagttgaccagtgccgttccggtgctcaccgcgcgcgacgtcgccggagcggtcgagttctggaccgaccggctcgggttctcccgggacttcgtggaggacgacttcgccggtgtggtccgggacgacgtgaccctgttcatcagcgcggtccaggaccaggtggtgccggacaacaccctggcctgggtgtgggtgcgcggcctggacgagctgtacgccgagtggtcggaggtcgtgtccacgaacttccgggacgcctccgggccggccatgaccgagatcggcgagcagccgtgggggcgggagttcgccctgcgcgacccggccggcaactgcgtgcacttcgtggccgaggagcaggactgacacgtccgacgcggcccgacgggtccgaggcctcgtagatccgtcccccttttcctttgtcgatatcatgtaattagttatgtcacgcttacattcacgccctccccccacatccgctctaaccgaaaaggaaggagttagacaacctgaagtctaggtccctatttatttttttatagttatgttagtattaagaacgttatttatatttcaaatttttcttttttttctgtacagacgcgtgtacgcatgtaacattatactgaaaaccttgcttgagaaggttttgggacgctcgaaggctttaatttgcggatccccgtgtaggtctacaaactggtattctcgtttgttgtgaatcttgtcttttctgcttactttctaatagattgagtgagttaggtctgtatttgacggctaaagttaaacttaactgttgtgtgggatagttgttagctgtcaagaaaccgttaaacgtagaaagagtaacttttctaaaccgtagtgtctactcaatattgagacaagcagctagtttaagagcagaagctgaagtctgtgagtttggaaactaaaaggtctgtgaacagctaactagaactaggcttgcttttagtattagaataacaacaaacgaggtctattagttgtgggatctagctagttgcctaagaacaaaggactaagtctgcagaaaaacgagtttcaataggtctgtgtgtaacagtgttaaggctactagcagaactgccacctaagtttcgtaagttccacctcagccttcaatattccagacttgaccaactttcagtttttaaactctaccgtactagtagtttctgcacagtattagacctacttacactgtctgtgcttgcaccacaaaccaaatagtagaatctcttttaagtagaacagtcttttcttgttagactaaaacaacgagagacaactgagtgtagggttacagttaaaaaactgtacagtctgctgtacagacgttacacggcagtcctgttccaaacttcacttaagggttagattggctaagttcaacttgtccttttaagttagttcagacctagttagcagacttaggtgggtaagctgagctacaagagttaaaaacaactgctaagactaaagtaagtgtacgtaagcttcaaactagtaagttaaacttagaaacagctaacgtctaaagctagaagtaagctcaatcttagtactgttaacctaagttgctcacgcaccttcgccaactcaacctgcagggcgatcgccgcgcgagacgaaagggcctcgtgatacgcctatttttataggttaatgtcatgataataatggtttcttagacgtcaggtggcacttttcggggaaatgtgcgcggaacccctatttgtttatttttctaaatacattcaaatatgtatccgctcatgagacaataaccctgataaatgcttcaataatattgaaaaaggaagagtatgagtattcaacatttccgtgtcgcccttattcccttttttgcggcattttgccttcctgtttttgctcacccagaaacgctggtgaaagtaaaagatgctgaagatcagttgggtgcacgagtgggttacatcgaactggatctcaacagcggtaagatccttgagagttttcgccccgaagaacgttttccaatgatgagcacttttaaagttctgctatgtggcgcggtattatcccgtattgacgccgggcaagagcaactcggtcgccgcatacactattctcagaatgacttggttgagtactcaccagtcacagaaaagcatcttacggatggcatgacagtaagagaattatgcagtgctgccataaccatgagtgataacactgcggccaacttacttctgacaacgatcggaggaccgaaggagctaaccgcttttttgcacaacatgggggatcatgtaactcgccttgatcgttgggaaccggagctgaatgaagccataccaaacgacgagcgtgacaccacgatgcctgtagcaatggcaacaacgttgcgcaaactattaactggcgaactacttactctagcttcccggcaacaattaatagactggatggaggcggataaagttgcaggaccacttctgcgctcggcccttccggctggctggtttattgctgataaatctggagccggtgagcgtgggtctcgcggtatcattgcagcactggggccagatggtaagccctcccgtatcgtagttatctacacgacggggagtcaggcaactatggatgaacgaaatagacagatcgctgagataggtgcctcactgattaagcattggtaactgtcagaccaagtttactcatatatactttagattgatttaaaacttcatttttaatttaaaaggatctaggtgaagatcctttttgataatctcatgaccaaaatcccttaacgtgagttttcgttccactgagcgtcagaccccgtagaaaagatcaaaggatcttcttgagatcctttttttctgcgcgtaatctgctgcttgcaaacaaaaaaaccaccgctaccagcggtggtttgtttgccggatcaagagctaccaactctttttccgaaggtaactggcttcagcagagcgcagataccaaatactgttcttctagtgtagccgtagttaggccaccacttcaagaactctgtagcaccgcctacatacctcgctctgctaatcctgttaccagtggctgctgccagtggcgataagtcgtgtcttaccgggttggactcaagacgatagttaccggataaggcgcagcggtcgggctgaacggggggttcgtgcacacagcccagcttggagcgaacgacctacaccgaact

***contig yDA122***

catccagcatccagcatccagcatccagcatccagcatccagcatccagcatccagcatccagcatccagcatccagcatccagcatccagcatccagcatccagcatccacgatcggaggaccgaaggagctaaccgcttttttgcacaacatgggggatcatgtaactcgccttgatcgttgggaaccggagctgaatgaagccataccaaacgacgagcgtgacaccacgatgcctgtagcaatggcaacaacgttgcgcaaactattaactggcgaactacttactctagcttcccggcaacaattaatagactggatggaggcggataaagttgcaggaccacttctgcgctcggcccttccggctggctggtttattgctgataaatctggagccggtgagcgtgggtctcgcggtatcattgcagcactggggccagatggtaagccctcccgtatcgtagttatctacacgacggggagtcaggcaactatggatgaacgaaatagacagatcgctgagataggtgcctcactgattaagcattggtaactgtcagaccaagtttactcatatatactttagattgatttaaaacttcatttttaatttaaaaggatctaggtgaagatcctttttgataatctcatgaccaaaatcccttaacgtgagttttcgttccactgagcgtcagaccccgtagaaaagatcaaaggatcttcttgagatcctttttttctgcgcgtaatctgctgcttgcaaacaaaaaaaccaccgctaccagcggtggtttgtttgccggatcaagagctaccaactctttttccgaaggtaactggcttcagcagagcgcagataccaaatactgttcttctagtgtagccgtagttaggccaccacttcaagaactctgtagcaccgcctacatacctcgctctgctaatcctgttaccagtggctgctgccagtggcgataagtcgtgtcttaccgggttggactcaagacgatagttaccggataaggcgcagcggtcgggctgaacggggggttcgtgcacacagcccagcttggagcgaacgacctacaccgaactgagatacctacagcgtgagctatgagaaagcgccacgcttcccgaagggagaaaggcggacaggtatccggtaagcggcagggtcggaacaggagagcgcacgagggagcttccagggggaaacgcctggtatctttatagtcctgtcgggtttcgccacctctgacttgagcgtcgatttttgtgatgctcgtcaggggggcggagcctatggaaaaacgccagcaacgcggcctttttacggttcctggccttttgctggccttttgctcacatgttctttcctgcgttatcccctgattctgtggataaccgtattaccgcctttgagtgagctgataccgctcgccgcagccgaacgaccgagcgcagcgagtcagtgagcgaggaagcggaagagcgcccaatacgcaaaccgcctctccccgcgcgttggccgattcattaatgcagttgtcattttgtttcttctcgtacgagcttgctcctgatcagcctatctcgcagctgtgttaaaccgtctttaagtcaacccacagtctactgcaatcgtattcagaactagccactagactgttacaagtcgaaacctcagttaaccaactagaaactctagttaccaagttagaactgtgagtacgaaaagtctgaaaagcagaaagattcaatagatttgtctgtgttacacaagagttcaacaagtagctgcgttcgtgctagttgtctaggtttaaacgtttaaaaagacaactagtagtttacttcacctctgtattactgacggtttgtgcctcaacagtttacgttaacaaactagtaagcgtctacttcgtggacttgtacttttagtagaaagattgagtgtctggcagttttaaacctaagtcagttagaatctcaatcctctgcctttggtttagaaacagtgtagctgttttgacctagactgagtacaactgttcttgctgcttattcgaaacttgcctaactcactgcaacagacaagtctagtttagaataaagtaggctcagttattcaagtctaacttcagacagtgtttctcagatttgacttaagtagacagtagtgtgtgaaaaaccagactagtttgaatcttagcctgttcagaaaaaactaactgacttgtgtcaggcctcccaaaaaaactaacttcaatcctctgtactattccacaactattagttgttgcaagtctgaagcctagtcttgctttgaccttacaggatagaggttggctcacaactgagtctgtgtgaaaacaaaagtaaagattgccacaagaagtctattctagttagtccttttacgttggatagcaatagtccaatcttagtttgcagctgatgaatatcttgtggtagggtttgggaaaatcattcgagtttgatgtttttcttggtatttcccactcctcttcagagtacagaagattaagtgagaccttcgtttgtgcggatccccacacaccatagcttcaaaatgtttctactccttttttactcttccagattttctcggactccgcgcatcgccgtaccacttcaaaacacccaagcacagcatactaaatttcccctctttcttcctctagggtgtcgttaattacccgtactaaaggtttggaaaagaaaaaagagaccgcctcgtttctttttcttcgaaaaaggcaataaaaatttttatcacgtttctttttcttgaaaatttttttttttgatttttttctctttcgatgacctcccattgatatttaagttaataaacggtcttcaatttctcaagtttcagtttcatttttcttgttctattacaactttttttacttcttgctcattagaaagaaagcatagcaatctaatctaagggcggtgttgacaattaatcatcggcatagtatatcggcatagtataatacgacaaggtgaggaactaaaccatggccaagttgaccagtgccgttccggtgctcaccgcgcgcgacgtcgccggagcggtcgagttctggaccgaccggctcgggttctcccgggacttcgtggaggacgacttcgccggtgtggtccgggacgacgtgaccctgttcatcagcgcggtccaggaccaggtggtgccggacaacaccctggcctgggtgtgggtgcgcggcctggacgagctgtacgccgagtggtcggaggtcgtgtccacgaacttccgggacgcctccgggccggccatgaccgagatcggcgagcagccgtgggggcgggagttcgccctgcgcgacccggccggcaactgcgtgcacttcgtggccgaggagcaggactgacacgtccgacgcggcccgacgggtccgaggcctcggagatccgtcccccttttcctttgtcgatatcatgtaattagttatgtcacgcttacattcacgccctcccccacatccgctctaaccgaaaaggaaggagttagacaacctgaagtctaggtccctatttatttttttatagttatgttagtattaagaacgttatttatatttcaaatttttcttttttttctgtacagacgcgtgtacgcatgtaacattatactgaaaaccttgcttgagaaggttttgggacgctcgaaggctttaatttgcaagctggagaccaacatgtgagcaaaaggccagcaaaaggccaggaaccgtaaaaaggccgcgaattgtgagcggataacaatttcacacaggaaacagctatgaccatgattacgccaagcttgcatgccgcagaaaggcccacccgaaggtgagccaggtgattacatttgggccctcattagaaaaactcatcgagcatcaagtgaaactgcaatttattcatatcaggattatcaataccatatttttgaaaaagccgtttctgtaatgaaggagaaaactcaccgaggcagttccataggatggcaagatcctggtatcggtctgcgattccgactcgtccaacatcaatacaacctattaatttcccctcgtcaaaaataaggttatcaagtgagaaatcaccatgagtgacgactgaatccggtgagaatggcaaaagcttatgcatttctttccagacttgttcaacaggccagccattacgctcgtcatcaaaatcactcgcaccaaccaaaccgttattcattcgtgattgcgcctgagcgagacgaaatacgcgatcgccgttaaaaggacaattacaaacaggaatcgaatgcaaccggcgcaggaacactgccagcgcatcaacaatattttcacctgaatcaggatattcttctaatacctggaatgctgttttccctgggatcgcagtggtgagtaaccatgcatcatcaggagtacggataaaatgcttgatggtcggaagaggcataaattccgtcagccagtttagcctgaccatctcatctgtaacatcattggcaacgctacctttgccatgtttcagaaacaactctggcgcatcgggcttcccatacaatcgatagattgtcgcacctgattgcccgacattatcgcgagcccatttatacccatataaatcagcatccatgttggaatttaatcgcggcctcgagcaagacgtttcccgttgaatatggctcattttagcttccttagctcctgaaaatctcgataactcaaaaaatacgcccggtagtgatcttatttcattatggtgaaagttggaacctcttacgtgccgatcagcatgcagcaccaccaaaaaaaaacgaaaagtagaagacccacgatttatgtacccatacgatgttcctgactatgcgggtatgaaaaacatcaaaaaaaaccaggtaatgaacctgggtccgaactctaaactgctgaaagaatacaaatcccagctgatcgaactgaacatcgaacagttcgaagcaggtatcggtctgatcctgggtgatgcttacatccgttctcgtgatgaaggtaaaacctactgtatgcagttcgagtggaaaaacaaagcatacatggaccacgtatgtctgctgtacgatcagtgggtactgtccccgccgcacaaaaaagaacgtgttaaccacctgggtaacctggtaatcacctggggcgcccagactttcaaacaccaagctttcaacaaactggctaacctgttcatcgttaacaacaaaaaaaccatcccgaacaacctggttgaaaactacctgaccccgatgtctctggcatactggttcatggatgatggtggtaaatgggattacaacaaaaactctaccaacaaatcgatcgtactgaacacccagtctttcactttcgaagaagtagaatacctggttaagggtctgcgtaacaaattccaactgaactgttacgtaaaaatcaacaaaaacaaaccgatcatctacatcgattctatgtcttacctgatcttctacaacctgatcaaaccgtacctgatcccgcagatgatgtacaaactgccgaacactatctcctccgaaactttcctgaaataaccgcggcggccgccagcttgggcccgaacaaaaactcatctcagaagaggatctgaatagcgccgtcgaccatcatcatcatcatcattgagttttagccttagacatgactgttcctcagttcaagttgggcacttacgagaagaccggtcttgctagattctaatcaagaggatgtcagaatgccatttgcctgagagatgcaggcttcatttttgatacttttttatttgtaacctatatagtataggattttttttgtcattttgtttcttctcgtacgagcttgctcctgatcagcctatctcgcagctgatgaatatcttgtggtaggggtttgggaaaatcattcgagtttgatgtttttcttggtatttcccactcctcttcagagtacagaagattaagtgagccaaccgtgtaggtctacaaactggtattctcgtttgttgtgaatcttgtcttttctgcttactttctaatagattgagtgagttaggtctgtatttgacggctaaagttaaacttaactgttgtgtgggatagttgttagctgtcaagaaaccgttaaacgtagaaagagtaacttttctaaaccgtagtgtctactcaatattgagacaagcagctagtttaagagcagaagctgaagtctgtgagtttggaaactaaaaggtctgtgaacagctaactagaactaggcttgcttttagtattagaataacaacaaacgaggtctattagttgtgggatctagctagttgcctaagaacaaaggactaagtctgcagaaaaacgagtttcaataggtctgtgtgtaacagtgttaaggctactagcagaactgccacctaagtttcgtaagttccacctcagccttcaatattccagacttgaccaactttcagtttttaaactctaccgtactagtagtttctgcacagtattagacctacttacactgtctgtgcttgcaccacaaaccaaatagtagaatctcttttaagtagaacagtcttttcttgttagactaaaacaacgagagacaactgagtgtagggttacagttaaaaaactgtacagtctgctgtacagacgttacacggcagtcctgttccaaacttcacttaagggttagattggctaagttcaacttgtccttttaagttagttcagacctagttagcagacttaggtgggtaagctgagctacaagagttaaaaacaactgctaagactaaagtaagtgtacgtaagcttcaaactagtaagttaaacttagaaacagctaacgtctaaagctagaagtaagctcaatcttagtactgttaacctaagttgctcacgcaccttcgccaactcaacctgcaggtcgactctagagggattcggcgctccaactgtattctaggcctttccacaatttaaaaaaagacctccgcgacggggaattgaaccccggtctaccgcgcgacaagcggtggttctaccactaaactatcacggatttcgatgctgctttcccacttccttactttacattctctaggcgtttagcctgtagaataaaaccttttttcaagactacaatactgctgctagtaataccgaattattacatgttttaacaacttgagagcgggtcgtatcctgtcttccgattgtacaatctcatcttatcagctcatctcatctcttcggaatccccgggtaccgagctcgaattcactggccgtcgttttacaacgtcgtgactgggaaaaccctggcgttacccaacttaatcgccttgcagcacatcccgggtaccgtagatcaagtgacttcttgagcttgccaagcatcaagtcccaccgacaagtggtacaccagtcagcgttatctttccatacattctttaattcaaattgtcccattcacaactcaatgtttttgtttctctcgacgaaacgtttttgcttagcatttcggtatcttaaaccgtgtgattttaagtctcataaccttctacatacaggatcaataataatattattgttgatacaatataggattagaatacatagattctgatgttcgaatcaatcaatgtcttaatattcacgtacttgtataccctctgattccacattggtccaacggaatcttctagcaacagccagttgcactagatgtggttctccaccagaaattatagtactgacagattttttggtttacggtgagaggctccaatttccaaagctaaaccaatttcttcataagcaacctccacgcgcgcttttagtatgatgtagaaactggacctaaagatcgaccaatacttgcttacacggcagcgcgaaataaaagacattgataatgtagagtaagtactatttcttcaaaagatagaaccactctcagaggctaactagaagatctaaaacggatgtacagctaatatgtttcgaggcaatcatgttcaggtgcgcctcaagtggctcttaatctattttaaagcataatccacgaaaatttcgatacagagagatacgtcataaactgagaacgatagttagaggcatagcttcggcaatgttcgataaacaagtcaacgagtcagaatttcatgtttcttttttttcttgtagtaatatgtaataaccgaatcagataacttgccagtaatgacctgagctctataaattgaaaccgctacaaaaagtaagggatcgtgttaagggcacagagaagaaagtgtaaagaagtaagctagcgttgtacaacaactcaaaagcatggagcccggtgggtactagataataagaacaaggtttcagtgagtttacaagttcataaaacacggaacctggcaacatagcataccagtaaatcgatttgaagaactctattaggctggcagcctctctcaatcaatagtctaaaacggttaagagacacaatcatctctaggtgtcgttgtggtctaccatcgaaacactggaaatcaatccaatgaacttcaagtgatatatttttcaagagtcttcaaagaccaagcaatgttcttattgattgataattttaaagttgaagtctaaatttgtagcagtgaagtctgagcagggggaatgttaagccatatctaatagtacctattagacgcttgattgccgtttgcaaatatgccatgtttctttgaaaccaaaaccacgtaagggggcttaagttttcggcttaggattgttacggagctcaaaccaaataaggcagagagcatacaaaacgtttattaaaaagaacaactacttctggatagctcaattatcttttgtttcttttgaggcgtgcccttcatgacatgacattgcatccatacaattaatagtagataggggaggataagtttgctagtgtgagcatttagagcaaggggtcagtttcctccatctgcctactctgctccatttaaaagcgagccgattgccataagcttgctcacgtagataggacttcaaaaagacgttaagaggctgctctctagaatacgatggaataaacaaatctcgttagtttcttgaaacggaaagtacatgtaggataattgctggatcattcggaagctaaaattggtcctttccccaaagacaatcgactatagttcctgaagcttcctgaggtgagctggatagaaccagaagatagtacctataacgtcaacaacaactgaaactacagatggaagagctaggcatatccataggggatagccatagaaaactttgattgacgcagcaaatcgagacgttaacctgaagctgccataacttcaggaacttgaacacaagagtggcattaaaattcctatgttgcatttcagaaataccacaagtaaactgagcaattaacttgttcataccgacactacaagttaactacaaaccgagacccttatatgcagactgataatacagaatgatacgtatcactcctaagactaggaacatacagagcttttgagcttttggtattttcaagttatttgaaaaattaaatctttatactagggacgaggttcgtgacaaaagaagacaatcatgagactcaccgtcttgtcatctacaagtttcaataagcttccaacttgagtaacctagccatgtgattagtaactttcgaagcatgattcagaacgttctgctctgccgcgtcaaaaagggcagctacttgaagttaaaagaaagatcaaaaattcctgagcattgttcaaacagctataaatcaaaacagaataagaaataaggccatcacttaccaaggaaaaacaaaaaagttcgagagatcaacaatgcacttcagggacgcacctgaacctttacattagtgtaaaaataaattattagattgttcggttcgagcgttagccatatacaaaatgcggccttcaaagtcattggaaaagagctgctttaggacctctccaaaagctaaattgacaaacaggctacctcatatctccatcatctttgccaaccttttcagtaacaaaaaacaaaagcaatatcgtttatttgtccattggacgatcatcaaatattgctaggtaggcacttttttcggtaagcgtgtctgcctcctttcttcagtaagatcaattgtcgccggatactatgaatcgatcattgtagaaactaacatgcttctaattacccattcaatcttgtagcactgagtgcatctcgcgtgagacaaattgaactcgcctacttgggtaggataaacggttgggtaacgtagctgcttcatccggctacttgcttggtctgctgcgtgctttacgcgctgattggtggtcgaataaaccggtacattatttgttcttatccacgacaatgagacgaacaagtaacaaaccagaactggtagctcgatgattacagttccacaatatcagccgaatacccacactttttccaacctgtgtcaattgccttagaaattgtattagcacaaaaatcatttcgctgacggctgaacatccgatatttactattcgtcaaggtatttagaattcttgtgcgaacttgtgggtcgtggtgaaaaagactattgtgagttgttcgtagctgagtccttgactatactgatacacattaaaagaagctgccctttaagcaactaatgtatttgttatgcgtatatctgtagaaagcaacagtgcgacagttgaagtgttcgaacacacttctcaattttgaatctttctcattctcattcagtgaacgatgaaccaagtacacaaatttggaggtgaataattatgctagggagagcttgctagatcatttcacaattagaatgtgaaaccagacctttagatgtaaatgctaaggctaggataggatcgtaattagtaggatcaacaaactttattgattagcctcacctagactaagtttgatcttttcttcatcgtccaatggacaaataaacgatattgcttttgttttttgttactgaaaaggttggcaaagatgatggagatatgaggtagcctgtttgtcaatttagcttttggagaggtcctaaagcagctcttttccaatgactttgaaggccgcattttgtatatggctaacgctcgaaccgaacaatctaataatttatttttacactaatgtaaaggttcaggtgcgtccctgaagtgcattgttgatctctcgaacttttttgtttttccttggtaagtgatggccttatttcttattctgttttgatttatagctgtttgaacaatgctcaggaatttttgatctttcttttaacttcaagtagctgccctttttgacgcggcagagcagaacgttctgaatcatgcttcgaaagttactaatcacatggctaggttactcaagttggaagcttattgaaacttgtagatgacaagacggtgagtctcatgattgtcttcttttgtcacgaacctcgtccctagtataaagatttaatttttcaaataacttgaaaataccaaaagctcaaaagctctgtatgttcctagtcttaggagtgatacgtatcattctgtattatcagtctgcatataagggtctcggtttgtagttaacttgtagtgtcggtatgaacaagttaattgctcagtttacttgtggtatttctgaaatgcaacataggaattttaatgccactcttgtgttcaagttcctgaagttatggcagcttcaggttaacgtctcgatttgctgcgtcaatcaaagttttctatggctatcccctatggatatgcctagctcttccatctgtagtttcagttgttgttgacgttataggtactatcttctggttctatccagctcacctcaggaagcttcaggaactatagtcgattgtctttggggaaaggaccaattttagcttccgaatgatccagcaattatcctacatgtactttccgtttcaagaaactaacgagatttgtttattccatcgtattctagagagcagcctcttaacgtctttttgaagtcctatctacgtgagcaagcttatggcaatcggctcgcttttaaatggagcagagtaggcagatggaggaaactgaccccttgctctaaatgctcacactagcaaacttatcctcccctatctactattaattgtatggatgcaatgtcatgtcatgaagggcacgcctcaaaagaaacaaaagataattgagctatccagaagtagttgttctttttaataaacgttttgtatgctctctgccttatttggtttgagctccgtaacaatcctaagccgaaaacttaagcccccttacgtggttttggtttcaaagaaacatggcatatttgcaaacggcaatcaagcgtctaataggtactattagatatggcttaacattccccctgctcagacttcactgctacaaatttagacttcaactttaaaattatcaatcaataagaacattgcttggtctttgaagactcttgaaaaatatatcacttgaagttcattggattgatttccagtgtttcgatggtagaccacaacgacacctagagatgattgtgtctcttaaccgttttagactattgattgagagaggctgccagcctaatagagttcttcaaatcgatttactggtatgctatgttgccaggttccgtgttttatgaacttgtaaactcactgaaaccttgttcttattatctagtacccaccgggctccatgcttttgagttgttgtacaacgctagcttacttctttacactttcttctctgtgcccttaacacgatcccttactttttgtagcggtttcaatttatagagctcaggtcattactggcaagttatctgattcggttattacatattactacaagaaaaaaagaaacatgaaattctgactcgttgacttgtttatcgaacattgccgaagctatgcctctaactatcgttctcagtttatgacgtatctctctgtatcgaaattttcgtggattatgctttaaaatagattaagagccacttgaggcgcacctgaacatgattgcctcgaaacatattagctgtacatccgttttagatcttctagttagcctctgagagtggttctatcttttgaagaaatagtacttactctacattatcaatgtcttttatttcgcgctgccgtgtaagcaagtattggtcgatctttaggtccagtttctacatcatactaaaagcgcgcgtggaggttgcttatgaagaaattggtttagctttggaaattggagcctctcaccgtaaaccaaaaaatctgtcagtactataatttctggtggagaaccacatctagtgcaactggctgttgctagaagattccgttggaccaatgtggaatcagagggtatacaagtacgtgaatattaagacattgattgattcgaacatcagaatctatgtattctaatcctatattgtatcaacaataatattattattgatcctgtatgtaaaaggttatgagacttaaaatcacacggtttaagataccgaaatgctaagcaaaaacgtttcgtcgagagaaacaaaaacattgagttgcgttataaaatagatagtaacagacttgctaactgagcattcaagactacgttctatttcaggggcttattcaagctgacggagcaagtgtgttgatgcttatatatctgtcatgagtttgtaatccccatctttttgatcacagttgctctaatgataggtgagtattctaactagacaccaagaacgagtcttcaagcgtacaaatccacgatctacaaatgtcgacgaaagtggttctgttaaaacgtgaccaagcttaggttgtctagtttgaaccattggaatcaagggcccgaacttcgttcaaagagacactcatcctcttgattccagtagaattgagattgacaaagtcgttgtctactgattcttaaaacattggcaaattggtttggcaagtttaaaaaaggtacatctcttatattgagcagttgcagttgaatactgaaacacacctgaaaggcccgtcgatatgaaccgcaaatactacttttcgaaagttaaataacaatataccataatcatgtcagggcatcaatagattcgttggttaaagctcatctcttttcttgatatccaccaattctgccttgaaaactagcggagttagtatcttttttcaaatcaatccaaataatagactcacctagtacagcatttgctggaattgggcctattccgcggctgccgtaacctaaatcaggaggaattgttagctttctctgctctccaatacacatgcttaatattagtattcagtaaaaaagttgagaaagaaaagaagctcaaaatcacgtacttttgtaaaccttgatcccaacctctgataacttgtccagatcctaaaacaaactccaatggttgatcacgttcataagatgaatcaaaaatggtgccgtcctccagagacccttcatagtgtatagcaactgtgtctcctgattgagtcttttggacacactcatctggtggtactttcctagtgattcctaaaaacacttattagcacacgtaattgtggaactcaagtaactacgtaccaatccttaattgacctgaatcaacagcgcaaacgagtctaactaataagaacacagccagaaatttggtcgtagaaactttcatggcgtggcggcgttcttgtcaagttggcgagaatacatggctcgctccgtctcctcagaagcaattcttcgcgactacagaaataagcattgtcactgaacattttagaccagaagttagaattgcatatttacctacagttattctacaacgattcgcagagatgagttccattgtgacgtccactactatgataaactgatttcgcaatggctttcactactagcttgagtaaatgtatcagcctgttaaccatttacgctcacttgcatcaaaaggaatcgccaatacacaaaggacttctcctaatttatatggtacttgcattgcttcaacatcatcatcaactattctttcggttttcatgttcgttcgctcacaagttaaatattcttgactcttttaaattctggaaagattccattaaaaaattcatcaaaactatcaaactgcgacaaatcgggcatttcaaaacttgattcctcgtgaggaaactcagggttcgtagaaaagtgacttggtgacaaaccgagattaggaactttgcaaacactttctgtaggaattgccgcttgtaacatctgttcctgaaaatccacagcggaagagccgttccgtatcttatgagaataaagattaacagtcggcgataaactctctgaatccttctcatctaactcatcagtcctaaagtcagcatgcttttgtgatggtgtactaactttgcttgctttgtctacaatgttcttgagaattttacccacagagtcggttgcttcatttgatccttctcctatctcgttcaaaaaccaatatactttagaaatcaaattcaaggaaaaagatttctgttcttgattcattataggtaaaaactcaaataatgataaacttgcataaataatttgagaccattggaacgaaggggatccccgaatcaaagtagttggtagcttcaaaagtctttctaaaattttcatcgaaaatttcgtaactgtgttgaatgcatatactaggttaggaagtctatgagaatccctgaaaacctgagttatggtacttttcttcggtgtgcacaacttgatgtatcttctcgataataccacgtaaagaaattcgtaagtaaaagaaagcaaaccactaatatctcgatcaatcaatgctatccaccgagttttccagtcgtccaaatcataaaatacgaatattgcattgtcgtcatgtgtaaacgtcttggtccagcccggtgactcgaatagtcgttgaaaaattaggtaaagctggagctcactaactttgatcgcatccccaatagtagccttcggatacatgatagtaagcttgtgtaccgagtaaaaatcgttagtgatcattattgggcgtcctgatccaatggaatgctgaaaatggaccgcacacatggcattccatattcgcagatggtacaagtcgtcggcttcaaacttcttctccttttcaactctggtaacaatctccttgaaattgaaagatatcatcaaatgtttgattccgattccagataacaaccatccatctagttgaggctgcttgggcttgcgggcaacgttgtaaagacaaaggtacaatattccttcaatatcatggtcagtaagtggctcagttatggcaactacaccaagcagctttgatgtaagccagtacaattgtttttgctgcgaagtataagtataacgttcatcaaagcacatcgccaaggcaacaaaaacacatgtgatgaatgggtgctctaacacgtactctctgttgatctcagtaatgcctcctggtataatccaaaaatgagcaacctcaaagtagagtttgaccagttccagacaaagtttctcgttttcaaggtagaacttaagaaactcctcacacgctaactgtaaaggatcatacctcgtgactcttttgaaaagagagtggtctatttgtttgataatttgcagtggagaatcttctataaactgatccagataatcctctgtccaagatatgttcttcgtgaagttcaaagtgtcgataaaagactcttcgcgtcgtctcataggtggactcaatctctcagcgtgattttctagcttccagagcactgaggtcaattgggttctcaattcatctattgctatttgcattttttctaatctcaagtccatgttaaaagtttctggcttcattgtgtttcgaatggccttaatttcggtttcattttcttgggtcaaggaacagctaagtcgaagcttagcgcacctatgacatttgccaagctctagttcaaaatcacaccttgtcttaagctttcgacatgattggcaactgagggctctccttcttttcctttctttacaagggtttgaggttttttccacgctgttagtgggttgaagcggttgatgtaatgatccttcagcatgatctttgccatcaactccagaattccgatcgaaactcagaaccttacggtcattttttgctcctctatattcattgtgctgattggtttgggccctatccgtatctctctcttgatccttttcaaacctggccgcttcttcaacaaagcttctgctgctcatggatgctaagagagcatcatgccgtgttgaatgcctcttaaatgaaatattatctgcactatgctttgcttcattttcatctagaagcttacggtgttccatgctggaggttttcgtctatgcaggtctagcctttagcctttgaccatatctaccagtttgggagatcgatttcttttagtgttgggataacatttcgcggacaaagcaacatttatctaaatagaaagactacttaatttataacccaggaatcatcagcagggccataaaaagggaaaacaggctgtattttagatttgcggcatttgatttggaagagaaagagttaatatcggttatattcttcacgactctaccctcttcgtttacatgaatggtctgtgaatgtttgttcgagaagcattcgatcactccccctcgaacgattgaggaggccccgaacttcagccaattattacagtcgaatgttccagatgaggtacgaaccaaggctgaccctctaaccacctgcaactcttctagctggatatctgaaaattcgccttcaaaacgaattgcacccccaataccttttagttcagggaaaaaactgatactctgaattcttgtgttattagcaataatgaggcctccttcaatttcgctaagttctggaaaatctaaaatctgaaccatatcattcttcataataccaaatgttccccccacatgttgtagtttcggtaaatgtaattctttgaaggagttttctatgaatcctgccgaaacttccacatttttcaagttccttagatcaactgattcagcatctctgactgtaatatttttagcattctttaagtttggaagatttatttgaacgtttttggcattggaagtgattgaaagctcgccagtaatctcttctacgtcagattccacaagttccatatacctattattgttgatattcaagacgttaaggctctgtgtatcaaatcctgtaaagtgtgtcaacgctgtatctgaaataataatggtttcaagcgagctgacctcactattgaacgaaacgctactcagaatgggcagaacctgccaatttatggactgcatagtttttagccgcggtaaacgaatagttgacaatgcggttaaaaacagaaaggaggaactaccgccgatgcgttccagttctaagccctcaacccttgccacattggagctgttttgaaccttgatatctccccagacatgttttgtggcgccaaaatctaacacggtgtcggtgtatccatcgaccacaatatctccctctattgttgtgcaatcttcaagctgtttcatatctccttcgttgcggatgatatgctcaggttttgaacacttttgagggatgcttttgcttttaggtcgttccccagtcgacactttggtgtccaatacagccaacaaatctctgcttggagttgttgccttttgtgggtaatcttcaaaggccttgttcccagtatgtttcgtcatcgaaaaagcattgatcttaaagcagatagaagagatgactaagatatagatgactataggtctcatctcaacaattgctttgattgtttcactaatagaaaattttctagatgattaatagactgcgttaggtgcgacaatcctgtttcgaagatacagagtacaaaagcaagcaaatcttgaagcggtccgaacactactatagcagcaggtacctagtcatgccataaccgtagtgcttgtcagtattatacgcttcagtatttgtttaagatgtagccaatatgttcaggaacatttataacagttgagttgatcaaaatatactcgcgatgggattcgaagcacaatcttttgacagtaagacaatcgctttgaccattacgccacacagataataaagttcaagcaaagtgtacaggtgtactttaccgcagcttttgttgctatactcattattctcccttttactttgctgaaaaaacgacacaataaactccatataggcctaatatttgatagggtggaagtcaaaagagaaaaaaggagcaacagctctgcactattaacaattggaaatgaaaaagtgaaaaaatgacagcaattaacttttacaagctctttgagtacaaaataagttcacctaagacggttttagtgttttttcacaataagaagatttggaagttatcaacggcaactaatttatttagctgtaaatcatcaaggaagtttaaagtttggagttcaatctttataactaccaatcaagagttgtataagtactatagtcctttgtttgtctcagaaatcaagttcagtcggtatacgatattcttcaaacatgacctgccatctcgtttattgcatcaattggtaaaaatacatctagttatattcacttatacattagaaaacccaaatttctctacattctagtctcccaaagcatctcccttaatatcaccgttatccgatacatcgtctgacaatttctgcaatctagcagcattcactaaaggaatcaacgaggatatagggtcagcattgagtatgaaaggatgctgcagcaactggatggcatccgcacgtcgttcaggattaacctctaaacatgattgaaggaacgacttcaaaccgccactcaaaatgtctggctcttccaatttaggagttccgttagtagttattaaaaacaaagccctcaaaggtgtctcattcaagtaaggaggttcaccttcaatcatttcaataaccattattcctaaagaccagaggtcgactttaggtccatattcttttctggacacaacctctggagccatccaataaggagtacctaccattgttgtacgctttaagttgtattccttgatctgcgcgcaaaacccaaaatctgtgatcttaatttctccacttagtgaaagtaaaatgttatcagacttgatatcgcgatgaatgatgcctttagaatgcaggaacaacaaaccttctaacgtctctttacaaacggcaccgatctgcccttcagacataacactatgcgtgacaatgtccgtcagggatccaccttccatatactccatcacaacccacaggttgttgtcctgaacataggaattaataaagttcacaatattcttgtgtttggatcctttcattaccaagatttcgtttattattagttctttcttgggctgcttttgtaagtacatcctttttatagccacgcaagaccgtgttcctatttcattcgcagtataaacatcaccggaagctcctttcccaactttgacaaagttttcgtacttagcagtagggtcacctgcagagcagatgctattcaacttattgagtatcaatttgtctctacatttcttttcttcctgttttcgttcactgcgttgtttaacttcctttgagctcctttgcttgcccgaattcgcacccaacggggttggaggtactttttctgttctgggtgacaatctaacaactggagggggtggagccggtctgttcggcctcattatacgagtgttggatagtggctgttcttgctgttgctgctggtttgattgttcgtgttccagctgctgttgcggctgcagattttgttgtgtgaactgagactgccccttaggaaccaaaggagatccaatgacaacatttcttggtgaggaagaggatgattttgtagacttggttgagatcttacgcgataaagagctaataaatgaacttggagaaggatgtacatttggcacgccattgtagtgtagacccagtctaggagaccctgaagttgcgaacgttgatgcagattttgtaatgggggttttggaagacggcggcttcggaggaggacgagaaggaatgaaagtcgaatcttctggaggaggatatatctgagcaggagacgccaactcgaaactagaactaaattttcccttatgggatagaacgggagtcgacagccttttcaagctagtctcttggattcccaaagcttcataattatcagaaatttcttcgttggaagtagaatgagtcgaatctaaactgtgtttggaggcttggtcaatgttaaactgcatgaaacgatcctcttcagtgttcatgttattcttgtaaaagttaacgatatccattactgcctgagggttagactgctgctccgtctttgagatcccactatcgcttaggacgcgttgccaggattctggaagtccttgaaatgatccagtggaattatcatactttacagattgcacatgcttgtaaccataaggtgtggaaatgctcaaagaaggagaatcagttgaatgttgaattaccccgttcaagttcaaactcctagattgagatcccaaataattttcacccttgattgagctcacgaagctacttaggacccctttcactttgtgctttctactaggaggccgagaacttgaagttaccctctgtttagtgggtgagagtgtttttgaggtatggtgtggtgaatatactggcgtagaatgaatcatggtttcctgcacatttgtatttgtgggtgaagcctgtaatgcacgtgattggaaggaagatttagaggaagggtcgttgtgaacttctgcaaagtcggccccagtagaattcgaagagtgatcgaaagtacccacttgtgaggactctttatcaaagtttttctctataccctcctttggttccgtaatattgttgagttcctttaatactgtttctggaagggatctttgaagagatttctgcctaaaatcaaactcctgatttccatcgttcggcgcactgcctggaacgggcatcacagagctgtcttggaacgtgtcattagatctaagaactggggtcaaaatttccaactcattatcataacatattttgttgagcttgtgaactacatcctccttgaaggcagagtcaggctggatcttgaatacctgttcttcagctaacgcattcacattcagcgtgccaggaaaacgtgggaggtctgcgtttaaagttgatgaagctgaaagttctggctgatcagcaactgaatcaggttccgccacgagggagttgtgtgacatgtaattcggatggttagttttttgcagagattaccatattcctgatacttattcgatcacgactgcgaagatcacacataggaagaaactaaagtgagtttagcactactgattacgtaagcaagattttgcaggtatataacatggctgaagacagttcgtgcgctcatttcggtttagaaagagagtaggcgattcactttttttcaatcagtcctttgctagtgaatactcgtggactgaatgctgctacccataaatcttgcatgtgcttagaaatttttatgctaatggcatccaataaccatgtgtaggaatggttacactctatttacaaacaaactatcagaaaaccctacataagctgcctgtattgctggactttctcgagacgccgacggcgtcctagttcaccagcttttgagaggaccccaagaatcaaatcagttgatcgattagaatcatcattgcctggaactggataagtaaccagtgacggctctgagtccgtgtccacgatgccaatcgtaggaacacgagcgtacattgcttctttcaaagcattcctgttctcaacagggtttaaaattacaataaggtcaggttttacaagggaacttaattcatcaccttccaaatttctaccagtaggagtgtcttctaggtctacctcatgtctctgccaggtgccactgatttcagtgcagttggtcaaagtcccgggtatccatcttgaggacacaaagtaacccttcactctacttgcagcttcctctaaagatcttctttgtccatctctggttcccaaaaacaaaatcaagccgcctttctcagcaacgccttgaacaacttttgaagctctctttaagtatatcaatgtctgttccaagtctataatgtgtaatcctttgtactcaccatatatgaagggttgcgtggatggcctataaagggatgtcgaatgtcctaaatggacccctgcagccagtaaagttttgatagacaactcatttgctttcttaggattcaaaacatcttcatggggtttgtaaacgttagttaaagaagcaccgagagaatttacctgctgtgaatgatggtattggcgtacaaatagttcttgtatcgaatattctttatttggaggggtggagttcaagaagggaaatttgttactgtttcttacgttggtcccaatgttagaacttccaacattcaatttgatttcttccttgttcagaaattcatccaaaaagtccttaagcatattcctgaaatctacgtggtccttctggtccaaattgtactctgtcaataagtcatttgcttgttcttctgaaagggatccaagagctcgtattttgcctcgtatttcatcatacttcaggcgctgatcaatgtctttatagctatgggatttatcaatgtccgaagagatactatatggtcttttaaaggcaaaagaccttagcacaccgattcttcgaaccaatgacatgttgctcataccgctgttgcagaaagtaaaattttcagcgaacccatattctaagcggtagaaaatgggaaaaagaatgctaccttcctctttaagctttattagctacatttttaaagaagtagtaaaaaaaaaaaaaaaaaaacaaaaaaacaaacgcagttaacgatctcaaaacacaaatgtcctacgagcagcaaaacaatcaacaacttgaccagctggctcacaaacttaacagttttaggagggtgacccaagacattagttttcaggctaacgaagatagctctacaatggattcgttatcgaacggcttcagctcattggcaagttctgtacgatcgacctcttcaaggtttgctagggttattagtcagaaccagagtatcagtagggtcataggattgtctcttctggtattcataatattttggacactgtttaaactgctatgactattcttttcttacatactggcggatactgctattctcttccacctccaatgtcttgccctagtttataagttccaatagactttccccattgaacttttgatctgttaattggaagctgcctacgcagattcctaaacacaatttcgttcaagttattgaaactactcaataggtttctttggtaatcatcctcaaaagatttgatgtgattgacaatgtctgtaaccgtagtttctgtgggattaagtcttttgtgactattgagtgtgacattcccgtcctcatagtagtggatttttattcttacctgtcccgaaattaagcctttgaccctgttaaatgtatagaacgattgccactttccgttgtaaaaattcttatcattgtaccggacgtccacaataatgacatggaatccttcctcaaacggagtaagcactaccttgtaggagtctgggaaatgctcttgggtgtacttctccagctccttgtcaacagaattatactcagggagaatatcttcgctaaattcttcagtatccatagacttctggtctacaaaatcatagtggaacttcaggttggatgcatagtctataaatttagtacctctttgagtatacctggaaactatagctaacaatccttgagaggtatcctgaaccaaatgaaaatcagaattgaggacatgtgtttcaaattcatttttgatttccgcaccaaattccgcagcaaccagtttcaagtctcggattacaggtttggcttccccaggaggaatgtcttggacaaattcagagattatggacatagatgaattggagtatgttttgaaatctagttcgatcatgccatatcgcgcttcgggctcgatttctaactttgaattgatagaagaatcagacaagcgcgactacactaccactgttgccgccgccccctttcaattggttctttcattttgggtaactatgccttccaagttggaatcatgtattataaaagcaactgatgagaaactgacaagtgacaactggggttatataattgaagtatgtgatactattaacgatgagcctgagacaggaccagcgactgctattatttacatcaacaagcgacttagcataaaggatgccaatgttctcttacgctcattgtcattgattatcgcaatggccgaaaattgtggttcaagaatgaagcaggctattgctaccaagggcttcatttcaacttttgtgaaattaatagaagactccagaattcaccacacaataaagcttaagattgctaacatgctacatcaactttcagaatcgttcatcgatgatccttcacttgcaataattgataagacttgctcgaggttaagatctcagtaccctgatctatttgctccagctccttccaagccttctaagagagagatttcacaggacgatagacagagagaacaggagatgcttgacagggctctcaaactttctttggaggaatacaatagaactgaatctccgccgttgaaaaaaaacgaaacattgaaatcaaaccccgagtttgtaacggtggataactcgtataagtggtcctcagcagattctcaccagaagctcgatgagaaagtagacagccatcctaagtttaaatcctccagtgtaacttctgtagaggataatgaacctctcagtgaaatagtgaatacaggtagtatagcaaaggtcgctaaagtcagggctttgtatgacttggtgagtaatgaagctggtgagctctcttttcacagaggtgatatcataaccgttctcgctagtgtttacaaagactggtggaaaggttctctgaaaggcaaagtaggaatttttcctctcaactatgttacaccaatcaaagaattaaccgttcatgaagcgatggaagaagccgctaaagagtatcgtatctggaaacaatctagacagattgacctctttctcaataaactgactgaagtacatgcccttgtgacaaatactggcaactacgatgccctcaacgatcttctcgaagatgaaaccatatccaaattatacaatagtcttacacccttgagacctcagctgacacgactgatcgagaagtatagtaacagaaaagaagaaatgttgtctttaaacgacaaacttatcaaaagcgagcggctctattcaaacttgttggaggcttctgtctctcggtttaagagtgcacatgattcagcctactccaacgatgatccacgacaagcttatagttgaagtataaatatacaatctgacttatttacaagagataattgttttagttgtacagcaattgatggctcgctatttatccaccaccaactacttcttggagtttctcttcttgcttgcaatatttcctcccaatttctgttcaacgtgattgctgccaatcaaatcaagaaagtcgttctcgctctgtgtaccttggacagtatcttcttggctattgtcggcaaatcccgagtatctctgcttgaatttgttcaaacgaccccttgtgcctgtgtcacccgaaactaaatcttttctactactattccagaaaggactatttctctggtcagaaatcattcgcatttccaccttggggtactgtgatttgcgggtaattactgatccatctgataattctacttgacaatcgaacttataataaattgcaggtcttgccttaccgatttgaattttcttcaaaggtctattggggatagaaactccttgttctaaggagggtctactcgcaaatcttttcaatgccaagttcacacccttcaacatagttacttgatgaagtaaaaagttaactatatatagttttgagttttggctattaggtcttgaaaaaattgtttctttcgcgaaattgttcagcttctgaaaccaaaagtaaatgaaccggggcacccatctagaggcgtaaacaattactgtcaacatcactggaaggaacaatttttgttcagttcagataggtagactcagttgtattggaacctaagtcaaaagacattaaatcctcttgcttaatgattccattaagttcagatactttggatacttcattagagttgtacaattccagcttcgttaataaatgtcgccgcccttgtaagagcttttcaggaagttcaccagtgaaacttatttcaggcccaggaaagttgtgctcggtagatagtccttcacattcttctctatcattttcttcctcctcttcctcaagctgttctaggctatcttcgtcaaccccttgtttagttgaaaagttgtacgcaacccggtctactatgttctctactttattcaagtttacctcctttccttgacttaaagcatctagcttatactttataattgagcgaactttttggagtgacaatctatcaaaccacccataacgctcagaaggcttctctaatacagaagcagcgtgaagtatgttggtaatcaaagggtcctgaagagcgtgcacctggaaggttacaactgatgggaaggtatcaccagtgaaaacctgatcttttggaataaatgcaaattttttttcatcttctgtcaagttattgtagtctagaaagtttaggggaactaactgggactcagagatgaaattagaaggttggtgttccagatctgcaacggcagtgaagctcctctctacagttcggaaagtctctatttggtatacttttgagctcgggtctaggcgaggatcatatccgaaaactataggcgaagatctccatggtccttttctcattagatacgaaacatatggcagagcaaacttgatactagacttaaatctgtcagggaacagcttaaccaaattacttctaagccagataggacgaaccctaaacgccacattcaatagctctatcgtatccaaaagtgtctgagcaggagattcccggatgaattcttcactgtttctcttgtcaatgcgcatttgctgcacctcttttctcaaatcttctaaattgttcaacagttcattaggaggacccagtggcgtagcttcgccgaaatctatactagtactaaacaataaaattctttttgatcgagtctgtaaagatatttgtccgttctcatccactatagttgatgagtacgggttggacttgtagccataattgaaaggcaatgaaactctgttgaatctcaaaggagggggtaaatccaaatcatcgttgttttgctgtctgcttatatcattttcgatatcactggcaatgcccttaatattaggcagatatccccttaatagcccttgtctgaacttacttgcaaatgcagaactcttcgtgatatactgaaaatctgccaactccctaaatttgaagttcttattcaagactccaataacgttgaactcagtactaaagccattctgcttgcagtgtcgcaatgtagctgctatgttgtttccatttctctttcgaaataccttcctagggactcgtacctttaccaatagattctcattctgatgaatgtaagagttgattggatgatgatggggatcttctctcagtcgaagctcaagagttgcttcattaagagcgttgccctgttgtcctaaacttagaaaagggcgattctccaatatgtcgttggatatgatacatttctgaagtttttctttcccgccaaccatctccagtgcccttgccgagttattaaccaccaacggaaattctactgacgtgagcttcggcaggtctagtaggtatttcttattttcagacattgaaacgtttttgcaattctttgaagatagtctagatccttgcttgacctgctcttaataaacaaataaacagaagcgcgaatatagcttaatagccagttgatacagaaactccaaattggaaaatatcaatttatcaagtatactgccaaagaccaaaagcactatcacagattcaaaagccaatgcgtgcaatgtcatataattagcgagatcttctataagctttagacgcacttttggtaggttttgtttgacgtactcggagcatcccaaacgagcgcgaattaagcttccatcagtgcttcattttatacgctaataattacacttcgttggtcaaggcattactatgtcagaaataagaaaatcaaagagggctactgccggcaaaagagcgcttacactggatgaggagttaatagtcgaagagagaaaaaagcaaaatttgaaacgaaattttctttccaggaaatctgagttaactcggcgaggtgaaacctacggaccaaagcagaggagaagaaaaccaaaacatacatcctcggaaatggaggatttagaagacgaagtcagatgcttagtttgtggtacaacggaagatgatggcgaagcaatggtccagtgcgatcgttgtcacacctggcagcataaccattgtatgtttcaggagaatactattcctgaatcatatatttgtaacgtatgcgacccccataatgaagcttatggtaacttgaaattcgaaatgagtttaagtgagtataaacaactcaaaagagagactctttccccgcatggtaagggaccagcactcaactttgatattcaagagtctattgttgcggctaagaaagcaccagatagtgattattcagaagagtcaatagcaaaaattcctgaggatgatagtgagagttttttttctgaggagagcgaccaggaacatttaattcactcaaaaccaacacaaaaaggtcacaacacaaaacacactaaagctttcgggcaagagcaagaaggttgcgagatcaagagcatggatgaggagtttaatcctgtaagaaagggaatagtgaaaacattctttaaaatgttcaacaatttgttaccttcgaacgctaagaatgccgaagaatgggctctgaaacttgaataccaattactgctttctcgtggagacaagaaaactggcaaaggtggcaagatagtacacattcctgggaacagctatacttctaaagctcgaactcttatgttcaatcttaaaaacccaaagactaaactcttagaacggctcattggcaaagagctaacatttgttcaactatgcaatttgactcccgaagagatgcaaacctctgacatcaagcaacttgccaaggatgtccacacgaaggctataagtcaaacagttatagagaatggaatgaactccaaacctcgagcgaagataactcacaaaggagaggaaattatagattttggtgaggcagatgaagggaaaacaagtgaactgtttaatgaagatcaattttctgattatgaaaggggagaagaagatagcccgcaacgaatgcagcatctcatcgaagaagaaggttctcatcagagagaggaggagcaggaaaaattgtctgaacggggagagccatataataagaatgtttctctatatcacaatgagtactattctggagaagaagatgagggaattaatcatgaacgccagcaaatgatcctcgatgagagcattagtgccatcataggagacaatttccaagatgaagatcagcctattgatgaagctggtattactgatagaagtaagggttttgtttgggctggtgaactaatattttcagagatttcaaatttcagagcatgtgcttactttgtaacttcttcaacgttaatggattttgacaaagaaagccaacaatgtatttcaatcataagacagtttcatcaaatccataagtatcttcaaattgagggtaggctcaaccgtaaaacagcgctcgcatacctcaaaaattgtatggaccgtaagaaatacaccctcatagttgaatttgttccagacgaatcaggagaagcattaacagatgaagagcaccgaaataacttcataaggttgtttaattacttctacaatcgtaacagatttggagtgatacaaggcaagggggattctgttaaagatgcatacttgataccttaccaggagggagatgtgccagaaaaagtttttgggtcgttcaagtttccgaatactatatttgaagttcgaagaagctcgttgtctcaaagactttttggagtgtttgtaaccaacttataatatcaacaagcatcaattcttatcctgatcatactatactaaaatgcttctagaacccatacacactagtagccctatttagcagcactaccgatagcaaggcttctttgtttttgcaaaagctgcacttgaaatctctcaaagtcgttcagctcacttcttctctgcttttgagcaatatccttagcccagtcagaagaggcccattgagaagagatatcagaagcatcccatgcctccttgacagcatcggagttgtctcccctggaaacggaatccaacactaagaaggttagttctatgtcttgaagacgagcaacttgtctttcaacatcagtagatggtccatcaacgagaactctcttctggtcgatgatgtcgacaatagcagcaagcttgttgttgatcaaaacaactctaccgacttccacgagtctccagttcaattcttcaacttcaacggttggggcggacatctttactttggttgatagaagaaggacaataattttttttgttgtactgcaaaaaacaagcgcgcaagaaccaatcacatgaccagggttgagccctagagaatattcgctttgagagtctcgaatctcgtttcaataagtgtttgttctacaggaaaaaagacgatgttcaggctcgtgtgattgtcagagtcttttagtaagttgctaatctatttttatctttcttatgatgaaataatttgcgattgagagaaaaggttcgaaacatgcctagtttcaatgtgttctcttttcgaaatgcgataaatttctttctgaataaaggatggctttttattctagaacttgattatctaactttaaaggtaatcagttttgcccctatgatcgatgtctagcaatgatgacgacaaggttgatttcaagatgattatggaaggtataaataaaaggaaaaccctggtggtattgatatgtgtgaattgcccctcagttgctatagttgataccattgaggaacatctgatttccggtggagctaagtttctatttttgaagccagatgaagtgactgccttatacttggtgctagttgcgcttcaggacttacgtcaggttcacagtgaatctttcaatgagcttgttatctatggcattgatagttgttacatagactctgtgcgcctggtaaagttgttgagccaagttgctctgctgggtggtagtttggcaggtccaaagaagtgcaaggcagaaatgacactaagtaagctaaatattcgcaaaatgagtagcctgtactaagatcgttaccaaaaacagagctacgagcttctctgctcgggtagatggtccaaatgatcacgatgacctgttaggctattgataataatctcacaagtgggagacgcagcgtttatctccggagatcgtccaagcgtgcctgtaagtcgtcgtcttccttaactgcgactctagctcctgtttgtccttctgtaacgacttctccaggaacatcgatcaaattattacttaaatcaaccccaatttcgtccagtaccttgctgattatttcgtctacctcctcttcgtcttcatcatctaactcttcatccattaaatcatccagagagtcatccatcatttcctctctctggctcatgatatccgtttgcctagagaattcttgagaaatatgagctatttggggcaaattcatagatttattcatcgttccaagtaattgcgcagcattacgcattgactgtgtcatttgctcattcgatcgaactgtttgtatacggagtgaaattgcttggagttgcaccttcatctttgcaaacttctgtatattcttcttcgtcctaataagatcttttgcctgaatcttacacgctctcatttgtccattacgaccagatttctttatttccccaattaattttctctcctgttgctctaactttaatcgctctctatcgagttctctttgagtcttctccagagctcgctggttcttgcgcaatctctcttggggagtttgctttttgccaaacatccattcaaacatcttggctaacctaaatgagacgctggagttagcaaactgcaatgtagaaaagacaacttggaggctcgattcagcgattaggagataaagctaacaaccataagagcagtacaccactaatttagaggttctcttttttcaatagttcgcagctctttctcgctttcacttaaaaagggtgatgttatacgagttagtgggaatcaccagggtgatcagaacttcacctaatgaggatgttaaagcgttcgtatgacttttgatttttgattgttcattactgactatgtagagtgatcaaaacagttggaaagttgataataaataataaaggagtgataagagaaatcaatcccatggggattaagtttctgccaaagattatgacaaaaactcaggagaagcattacaaaggcaatcatttcatgctgttgtttgattcctctcccacagttcagtccgaaattttgaggacattgaggaatgatccacgagtagtccgtgctaatgtctttaaggtaacccactcgaaaggggccttagacattgcatcatgtttttctagaacggactcttaacgaatatttatattctctgtatatatgtaaatgtgtatatcatatgatcaaatcatattattttttcagtcgacttcttcagtttctggtttgctagcattacgttctagtagggtcgggttccacattctcagcatattcgaagctaaagggagtgcgctctgaagattactataattaggtaagtctaccacctctgacctcaagctgattccctttagaatggagctgtccattttggttcttggggtgaggctctggtttagtaaacttatccgtttaagatcatctatagtgtgagacatctcagtacttggaaaatgaacaaccgaggaatcaccagtagagtggagtcgctgattgttaacagtatcggcattaccagaaactacgcccaggagctcaacctgtgccattttggcatcgtcaaaagcgcccatcaaatagggtagctcttccgagatagtggatctttgcaaagagctcataattttctcagatggagcattgaaagaggagttgctcttccctctggtcttgagatgatttttggggtcatgttcaaaattaatttcagaagaaacagaatcagttgggtaaagaaattctttgggtctaacagcatcaaacatcctttcaacatctttaaccacagagagacattccatggttccaatagtgtcattgttgagtatacaatcttccgcctgtttcagaagatcagttatgtagcttctgtatcctaaaatagttttctctgtgaggctttcgttcaaatttagtgccattatgcgttccaaatctatgcctattaccactaacttactcttgcaatacttaatatgcaacttgtattctagactcttttgtagtcggagacatcgaaaagtcaactgtgccaaaattgtacctagatcactaatatagtcagacggtgacattttcaataaaaactcattgattctttctacagctccgtttgatacgttcaattgctcataaagttccatgcaagtcttcaggactttgcttgctagttccttttgaaaaaactgtctttcaactaaatcctgctcaattttatggttcgcagtatcataagatgagttctcatcaccaacggtagacacatagataccatctaccaaatggtattccaaggttttgaactcccaattgtctaatcgctgagcaatgtgctctagatcccgcaaattctggaatcccttcaaacagacatgcttcaatttgcgatcctttgtagctttgatgatatggaaaagcttttgacacttcactttgtctgaattcagcgtcggatctctgagtagatccaccaagagtctccacatgttgagacttccatggcatctgtcaaggaaagagtgtcacgccagatgctatagacaactaggtattcgcgaaggaggcagatcaaagtccctttcctgaaactccttctgtaaaggacgaaataagtagtaataaaagattcgccctattatgtgttctctgctagtatttcagacgaaagcttcttttgccattggttaactgtccttaaaggatgcaatgcgaccaagattaacgtcatgttatcaaacccaactccagttatggtgttcgcttttcttatgcataagtccagcagtttttctacaacaacgttcaaagacagatccaaacttaggaaatgtcgagtttggcgcactaaaggctcgttattatagcaatcccaaactccatcacacgctaggattaaaaaatggtcttccgatgtgatcgtatggataactatgtctggttcagcagtgacttgaaattcttcaggaacggtaagactctccaacgaggatttatttgttgcctccaaagaaacgctcatattattgtagtttctttgctcaaggttagcccttgcttgcattttcttgttgttcaatagtcgggcataattggcttggttgaaacatttgaatgtaaaatctccaaatgctcggcttagtgctaatacacccccaaccctgttggcttgaacgtggccaccgtcgctatgaatacgaaataattcaccgtagtgttttggtttatgatcgaaggacaagttctttacggtgtcatgcttgcctgtgtgtagtatacaccgagagtcgcctatgttggcggtaaacacaattccgttcaaaataatcacaacaactgcggtagaaccagacttaatccttcgagagtatagttcagagtcacatttgaagaaagcattcttaattatgtgggtgaaatttatagtttcaactaaacagtcagtttccgcatatcgagtgccaatagcgttcttgtgctccaccaattgttccttcagaagtccaaatatgatgtgtggtaatctttctcctacatattgtgcagcatttttacccccatgaccatcaaaaacaccaacaatatcaatctcgtatttttgcaagaagccaccgaaggaagtgatctcaagggttttatgccaacagcaatgggcatcctccattgaaacacgatatccttgcatctgaccgactaccatggaaatatgcggccttccatatgtctcaacgaacttttcagttacaggttgtgatataaattggcccataactgaactgctgatgttgctaggctaaatgatgttatggatcgcgcaagaaaccagaagcacttctggaacacgaggtaaaaatgtttacgagtatatactactactgcaagtgcatccaacatctaactcagtcagtcatcaccactattaaaaccaagcaacgaacatcattgtgaatcgccatcacttaaggaacaaaagtcaaaagtgcagtttgattgataattaaagtaaaaatctatagaggtacgagaactcgaaatctctcttacatgttgttacctacccttgtgagttccttgacgaagttttctccccaatatgcagctgtgtatttgctgatatacttgaacatagactcaaaactagattctctcttttcttcgggtagggatagagcttcataaattgcttcacttaactcttctgtattccacggattaaccacgattgcaccattcaatgactgggcagctccagcgaactcactcaatatcagagatcccttcttttgctgctggcatgcaatatactcgtaggatacaagattcattccatcccgtgtacttgaaacgagacagatatccgagacagcgtataaagagatcagctcctcgaagggaacagacttgtgcataaagtgaataggaacaaattctatggtgccaaattggccattgatacgtcccacaagttcattaacagtggcacgaagtgtttgatactcctctacatcgcctcttgagggtacagcaacttgaactaagacaactttgccaatccattctggatgctcctgcaagaatatttcaaaagcatgtagcttttgtggaactcccttgatataatcaagcctgtccaccccgacgattaacttcactccttgaccaaaacgtttttgtaatgagtgaattcgttcctgcaccaagtgctgcttgatccctgtagtgaacttgtccacatctattccgataggatacgctccaacggacacattgtgggaaccatacgtaagacctgttggagttgttctgcagcccagaattcgttccacagacgacgtaaaatgacgagcgtaatcataggtgtggaatccaatcaaatcacagctcaacaccccttccagaatctccttacgtacaggaagaatacgataaatctctgatgaagggaatggcgtgtgcaagaaaaaacctaaacgtacattcttgagtccttgttcatcaatttgacgtcgaagcatttgtgggagtaacattaaatggtagtcatgaatccaaacaagatcgttatcagcgatcttaccaactataactttgcagaactgtctgttggcctcaatgtaagctgcccatgcagcctcatcaaaattcatttctcctggatgataatgaaacaagggccacaaaatagagttgctaaatccattgtagtgtaattctgcaatttcatcactcagatagataggaaagcaattgaacttctctttaaggtcactgttcaccttttcctggtcaatagctggaacttccaaaccaggccagccaaaccattgaaatttttgctttagcccttgaagcgcagtaaccaatccaccacttgacatgttgtattgatagttgccgtcagaaccttttttgattgtaactggaactctattagagatgactagaacatttccttccaccatttcctgaaagtctagaagctttgttgagagagacgaaactaaggaggtaactggtaaaaatatttagacacttgctattttcaaggtaggggaatgtcagttatacttatactacataccaaatgtcgcgcaacaatttccaggagatgaaactgaaactactgtatgcagctaaaaacgagcaacagttcagtttccttacggttgtacgtgccacttcatagcttttgcaataaatctgtctacatatttatcacttatcagtcttgtctgcgctcttaacaatatcttcttcttccccatggcgctctctatcgtagaactcatccagaatctttggtgagattctattcagcatttccttagggtagattctcaacagagaccaagcttgatctaacgactcaaagacagtcctgttatcataggcaccctgagaaataaatgtcttctcaaacttgtctaaaaactctaaggagagtttgtcttcgatcgacaaagcctcttcaccaacaacagctttcattgcagcggcatcacgacctatagcgtatttagcgtataattggttagaaacatcaccatgatccttacgtgtcataccctcgccaatagcgctcttcattaagcgggacaaagatggcaatacgttgatagggggatagatacctcgattgtgcagttgacgatcgactgatatctggccttcggtaatgtaaccagtaagatcgggaattggatgggtgatatcatcgtttggcatggtcaaaatggggatttgagtaatggaaccgttacgaccttccactcgccctgctctctcgtaaatagttgataaatctgtatacatataccccggataacctcgacggccgggaacttcttctctggcagccgaaacttcacgcaacgcatctgcatacgaggacatatctgttaaaattgttagcacatgtctttctgtttgatatgccaaatattcagctgttgtaagagctagacgaggtgtgatgattctttcgatagtgggatcgtttgccaggttcaaaaacaatgatgttctttccagtgacccattctcctcaaagtcttgtttgaagaaccgggaggtttccaagttaacacccatagccgcaaaaacaatagaaaagttctcttcatgaccatcgtgtacatccttagtaggcctgaccaaacctgcctgtctacaaatctgagccgcaatttcgttatgaggcaaccctgaagcggaaaaaataggaatcttttgcccccgagcgatagaattcatagtatcaatagctgatatacccgtggaaatcatttcctctggatagatacgagaatagggattgatgggagacccgtttatatctaaaaagtcttcagcaaaaaccttcggtccattgtctatgggcctaccagacccatcaaaaactcgtcctagcatatcttctgacacagggattttcaaattgtccccagtaaactcgacacgagttttcttgacatcaatgccagaagttccttcaaagacctgaacaatcgctctattgccacggacttctagaacctgtcctgctctgatagttccatcgggtagggtaaggttaacaatttcattaaacctaggaaatttgacattttcaaggatcactaaagggccattaaccccgccaacagtattataattgatacgaggcttaaccttaaatgcttcggccacggccttcttgtttatttcaactcaaacgctagttagtatcattaggagatagaaaagtatagtaacagaagcttacatagttcctggtcggtgagcattgcttcttattgaagtcccttttcaaaaacagtgataacaataaggagttgaattcactgcggtgtatttgttcgtatagattgcgatggctgcttcactggcaagatgacttaggatggaaattacgcagacgtaccatgaattacatagaaggagtaactattgaagttcaactatctcccatttttcatcttgtgccttcaaaatattggaatcagccccaattctgtcagattccttggaaacattgaagtcattagtaggagcttccaaggtactgtttaagtgaaatcttggactattgctgacctcttcacctgaagtatgcttcaatgacgcaacaactttgataccaggagtggcctcagtgataattcgcgatgaatttgcagttttacttacgtctttcaaagaatacttgtctgtcatatcatcaataacttttaattgatgttgaaagaaatcgtagaccttggcattcaatgatggcaagtcaccacattcctttattgaaatcatttttgatttcaaataaagtttattatcgtggtataggttttcggtgaggcattgctcagattcagtgtcctctggcaaaggttcccttggttttggaactaaaccaaagctcctttgtaaggatttctctccattagtttcttcagtcaagcactcgattttgttaccgatgatcctccaaataaaggagtttatatcgaataagtcataaagggttatatctgaattccaagtagactcctgtttgttgagggttaagaagtttaaaacaatgtacgaaacggctttagagagaatgtatagtatctctggatggttcatggtcttctggaatgccagcaaaataccaataggactccctacctatcatggtttgaaattcacactcctggttctgattatcgaagctgtatatctttatgccgaaagggagctttatgcccaaatacctggaagtcaattgctgaaagatcaaaatagattgaaatgacaaattagtcgtatttcgatcatatagcagtatatcttgtatatgtaacacagggacttgccttattctcagctgtgtattggaacttttacttacctgaaggacacttaccaaagtttgtaataagtttcctctggccatgctggtagacttttccaaaaactcaagtttctgatgaatataacatttgctcagctcttggtccatcctacgttttttgtgagcgttctctactgacgacctcaaattaataatatctgcctctctgattcttttcttagagagtagcttctcgattcgttgtttcaactgagaattgcgtatctcctgcttatagcacctactttcaatccttagaatccgtcccaaaattatatggctttccacctgcaataaccctaaatataggttctgcactccatgagctaaatcactttcaatttgattctcaccacgcaaattgtaatgtaagcaattttttagcacggtgttgatacggtataccaattccgatgattcagataccaactctaccaaacctgtttccaatttaagtagtttatagtttatacatcctgagcatagcaagttggggattagtttcgctccatggtgagaagacttgacggtaggtatcagaccgacgagaacagattctgacttagttctcatcccctaaagtaaggatcgtataggagtaatcacatattccgcatttaggcgactctttcataccagcttgtttagaacttaggaacttacatgcattcgtatgatttctaggatatgcgtgtgtggttcacgagcgctgctcccaccttatcgcgaattcgatggttcagtgtgtctattcctttctttgtaatcaaaataatcagactagctaagcagcattcattcagtataataacgggagcaatgtttcatggatacattaaattactgttttacgatacacttcttttgaccaaaggaatgagttggtatgatggttgattagtcagtcacttttgtttttctccagcattagatgttaacgcaatattccacaggtagcccccgttatattagatgaagtatacttttaccgtcatcaaaaagcaactgtggacctgagtaagaactttcaacggttaaaactacaaaatagcatactttactcaattgtcgatgacaggaagccatttttcagtttttcaggcgtcatttgctgacgaaatttttgagctgcagaataaaagatcatttcgtacacccacaaattagaagcctgataagcaatcgatgggcactaagccatttatttgtgctttagtcactgcaagaagcgtaggtacaaaagcactgtgaggattgagcttttcagctttcttttctactgctatccctctatgttttcgtacaccgacagtagcctgctgctaccacaagattaaatccaacaatgtactcgggccagctaattgtgattgtatgccttgatatcagttgtatcatgaaaaatgacgatttctcgatgtagtcatagatctgaaggttgtgccgtgctgggagcgtaggaatcactgggtcgtacaagttatgtaagggcatcaatacgtttttttgttgatgcaaatgaaccacgtatttatctcctcgttcccctacttataataagagagtttggttctggtttttactagtggagtatactgattgcatagctaaactgtttccacttattgaccaaagtcattcgaagattttcaggcgttgtagacagagaaagcgttccgaaaaccatattaaaatagataagtgttcatcccctttaatgttctaccgtgccacctttgaaatggacttagtgtctctcaggctcggagtgcgtcaactatcttgataaattttggagaacgttgatagtgggattcagtcatcacacccgtccaaagaagatttgctcaagatagtccctctagtctcagatactttgcaaggatagggtgagagataggtttgtattcaaccccaactgcctggcacggaatttcaaatgggggaaggggtgaacgtaactaagattctcttcaaacacaattgtcagttggttaatgataaactcatatccgtttgagcccgctaccatgggttgttaacctcgaagtcagctgcagcctaaacacctagatgatgttttgatttttgttgtaaaggcggctcacctcttcggtgatttcgtgtcaaacgagattctcgttgaggcctgacacaaagagatgtaatttttgacgcgtacttttaggaacaatttttctgctatcttagcagcgccatggctattgctactggttcatactcactggtagtaaccaagacttcaagtggcacatgattacaagatgttgagatttcagtgaagttctccttggttatttcgttggcaaaatatgatccccaacaccgagacgagaattgtcttgctgcacaatcgttgaactaagtcgtctgcccctccattttcaccggctgccacctctcttccagctcatcgcaagtccggatgatagagaggtttgctgaacacaactaccagagagcaatagtttaattgatagcatggaccaagacagaggaggatctcgaagtgttttctcgtagctgttattttggatcagaacaactaaagtttcgcaagtgacaacattgtctgtttacaccatcgtaccagttagtggacaaccgtatttatcaacgttttgttgtgagatatttgctccattgttttcattggaaaaatttgcaccatgatattttgtgttttgtatgttctgaaaaatcttctaggttcttatccaattatcacattaacaaccatttggtttctctactgttgcggatggggattttctgatcagaatgtgccagtttatgctagtatcaacagatattcatatcaagcattggggtttttaccgaaacgaatacataaagtttttttacaaagctcagagagtcagcaacccggtcttaagaatcctgacctaactaaaaaatttcccttttactctacttcaagtggatcaaacagttataatctccgtttttgttggaacttgcctatatgcatgcaatcagttttaagacagcagggtgtttggttggctctgtctaacttgatactaaacttgtactgtggtataaaagagtcattgaaattcataaataatcagaaggttgacgattataagttttggccggggtgttggttggtacgattgcttcacactttacaacaggtgaacacgacatcattttactgtaaatacaaaggtgtagaggaaaccgtgggacctaaaaacaaatctgaacagtgtataagaaccaagtgacttactgatcctgcgatggcatgaaaagtttctctgtttgctgagcacgcgtcacttcttcttggcagcaagtaaatgtctgatttgagcttgagcagttgtttgtgaaaacaactgaattcaaaatttgccagctatcacgaatttcagccaaggcagttcctctacaggttaatggtttatgcccctcccacaccagttggagtctggcaagacctagaaaaatataatagacataaaccgttgaaaggtcaaatatagaagctccgtgaagagtattggtaagaacaataaacgttgataagctactacaaccaaaaaaaagacgttttcagcgttagggaatgaatagatttatgaatacagtgaaaaatcagttgcacaacagaacgtatgaaaagtggagatcaggccattaaggcagaactctgtcattaaaaatggagaaactacgaaagttatcacacactggacatggttaatgcatagtaggtagtcttcaattgcgaaagactgaagttgctactggttgaagaaattgccttccagccgaaggattcacttgcaaaataaaaatattattcgatgatgaattaaatagaaaaatacaaattttgaataatttaataaatcacatcttaatctaggtaccctgcagctctaacttaaacaaatgagaatgttaaaaaattagctttcaataaaaaaaatggtagaacttaaactatgaaaaaagtaaataaaattgaacaaatttctaaaatatagtacaataacctcgtgacgggagtaagaagacattaagcattttatctagagattcattgtaactttatatttctttcttctagtgttttccgcatactcccaagataaaccattatcatttctcggctatgaatgttccgttagtttgatgatttgatcaagttgtgcatttttatgtatcactttagattctgcatctagttaagtaatcaaaaaactcgtcttctacttgggtaccaatgtcttgcatattagtacctaatgttcctcttttatggatgtttttcaagcggattagttatggttgaattaatctcttcatctatagacttgattatagaatcgaaacaaaggttttttatggaaacgttgacattttgtcatttcaatcggtaatctctcactctcagaatgaatactagaaaaaaagaaaagaaaactaacaactatagaaatctgcctaagttaatgtaaggatcactcaagtactgaaaacgaggaaacccggtactttctagggatcatcataaacattataaactactgattgacttggccagtagcgacaagaaatgaccaattcttaggataacttgccttgtacatcttcttggctcgcttggagctgcttctgcgtacttttaagaattcttcgacttccgacattgtcttgaagctggcagatctttcgcccatgcggaagagcggaccgcgacagatgatgtcaggatgcgcgacacgagtgtggacaaatttttcgccctcggctttgacctctctgtaagaaccaatccagatgtggcgaattgcggcttgatgcggcttctttcggcgctcgtaagcacctcgggcagtctcatctctatggaactcgcgtatatcgccaaggttatatttagccatgagctctcggcaggaggtaatcagttcgtacagtgatggcctgttgtcttcaccattaggagcctcgagtctaaaggaatagttttgtatacggtcatatttttgtgcttttgctggtggacaaagtgttttgtagtaggcgtcatggtttgtacttttgtcgttcgcataaccgctgaaggggctgtgggcaattagactaggtaggagcaaggcccaaacgggatctagattatcgctttcaatttcggcgtaatactcgctgtcattggcacttgatagccctgtaccgttctggtttttgatgtagtaactaaaaagatcgaagcacattcctaatgtttcttgctgttcctctggaatagagctggtgagcttaaagcgggatgtcttcttgtttgtcattcctatgaaggtacgggcatttgtaatcatgtagttgaactcttgtggaggatggcttttttcgttttgtgtttgcgtatgttcaatgggggtcatcatgatacgtcggatcctgacgttaatgggatgtctttttcgataagcattttgtaggacgacgagcgtttgtggcacaatagcttgcagcggaatagcaggtccttctttttgtggaatgtgagaagtaacctggaaaattgctgctgctagttggtcttcaaaggtctcgtggactgaacttctttcacgttcggtaagtttgttgaagttatgcgtggctgcattaagcgcgcattctttgataagatcagtacagttagaacctaatgcagcgcgcaagacataagtttgttggttgagtgtcgaggactggtggttagattgtggtgtttcgacattgtagtctaaatcggtaatatcggagaagtagcctccgttttcatctacgaagaaaaggctattttcagtattggaatagcataatccacatgtagtaaatgctttttcgccaattagtttttgaaaaaaattgaggtagttgttgagttctgtagatggcgggttttcgccaaatacatcgtcagtggttattatttttttgagtggcttttcgtctgttggacggtttagaaactcccttgctagttctatgaaggtattcaattttttttcttttgtgagttcttctttgattttttgtgcaacatttagaagtaccatggttcgagattgacagctagtatcgccggtaagtgcgtctaatttgcgactgtctgcttctggaacaagaagggtgcataaggccatggccggatgttttttgctggttagaagtatgtatgcgtcatgctgcatgacagataatgtatttgtgattttgtttgtttttttggtcattgtggctactttttgtggaatttgtgtatgggattggaattgggagaggtaggggccttagaagtcgttatatggaagtgttgttttgtagaaagatatgaataaggaagatgaaaaaaattggcggtggtctgggtatttatagtttcttatatgctggatgctggatgctggatgctggatgctggatgctggatgctggatgctggatgctggatgctggatgctggatgctggatgctggatgctggatgctggatgctggatgctggatgctgga

***Contig yDA174:***

tccagcatccagcatccagcatccagcatccagcatccagcatccagcatccagcatccagcatccagcatccagcatccagcatccagcatccagcatccaacatccagcatccagcatccagcatccagcatccagcatccagcatccagcatccagcatccataacagggtaatgcatccagcatccagcatccagcatccagcatccagcatccacgatcggaggaccgaaggagctaaccgcttttttgcacaacatgggggatcatgtaactcgccttgatcgttgggaaccggagctgaatgaagccataccaaacgacgagcgtgacaccacgatgcctgtagcaatggcaacaacgttgcgcaaactattaactggcgaactacttactctagcttcccggcaacaattaatagactggatggaggcggataaagttgcaggaccacttctgcgctcggcccttccggctggctggtttattgctgataaatctggagccggtgagcgtgggtctcgcggtatcattgcagcactggggccagatggtaagccctcccgtatcgtagttatctacacgacggggagtcaggcaactatggatgaacgaaatagacagatcgctgagataggtgcctcactgattaagcattggtaactgtcagaccaagtttactcatatatactttagattgatttaaaacttcatttttaatttaaaaggatctaggtgaagatcctttttgataatctcatgaccaaaatcccttaacgtgagttttcgttccactgagcgtcagaccccgtagaaaagatcaaaggatcttcttgagatccttttctgcgcgtaatctgctgcttgcaaacaaaaaaaccaccgctaccagcggtggtttgtttgccggatcaagagctaccaactctttttccgaaggtaactggcttcagcagagcgcagataccaaatactgttcttctagtgtagccgtagttaggccaccacttcaagaactctgtagcaccgcctacatacctcgctctgctaatcctgttaccagtggctgctgccagtggcgataagtcgtgtcttaccgggttggactcaagacgatagttaccggataaggcgcagcggtcgggctgaacggggggttcgtgcacacagcccagcttggagcgaacgacctacaccgaactgagatacctacagcgtgagctatgagaaagcgccacgcttcccgaagggagaaaggcggacaggtatccggtaagcggcagggtcggaacaggagagcgcacgagggagcttccagggggaaacgcctggtatctttatagtcctgtcgggtttcgccacctctgacttgagcgtcgatttttgtgatgctcgtcaggggggcggagcctatggaaaaacgccagcaacgcggcctttttacggttcctggccttttgctggccttttgctcacatgttctttcctgcgttatcccctgattctgtggataaccgtattaccgcctttgagtgagctgataccgctcgccgcagccgaacgaccgagcgcagcgagtcagtgagcgaggaagcggaagagcgcccaatacgcaaaccgcctctccccgcgcgttggccgattcattaatgcagttgtcattttgtttcttctcgtacgagcttgctcctgatcagcctatctcgcagctgtgttaaaccgtctttaagtcaacccacagtctactgcaatcgtattcagaactagccactagactgttacaagtcgaaacctcagttaaccaactagaaactctagttaccaagttagaactgtgagtacgaaaagtctgaaaagcagaaagattcaatagatttgtctgtgttacacaagagttcaacaagtagctgcgttcgtgctagttgtctaggtttaaacgtttaaaaagacaactagtagtttacttcacctctgtattactgacggtttgtgcctcaacagtttacgttaacaaactagtaagcgtctacttcgtggacttgtacttttagtagaaagattgagtgtctggcagttttaaacctaagtcagttagaatctcaatcctctgcctttggtttagaaacagtgtagctgttttgacctagactgagtacaactgttcttgctgcttattcgaaacttgcctaactcactgcaacagacaagtctagtttagaataaagtaggctcagttattcaagtctaacttcagacagtgtttctcagatttgacttaagtagacagtagtgtgtgaaaaaccagactagtttgaatcttagcctgttcagaaaaaactaactgacttgtgtcaggcctcccaaaaaaactaacttcaatcctctgtactattccacaactattagttgttgcaagtctgaagcctagtcttgctttgaccttacaggatagaggttggctcacaactgagtctgtgtgaaaacaaaagtaaagattgccacaagaagtctattctagttagtccttttacgttggatagcaatagtccaatcttagtttgcagctgatgaatatcttgtggtaggggtttgggaaaatcattcgagtttgatgtttttcttggtatttcccactcctcttcagagtacagaagattaagtgagaccttcgtttgtgcggatcccccacacaccatagcttcaaaatgtttctactccttttttactcttccagattttctcggactccgcgcatcgccgtaccacttcaaaacacccaagcacagcatactaaatttcccctctttcttcctctagggtgtcgttaattacccgtactaaaggtttggaaaagaaaaaagagaccgcctcgtttctttttcttcgtcgaaaaaggcaataaaaatttttatcacgtttctttttcttgaaaattttttttttgatttttttctctttcgatgacctcccattgatatttaagttaataaacggtcttcaatttctcaagtttcagtttcatttttcttgttctattacaactttttttacttcttgctcattagaaagaaagcatagcaatctaatctaagggcggtgttgacaattaatcatcggcatagtatatcggcatagtataatacgacaaggtgaggaactaaaccatggccaagttgaccagtgccgttccggtgctcaccgcgcgcgacgtcgccggagcggtcgagttctggaccgaccggctcgggttctcccgggacttcgtggaggacgacttcgccggtgtggtccgggacgacgtgaccctgttcatcagcgcggtccaggaccaggtggtgccggacaacaccctggcctgggtgtgggtgcgcggcctggacgagctgtacgccgagtggtcggaggtcgtgtccacgaacttccgggacgcctccgggccggccatgaccgagatcggcgagcagccgtgggggcgggagttcgccctgcgcgacccggccggcaactgcgtgcacttcgtggccgaggagcaggactgacacgtccgacgcggcccgacgggtccgaggcctcggagatccgtcccccttttcctttgtcgatatcatgtaattagttatgtcacgcttacattcacgccctcccccacatccgctctaaccgaaaaggaaggagttagacaacctgaagtctaggtccctatttatttttttatagttatgttagtattaagaacgttatttatatttcaaatttttctttttttctgtacagacgcgtgtacgcatgtaacattatactgaaaaccttgcttgagaaggttttgggacgctcgaaggctttaatttgcaagctggagaccaacatgtgagcaaaaggccagcaaaaggccaggaaccgtaaaaaggccgcgaattgtgagcggataacaatttcacacaggaaacagctatgaccatgattacgccaagcttgcatgccgcagaaaggcccacccgaaggtgagccaggtgattacatttgggccctcattagaaaaactcatcgagcatcaagtgaaactgcaatttattcatatcaggattatcaataccatatttttgaaaaagccgtttctgtaatgaaggagaaaactcaccgaggcagttccataggatggcaagatcctggtatcggtctgcgattccgactcgtccaacatcaatacaacctattaatttcccctcgtcaaaaataaggttatcaagtgagaaatcaccatgagtgacgactgaatccggtgagaatggcaaaagcttatgcatttctttccagacttgttcaacaggccagccattacgctcgtcatcaaaatcactcgcaccaaccaaaccgttattcattcgtgattgcgcctgagcgagacgaaatacgcgatcgccgttaaaaggacaattacaaacaggaatcgaatgcaaccggcgcaggaacactgccagcgcatcaacaatattttcacctgaatcaggatattcttctaatacctggaatgctgttttccctgggatcgcagtggtgagtaaccatgcatcatcaggagtacggataaaatgcttgatggtcggaagaggcataaattccgtcagccagtttagcctgaccatctcatctgtaacatcattggcaacgctacctttgccatgtttcagaaacaactctggcgcatcgggcttcccatacaatcgatagattgtcgcacctgattgcccgacattatcgcgagcccatttatacccatataaatcagcatccatgttggaatttaatcgcggcctcgagcaagacgtttcccgttgaatatggctcattttagcttccttagctcctgaaaatctcgataactcaaaaaatacgcccggtagtgatcttatttcattatggtgaaagttggaacctcttacgtgccgatcagcatgcagcaccaccaaaaaaaaaacgaaaagtagaagacccacgatttatgtacccatacgatgttcctgactatgcgggtatgaaaaacatcaaaaaaaaccaggtaatgaacctgggtccgaactctaaactgctgaaagaatacaaatcccagctgatcgaactgaacatcgaacagttcgaagcaggtatcggtctgatcctgggtgatgcttacatccgttctcgtgatgaaggtaaaacctactgtatgcagttcgagtggaaaaacaaagcatacatggaccacgtatgtctgctgtacgatcagtgggtactgtccccgccgcacaaaaaagaacgtgttaaccacctgggtaacctggtaatcacctggggcgcccagactttcaaacaccaagctttcaacaaactggctaacctgttcatcgttaacaacaaaaaaaccatcccgaacaacctggttgaaaactacctgaccccgatgtctctggcatactggttcatggatgatggtggtaaatgggattacaacaaaaactctaccaacaaatcgatcgtactgaacacccagtctttcactttcgaagaagtagaatacctggttaagggtctgcgtaacaaattccaactgaactgttacgtaaaaatcaacaaaaacaaaccgatcatctacatcgattctatgtcttacctgatcttctacaacctgatcaaaccgtacctgatcccgcagatgatgtacaaactgccgaacactatctcctccgaaactttcctgaaataaccgcggcggccgccagcttgggcccgaacaaaaactcatctcagaagaggatctgaatagcgccgtcgaccatcatcatcatcatcattgagttttagccttagacatgactgttcctcagttcaagttgggcacttacgagaagaccggtcttgctagattctaatcaagaggatgtcagaatgccatttgcctgagagatgcaggcttcatttttgatacttttttatttgtaacctatatagtataggattttttttgtcattttgtttcttctcgtacgagcttgctcctgatcagcctatctcgcagctgatgaatatcttgtggtaggggtttgggaaaatcattcgagtttgatgtttttcttggtatttcccactcctcttcagagtacagaagattaagtgagccaaccgtgtaggtctacaaactggtattctcgtttgttgtgaatcttgtcttttctgcttactttctaatagattgagtgagttaggtctgtatttgacggctaaagttaaacttaactgttgtgtgggatagttgttagctgtcaagaaaccgttaaacgtagaaagagtaacttttctaaaccgtagtgtctactcaatattgagacaagcagctagtttaagagcagaagctgaagtctgtgagtttggaaactaaaaggtctgtgaacagctaactagaactaggcttgcttttagtattagaataacaacaaacgaggtctattagttgtgggatctagctagttgcctaagaacaaaggactaagtctgcagaaaaacgagtttcaataggtctgtgtgtaacagtgttaaggctactagcagaactgccacctaagtttcgtaagttccacctcagccttcaatattccagacttgaccaactttcagtttttaaactctaccgtactagtagtttctgcacagtattagacctacttacactgtctgtgcttgcaccacaaaccaaatagtagaatctcttttaagtagaacagtcttttcttgttagactaaaacaacgagagacaactgagtgtagggttacagttaaaaaactgtacagtctgctgtacagacgttacacggcagtcctgttccaaacttcacttaagggttagattggctaagttcaacttgtccttttaagttagttcagacctagttagcagacttaggtgggtaagctgagctacaagagttaaaaacaactgctaagactaaagtaagtgtacgtaagcttcaaactagtaagttaaacttagaaacagctaacgtctaaagctagaagtaagctcaatcttagtactgttaacctaagttgctcacgcaccttcgccaactcaacctgcaggtcgactctagagggattcggcgctccaactgtattctaggcctttccacaatttaaaaaaagacctccgcgacggggaattgaaccccggtctaccgcgcgacaagcggtggttctaccactaaactatcacggatttcgatgctgctttcccacttccttactttacattctctaggcgtttagcctgtagaataaaaccttttttcaagactacaatactgctgctagtaataccgaattattacatgttttaacaacttgagagcgggtcgtatcctgtcttccgattgtacaatctcatcttatcagctcatctcatctcttcggaatcccccgggtaccgagctcgaattcactggccgtcgttttacaacgtcgtgactgggaaaaccctggcgttacccaacttaatcgccttgcagcacatcccgggtaccgtagatcaagtgacttcttgagcttgccaagcatcaagtcccaccgacaagtggtacaccagtcagcgttatctttccatacattctttaattcaaattgtcccattcacaactcaatgtttttgtttctctcgacgaaacgtttttgcttagcatttcggtatcttaaaccgtgtgattttaagtctcataaccttctacatacaggatcaataataatattattgttgatacaatataggattagaatacatagattctgatgttcgaatcaatcaatgtcttaatattcacgtacttgtataccctctgattccacattggtccaacggaatcttctagcaacagccagttgcactagatgtggttctccaccagaaattatagtactgacagattttttggtttacggtgagaggctccaatttccaaagctaaaccaatttcttcataagcaacctccacgcgcgcttttagtatgatgtagaaactggacctaaagatcgaccaatacttgcttacacggcagcgcgaaataaaagacattgataatgtagagtaagtactatttcttcaaaagatagaaccactctcagaggctaactagaagatctaaaacggatgtacagctaatatgtttcgaggcaatcatgttcaggtgcgcctcaagtggctcttaatctattttaaagcataatccacgaaaatttcgatacagagagatacgtcataaactgagaacgatagttagaggcatagcttcggcaatgttcgataaacaagtcaacgagtcagaatttcatgtttctttttttcttgtagtaatatgtaataaccgaatcagataacttgccagtaatgacctgagctctataaattgaaaccgctacaaaaagtaagggatcgtgttaagggcacagagaagaaagtgtaaagaagtaagctagcgttgtacaacaactcaaaagcatggagcccggtgggtactagataataagaacaaggtttcagtgagtttacaagttcataaaacacggaacctggcaacatagcataccagtaaatcgatttgaagaactctattaggctggcagcctctctcaatcaatagtctaaaacggttaagagacacaatcatctctaggtgtcgttgtggtctaccatcgaaacactggaaatcaatccaatgaacttcaagtgatatatttttcaagagtcttcaaagaccaagcaatgttcttattgattgataattttaaagttgaagtctaaatttgtagcagtgaagtctgagcagggggaatgttaagccatatctaatagtacctattagacgcttgattgccgtttgcaaatatgccatgtttctttgaaaccaaaaccacgtaagggggcttaagttttcggcttaggattgttacggagctcaaaccaaataaggcagagagcatacaaaacgtttattaaaaagaacaactacttctggatagctcaattatcttttgtttcttttgaggcgtgcccttcatgacatgacattgcatccatacaattaatagtagataggggaggataagtttgctagtgtgagcatttagagcaaggggtcagtttcctccatctgcctactctgctccatttaaaagcgagccgattgccataagcttgctcacgtagataggacttcaaaaagacgttaagaggctgctctctagaatacgatggaataaacaaatctcgttagtttcttgaaacggaaagtacatgtaggataattgctggatcattcggaagctaaaattggtcctttccccaaagacaatcgactatagttcctgaagcttcctgaggtgagctggatagaaccagaagatagtacctataacgtcaacaacaactgaaactacagatggaagagctaggcatatccataggggatagccatagaaaactttgattgacgcagcaaatcgagacgttaacctgaagctgccataacttcaggaacttgaacacaagagtggcattaaaattcctatgttgcatttcagaaataccacaagtaaactgagcaattaacttgttcataccgacactacaagttaactacaaaccgagacccttatatgcagactgataatacagaatgatacgtatcactcctaagactaggaacatacagagcttttgagcttttggtattttcaagttatttgaaaaattaaatctttatactagggacgaggttcgtgacaaaagaagacaatcatgagactcaccgtcttgtcatctacaagtttcaataagcttccaacttgagtaacctagccatgtgattagtaactttcgaagcatgattcagaacgttctgctctgccgcgtcaaaaagggcagctacttgaagttaaaagaaagatcaaaaattcctgagcattgttcaaacagctataaatcaaaacagaataagaaataaggccatcacttaccaaggaaaaacaaaaaagttcgagagatcaacaatgcacttcagggacgcacctgaacctttacattagtgtaaaaataaattattagattgttcggttcgagcgttagccatatacaaaatgcggccttcaaagtcattggaaaagagctgctttaggacctctccaaaagctaaattgacaaacaggctacctcatatctccatcatctttgccaaccttttcagtaacaaaaaacaaaagcaatatcgtttatttgtccattggacgatcatcaaatattgctaggtaggcacttttttcggtaagcgtgtctgcctcctttcttcagtaagatcaattgtcgccggatactatgaatcgatcattgtagaaactaacatgcttctaattacccattcaatcttgtagcactgagtgcatctcgcgtgagacaaattgaactcgcctacttgggtaggataaacggttgggtaacgtagctgcttcatccggctacttgcttggtctgctgcgtgctttacgcgctgattggtggtcgaataaaccggtacattatttgttcttatccacgacaatgagacgaacaagtaacaaaccagaactggtagctcgatgattacagttccacaatatcagccgaatacccacactttttccaacctgtgtcaattgccttagaaattgtattagcacaaaaatcatttcgctgacggctgaacatccgatatttactattcgtcaaggtatttagaattcttgtgcgaacttgtgggtcgtggtgaaaaagactattgtgagttgttcgtagctgagtccttgactatactgatacacattaaaagaagctgccctttaagcaactaatgtatttgttatgcgtatatctgtagaaagcaacagtgcgacagttgaagtgttcgaacacacttctcaattttgaatctttctcattctcattcagtgaacgatgaaccaagtacacaaatttggaggtgaataattatgctagggagagcttgctagatcatttcacaattagaatgtgaaaccagacctttagatgtaaatgctaaggctaggataggatcgtaattagtaggatcaacaaactttattgattagcctcacctagactaagtttgatcttttcttcatcgtccaatggacaaataaacgatattgcttttgttttttgttactgaaaaggttggcaaagatgatggagatatgaggtagcctgtttgtcaatttagcttttggagaggtcctaaagcagctcttttccaatgactttgaaggccgcattttgtatatggctaacgctcgaaccgaacaatctaataatttatttttacactaatgtaaaggttcaggtgcgtccctgaagtgcattgttgatctctcgaacttttttgtttttccttggtaagtgatggccttatttcttattctgttttgatttatagctgtttgaacaatgctcaggaatttttgatctttcttttaacttcaagtagctgccctttttgacgcggcagagcagaacgttctgaatcatgcttcgaaagttactaatcacatggctaggttactcaagttggaagcttattgaaacttgtagatgacaagacggtgagtctcatgattgtcttcttttgtcacgaacctcgtccctagtataaagatttaatttttcaaataacttgaaaataccaaaagctcaaaagctctgtatgttcctagtcttaggagtgatacgtatcattctgtattatcagtctgcatataagggtctcggtttgtagttaacttgtagtgtcggtatgaacaagttaattgctcagtttacttgtggtatttctgaaatgcaacataggaattttaatgccactcttgtgttcaagttcctgaagttatggcagcttcaggttaacgtctcgatttgctgcgtcaatcaaagttttctatggctatcccctatggatatgcctagctcttccatctgtagtttcagttgttgttgacgttataggtactatcttctggttctatccagctcacctcaggaagcttcaggaactatagtcgattgtctttggggaaaggaccaattttagcttccgaatgatccagcaattatcctacatgtactttccgtttcaagaaactaacgagatttgtttattccatcgtattctagagagcagcctcttaacgtctttttgaagtcctatctacgtgagcaagcttatggcaatcggctcgcttttaaatggagcagagtaggcagatggaggaaactgaccccttgctctaaatgctcacactagcaaacttatcctcccctatctactattaattgtatggatgcaatgtcatgtcatgaagggcacgcctcaaaagaaacaaaagataattgagctatccagaagtagttgttctttttaataaacgttttgtatgctctctgccttatttggtttgagctccgtaacaatcctaagccgaaaacttaagcccccttacgtggttttggtttcaaagaaacatggcatatttgcaaacggcaatcaagcgtctaataggtactattagatatggcttaacattccccctgctcagacttcactgctacaaatttagacttcaactttaaaattatcaatcaataagaacattgcttggtctttgaagactcttgaaaaatatatcacttgaagttcattggattgatttccagtgtttcgatggtagaccacaacgacacctagagatgattgtgtctcttaaccgttttagactattgattgagagaggctgccagcctaatagagttcttcaaatcgatttactggtatgctatgttgccaggttccgtgttttatgaacttgtaaactcactgaaaccttgttcttattatctagtacccaccgggctccatgcttttgagttgttgtacaacgctagcttacttctttacactttcttctctgtgcccttaacacgatcccttactttttgtagcggtttcaatttatagagctcaggtcattactggcaagttatctgattcggttattacatattactacaagaaaaaaagaaacatgaaattctgactcgttgacttgtttatcgaacattgccgaagctatgcctctaactatcgttctcagtttatgacgtatctctctgtatcgaaattttcgtggattatgctttaaaatagattaagagccacttgaggcgcacctgaacatgattgcctcgaaacatattagctgtacatccgttttagatcttctagttagcctctgagagtggttctatcttttgaagaaatagtacttactctacattatcaatgtcttttatttcgcgctgccgtgtaagcaagtattggtcgatctttaggtccagtttctacatcatactaaaagcgcgcgtggaggttgcttatgaagaaattggtttagctttggaaattggagcctctcaccgtaaaccaaaaaatctgtcagtactataatttctggtggagaaccacatctagtgcaactggctgttgctagaagattccgttggaccaatgtggaatcagagggtatacaagtacgtgaatattaagacattgattgattcgaacatcagaatctatgtattctaatcctatattgtatcaacaataatattattattgatcctgtatgtaaaaggttatgagacttaaaatcacacggtttaagataccgaaatgctaagcaaaaacgtttcgtcgagagaaacaaaaacattgagttgcgttataaaatagatagtaacagacttgctaactgagcattcaagactacgttctatttcaggggcttattcaagctgacggagcacatatgcaacaatactacggaaacgacccataacgctgaaacagccgtggaactcgatggaactttcttgccgatttgttgcccaatgtgctccaaaacggagggaagtctttccgcggacagtctggcttgaacgggctagagatggcagagatcggtttgacttagcactcttaaattcatgctttagggttatgtcgcgagacattaatattgggcaagccagggcagatttggctatattcaggctggtttaggcgatccaaaaaggattatgtaagggattttgactgggatggataaaatagttctgattggggcttaatggggctcaagtggctgtgttttctgattgatattggaacctgctgtcatttcgacaattaatatttacttattttggtcaaccccaaataggttgatttcatacttggttcattcaaaaataagtagtcttttgagatctttcaatattataataaatatactataacagccgacttgtttcattttcgcgaatgttcccccagcttatcggatcccgagctcgaattcgctagccagatcggcttaagacaaccaaacccagaactagtcgaatagtctgtgctaaaaaaccaatcggttaaaactagaatcgatctttaagtttaagattcaggcttgttgttaaacctattgccagaaaggacagttgactttggaacaaactcagctaacttgaaactcaaacaacagttttgttacttgccggcttacctcacttgctcagctagtctagcaagaaaacagtgtttcttgtcaacttagaactaatagcagagttaagcagaatcttgaaggctactctgtatttgaaaaccaattgacttgctttttacttagccttacttgctagtcaatcttcagggcagaagtaacttgaaaagttagaaaccaagtttaaaaagacaacaagaaacttgcacacaggactgagaaagtaactaagatttgaaactgagtagactaactctaccagtggttcacaagggttccaccctactttaacttttgacaagagttcaaagtcgaactttaaactgagcttacctctgtggactgtaaactcttgagttaccaaacttttgagctagcctctcagactgttactggttagctctacacagtcttttactattctaacgtaagcttagactagttcttggaataagtgaatctaagaagttgaggctaaactctcaaatagtgtagactaggcacgaactagaagttagtttcttacgttggtagaaaaaaacagttgagtaggctgaaaaggtcaactaagccaaacaagtttcttgtcagaagtctaactgagtgaatacagttgtattcagaaactgtagctagttcgtcttctcaaaccaacttaccagacttttttaaacttgcagtttttactgtacgagacagtttgttactcttcacagtcagttagacgtaaagtaacttgaataagctgaatactaacgtccacgaggtaggcagactatcctgttactgtcttttttagtctgcagtacaaaaagttaaactgttagcaatcctcggttaggcaactttgagttaacttagaagttaccactgcaagctcaatagattcgccctgcagattctaaaaacaaaagctgagctaaagctgctaactgcagttaggtttagatttgagaatagacttggtaaactgcagtaacttacgagaaagtaactaaaactacagttttcaacttcagttttcgaactgtagttgtcttagtctgagttgaaagactgagttaagtaagttaactcactaggttacgctaaagtgtagctactttggattcaggctcactctgcactagcatatggtgcactctcagtacaatctgctctgatgccgcatagttaagccagccccgacacccgccaacacccgctgacgcgccctgacgggcttgtctgctcccggcatccgcttacagacaagctgtgaccgtctccgggagctgcatgtgtcagaggttttcaccgtcatcaccgaaacgcgcgagacgaaagggcctcgtgatacgcctatttttataggttaatgtcatgataataatggtttcttagacgtcaggtggcacttttcggggaaatgtgcgcggaacccctatttgtttatttttctaaatacattcaaatatgtatccgctcatgagacaataaccctgataaatgcttcaataatattgaaaaaggaagagtatgagtattcaacatttccgtgtcgcccttattcccttttttgcggcattttgccttcctgtttttgctcacccagaaacgctggtgaaagtaaaagatgctgaagatcagttgggtgcacgagtgggttacatcgaactggatctcaacagcggtaagatccttgagagttttcgccccgaagaacgttttccaatgatgagcacttttaaagttctgctatgtggcgcggtattatcccgtattgacgccgggcaagagcaactcggtcgccgcatacactattctcagaatgacttggttgagtactcaccagtcacagaaaagcatcttacggatggcatgacagtaagagaattatgcagtgctgccataaccatgagtgataacactgcggccaacttacttctgacaacgatcggaggaccgaaggagctaaccgcttttttgcacaacatgggggatcatgtaactcgccttgatcgttgggaaccggagctgaatgaagccataccaaacgacgagcgtgacaccacgatgcctgtagcaatggcaacaacgttgcgcaaactattaactggcgaactacttactctagcttcccggcaacaattaatagactggatggaggcggataaagttgcaggaccacttctgcgctcggcccttccggctggctggtttattgctgataaatctggagccggtgagcgtgggtctcgcggtatcattgcagcactggggccagatggtaagccctcccgtatcgtagttatctacacgacggggagtcaggcaactatggatgaacgaaatagacagatcgctgagataggtgcctcactgattaagcattggtaactgtcagaccaagtttactcatatatactttagattgatttaaaacttcatttttaatttaaaaggatctaggtgaagatcctttttgataatctcatgaccaaaatcccttaacgtgagttttcgttccactgagcgtcagaccccgtagaaaagatcaaaggatcttcttgagatccttttctgcgcgtaatctgctgcttgcaaacaaaaaaaccaccgctaccagcggtggtttgtttgccggatcaagagctaccaactctttttccgaaggtaactggcttcagcagagcgcagataccaaatactgttcttctagtgtagccgtagttaggccaccacttcaagaactctgtagcaccgcctacatacctcgctctgctaatcctgttaccagtggctgctgccagtggcgataagtcgtgtcttaccgggttggactcaagacgatagttaccggataaggcgcagcggtcgggctgaacggggggttcgtgcacacagcccagcttggagcgaacgacctacaccgaactgagatacctacagcgtgagctatgagaaagcgccacgcttcccgaagggagaaaggcggacaggtatccggtaagcggcagggtcggaacaggagagcgcacgagggagcttccagggggaaacgcctggtatctttatagtcctgtcgggtttcgccacctctgacttgagcgtcgatttttgtgatgctcgtcaggggggcggagcctatggaaaaacgccagcaacgcggcctttttacggttcctggccttttgctggccttttgctcacatgttctttcctgcgttatcccctgattctgtggataaccgtattaccgcctttgagtgagctgataccgctcgccgcagccgaacgaccgagcgcagcgagtcagtgagcgaggaagcggaagagcgcccaatacgcaaaccgcctctccccgcgcgttggccgattcattaatgcagttgtcattttgtttcttctcgtacgagcttgctcctgatcagcctatctcgcagctgtgttaaaccgtctttaagtcaacccacagtctactgcaatcgtattcagaactagccactagactgttacaagtcgaaacctcagttaaccaactagaaactctagttaccaagttagaactgtgagtacgaaaagtctgaaaagcagaaagattcaatagatttgtctgtgttacacaagagttcaacaagtagctgcgttcgtgctagttgtctaggtttaaacgtttaaaaagacaactagtagtttacttcacctctgtattactgacggtttgtgcctcaacagtttacgttaacaaactagtaagcgtctacttcgtggacttgtacttttagtagaaagattgagtgtctggcagttttaaacctaagtcagttagaatctcaatcctctgcctttggtttagaaacagtgtagctgttttgacctagactgagtacaactgttcttgctgcttattcgaaacttgcctaactcactgcaacagacaagtctagtttagaataaagtaggctcagttattcaagtctaacttcagacagtgtttctcagatttgacttaagtagacagtagtgtgtgaaaaaccagactagtttgaatcttagcctgttcagaaaaaactaactgacttgtgtcaggcctcccaaaaaaactaacttcaatcctctgtactattccacaactattagttgttgcaagtctgaagcctagtcttgctttgaccttacaggatagaggttggctcacaactgagtctgtgtgaaaacaaaagtaaagattgccacaagaagtctattctagttagtccttttacgttggatagcaatagtccaatcttagtttgcagctgatgaatatcttgtggtaggggtttgggaaaatcattcgagtttgatgtttttcttggtatttcccactcctcttcagagtacagaagattaagtgagaccttcgtttgtgcggatcccccacacaccatagcttcaaaatgtttctactccttttttactcttccagattttctcggactccgcgcatcgccgtaccacttcaaaacacccaagcacagcatactaaatttcccctctttcttcctctagggtgtcgttaattacccgtactaaaggtttggaaaagaaaaaagagaccgcctcgtttctttttcttcgtcgaaaaaggcaataaaaatttttatcacgtttctttttcttgaaaattttttttttgatttttttctctttcgatgacctcccattgatatttaagttaataaacggtcttcaatttctcaagtttcagtttcatttttcttgttctattacaactttttttacttcttgctcattagaaagaaagcatagcaatctaatctaagggcggtgttgacaattaatcatcggcatagtatatcggcatagtataatacgacaaggtgaggaactaaaccatggccaagttgaccagtgccgttccggtgctcaccgcgcgcgacgtcgccggagcggtcgagttctggaccgaccggctcgggttctcccgggacttcgtggaggacgacttcgccggtgtggtccgggacgacgtgaccctgttcatcagcgcggtccaggaccaggtggtgccggacaacaccctggcctgggtgtgggtgcgcggcctggacgagctgtacgccgagtggtcggaggtcgtgtccacgaacttccgggacgcctccgggccggccatgaccgagatcggcgagcagccgtgggggcgggagttcgccctgcgcgacccggccggcaactgcgtgcacttcgtggccgaggagcaggactgacacgtccgacgcggcccgacgggtccgaggcctcggagatccgtcccccttttcctttgtcgatatcatgtaattagttatgtcacgcttacattcacgccctcccccacatccgctctaaccgaaaaggaaggagttagacaacctgaagtctaggtccctatttatttttttatagttatgttagtattaagaacgttatttatatttcaaatttttctttttttctgtacagacgcgtgtacgcatgtaacattatactgaaaaccttgcttgagaaggttttgggacgctcgaaggctttaatttgcaagctggagaccaacatgtgagcaaaaggccagcaaaaggccaggaaccgtaaaaaggccgcgaattgtgagcggataacaatttcacacaggaaacagctatgaccatgattacgccaagcttgcatgccgcagaaaggcccacccgaaggtgagccaggtgattacatttgggccctcattagaaaaactcatcgagcatcaagtgaaactgcaatttattcatatcaggattatcaataccatatttttgaaaaagccgtttctgtaatgaaggagaaaactcaccgaggcagttccataggatggcaagatcctggtatcggtctgcgattccgactcgtccaacatcaatacaacctattaatttcccctcgtcaaaaataaggttatcaagtgagaaatcaccatgagtgacgactgaatccggtgagaatggcaaaagcttatgcatttctttccagacttgttcaacaggccagccattacgctcgtcatcaaaatcactcgcaccaaccaaaccgttattcattcgtgattgcgcctgagcgagacgaaatacgcgatcgccgttaaaaggacaattacaaacaggaatcgaatgcaaccggcgcaggaacactgccagcgcatcaacaatattttcacctgaatcaggatattcttctaatacctggaatgctgttttccctgggatcgcagtggtgagtaaccatgcatcatcaggagtacggataaaatgcttgatggtcggaagaggcataaattccgtcagccagtttagcctgaccatctcatctgtaacatcattggcaacgctacctttgccatgtttcagaaacaactctggcgcatcgggcttcccatacaatcgatagattgtcgcacctgattgcccgacattatcgcgagcccatttatacccatataaatcagcatccatgttggaatttaatcgcggcctcgagcaagacgtttcccgttgaatatggctcattttagcttccttagctcctgaaaatctcgataactcaaaaaatacgcccggtagtgatcttatttcattatggtgaaagttggaacctcttacgtgccgatcagcatgcagcaccaccaaaaaaaaaacgaaaagtagaagacccacgatttatgtacccatacgatgttcctgactatgcgggtatgaaaaacatcaaaaaaaaccaggtaatgaacctgggtccgaactctaaactgctgaaagaatacaaatcccagctgatcgaactgaacatcgaacagttcgaagcaggtatcggtctgatcctgggtgatgcttacatccgttctcgtgatgaaggtaaaacctactgtatgcagttcgagtggaaaaacaaagcatacatggaccacgtatgtctgctgtacgatcagtgggtactgtccccgccgcacaaaaaagaacgtgttaaccacctgggtaacctggtaatcacctggggcgcccagactttcaaacaccaagctttcaacaaactggctaacctgttcatcgttaacaacaaaaaaaccatcccgaacaacctggttgaaaactacctgaccccgatgtctctggcatactggttcatggatgatggtggtaaatgggattacaacaaaaactctaccaacaaatcgatcgtactgaacacccagtctttcactttcgaagaagtagaatacctggttaagggtctgcgtaacaaattccaactgaactgttacgtaaaaatcaacaaaaacaaaccgatcatctacatcgattctatgtcttacctgatcttctacaacctgatcaaaccgtacctgatcccgcagatgatgtacaaactgccgaacactatctcctccgaaactttcctgaaataaccgcggcggccgccagcttgggcccgaacaaaaactcatctcagaagaggatctgaatagcgccgtcgaccatcatcatcatcatcattgagttttagccttagacatgactgttcctcagttcaagttgggcacttacgagaagaccggtcttgctagattctaatcaagaggatgtcagaatgccatttgcctgagagatgcaggcttcatttttgatacttttttatttgtaacctatatagtataggattttttttgtcattttgtttcttctcgtacgagcttgctcctgatcagcctatctcgcagctgatgaatatcttgtggtaggggtttgggaaaatcattcgagtttgatgtttttcttggtatttcccactcctcttcagagtacagaagattaagtgagccaaccgtgtaggtctacaaactggtattctcgtttgttgtgaatcttgtcttttctgcttactttctaatagattgagtgagttaggtctgtatttgacggctaaagttaaacttaactgttgtgtgggatagttgttagctgtcaagaaaccgttaaacgtagaaagagtaacttttctaaaccgtagtgtctactcaatattgagacaagcagctagtttaagagcagaagctgaagtctgtgagtttggaaactaaaaggtctgtgaacagctaactagaactaggcttgcttttagtattagaataacaacaaacgaggtctattagttgtgggatctagctagttgcctaagaacaaaggactaagtctgcagaaaaacgagtttcaataggtctgtgtgtaacagtgttaaggctactagcagaactgccacctaagtttcgtaagttccacctcagccttcaatattccagacttgaccaactttcagtttttaaactctaccgtactagtagtttctgcacagtattagacctacttacactgtctgtgcttgcaccacaaaccaaatagtagaatctcttttaagtagaacagtcttttcttgttagactaaaacaacgagagacaactgagtgtagggttacagttaaaaaactgtacagtctgctgtacagacgttacacggcagtcctgttccaaacttcacttaagggttagattggctaagttcaacttgtccttttaagttagttcagacctagttagcagacttaggtgggtaagctgagctacaagagttaaaaacaactgctaagactaaagtaagtgtacgtaagcttcaaactagtaagttaaacttagaaacagctaacgtctaaagctagaagtaagctcaatcttagtactgttaacctaagttgctcacgcaccttcgccaactcaacctgcaggtcgactctagagggattcggcgctccaactgtattctaggcctttccacaatttaaaaaaagacctccgcgacggggaattgaaccccggtctaccgcgcgacaagcggtggttctaccactaaactatcacggatttcgatgctgctttcccacttccttactttacattctctaggcgtttagcctgtagaataaaaccttttttcaagactacaatactgctgctagtaataccgaattattacatgttttaacaacttgagagcgggtcgtatcctgtcttccgattgtacaatctcatcttatcagctcatctcatctcttcggaatcccccgggtaccgagctcgaattcactggccgtcgttttacaacgtcgtgactgggaaaaccctggcgttacccaacttaatcgccttgcagcacatcccgggtaccgtagatcaagtgacttcttgagcttgccaagcatcaagtcccaccgacaagtggtacaccagtcagcgttatctttccatacattctttaattcaaattgtcccattcacaactcaatgtttttgtttctctcgacgaaacgtttttgcttagcatttcggtatcttaaaccgtgtgattttaagtctcataaccttctacatacaggatcaataataatattattgttgatacaatataggattagaatacatagattctgatgttcgaatcaatcaatgtcttaatattcacgtacttgtataccctctgattccacattggtccaacggaatcttctagcaacagccagttgcactagatgtggttctccaccagaaattatagtactgacagattttttggtttacggtgagaggctccaatttccaaagctaaaccaatttcttcataagcaacctccacgcgcgcttttagtatgatgtagaaactggacctaaagatcgaccaatacttgcttacacggcagcgcgaaataaaagacattgataatgtagagtaagtactatttcttcaaaagatagaaccactctcagaggctaactagaagatctaaaacggatgtacagctaatatgtttcgaggcaatcatgttcaggtgcgcctcaagtggctcttaatctattttaaagcataatccacgaaaatttcgatacagagagatacgtcataaactgagaacgatagttagaggcatagcttcggcaatgttcgataaacaagtcaacgagtcagaatttcatgtttctttttttcttgtagtaatatgtaataaccgaatcagataacttgccagtaatgacctgagctctataaattgaaaccgctacaaaaagtaagggatcgtgttaagggcacagagaagaaagtgtaaagaagtaagctagcgttgtacaacaactcaaaagcatggagcccggtgggtactagataataagaacaaggtttcagtgagtttacaagttcataaaacacggaacctggcaacatagcataccagtaaatcgatttgaagaactctattaggctggcagcctctctcaatcaatagtctaaaacggttaagagacacaatcatctctaggtgtcgttgtggtctaccatcgaaacactggaaatcaatccaatgaacttcaagtgatatatttttcaagagtcttcaaagaccaagcaatgttcttattgattgataattttaaagttgaagtctaaatttgtagcagtgaagtctgagcagggggaatgttaagccatatctaatagtacctattagacgcttgattgccgtttgcaaatatgccatgtttctttgaaaccaaaaccacgtaagggggcttaagttttcggcttaggattgttacggagctcaaaccaaataaggcagagagcatacaaaacgtttattaaaaagaacaactacttctggatagctcaattatcttttgtttcttttgaggcgtgcccttcatgacatgacattgcatccatacaattaatagtagataggggaggataagtttgctagtgtgagcatttagagcaaggggtcagtttcctccatctgcctactctgctccatttaaaagcgagccgattgccataagcttgctcacgtagataggacttcaaaaagacgttaagaggctgctctctagaatacgatggaataaacaaatctcgttagtttcttgaaacggaaagtacatgtaggataattgctggatcattcggaagctaaaattggtcctttccccaaagacaatcgactatagttcctgaagcttcctgaggtgagctggatagaaccagaagatagtacctataacgtcaacaacaactgaaactacagatggaagagctaggcatatccataggggatagccatagaaaactttgattgacgcagcaaatcgagacgttaacctgaagctgccataacttcaggaacttgaacacaagagtggcattaaaattcctatgttgcatttcagaaataccacaagtaaactgagcaattaacttgttcataccgacactacaagttaactacaaaccgagacccttatatgcagactgataatacagaatgatacgtatcactcctaagactaggaacatacagagcttttgagcttttggtattttcaagttatttgaaaaattaaatctttatactagggacgaggttcgtgacaaaagaagacaatcatgagactcaccgtcttgtcatctacaagtttcaataagcttccaacttgagtaacctagccatgtgattagtaactttcgaagcatgattcagaacgttctgctctgccgcgtcaaaaagggcagctacttgaagttaaaagaaagatcaaaaattcctgagcattgttcaaacagctataaatcaaaacagaataagaaataaggccatcacttaccaaggaaaaacaaaaaagttcgagagatcaacaatgcacttcagggacgcacctgaacctttacattagtgtaaaaataaattattagattgttcggttcgagcgttagccatatacaaaatgcggccttcaaagtcattggaaaagagctgctttaggacctctccaaaagctaaattgacaaacaggctacctcatatctccatcatctttgccaaccttttcagtaacaaaaaacaaaagcaatatcgtttatttgtccattggacgatgaagaaaagatcaaacttagtctaggtgaggctaatcaataaagtttgttgatcctactaattacgatcctatcctagccttagcatttacatctaaaggtctggtttcacattctaattgtgaaatgatctagcaagctctccctagcataattattcacctccaaatttgtgtacttggttcatcgttcactgaatgagaatgagaaagattcaaaattgagaagtgtgttcgaacacttcaactgtcgcactgttgctttctacagatatacgcataacaaatacattagttgcttaaagggcagcttcttttaatgtgtatcagtatagtcaaggactcagctacgaacaactcacaatagtctttttcaccacgacccacaagttcgcacaagaattctaaataccttgacgaatagtaaatatcggatgttcagccgtcagcgaaatgatttttgtgctaatacaatttctaaggcaattgacacaggttggaaaaagtgtgggtattcggctgatattgtggaactgtaatcatcgagctaccagttctggtttgttacttgttcgtctcattgtcgtggataagaacaaataatgtaccggtttattcgaccaccaatcagcgcgtaaagcacgcagcagaccaagcaagtagccggatgaagcagctacgttacccaaccgtttatcctacccaagtaggcgagttcaatttgtctcacgcgagatgcactcagtgctacaagattgaatgggtaattagaagcatgttagtttctacaatgatcgattcatagtatccggcgacaattgatcttactgaagaaaggaggcagacacgcttaccgaaaaaagtgcctacctagcaatatttgatgatcgtccaatggacaaataaacgatattgcttttgttttttgttactgaaaaggttggcaaagatgatggagatatgaggtagcctgtttgtcaatttagcttttggagaggtcctaaagcagctcttttccaatgactttgaaggccgcattttgtatatggctaacgctcgaaccgaacaatctaataatttatttttacactaatgtaaaggttcaggtgcgtccctgaagtgcattgttgatctctcgaacttttttgtttttccttggtaagtgatggccttatttcttattctgttttgatttatagctgtttgaacaatgctcaggaatttttgatctttcttttaacttcaagtagctgccctttttgacgcggcagagcagaacgttctgaatcatgcttcgaaagttactaatcacatggctaggttactcaagttggaagcttattgaaacttgtagatgacaagacggtgagtctcatgattgtcttcttttgtcacgaacctcgtccctagtataaagatttaatttttcaaataacttgaaaataccaaaagctcaaaagctctgtatgttcctagtcttaggagtgatacgtatcattctgtattatcagtctgcatataagggtctcggtttgtagttaacttgtagtgtcggtatgaacaagttaattgctcagtttacttgtggtatttctgaaatgcaacataggaattttaatgccactcttgtgttcaagttcctgaagttatggcagcttcaggttaacgtctcgatttgctgcgtcaatcaaagttttctatggctatcccctatggatatgcctagctcttccatctgtagtttcagttgttgttgacgttataggtactatcttctggttctatccagctcacctcaggaagcttcaggaactatagtcgattgtctttggggaaaggaccaattttagcttccgaatgatccagcaattatcctacatgtactttccgtttcaagaaactaacgagatttgtttattccatcgtattctagagagcagcctcttaacgtctttttgaagtcctatctacgtgagcaagcttatggcaatcggctcgcttttaaatggagcagagtaggcagatggaggaaactgaccccttgctctaaatgctcacactagcaaacttatcctcccctatctactattaattgtatggatgcaatgtcatgtcatgaagggcacgcctcaaaagaaacaaaagataattgagctatccagaagtagttgttctttttaataaacgttttgtatgctctctgccttatttggtttgagctccgtaacaatcctaagccgaaaacttaagcccccttacgtggttttggtttcaaagaaacatggcatatttgcaaacggcaatcaagcgtctaataggtactattagatatggcttaacattccccctgctcagacttcactgctacaaatttagacttcaactttaaaattatcaatcaataagaacattgcttggtctttgaagactcttgaaaaatatatcacttgaagttcattggattgatttccagtgtttcgatggtagaccacaacgacacctagagatgattgtgtctcttaaccgttttagactattgattgagagaggctgccagcctaatagagttcttcaaatcgatttactggtatgctatgttgccaggttccgtgttttatgaacttgtaaactcactgaaaccttgttcttattatctagtacccaccgggctccatgcttttgagttgttgtacaacgctagcttacttctttacactttcttctctgtgcccttaacacgatcccttactttttgtagcggtttcaatttatagagctcaggtcattactggcaagttatctgattcggttattacatattactacaagaaaaaaagaaacatgaaattctgactcgttgacttgtttatcgaacattgccgaagctatgcctctaactatcgttctcagtttatgacgtatctctctgtatcgaaattttcgtggattatgctttaaaatagattaagagccacttgaggcgcacctgaacatgattgcctcgaaacatattagctgtacatccgttttagatcttctagttagcctctgagagtggttctatcttttgaagaaatagtacttactctacattatcaatgtcttttatttcgcgctgccgtgtaagcaagtattggtcgatctttaggtccagtttctacatcatactaaaagcgcgcgtggaggttgcttatgaagaaattggtttagctttggaaattggagcctctcaccgtaaaccaaaaaatctgtcagtactataatttctggtggagaaccacatctagtgcaactggctgttgctagaagattccgttggaccaatgtggaatcagagggtatacaagtacgtgaatattaagacattgattgattcgaacatcagaatctatgtattctaatcctatattgtatcaacaataatattattattgatcctgtatgtagaaggttatgagacttaaaatcacacggtttaagataccgaaatgctaagcaaaaacgtttcgtcgagagaaacaaaaacattgagttgtgaatgggacaatttgaattaaagaatgtatggaaagataacgctgactggtgtaccacttgtcggtgggacttgatgcttggcaagctcaagaagtcacttgatctacggtacccgggatgtgctgcaaggcgattaagttgggtaacgccagggttttcccagtcacgacgttgtaaaacgacggccagtgaattcgagctcggtacccgggggattccgaagagatgagatgagctgataagatgagattgtacaatcggaagacaggatacgacccgctctcaagttgttaaaacatgtaataattcggtattactagcagcagtattgtagtcttgaaaaaaggttttattctacaggctaaacgcctagagaatgtaaagtaaggaagtgggaaagcagcatcgaaatccgtgatagtttagtggtagaaccaccgcttgtcgcgcggtagaccggggttcaattccccgtcgcggaggtctttttttaaattgtggaaaggcctagaatacagttggagcgccgaatccctctagagtcgacctgcaggttgagttggcgaaggtgcgtgagcaacttaggttaacagtactaagattgagcttacttctagctttagacgttagctgtttctaagtttaacttactagtttgaagcttacgtacacttactttagtcttagcagttgtttttaactcttgtagctcagcttacccacctaagtctgctaactaggtctgaactaacttaaaaggacaagttgaacttagccaatctaacccttaagtgaagtttggaacaggactgccgtgtaacgtctgtacagcagactgtacagttttttaactgtaaccctacactcagttgtctctcgttgttttagtctaacaagaaaagactgttctacttaaaagagattctactatttggtttgtggtgcaagcacagacagtgtaagtaggtctaatactgtgcagaaactactagtacggtagagtttaaaaactgaaagttggtcaagtctggaatattgaaggctgaggtggaacttacgaaacttaggtggcagttctgctagtagccttaacactgttacacacagacctattgaaactcgtttttctgcagacttagtcctttgttcttaggcaactagctagatcccacaactaatagacctcgtttgttgttattctaatactaaaagcaagcctagttctagttagctgttcacagaccttttagtttccaaactcacagacttcagcttctgctcttaaactagctgcttgtctcaatattgagtagacactacggtttagaaaagttactctttctacgtttaacggtttcttgacagctaacaactatcccacacaacagttaagtttaactttagccgtcaaatacagacctaactcactcaatctattagaaagtaagcagaaaagacaagattcacaacaaacgagaataccagtttgtagacctacacggttggctcacttaatcttctgtactctgaagaggagtgggaaataccaagaaaaacatcaaactcgaatgattttcccaaacccctaccacaagatattcatcagctgcgagataggctgatcaggagcaagctcgtacgagaagaaacaaaatgacaaaaaaaatcctatactatataggttacaaataaaaaagtatcaaaaatgaagcctgcatctctcaggcaaatggcattctgacatcctcttgattagaatctagcaagaccggtcttctcgtaagtgcccaacttgaactgaggaacagtcatgtctaaggctaaaactcaatgatgatgatgatgatggtcgacggcgctattcagatcctcttctgagatgagtttttgttcgggcccaagctggcggccgccgcggttatttcaggaaagtttcggaggagatagtgttcggcagtttgtacatcatctgcgggatcaggtacggtttgatcaggttgtagaagatcaggtaagacatagaatcgatgtagatgatcggtttgtttttgttgatttttacgtaacagttcagttggaatttgttacgcagacccttaaccaggtattctacttcttcgaaagtgaaagactgggtgttcagtacgatcgatttgttggtagagtttttgttgtaatcccatttaccaccatcatccatgaaccagtatgccagagacatcggggtcaggtagttttcaaccaggttgttcgggatggtttttttgttgttaacgatgaacaggttagccagtttgttgaaagcttggtgtttgaaagtctgggcgccccaggtgattaccaggttacccaggtggttaacacgttcttttttgtgcggcggggacagtacccactgatcgtacagcagacatacgtggtccatgtatgctttgtttttccactcgaactgcatacagtaggttttaccttcatcacgagaacggatgtaagcatcacccaggatcagaccgatacctgcttcgaactgttcgatgttcagttcgatcagctgggatttgtattctttcagcagtttagagttcggacccaggttcattacctggttttttttgatgtttttcatacccgcatagtcaggaacatcgtatgggtacataaatcgtgggtcttctacttttcgtttttttttggtggtgctgcatgctgatcggcacgtaagaggttccaactttcaccataatgaaataagatcactaccgggcgtattttttgagttatcgagattttcaggagctaaggaagctaaaatgagccatattcaacgggaaacgtcttgctcgaggccgcgattaaattccaacatggatgctgatttatatgggtataaatgggctcgcgataatgtcgggcaatcaggtgcgacaatctatcgattgtatgggaagcccgatgcgccagagttgtttctgaaacatggcaaaggtagcgttgccaatgatgttacagatgagatggtcaggctaaactggctgacggaatttatgcctcttccgaccatcaagcattttatccgtactcctgatgatgcatggttactcaccactgcgatcccagggaaaacagcattccaggtattagaagaatatcctgattcaggtgaaaatattgttgatgcgctggcagtgttcctgcgccggttgcattcgattcctgtttgtaattgtccttttaacggcgatcgcgtatttcgtctcgctcaggcgcaatcacgaatgaataacggtttggttggtgcgagtgattttgatgacgagcgtaatggctggcctgttgaacaagtctggaaagaaatgcataagcttttgccattctcaccggattcagtcgtcactcatggtgatttctcacttgataaccttatttttgacgaggggaaattaataggttgtattgatgttggacgagtcggaatcgcagaccgataccaggatcttgccatcctatggaactgcctcggtgagttttctccttcattacagaaacggctttttcaaaaatatggtattgataatcctgatatgaataaattgcagtttcacttgatgctcgatgagtttttctaatgagggcccaaatgtaatcacctggctcaccttcgggtgggcctttctgcggcatgcaagcttggcgtaatcatggtcatagctgtttcctgtgtgaaattgttatccgctcacaattcgcggcctttttacggttcctggccttttgctggccttttgctcacatgttggtctccagcttgcaaattaaagccttcgagcgtcccaaaaccttctcaagcaaggttttcagtataatgttacatgcgtacacgcgtctgtacagaaaaaaagaaaaatttgaaatataaataacgttcttaatactaacataactataaaaaaataaatagggacctagacttcaggttgtctaactccttccttttcggttagagcggatgtggggagggcgtgaatgtaagcgtgacataactaattacatgatatcgacaaaggaaaagggggacggatctccgaggcctcggacccgtcgggccgcgtcggacgtgtcagtcctgctcctcggccacgaagtgcacgcagttgccggccgggtcgcgcagggcgaactcccgcccccacggctgctcgccgatctcggtcatggccggcccggaggcgtcccggaagttcgtggacacgacctccgaccactcggcgtacagctcgtccaggccgcgcacccacacccaggccagggtgttgtccggcaccacctggtcctggaccgcgctgatgaacagggtcacgtcgtcccggaccacaccggcgaagtcgtcctccacgaagtcccgggagaacccgagccggtcggtccagaactcgaccgctccggcgacgtcgcgcgcggtgagcaccggaacggcactggtcaacttggccatggtttagttcctcaccttgtcgtattatactatgccgatatactatgccgatgattaattgtcaacaccgcccttagattagattgctatgctttctttctaatgagcaagaagtaaaaaaagttgtaatagaacaagaaaaatgaaactgaaacttgagaaattgaagaccgtttattaacttaaatatcaatgggaggtcatcgaaagagaaaaaaatcaaaaaaaaaattttcaagaaaaagaaacgtgataaaaatttttattgcctttttcgacgaagaaaaagaaacgaggcggtctcttttttcttttccaaacctttagtacgggtaattaacgacaccctagaggaagaaagaggggaaatttagtatgctgtgcttgggtgttttgaagtggtacggcgatgcgcggagtccgagaaaatctggaagagtaaaaaaggagtagaaacattttgaagctatggtgtgtgggggatccgcacaaacgaaggtctcacttaatcttctgtactctgaagaggagtgggaaataccaagaaaaacatcaaactcgaatgattttcccaaacccctaccacaagatattcatcagctgcaaactaagattggactattgctatccaacgtaaaaggactaactagaatagacttcttgtggcaatctttacttttgttttcacacagactcagttgtgagccaacctctatcctgtaaggtcaaagcaagactaggcttcagacttgcaacaactaatagttgtggaatagtacagaggattgaagttagtttttttgggaggcctgacacaagtcagttagttttttctgaacaggctaagattcaaactagtctggtttttcacacactactgtctacttaagtcaaatctgagaaacactgtctgaagttagacttgaataactgagcctactttattctaaactagacttgtctgttgcagtgagttaggcaagtttcgaataagcagcaagaacagttgtactcagtctaggtcaaaacagctacactgtttctaaaccaaaggcagaggattgagattctaactgacttaggtttaaaactgccagacactcaatctttctactaaaagtacaagtccacgaagtagacgcttactagtttgttaacgtaaactgttgaggcacaaaccgtcagtaatacagaggtgaagtaaactactagttgtctttttaaacgtttaaacctagacaactagcacgaacgcagctacttgttgaactcttgtgtaacacagacaaatctattgaatctttctgcttttcagacttttcgtactcacagttctaacttggtaactagagtttctagttggttaactgaggtttcgacttgtaacagtctagtggctagttctgaatacgattgcagtagactgtgggttgacttaaagacggtttaacacagctgcgagataggctgatcaggagcaagctcgtacgagaagaaacaaaatgacaactgcattaatgaatcggccaacgcgcggggagaggcggtttgcgtattgggcgctcttccgcttcctcgctcactgactcgctgcgctcggtcgttcggctgcggcgagcggtatcagctcactcaaaggcggtaatacggttatccacagaatcaggggataacgcaggaaagaacatgtgagcaaaaggccagcaaaaggccaggaaccgtaaaaaggccgcgttgctggcgtttttccataggctccgccccctgacgagcatcacaaaaatcgacgctcaagtcagaggtggcgaaacccgacaggactataaagataccaggcgtttccccctggaagctccctcgtgcgctctcctgttccgaccctgccgcttaccggatacctgtccgcctttctcccttcgggaagcgtggcgctttctcatagctcacgctgtaggtatctcagttcggtgtaggtcgttcgctccaagctgggctgtgtgcacgaaccccccgttcagcccgaccgctgcgccttatccggtaactatcgtcttgagtccaacccggtaagacacgacttatcgccactggcagcagccactggtaacaggattagcagagcgaggtatgtaggcggtgctacagagttcttgaagtggtggcctaactacggctacactagaagaacagtatttggtatctgcgctctgctgaagccagttaccttcggaaaaagagttggtagctcttgatccggcaaacaaaccaccgctggtagcggtggtttttttgtttgcaagcagcagattacgcgcagaaaaggatctcaagaagatcctttgatcttttctacggggtctgacgctcagtggaacgaaaactcacgttaagggattttggtcatgagattatcaaaaaggatcttcacctagatccttttaaattaaaaatgaagttttaaatcaatctaaagtatatatgagtaaacttggtctgacagttaccaatgcttaatcagtgaggcacctatctcagcgatctgtctatttcgttcatccatagttgcctgactccccgtcgtgtagataactacgatacgggagggcttaccatctggccccagtgctgcaatgataccgcgagacccacgctcaccggctccagatttatcagcaataaaccagccagccggaagggccgagcgcagaagtggtcctgcaactttatccgcctccatccagtctattaattgttgccgggaagctagagtaagtagttcgccagttaatagtttgcgcaacgttgttgccattgctacaggcatcgtggtgtcacgctcgtcgtttggtatggcttcattcagctccggttcccaacgatcaaggcgagttacatgatcccccatgttgtgcaaaaaagcggttagctccttcggtcctccgatcgtggatgctggatgctggatgctggatgctggatgctggatgcattaccctgttatggatgctggatgctggatgctggatgctggatgctggatgctggatgctggatgctggatgttggatgctggatgctggatgctggatgctggatgctggatgctggatgctggatgctggatgctggatgctgga

***Contig yDA175:***

cagcatccagcatccagcatccagcatccagcatccagcatccagcatccagcatccagcatccagcatccagcatccagcatccagcatccagcatccagcatccagcatccagcatccagcatccagcatccagcatccagcatccacgatcggaggaccgaaggagctaaccgcttttttgcacaacatgggggatcatgtaactcgccttgatcgttgggaaccggagctgaatgaagccataccaaacgacgagcgtgacaccacgatgcctgtagcaatggcaacaacgttgcgcaaactattaactggcgaactacttactctagcttcccggcaacaattaatagactggatggaggcggataaagttgcaggaccacttctgcgctcggcccttccggctggctggtttattgctgataaatctggagccggtgagcgtgggtctcgcggtatcattgcagcactggggccagatggtaagccctcccgtatcgtagttatctacacgacggggagtcaggcaactatggatgaacgaaatagacagatcgctgagataggtgcctcactgattaagcattggtaactgtcagaccaagtttactcatatatactttagattgatttaaaacttcatttttaatttaaaaggatctaggtgaagatcctttttgataatctcatgaccaaaatcccttaacgtgagttttcgttccactgagcgtcagaccccgtagaaaagatcaaaggatcttcttgagatccttttctgcgcgtaatctgctgcttgcaaacaaaaaaaccaccgctaccagcggtggtttgtttgccggataagagctaccaactctttttccgaaggtaactggcttcagcagagcgcagataccaaatactgttcttctagtgtagccgtagttaggccaccacttcaagaactctgtagcaccgcctacatacctcgctctgctaatcctgttaccagtggctgctgccagtggcgataagtcgtgtcttaccgggttggactcaagacgatagttaccggataaggcgcagcggtcgggctgaacggggggttcgtgcacacagcccagcttggagcgaacgacctacaccgaactgagatacctacagcgtgagctatgagaaagcgccacgcttcccgaagggagaaaggcggacaggtatccggtaagcggcagggtcggaacaggagagcgcacgagggagcttccagggggaaacgcctggtatctttatagtcctgtcgggtttcgccacctctgacttgagcgtcgatttttgtgatgctcgtcaggggggcggagcctatggaaaaacgccagcaacgcggcctttttacggttcctggccttttgctggccttttgctcacatgttctttcctgcgttatcccctgattctgtggataaccgtattaccgcctttgagtgagctgataccgctcgccgcagccgaacgaccgagcgcagcgagtcagtgagcgaggaagcggaagagcgcccaatacgcaaaccgcctctccccgcgcgttggccgattcattaatgcagttgtcattttgtttcttctcgtacgagcttgctcctgatcagcctatctcgcagctgtgttaaaccgtctttaagtcaacccacagtctactgcaatcgtattcagaactagccactagactgttacaagtcgaaacctcagttaaccaactagaaactctagttaccaagttagaactgtgagtacgaaaagtctgaaaagcagaaagattcaatagatttgtctgtgttacacaagagttcaacaagtagctgcgttcgtgctagttgtctaggtttaaacgtttaaaaagacaactagtagtttacttcacctctgtattactgacggtttgtgcctcaacagtttacgttaacaaactagtaagcgtctacttcgtggacttgtacttttagtagaaagattgagtgtctggcagttttaaacctaagtcagttagaatctcaatcctctgcctttggtttagaaacagtgtagctgttttgacctagactgagtacaactgttcttgctgcttattcgaaacttgcctaactcactgcaacagacaagtctagtttagaataaagtaggctcagttattcaagtctaacttcagacagtgtttctcagatttgacttaagtagacagtagtgtgtgaaaaaccagactagtttgaatcttagcctgttcagaaaaaactaactgacttgtgtcaggcctcccaaaaaaactaacttcaatcctctgtactattccacaactattagttgttgcaagtctgaagcctagtcttgctttgaccttacaggatagaggttggctcacaactgagtctgtgtgaaaacaaaagtaaagattgccacaagaagtctattctagttagtccttttacgttggatagcaatagtccaatcttagtttgcagctgatgaatatcttgtggtaggggtttgggaaaatcattcgagtttgatgtttttcttggtatttcccactcctcttcagagtacagaagattaagtgagaccttcgtttgtgcggatcccccacacaccatagcttcaaaatgtttctactccttttttactcttccagattttctcggactccgcgcatcgccgtaccacttcaaaacacccaagcacagcatactaaatttcccctctttcttcctctagggtgtcgttaattacccgtactaaaggtttggaaaagaaaaaagagaccgcctcgtttctttttcttcgtcgaaaaaggcaataaaaatttttatcacgtttctttttcttgaaaattttttttttgatttttttctctttcgatgacctcccattgatatttaagttaataaacggtcttcaatttctcaagtttcagtttcatttttcttgttctattacaactttttttacttcttgctcattagaaagaaagcatagcaatctaatctaagggcggtgttgacaattaatcatcggcatagtatatcggcatagtataatacgacaaggtgaggaactaaaccatggccaagttgaccagtgccgttccggtgctcaccgcgcgcgacgtcgccggagcggtcgagttctggaccgaccggctcgggttctcccgggacttcgtggaggacgacttcgccggtgtggtccgggacgacgtgaccctgttcatcagcgcggtccaggaccaggtggtgccggacaacaccctggcctgggtgtgggtgcgcggcctggacgagctgtacgccgagtggtcggaggtcgtgtccacgaacttccgggacgcctccgggccggccatgaccgagatcggcgagcagccgtgggggcgggagttcgccctgcgcgacccggccggcaactgcgtgcacttcgtggccgaggagcaggactgacacgtccgacgcggcccgacgggtccgaggcctcggagatccgtcccccttttcctttgtcgatatcatgtaattagttatgtcacgcttacattcacgccctcccccacatccgctctaaccgaaaaggaaggagttagacaacctgaagtctaggtccctatttatttttttatagttatgttagtattaagaacgttatttatatttcaaatttttcttttttttctgtacagacgcgtgtacgcatgtaacattatactgaaaaccttgcttgagaaggttttgggacgctcgaaggctttaatttgcaagctggagaccaacatgtgagcaaaaggccagcaaaaggccaggaaccgtaaaaaggccgcgaattgtgagcggataacaatttcacacaggaaacagctatgaccatgattacgccaagcttgcatgccgcagaaaggcccacccgaaggtgagccaggtgattacatttgggccctcattagaaaaactcatcgagcatcaagtgaaactgcaatttattcatatcaggattatcaataccatatttttgaaaaagccgtttctgtaatgaaggagaaaactcaccgaggcagttccataggatggcaagatcctggtatcggtctgcgattccgactcgtccaacatcaatacaacctattaatttcccctcgtcaaaaataaggttatcaagtgagaaatcaccatgagtgacgactgaatccggtgagaatggcaaaagcttatgcatttctttccagacttgttcaacaggccagccattacgctcgtcatcaaaatcactcgcaccaaccaaaccgttattcattcgtgattgcgcctgagcgagacgaaatacgcgatcgccgttaaaaggacaattacaaacaggaatcgaatgcaaccggcgcaggaacactgccagcgcatcaacaatattttcacctgaatcaggatattcttctaatacctggaatgctgttttccctgggatcgcagtggtgagtaaccatgcatcatcaggagtacggataaaatgcttgatggtcggaagaggcataaattccgtcagccagtttagcctgaccatctcatctgtaacatcattggcaacgctacctttgccatgtttcagaaacaactctggcgcatcgggcttcccatacaatcgatagattgtcgcacctgattgcccgacattatcgcgagcccatttatacccatataaatcagcatccatgttggaatttaatcgcggcctcgagcaagacgtttcccgttgaatatggctcattttagcttccttagctcctgaaaatctcgataactcaaaaaatacgcccggtagtgatcttatttcattatggtgaaagttggaacctcttacgtgccgatcagcatgcagcaccaccaaaaaaaaacgaaaagtagaagacccacgatttatgtacccatacgatgttcctgactatgcgggtatgaaaaacatcaaaaaaaaccaggtaatgaacctgggtccgaactctaaactgctgaaagaatacaaatcccagctgatcgaactgaacatcgaacagttcgaagcaggtatcggtctgatcctgggtgatgcttacatccgttctcgtgatgaaggtaaaacctactgtatgcagttcgagtggaaaaacaaagcatacatggaccacgtatgtctgctgtacgatcagtgggtactgtccccgccgcacaaaaaagaacgtgttaaccacctgggtaacctggtaatcacctggggcgcccagactttcaaacaccaagctttcaacaaactggctaacctgttcatcgttaacaacaaaaaaaccatcccgaacaacctggttgaaaactacctgaccccgatgtctctggcatactggttcatggatgatggtggtaaatgggattacaacaaaaactctaccaacaaatcgatcgtactgaacacccagtctttcactttcgaagaagtagaatacctggttaagggtctgcgtaacaaattccaactgaactgttacgtaaaaatcaacaaaaacaaaccgatcatctacatcgattctatgtcttacctgatcttctacaacctgatcaaaccgtacctgatcccgcagatgatgtacaaactgccgaacactatctcctccgaaactttcctgaaataaccgcggcggccgccagcttgggcccgaacaaaaactcatctcagaagaggatctgaatagcgccgtcgaccatcatcatcatcatcattgagttttagccttagacatgactgttcctcagttcaagttgggcacttacgagaagaccggtcttgctagattctaatcaagaggatgtcagaatgccatttgcctgagagatgcaggcttcatttttgatacttttttatttgtaacctatatagtataggattttttttgtcattttgtttcttctcgtacgagcttgctcctgatcagcctatctcgcagctgatgaatatcttgtggtaggggtttgggaaaatcattcgagtttgatgtttttcttggtatttcccactcctcttcagagtacagaagattaagtgagccaaccgtgtaggtctacaaactggtattctcgtttgttgtgaatcttgtcttttctgcttactttctaatagattgagtgagttaggtctgtatttgacggctaaagttaaacttaactgttgtgtgggatagttgttagctgtcaagaaaccgttaaacgtagaaagagtaacttttctaaaccgtagtgtctactcaatattgagacaagcagctagtttaagagcagaagctgaagtctgtgagtttggaaactaaaaggtctgtgaacagctaactagaactaggcttgcttttagtattagaataacaacaaacgaggtctattagttgtgggatctagctagttgcctaagaacaaaggactaagtctgcagaaaaacgagtttcaataggtctgtgtgtaacagtgttaaggctactagcagaactgccacctaagtttcgtaagttccacctcagccttcaatattccagacttgaccaactttcagtttttaaactctaccgtactagtagtttctgcacagtattagacctacttacactgtctgtgcttgcaccacaaaccaaatagtagaatctcttttaagtagaacagtcttttcttgttagactaaaacaacgagagacaactgagtgtagggttacagttaaaaaactgtacagtctgctgtacagacgttacacggcagtcctgttccaaacttcacttaagggttagattggctaagttcaacttgtccttttaagttagttcagacctagttagcagacttaggtgggtaagctgagctacaagagttaaaaacaactgctaagactaaagtaagtgtacgtaagcttcaaactagtaagttaaacttagaaacagctaacgtctaaagctagaagtaagctcaatcttagtactgttaacctaagttgctcacgcaccttcgccaactcaacctgcaggtcgactctagaggggattcggcgctccaactgtattctaggcctttccacaatttaaaaaaagacctccgcgacggggaattgaaccccggtctaccgcgcgacaagcggtggttctaccactaaactatcacggatttcgatgctgctttccacttccttactttacattctctaggcgtttagcctgtagaataaaaccttttttcaagactacaatactgctgctagtaataccgaattattacatgttttaacaacttgagagcgggtcgtatcctgtcttccgattgtacaatctcatcttatcagctcatctcatctcttcggaatcccccgggtaccgagctcgaattcactggccgtcgttttacaacgtcgtgactgggaaaaccctggcgttacccaacttaatcgccttgcagcacatcccggtaccgtagatcaagtgacttcttgagcttgccaagcatcaagtcccaccgacaagtggtacaccagtcagcgttatctttccatacattctttaattcaaattgtcccattcacaactcaatgtttttgtttctctcgacgaaacgtttttgcttagcatttcggtatcttaaaccgtgtgattttaagtctcataaccttctacatacaggatcaataataatattattgttgatacaatataggattagaatacatagattctgatgttcgaatcaatcaatgtcttaatattcacgtacttgtataccctctgattccacattggtccaacggaatcttctagcaacagccagttgcactagatgtggttctccacgaaattatagtactgacagattttttggtttacggtgagaggctccaattttccaaagctaaaccaatttcttcataagcaacctccacgcgcgcttttagtatgatgtagaaactggacctaaagatcgaccaatacttgcttacacggcagcgcgaaataaaagacattgataatgtagagtaagtactatttcttcaaaagatagaaccactctcagaggctaactagaagatctaaaacggatgtacagctaatatgtttcgaggcaatcatgttcaggtgcgcctcaagtggctcttaatctattttaaagcataatccacgaaaatttcgatacagagagatacgtcataaactgagaacgatagttagaggcatagcttcggcaatgttagataaacaagtcaacgagtcagaatttcatgtttctttttttcttgtagtaatatgtaataaccgaatcagataacttgccagtaatgacctgagctctataaattgaaaccgctacaaaaagtaagggatcgtgttaagggcacagagaagaaagtgtaaagaagtaagctagcgttgtacaacaactcaaaagcatggagcccggtgggtactagataataagaacaaggtttcagtgagtttacaagttcataaaacacggaacctggcaacatagcatacagtaaatcgatttgaagaactctattaggctggcagcctctctcaatcaatagtctaaaacggttaagagacacaatcatctctaggtgtcgttgtggtctaccatcgaaacactggaaatcaatccaatgaacttcaagtgatatattttcaagagtcttcaaagaccaagcaatgttcttattgattgataattttaaagttgaagtctaaatttgtagcagtgaagtctgagcaggggaatgttaagccatatgtaatagtacctattagacgcttgattgccgtttgcaaatatgccatgtttctttgaaaccaaaaccacgtaagggggcttaagttttcggcttaggagttattcaggagctcaaaccaaataaggcagagagcatacaaaacgtttattaaaaagaacaactacttctggatagctcaattatcttttgtttcttttggcgtgcccttcatgacatgacattgcatccatacaattaatagtagatcaggaaggataagtttgctagtgtgagcatttagagcaaggggtcagtttcctccatctgcctactctgctccatttaaaagcgagccgattgccataagcttgctcacgtagataggacttcaaaaagacgttaagaggctgctctctagaatacgatggaataaacaaatctcgttagtttcttgaaacggaaagtacatgtaggataattgctggatcattcggaagctaaaattggtcctttccccaaagacaatcgactatagttcctgaagcttcctgaggtgagctggatagaaccagaagatagtacctataacgtcaacaacaactgaaactacagatggaagagctaggcatatccataggggatagccatagaaaactttgattgacgcagcaaatcgagacgttaacctgaagctgccataacttcaggaacttgaacacaagagtggcattaaaattcctatgttgcatttcagaaataccacaagtaaactgagcaattaacttgttcataccgacactacaagttaactacaaaccgagacccttatatgcagactgataatacagaatgatacgtatcactcctaagactaggaacatacagagcttttgagcttttggtattttcaagttatttgaaaaattaaatctttatactagggacgaggttcgtgacaaaagaagacaatcatgagactcaccgtcttgtcatctacaagtttcaataagcttccaacttgagtaacctagccatgtgattagtaactttcgaagcatgattcagaacgttctgctctgccgcgtcaaaaagggcagctacttgaagttaaaagaaagatcaaaaattcctgagcattgttcaaacagctataaatcaaaacagaataagaaataaggccatcacttaccaaggaaaaacaaaaaagttcgaagatcaacaatgcacttcagggacgcacctgaacctttacattagtgtaaaaataaattattagattgggttcgagcattagccatatacaaatgcggccttcaaagtcattggaaaagagctgctttaggacctctccaaaagctaaattgacaaacaggctacctcatatctccatcatctttgccaaccttttcagtaacaaaaaacaaaagcaatatcgtttatttgtccattggacgatcatcaaatattgctaggtaggcacttttttcggtaagcgtgtctgcctcctttcttcagtaagatcaattgtcgccggatactatgaatcgatcattgtagaaactaacatgcttctaattacccattcaatcttgtagcactgagtgcatctcgcgtgagacaaattgaactcgcctattgggtagtgaaacggttgggtaacgtgctgcttcatccggctacttgcttggtctgctgcgtgctttacgcgctgattggtggtcgaataaaccggtacattattttgttcttatccacgacaatgagacgaacaagtaacaaaccagaactggtagctcgatgattacagttccacaatatcagccgaatacccacactttttccaacctgtgtcaattgccttagaaattgtattagcacaaaaatcatttcgctgacggctgaacatccgatatttactattcgtcaaggtatttagaattcttgtgcgaacttgtgggtcgtggtgaaaaagactattgtgagttgttcgtagctgagtccttgactatactgatacacattaaaagaagctgccctttaagcaactaatgtatttgttatgcgtatatctgtagaaagcaacagtgcgacagttgaagtgttcgaacacacttctcaattttgaatctttctcattctcattcagtgaacgatgaaccaagtacacaaatttggaggtgaataattatgctagggagagcttgctagatcatttcacaattagaatgtgaaaccagacctttagatgtaaatgctaaggctaggataggatcgtaattagtaggatcaacaaactttattgattagcctcacctagactaagtttgatcttttcttcatcgtccaatggacaaataaacgatattgcttttgttttttgttactgaaaaggttggcaaagatgatggagatatgaggtagcctgtttgtcaatttagcttttggagaggtcctaaagcagctctttccaatgactttgaaggccgcattttgtatatggctaacgctcgaaccgaacaatctaataatttatttttacactaatgtaaaggttcaggtgcgtccctgaagtgcattgttgatctctcgaacttttttgtttttccttggtaagtgatggccttatttcttattctgttttgatttatagctgtttgaacaatgctcaggaatttttgatctttcttttaacttcaagtagctgccctttttgacgcggcagagcagaacgttctgaatcatgcttcgaaagttactaatcacatggctaggttactcaagttggaagcttattgaaacttgtagatgacaagacggtgagtctcatgattgtcttcttttgtcacgaacctcgtccctagtataaagatttaatttttcaaataacttgaaaataccaaaagctcaaaagctctgtatgttcctagtcttaggagtgatacgtatcattctgtattatcagtctgcatataagggtctcggtttgtagttaacttgtagtgtcggtatgaacaagttaattgctcagtttacttgtggtatttctgaaatgcaacataggaattttaatgccactcttgtgttcaagttcctgaagttatggcagcttcaggttaacgtctcgatttgctgcgtcaatcaaagttttctatggctatcccctatggatatgcctagctcttccatctgtagtttcagttgttgttgacgttataggtactatcttctggttctatccagctcacctcaggaagcttcaggaactatagtcgattgtctttgggaaaggaccaattttagcttccgaatgatccagcaattatcctacatgtactttccgtttcaagaaactaacgagatttgtttattccatcgtattctagagagcagcctcttaacgtctttttgaagtcctatctacgtgagcaagcttatggcaatcggctcgcttttaaatggagcagagtaggcagatggaggaaactgaccccttgctctaaatgctcacactagcaaacttatcctcccctatctactattaattgtatggatgcaatgtcatgtcatgaagggcacgcctcaaaagaaacaaaagataattgagctatccagaagtagttgttctttttaataaacgttttgtatgctctctgccttatttggtttgagctccgtaacaatcctaagccgaaaacttaagcccccttacgtggttttggtttcaaagaaacatggcatatttgcaaacggcaatcaagcgtctaataggtactattagatatggcttaacattccccctgctcagacttcactgctacaaatttagacttcaactttaaaattatcaatcaataagaacattgcttggtctttgaagactcttgaaaaatatatcacttgaagttcattggattgatttccagtgtttcgatggtagaccacaacgacacctagagatgattgtgtctcttaaccgttttaactatttgattgagagaggctgccagcctaatagagttcttcaaatcgatttactggtatgctatgttgccaggttccgtgttttatgaacttgtaaactcactgaaaccttgttcttattatctagtccaccacggctccatgcttttgagttgttgtacaacgctagcttctctttacactttcttctctgtgcccttaacacgatcccttactttttgtagcggtttcaatttatagagctcaggtcattactggcaagttatctgattcggttattacatattactacaagaaaaagaaacatgaaattctgactcgttgacttgtttatcgaacattgccgaagctatgcctctaactatcgttctcagtttatgacgtatctctctgtatcgaaattttcgtggattatgctttaaaatagattagagccacttgaggcgcacctgaacatgattgcctcgaaacatattagctgtacatccgttttagatcttctagttagcctctgagagtggttctatcttttgaagaaatagtacttactctacattatcaatgtcttttatttcgcgctgccgtgtaagtttggtcgatctttaggtccagtttctacatcatactaaaagcgcgcgtggaggttgcttatgaagaaattggtttagctttggaaattggagcctctcaccgtaaaccaaaaaatctgtcagtactataatttctggtggagaccacatctagtgcaactggctgttgctagaagattccgttggaccaatgtggaatcagagggtatacaagtacgtgaatattaagacattgattgattcaaacatcagaatctatgtattctaatcctatattgtatcaacaataatattattattgatcctgtatgtaaaaggttataaacttaaaatcacacggtttaagataccgaaatgctaagcaaaaacgtttcgtcgagagaaacaaaaacattgagttcatgaaatggaattaacaattgctaactgaaagataacgctgactggtgtaccacttgtcgggggcttattcaagctgacggagtcacttgatctacggtacccgggatgacccataacgcgattaagttgggtaacgccagggtttttccagtcacgcccaatgtgctaaaacggagggaagtgaattccgcggacagtctggcttgaacgagctagagatgagctgagatcggtttacacttagcactcttaaactcatgctttaggttgttaaaacatgtaataattgggtattactaggcagtattgtggtcttgaaaaaaggttttattctacaggctaaacgcctagagaatgtaaagtaaggaagtgggaaagggatggataaaatagttctgattggggcttaatggggctcaagtggctgtgttttctgattgatattggaacctgctgtcatttcgacaattaatatttacttatttggtcaaccccaaataggttgagtcgacttggttgattcaaaaataagtagtcttttgagatctttcaatattagcttaaatattgctatagacagccgacttgtttcattttaacttatgtttgaagcttatcggatcacgagcttatcactagcagttggctttaactccagtaattacaagttagtcgaatagtctgtgctaaaaaaccaatcggttaaaactagaatcgaatcttaagtttaagattcaggcttgttgttaaacctattaccagaaaggacagttggctttggaacaaactcagctaacttgaaactcaaacaacagttttgttcttgccggcttacctcacttgctcagctagtctagcaagaaaacagtgttcttgtcaacttagaactactagcagagttaagcagaatcttgaaggctactctgtgttgaaacaattgacttgcttttcttagcttaaagctagtcaatcttcaggcagaagtaacttgaaaagttagaaacttaagtttaaaaagacaacaagaaacttgcacacaggactgagaaagtaactaagatttgaaactgagtagactaactctacagtagttcacaagggttccaccctactttaactttgacaaggttcaaagtcgaactttaaactgagcttacctctgtggactgtaaactcttgagttaccaaactcagctagcctctcagactgttactggttagcttacacagtcttttactattctaacgtaagcttagactagttcttggaataagtgaatctaagaagttgaggctaaactctcaaatagtgtagactaggcacgaactagaagttagttcacttacgttggtagaaagaaacagttaatagctgaaaagaggtcaactaagccaaacaagtttcttgtcagaagtctaactgagtgaatacagtttatcagaaactgtagctagttcgtcttctcaaaccaacttaccagacttttttaaacttgcagtttttactgtacgagacagtttgttactcttcacagtcagttagacgtaaagtaacttgaataagctgaatactaacgtccacgaggtaggcagactatcctgttactgtctttcagtatgcagtacaaaaagttaaactgttacaatcctcggttaggcaactttgagttaacttagaagttaccactgcaagctcaatcagattcgccctgcagattctaaaaacaaaagctgagctaaagctgctaactgcagttaggtttagatttgagaatagacttggtaaactgcagtaagttttgagaaaagtaactaaaactacagttttcaacttcagtttcgaattgtgagtgttgagtctgagttgaaagactgagttaagtaagttaactcactaggttacgctaaagtgtagctactttggattcaggctcactctgcactagcatatggtgcactctcagtacaatctgctctgatgccgcatagttaagccagccccgacacccgccaacacccgctgacgcgccctgacgggcttgtctgctcccggcatcgcttacagacaagtatgaccgtctcccgggagctgcatgtgtcagaggttttcaccgtcatcaccgaaacgcgcgagacgaaagggcctcgtgatacgcctatttttataggttaatgtcatgataataatggtttcttagacgtcaggtggcacttttcggggaaatgtgcgcggaacccctatttgtttatttttctaaatacattcaaatatgtatccgctaataaagcaataaccctgataaatgcttcaataatattgaaaaaggaagagtatgttactgtcagacattttccgtgtcgcccttattccctttttgcggcattttgccttcctgttttgcacccagaaacgctggtgaaagtaaaagatgctgaagatcagttgggtgcgcattcggttacatcgaactggatctcaacagcggtaagatccttgagagttttcgccccgaagaacgttttccaatgatgagcacttttaaagttctgctatgtggcgcggtattatccattgacgccgggcaagagcaactcggtcgccgcatacaattctcagaatgctgatttggttgagtatcaccagtcacagaaaagcatcttacggatggcatgacagtaagagattatgcaggctgccataaccatgagtgataacactgcggccaacttacttctgacaacgatcggaggaccgaaggagctaaccgcttttttgcacaacatgggatcatgtaactcgccttgatcgttgggaaccggagctgaatgaagccataccaaacgacgagcgtgacaccacgatgcctgtagcaatggcaacaacgttgcgcaaactattaactggcgaactacttactctagcttcccggcaacaattaatagactggatggaggcggataaagttgcaggaccacttctgcgctcggcccttccggctggctggtttattgctgataaatctggagccggtgagcgtgggtctcgcggtatcattgcagcactggggccagatggtaagccctcccgtatcgtagttatctacacgacggggagtcaggcaactatggatgaacgaaatagacagatcgctgagataggtgcctcactgattaagcattggtaactgtcagaccaagtttactcatatatactttagattgatttaaaacttcatttttaatttaaaaggatctaggtgaagatcctttttgataatctcatgaccaaaatcccttaacgtgagttttcgttccactgagcgtcagaccccgtagaaaagatcaaaggatcttcttgagatccttttttgcgcgtaatctgctgcttgcaaacaaaaaaaccaccgctaccagcggtggtttgtttgccggatcaagagctaccaactctttttccgaaggtaactggcttcagcagagcgcagataccaaatactgttcttctagtgtagccgtagttaggccaccacttcaagaactctgtagcaccgcctacatacctcgctctgctaatcctgttaccagtggctgctgccagtggcgataagtcgtgtcttaccgggttggactcaagacgatagttaccggataaggcgcagcggtcgggctgaacggggggttcgtgcacacagcccagcttggagcgaacgacctacaccgaactgagatacctacagcgtgagctatgagaaagcgccacgcttcccgaagggagaaaggcggacaggtatccggtaagcggcagggtcggaacaggagagcgcacgagggagcttccagggggaaacgcctggtatctttatagtcctgtcgggtttcgccacctctgacttgagcgtcgatttttgtgatgctcgtcaggggggcggagcctatggaaaaacgccagcaacgcggcctttttacggttcctggccttttgctggccttttgctcacatgttctttcctgcgttatcccctgattctgtggataaccgtattaccgcctttgagtgagctgataccgctcgccgcagccgaacgaccgagcgcagcgagtcagtgagcgaggaagcggaagagcgcccaatacgcaaaccgcctctccccgcgcgttggccgattcattaatgcagttgtcattttgtttcttctcgtacgagcttgctcctgatcagcctatctcgcagctgtgttaaaccgtctttaagtcaacccacagtctactgcaatcgtattcagaactagccactagactgttacaagtcgaaacctcagttaaccaactagaaactctagttaccaagttagaactgtgagtacgaaaagtctgaaaagcagaaagattcaatagatttgtctgtgttacacaagagttcaacaagtagctgcgttcgtgctagttgtctaggtttaaacgtttaaaaagacaactagtagtttacttcacctctgtattactgacggtttgtgcctcaacagtttacgttaacaaactagtaagcgtctacttcgtggacttgtacttttagtagaaagattgagtgtctggcagttttaaacctaagtcagttagaatctcaatcctctgcctttggtttagaaacagtgtagctgttttgacctagactgagtacaactgttcttgctgcttattcgaaacttgcctaactcactgcaacagacaagtctagtttagaataaagtaggctcagttattcaagtctaacttcagacagtgtttctcagatttgacttaagtagacagtagtgtgtgaaaaaccagactagtttgaatcttagcctgttcagaaaaaactaactgacttgtgtcaggcctcccaaaaaaactaacttcaatcctctgtactattccacaactattagttgttgcaagtctgaagcctagtcttgctttgaccttacaggatagaggttggctcacaactgagtctgtgtgaaaacaaaagtaaagattgccacaagaagtctattctagttagtccttttacgttggatagcaatagtccaatcttagtttgcagctgatgaatatcttgtggtaggggtttgggaaaatcattcgagtttgatgtttttcttggtatttcccactcctcttcagagtacagaagattaagtgagaccttcgtttgtgcggatcccccacacaccatagcttcaaaatgtttctactccttttttactcttccagattttctcggactccgcgcatcgccgtaccacttcaaaacacccaagcacagcatactaaatttcccctctttcttcctctagggtgtcgttaattacccgtactaaaggtttggaaaagaaaaaagagaccgcctcgtttctttttcttcgtcgaaaaaggcaataaaaatttttatcacgtttctttttcttgaaaattttttttttgatttttttctctttcgatgacctcccattgatatttaagttaataaacggtcttcaatttctcaagtttcagtttcatttttcttgttctattacaactttttttacttcttgctcattagaaagaaagcatagcaatctaatctaagggcggtgttgacaattaatcatcggcatagtatatcggcatagtataatacgacaaggtgaggaactaaaccatggccaagttgaccagtgccgttccggtgctcaccgcgcgcgacgtcgccggagcggtcgagttctggaccgaccggctcgggttctcccgggacttcgtggaggacgacttcgccggtgtggtccgggacgacgtgaccctgttcatcagcgcggtccaggaccaggtggtgccggacaacaccctggcctgggtgtgggtgcgcggcctggacgagctgtacgccgagtggtcggaggtcgtgtccacgaacttccgggacgcctccgggccggccatgaccgagatcggcgagcagccgtgggggcgggagttcgccctgcgcgacccggccggcaactgcgtgcacttcgtggccgaggagcaggactgacacgtccgacgcggcccgacgggtccgaggcctcggagatccgtcccccttttcctttgtcgatatcatgtaattagttatgtcacgcttacattcacgccctcccccacatccgctctaaccgaaaaggaaggagttagacaacctgaagtctaggtccctatttatttttttatagttatgttagtattaagaacgttatttatatttcaaatttttcttttttttctgtacagacgcgtgtacgcatgtaacattatactgaaaaccttgcttgagaaggttttgggacgctcgaaggctttaatttgcaagctggagaccaacatgtgagcaaaaggccagcaaaaggccaggaaccgtaaaaaggccgcgaattgtgagcggataacaatttcacacaggaaacagctatgaccatgattacgccaagcttgcatgccgcagaaaggcccacccgaaggtgagccaggtgattacatttgggccctcattagaaaaactcatcgagcatcaagtgaaactgcaatttattcatatcaggattatcaataccatatttttgaaaaagccgtttctgtaatgaaggagaaaactcaccgaggcagttccataggatggcaagatcctggtatcggtctgcgattccgactcgtccaacatcaatacaacctattaatttcccctcgtcaaaaataaggttatcaagtgagaaatcaccatgagtgacgactgaatccggtgagaatggcaaaagcttatgcatttctttccagacttgttcaacaggccagccattacgctcgtcatcaaaatcactcgcaccaaccaaaccgttattcattcgtgattgcgcctgagcgagacgaaatacgcgatcgccgttaaaaggacaattacaaacaggaatcgaatgcaaccggcgcaggaacactgccagcgcatcaacaatattttcacctgaatcaggatattcttctaatacctggaatgctgttttccctgggatcgcagtggtgagtaaccatgcatcatcaggagtacggataaaatgcttgatggtcggaagaggcataaattccgtcagccagtttagcctgaccatctcatctgtaacatcattggcaacgctacctttgccatgtttcagaaacaactctggcgcatcgggcttcccatacaatcgatagattgtcgcacctgattgcccgacattatcgcgagcccatttatacccatataaatcagcatccatgttggaatttaatcgcggcctcgagcaagacgtttcccgttgaatatggctcattttagcttccttagctcctgaaaatctcgataactcaaaaaatacgcccggtagtgatcttatttcattatggtgaaagttggaacctcttacgtgccgatcagcatgcagcaccaccaaaaaaaaacgaaaagtagaagacccacgatttatgtacccatacgatgttcctgactatgcgggtatgaaaaacatcaaaaaaaaccaggtaatgaacctgggtccgaactctaaactgctgaaagaatacaaatcccagctgatcgaactgaacatcgaacagttcgaagcaggtatcggtctgatcctgggtgatgcttacatccgttctcgtgatgaaggtaaaacctactgtatgcagttcgagtggaaaaacaaagcatacatggaccacgtatgtctgctgtacgatcagtgggtactgtccccgccgcacaaaaaagaacgtgttaaccacctgggtaacctggtaatcacctggggcgcccagactttcaaacaccaagctttcaacaaactggctaacctgttcatcgttaacaacaaaaaaaccatcccgaacaacctggttgaaaactacctgaccccgatgtctctggcatactggttcatggatgatggtggtaaatgggattacaacaaaaactctaccaacaaatcgatcgtactgaacacccagtctttcactttcgaagaagtagaatacctggttaagggtctgcgtaacaaattccaactgaactgttacgtaaaaatcaacaaaaacaaaccgatcatctacatcgattctatgtcttacctgatcttctacaacctgatcaaaccgtacctgatcccgcagatgatgtacaaactgccgaacactatctcctccgaaactttcctgaaataaccgcggcggccgccagcttgggcccgaacaaaaactcatctcagaagaggatctgaatagcgccgtcgaccatcatcatcatcatcattgagttttagccttagacatgactgttcctcagttcaagttgggcacttacgagaagaccggtcttgctagattctaatcaagaggatgtcagaatgccatttgcctgagagatgcaggcttcatttttgatacttttttatttgtaacctatatagtataggattttttttgtcattttgtttcttctcgtacgagcttgctcctgatcagcctatctcgcagctgatgaatatcttgtggtaggggtttgggaaaatcattcgagtttgatgtttttcttggtatttcccactcctcttcagagtacagaagattaagtgagccaaccgtgtaggtctacaaactggtattctcgtttgttgtgaatcttgtcttttctgcttactttctaatagattgagtgagttaggtctgtatttgacggctaaagttaaacttaactgttgtgtgggatagttgttagctgtcaagaaaccgttaaacgtagaaagagtaacttttctaaaccgtagtgtctactcaatattgagacaagcagctagtttaagagcagaagctgaagtctgtgagtttggaaactaaaaggtctgtgaacagctaactagaactaggcttgcttttagtattagaataacaacaaacgaggtctattagttgtgggatctagctagttgcctaagaacaaaggactaagtctgcagaaaaacgagtttcaataggtctgtgtgtaacagtgttaaggctactagcagaactgccacctaagtttcgtaagttccacctcagccttcaatattccagacttgaccaactttcagtttttaaactctaccgtactagtagtttctgcacagtattagacctacttacactgtctgtgcttgcaccacaaaccaaatagtagaatctcttttaagtagaacagtcttttcttgttagactaaaacaacgagagacaactgagtgtagggttacagttaaaaaactgtacagtctgctgtacagacgttacacggcagtcctgttccaaacttcacttaagggttagattggctaagttcaacttgtccttttaagttagttcagacctagttagcagacttaggtgggtaagctgagctacaagagttaaaaacaactgctaagactaaagtaagtgtacgtaagcttcaaactagtaagttaaacttagaaacagctaacgtctaaagctagaagtaagctcaatcttagtactgttaacctaagttgctcacgcaccttcgccaactcaacctgcaggtcgactctagagggattcggcgctccaactgtattctaggcctttccacaatttaaaaaaagacctccgcgacggggaattgaaccccggtctaccgcgcgacaagcggtggttctaccactaaactatcacggatttcgatgctgctttcccacttccttactttacattctctaggcgtttagcctgtagaataaaaccttttttcaagactacaatactgctgctagtaataccgaattattacatgttttaacaacttgagagcgggtcgtatcctgtcttccgattgtacaatctcatcttatcagctcatctcatctcttcggaatccccccgggtaccgagctcgaattcactggccgtcgttttacaacgtcgtgactgggaaaaccctggcgttacccaacttaatcgccttgcagcacatcccgggtaccgtagatcaagtgacttcttgagcttgccaagcatcaagtcccaccgacaagtggtacaccagtcagcgttatctttccatacattctttaattcaaattgtcccattcacaactcaatgtttttgtttctctcgacgaaacgtttttgcttagcatttcggtatcttaaaccgtgtgattttaagtctcataaccttctacatacaggatcaataataatattattgttgatacaatataggattagaatacatagattctgatgttcgaatcaatcaatgtcttaatattcacgtacttgtataccctctgattccacattggtccaacggaatcttctagcaacagccagttgcactagatgtggttctccaccagaaattatagtactgacagattttttggtttacggtgagaggctccaatttccaaagctaaaccaatttcttcataagcaacctccacgcgcgcttttagtatgatgtagaaactggacctaaagatcgaccaatacttgcttacacggcagcgcgaaataaaagacattgataatgtagagtaagtactatttcttcaaaagatagaaccactctcagaggctaactagaagatctaaaacggatgtacagctaatatgtttcgaggcaatcatgttcaggtgcgcctcaagtggctcttaatctattttaaagcataatccacgaaaatttcgatacagagagatacgtcataaactgagaacgatagttagaggcatagcttcggcaatgttcgataaacaagtcaacgagtcagaatttcatgtttctttttttcttgtagtaatatgtaataaccgaatcagataacttgccagtaatgacctgagctctataaattgaaaccgctacaaaaagtaagggatcgtgttaagggcacagagaagaaagtgtaaagaagtaagctagcgttgtacaacaactcaaaagcatggagcccggtgggtactagataataagaacaaggtttcagtgagtttacaagttcataaaacacggaacctggcaacatagcataccagtaaatcgatttgaagaactctattaggctggcagcctctctcaatcaatagtctaaaacggttaagagacacaatcatctctaggtgtcgttgtggtctaccatcgaaacactggaaatcaatccaatgaacttcaagtgatatatttttcaagagtcttcaaagaccaagcaatgttcttattgattgataattttaaagttgaagtctaaatttgtagcagtgaagtctgagcagggggaatgttaagccatatctaatagtacctattagacgcttgattgccgtttgcaaatatgccatgtttctttgaaaccaaaaccacgtaagggggcttaagttttcggcttaggattgttacggagctcaaaccaaataaggcagagagcatacaaaacgtttattaaaaagaacaactacttctggatagctcaattatcttttgtttcttttgaggcgtgcccttcatgacatgacattgcatccatacaattaatagtagatagggaggataagtttgctagtgtgagcatttagagcaaggggtcagtttcctccatctgcctactctgctccatttaaaagcgagccgattgccataagcttgctcacgtagataggacttcaaaaagacgttaagaggctgctctctagaatacgatggaataaacaaatctcgttagtttcttgaaacggaaagtacatgtaggataattgctggatcattcggaagctaaaattggtcctttccccaaagacaatcgactatagttcctgaagcttcctgaggtgagctggatagaaccagaagatagtacctataacgtcaacaacaactgaaactacagatggaagagctaggcatatccataggggatagccatagaaaactttgattgacgcagcaaatcgagacgttaacctgaagctgccataacttcaggaacttgaacacaagagtggcattaaaattcctatgttgcatttcagaaataccacaagtaaactgagcaattaacttgttcataccgacactacaagttaactacaaaccgagacccttatatgcagactgataatacagaatgatacgtatcactcctaagactaggaacatacagagcttttgagcttttggtattttcaagttatttgaaaaattaaatctttatactagggacgaggttcgtgacaaaagaagacaatcatgagactcaccgtcttgtcatctacaagtttcaataagcttccaacttgagtaacctagccatgtgattagtaactttcgaagcatgattcagaacgttctgctctgccgcgtcaaaaagggcagctacttgaagttaaaagaaagatcaaaaattcctgagcattgttcaaacagctataaatcaaaacagaataagaaataaggccatcacttaccaaggaaaaacaaaaaagttcgagagatcaacaatgcacttcagggacgcacctgaacctttacattagtgtaaaaataaattattagattgttcggttcgagcgttagccatatacaaaatgcggccttcaaagtcattggaaaagagctgctttaggacctctccaaaagctaaattgacaaacaggctacctcatatctccatcatctttgccaaccttttcagtaacaaaaaacaaaagcaatatcgtttatttgtccattggacgatgaagaaaagatcaaacttagtctaggtgaggctaatcaataaagtttgttgatcctactaattacgatcctatcctagccttagcatttacatctaaaggtctggtttcacattctaattgtgaaatgatctagcaagctctccctagcataattattcacctccaaatttgtgtacttggttcatcgttcactgaatgagaatgagaaagattcaaaattgagaagtgtgttcgaacacttcaactgtcgcactgttgctttctacagatatacgcataacaaatacattagttgcttaaagggcagcttcttttaatgtgtatcagtatagtcaaggactcagctacgaacaactcacaatagtctttttcaccacgacccacaagttcgcacaagaattctaaataccttgacgaatagtaaatatcggatgttcagccgtcagcgaaatgatttttgtgctaatacaatttctaaggcaattgacacaggttggaaaaagtgtgggtattcggctgatattgtggaactgtaatcatcgagctaccagttctggtttgttacttgttcgtctcattgtcgtggataagaacaaataatgtaccggtttattcgaccaccaatcagcgcgtaaagcacgcagcagaccaagcaagtagccggatgaagcagctacgttacccaaccgtttatcctacccaagtaggcgagttcaatttgtctcacgcgagatgcactcagtgctacaagattgaatgggtaattagaagcatgttagtttctacaatgatcgattcatagtatccggcgacaattgatcttactgaagaaaggaggcagacacgcttaccgaaaaaagtgcctacctagcaatatttgatgatcgtccaatggacaaataaacgatattgcttttgttttttgttactgaaaaggttggcaaagatgatggagatatgaggtagcctgtttgtcaatttagcttttggagaggtcctaaagcagctcttttccaatgactttgaaggccgcattttgtatatggctaacgctcgaaccgaacaatctaataatttatttttacactaatgtaaaggttcaggtgcgtccctgaagtgcattgttgatctctcgaacttttttgtttttccttggtaagtgatggccttatttcttattctgttttgatttatagctgtttgaacaatgctcaggaatttttgatctttcttttaacttcaagtagctgccctttttgacgcggcagagcagaacgttctgaatcatgcttcgaaagttactaatcacatggctaggttactcaagttggaagcttattgaaacttgtagatgacaagacggtgagtctcatgattgtcttcttttgtcacgaacctcgtccctagtataaagatttaatttttcaaataacttgaaaataccaaaagctcaaaagctctgtatgttcctagtcttaggagtgatacgtatcattctgtattatcagtctgcatataagggtctcggtttgtagttaacttgtagtgtcggtatgaacaagttaattgctcagtttacttgtggtatttctgaaatgcaacataggaattttaatgccactcttgtgttcaagttcctgaagttatggcagcttcaggttaacgtctcgatttgctgcgtcaatcaaagttttctatggctatcccctatggatatgcctagctcttccatctgtagtttcagttgttgttgacgttataggtactatcttctggttctatccagctcacctcaggaagcttcaggaactatagtcgattgtctttggggaaaggaccaattttagcttccgaatgatccagcaattatcctacatgtactttccgtttcaagaaactaacgagatttgtttattccatcgtattctagagagcagcctcttaacgtctttttgaagtcctatctacgtgagcaagcttatggcaatcggctcgcttttaaatggagcagagtaggcagatggaggaaactgaccccttgctctaaatgctcacactagcaaacttatcctcccctatctactattaattgtatggatgcaatgtcatgtcatgaagggcacgcctcaaaagaaacaaaagataattgagctatccagaagtagttgttctttttaataaacgttttgtatgctctctgccttatttggtttgagctccgtaacaatcctaagccgaaaacttaagcccccttacgtggttttggtttcaaagaaacatggcatatttgcaaacggcaatcaagcgtctaataggtactattagatatggcttaacattccccctgctcagacttcactgctacaaatttagacttcaactttaaaattatcaatcaataagaacattgcttggtctttgaagactcttgaaaaatatatcacttgaagttcattggattgatttccagtgtttcgatggtagaccacaacgacacctagagatgattgtgtctcttaaccgttttagactattgattgagagaggctgccagcctaatagagttcttcaaatcgatttactggtatgctatgttgccaggttccgtgttttatgaacttgtaaactcactgaaaccttgttcttattatctagtacccaccgggctccatgcttttgagttgttgtacaacgctagcttacttctttacactttcttctctgtgcccttaacacgatcccttactttttgtagcggtttcaatttatagagctcaggtcattactggcaagttatctgattcggttattacatattactacaagaaaaaaagaaacatgaaattctgactcgttgacttgtttatcgaacattgccgaagctatgcctctaactatcgttctcagtttatgacgtatctctctgtatcgaaattttcgtggattatgctttaaaatagattaagagccacttgaggcgcacctgaacatgattgcctcgaaacatattagctgtacatccgttttagatcttctagttagcctctgagagtggttctatcttttgaagaaatagtacttactctacattatcaatgtcttttatttcgcgctgccgtgtaagcaagtattggtcgatctttaggtccagtttctacatcatactaaaagcgcgcgtggaggttgcttatgaagaaattggtttagctttggaaattggagcctctcaccgtaaaccaaaaaatctgtcagtactataatttctggtggagaaccacatctagtgcaactggctgttgctagaagattccgttggaccaatgtggaatcagagggtatacaagtacgtgaatattaagacattgattgattcgaacatcagaatctatgtattctaatcctatattgtatcaacaataatattattattgatcctgtatgtaaaaggttatgagacttaaaatcacacggtttaagataccgaaatgctaagcaaaaacgtttcgtcgagagaaacaaaaacattgagttgcgttataaaatagatagtaacagacttgctaactgagcattcaagactacgttctatttcaggggcttattcaagctgacggagcacatatgcaacaatactacggaaacgacccataacgctgaaacagccgtggaactcgatggaactttcttgccgatttgttgcccaatgtgctccaaaacggagggaagtctttccgcggacagtctggcttgaacgggctagagatggcagagatcggtttgacttagcactcttaaattcatgctttagggttatgtcgcgagacattaatattgggcaagccagggcagatttggctatattcaggctggtttaggcgatccaaaaaggattatgtaagggattttgactgggatggataaaatagttctgattggggcttaatggggctcaagtggctgtgttttctgattgatattggaacctgctgtcatttcgacaattaatatttacttattttggtcaaccccaaataggttgatttcatacttggttcattcaaaaataagtagtcttttgagatctttcaatattataataaatatactataacagccgacttgtttcattttcgcgaatgttcccccagcttatcggatcccgagctcgaattcgctagccagatcggcttaagacaaccaaacccagaactagtcgaatagtctgtgctaaaaaaccaatcggttaaaactagaatcgatctttaagtttaagattcaggcttgttgttaaacctattgccagaaaggacagttgactttggaacaaactcagctaacttgaaactcaaacaacagttttgttacttgccggcttacctcacttgctcagctagtctagcaagaaaacagtgtttcttgtcaacttagaactaatagcagagttaagcagaatcttgaaggctactctgtatttgaaaaccaattgacttgctttttacttagccttacttgctagtcaatcttcagggcagaagtaacttgaaaagttagaaaccaagtttaaaaagacaacaagaaacttgcacacaggactgagaaagtaactaagatttgaaactgagtagactaactctaccagtggttcacaagggttccaccctactttaacttttgacaagagttcaaagtcgaactttaaactgagcttacctctgtggactgtaaactcttgagttaccaaacttttgagctagcctctcagactgttactggttagctctacacagtcttttactattctaacgtaagcttagactagttcttggaataagtgaatctaagaagttgaggctaaactctcaaatagtgtagactaggcacgaactagaagttagtttcttacgttggtagaaaaaaacagttgagtaggctgaaaaggtcaactaagccaaacaagtttcttgtcagaagtctaactgagtgaatacagttgtattcagaaactgtagctagttcgtcttctcaaaccaacttaccagacttttttaaacttgcagtttttactgtacgagacagtttgttactcttcacagtcagttagacgtaaagtaacttgaataagctgaatactaatgtccacgaggtaggcagactatcctgttactgtcttttttagtctgcagtacaaaaagttaaactgttagcaatcctcggttaggcaactttgagttaacttagagttaccactgcaagctcaatagattcgccctgcagattctaaaacaaaagctgagctaaagctgctaactgcagttaggtttagatttgagaatagacttggtaaactgcagtaacttacgagaaagtaactaaaactacagttttcaacttcagttttcgaactgtagttgtcttagtctgagttgaaagactgagttaagtaagttaactcactaggttacgctaaagtgtagctactttggattcaggctcactctgcactagcatatggtgcactctcagtacaatctgctctgatgccgcatagttaagccagccccgacacccgccaacacccgctgacgcgccctgacgggcttgtctgctcccggcatccgcttacagacaagctgtgaccgtctccgggagctgcatgtgtcagaggttttcaccgtcatcaccgaaacgcgcgagacgaaagggcctcgtgatacgcctatttttataggttaatgtcatgataataatggtttcttagacgtcaggtggcacttttcggggaaatgtgcgcggaacccctatttgtttatttttctaaatacattcaaatatgtatccgctcatgagacaataaccctgataaatgcttcaataatattgaaaaaggaagagtatgagtattcaacatttccgtgtcgcccttattcccttttttgcggcattttgccttcctgtttttgctcacccagaaacgctggtgaaagtaaaagatgctgaagatcagttgggtgcacgagtgggttacatcgaactggatctcaacagcggtaagatccttgagagttttcgcccgaagaacgttttccaatgatgagcacttttaaagttctgctatgtggcgcggtattatcccgtattgacgccgggcaagagcaactcggtcgccgcatacactattctcagaatgacttggttgagtactcaccagtcacagaaaagcatcttacggatggcatgacagtaagagaattatgcagtgctgccataaccatgagtgataacactgcggccaacttacttctgacaacgatcggaggaccgaaggagctaaccgcttttttgcacaacatgggggatcatgtaactcgccttgatcgttgggaaccggagctgaatgaagccataccaaacgacgagcgtgacaccacgatgcctgtagcaatggcaacaacgttgcgcaaactattaactggcgaactacttactctagcttcccggcaacaattaatagactggatggaggcggataaagttgcaggaccacttctgcgctcggcccttccggctggctggtttattgctgataaatctggagccggtgagcgtgggtctcgcggtatcattgcagcactggggccagatggtaagccctcccgtatcgtagttatctacacgacggggagtcaggcaactatggatgaacgaaatagacagatcgctgagataggtgcctcactgattaagcattggtaactgtcagaccaagtttactcatatatactttagattgatttaaaacttcatttttaatttaaaaggatctaggtgaagatcctttttgataatctcatgaccaaaatcccttaacgtgagttttcgttccactgagcgtcagaccccgtagaaaagatcaaaggatcttcttgagatcctttttttctgcgcgtaatctgctgcttgcaaacaaaaaaaccaccgctaccagcggtggtttgtttgccggatcaagagctaccaactctttttccgaaggtaactggcttcagcagagcgcagataccaaatactgttcttctagtgtagccgtagttaggccaccacttcaagaactctgtagcaccgcctacatacctcgctctgctaatcctgttaccagtggctgctgccagtggcgataagtcgtgtcttaccgggttggactcaagacgatagttaccggataaggcgcagcggtcgggctgaacggggggttcgtgcacacagcccagcttggagcgaacgacctacaccgaactgagatacctacagcgtgagctatgagaaagcgccacgcttcccgaagggagaaaggcggacaggtatccggtaagcggcagggtcggaacaggagagcgcacgagggagcttccaggggaaacgcctggtatctttatagtcctgtcgggtttcgccacctctgacttgagcgtcgatttttgtgatgctcgtcaggggggcggagcctatggaaaaacgccagcaacgcggcctttttacggttcctggccttttgctggccttttgctcacatgttctttcctgcgttatcccctgattctgtggataaccgtattaccgcctttgagtgagctgataccgctcgccgcagccgaacgaccgagcgcagcgagtcagtgagcgaggaagcggaagagcgcccatacgcaaaccgcctctccccgcgcgttggccgattcattaatgcagttgtcattttgtttcttctcgtacgagcttgctcctgatcagcctatctcgcagctgtgttaaaccgtctttaagtcaacccacagtctactgcaatcgtattcagaactagccactagactgttacaagtcgaaacctcagttaaccaactagaaactctagttaccaagttagaactgtgagtacgaaaagtctgaaaagcagaaagattcaatagatttgtctgtgttacacaagagttcaacaagtagctgcgttcgtgctagttgtctaggtttaaacgtttaaaaagacaactagtagtttacttcacctctgtattactgacggtttgtgcctcaacagtttacgttaacaaactagtaagcgtctacttcgtggacttgtacttttagtagaaagattgagtgtctggcagttttaaacctaagtcagttagaatctcaatcctctgcctttggtttagaaacagtgtagctgttttgacctagactgagtacaactgttcttgctgcttattcgaaacttgcctaactcactgcaacagacaagtctagtttagaataaagtaggctcagttattcaagtctaacttcagacagtgtttctcagatttgacttaagtagacagtagtgtgtgaaaaaccagactagtttgaatcttagcctgttcagaaaaaactaactgacttgtgtcaggcctcccaaaaaaactaacttcaatcctctgtactattccacaactattagttgttgcaagtctgaagcctagtcttgctttgaccttacaggatagaggttggctcacaactgagtctgtgtgaaaacaaaagtaaagattgccacaagaagtctattctagttagtccttttacgttggatagcaatagtccaatcttagtttgcagctgatgaatatcttgtggtaggggtttgggaaaatcattcgagtttgatgtttttcttggtatttcccactcctcttcagagtacagaagattaagtgagaccttcgtttgtgcggatcccccacacaccatagcttcaaaatgtttctactccttttttactcttccagattttctcggactccgcgcatcgccgtaccacttcaaaacacccaagcacagcatactaaatttcccctctttcttcctctagggtgtcgttaattacccgtactaaaggtttggaaaagaaaaaagagaccgcctcgtttctttttcttcgtcgaaaaaggcaataaaaatttttatcacgtttctttttcttgaaaattttttttttgatttttttctcttcgatgacctcccattgatatttaagttaataaacggtcttcaatttctcaagtttcagtttcatttttcttgttctattacaactttttttacttcttgctcattagaaagaaagcatagcaatctaatctaagggcggtgttgacaattaatcatcggcatagtatatcggcatagtataatacgacaaggtgaggaactaaaccatggccaagttgaccagtgccgttccggtgctcaccgcgcgcgacgtcgccggagcggtcgagttctggaccgaccggctcgggttctcccgggacttcgtggaggacgacttcgccggtgtggtccgggacgacgtgaccctgttcatcagcgcggtccaggaccaggtggtgccggacaacaccctggcctgggtgtgggtgcgcggcctggacgagctgtacgccgagtggtcggaggtcgtgtccacgaacttccgggacgcctccgggccggccatgaccgagatcggcgagcagccgtgggggcgggagttcgccctgcgcgacccggccggcaactgcgtgcacttcgtggccgaggagcaggactgacacgtccgacgcggcccgacgggtccgaggcctcggagatccgtcccccttttcctttgtcgatatcatgtaattagttatgtcacgcttacattcacgccctccccccacatccgctctaaccgaaaaggaaggagttagacaacctgaagtctaggtccctatttatttttttatagttatgttagtattaagaacgttatttatatttcaaatttttcttttttttctgtacagacgcgtgtacgcatgtaacattatactgaaaaccttgcttgagaaggttttgggacgctcgaaggctttaatttgcaagctggagaccaacatgtgagcaaaaggccagcaaaaggccaggaaccgtaaaaaggccgcgaattgtgagcggataacaatttcacacaggaaacagctatgaccatgattacgccaagcttgcatgccgcagaaaggcccacccgaaggtgagccaggtgattacatttgggccctcattagaaaaactcatcgagcatcaagtgaaactgcaatttattcatatcaggattatcaataccatatttttgaaaaagccgtttctgtaatgaaggagaaaactcaccgaggcagttccataggatggcaagatcctggtatcggtctgcgattccgactcgtccaacatcaatacaacctattaatttcccctcgtcaaaaataaggttatcaagtgagaaatcaccatgagtgacgactgaatccggtgagaatggcaaaagcttatgcatttctttccagacttgttcaacaggccagccattacgctcgtcatcaaaatcactcgcaccaaccaaaccgttattcattcgtgattgcgcctgagcgagacgaaatacgcgatcgccgttaaaaggacaattacaaacaggaatcgaatgcaaccggcgcaggaacactgccagcgcatcaacaatattttcacctgaatcaggatattcttctaatacctggaatgctgttttccctgggatcgcagtggtgagtaaccatgcatcatcaggagtacggataaaatgcttgatggtcggaagaggcataaattccgtcagccagtttagcctgaccatctcatctgtaacatcattggcaacgctacctttgccatgtttcagaaacaactctggcgcatcgggcttcccatacaatcgatagattgtcgcacctgattgcccgacattatcgcgagcccatttatacccatataaatcagcatccatgttggaatttaatcgcggcctcgagcaagacgtttcccgttgaatatggctcattttagcttccttagctcctgaaaatctcgataactcaaaaatacgcccggtagtgatcttatttcattatggtgaaagttggaacctcttacgtgccgatcagcatgcagcaccaccaaaaaaacgaaaagtagaagacccacgatttatgtacccatacgatgttcctgactatgcgggtatgaaaaacatcaaaaaaaaccaggtaatgaacctgggtccgaactctaaactgctgaaagaatacaaatcccagctgatcgaactgaacatcgaacagttcgaagcaggtatcggtctgatctgggtgatgcttacatccgttctcgtgatgaaggtaaaacctactgtatgcagttcgagtggaaaaacaaagcatacatggaccacgtatgtctgctgtacgatcagtgggtactgtccccgccgcacaaaaaagaacgtgttaaccacctgggtaacctggtaatcaccggcgcccagactttcaaacaccaagctttcaacaaactggctaacctgttcatcgttaacaacaaaaaaaccatcccgaacaacctggttgaaaactacctgaccccgatgtctctggcatactggttcatggatgatggtggtaaatgggattacaacaaaaactctaccaacaaatcgatcgtactgaacacccagtctttcactttcgaagaagtagaatacctggttaagggtctgcgtaacaaattccaactgaactgttacgtaaaaatcaacaaaaacaaaccgatcatctacatcgattctatgtcttacctgatcttctacaacctgatcaaaccgtacctgatcccgcagatgatgtacaaactgccgaacactatctcctccgaaactttcctgaaataaccgcggcggccgccagcttgggcccgaacaaaaactcatctcagaagaggatctgaatagcgccgtcgaccatcatcatcatcatcattgagttttagccttagacatgactgttcctcagttcaagttgggcacttacgagaagaccggtcttgctagattctaatcaagaggatgtcagaatgccatttgcctgagagatgcaggcttcatttttgatacttttttatttgtaacctatatagtataggattttttttgtcattttgtttcttctcgtacgagcttgctcctgatcagcctatctcgcagctgatgaatatcttgtggtaggggtttgggaaaatcattcgagtttgatgtttttcttggtatttcccactcctcttcagagtacagaagattaagtgagccaaccgtgtaggtctacaaactggtattctcgtttgttgtgaatcttgtcttttctgcttactttctaatagattgagtgagttaggtctgtatttgacggctaaagttaaacttaactgttgtgtgggatagttgttagctgtcaagaaaccgttaaacgtagaaagagtaacttttctaaaccgtagtgtctactcaatattgagacaagcagctagtttaagagcagaagctgaagtctgtgagtttggaaactaaaaggtctgtgaacagctaactagaactaggcttgcttttagtattagaataacaacaaacgaggtctattagttgtgggatctagctagttgcctaagaacaaaggactaagtctgcagaaaaacgagtttcaataggtctgtgtgtaacagtgttaaggctactagcagaactgccacctaagtttcgtaagttccacctcagccttcaatattccagacttgaccaactttcagtttttaaactctaccgtactagtagtttctgcacagtattagacctacttacactgtctgtgcttgcaccacaaaccaaatagtagaatctcttttaagtagaacagtcttttcttgttagactaaaacaacgagagacaactgagtgtagggttacagttaaaaaactgtacagtctgctgtacagacgttacatgcagtcctgttccaaacttcacttaagggttagattggctaagttcaacttgtccttttaagttagttcagacctagttagcagacttaggtgggtaagctgagctacaagagttaaaaacaactgctaagactaaagaagtgtacgtaagcttcaaactagtaagttaaacttagaaacagctaacgtctaaagctagaagtaagctcaatcttagtactgttaacctaagttgctcacgcaccttcgccaactcaacctgcaggtcgactctagaggggattcggcgctccaactgtattctaggcctttccacaatttaaaaaaagacctccgcgacgggaattgaaccccggtctaccgcgcgacaagcggtggttctaccactaaactatcacggatttcgatgctgctttcccacttccttactttacattctctaggcgtttagcctgtagaataaaaccttttttcaagactacaatactgctgctagtaataccgaattattacatgttttaacaacttgagagcgggtcgtatcctgtcttccgattgtacaatctcatcttatcagctcatctcatctcttcggaatccccccgggtaccgagctcgaattcactggccgtcgttttacaacgtcgtgactggaaaaccctggcgttacccaacttaatcgccttgcagcacatcccgggtaccgtagatcaagtgacttcttgagcttgccaagcatcaagtcccaccgacaagtggtacaccagtcagcgttatctttccatacattctttaattcaaattgtcccattcacaactcaatgtttttgtttctctcgacgaaacgtttttgcttagcatttcggtatcttaaaccgtgtgattttaagtctcataaccttctacatacaggatcaataataatattattgttgatacaatataggattagaatacatagattctgatgttcgaatcaatcaatgtcttaatattcacgtacttgtataccctctgattccacattggtccaacggaatcttctagcaacagccagttgcactagatgtggttctccaccagaaattatagtactgacagattttttggtttacggtgagaggctccaatttccaaagctaaaccaatttcttcataagcaacctccacgcgcgcttttagtatgatgtagaaactggacctaaagatcgaccaatacttgcttacacggcagcgcgaaataaaagacattgataatgtagagtaagtactatttcttcaaaagatagaaccactctcagaggctaactagaagatctaaaacggatgtacagctaatatgtttcgaggcaatcatgttcaggtgcgcctcaagtggctcttaatctattttaaagcataatccacgaaaatttcgatacagagagatacgtcataaactgagaacgatagttagaggcatagcttcggcaatgttcgataaacaagtcaacgagtcagaatttcatgtttctttttcttgtagtaatatgtaataaccgaatcagataacttgccagtaatgacctgagctctataaattgaaaccgctacaaaaagtaagggatcgtgttaagggcacagagaagaaagtgtaaagaagtaagctagcgttgtacaacaactcaaaagcatggagcccggtgggtactagataataagaacaaggtttcagtgagtttacaagttcataaaacacggaacctggcaacatagcataccagtaaatcgatttgaagaactctattaggctggcagcctctctcaatcaatagtctaaaacggttaagagacacaatcatctctaggtgtcgttgtggtctaccatcgaaacactggaaatcaatccaatgaacttcaagtgatatatttttcaagagtcttcaaagaccaagcaatgttcttattgattgataattttaaagttgaagtctaaatttgtagcagtgaagtctgagcagggggaatgttaagccatatctaatagtacctattagacgcttgattgccgtttgcaaatatgccatgtttctttgaaaccaaaaccacgtaagggggcttaagttttcggcttaggattgttacggagctcaaaccaaataaggcagagagcatacaaaacgtttattaaaaagaacaactacttctggatagctcaattatcttttgtttcttttgaggcgtgcccttcatgacatgacattgcatccatacaattaatagtagataggggaggataagtttgctagtgtgagcatttagagcaaggggtcagtttcctccatctgcctactctgctccatttaaaagcgagccgattgccataagcttgctcacgtagataggacttcaaaaagacgttaagaggctgctctctagaatacgatggaataaacaaatctcgttagtttcttgaaacggaaagtacatgtaggataattgctggatcattcggaagctaaaattggtcctttccccaaagacaatcgactatagttcctgaagcttcctgaggtgagctggatagaaccagaagatagtacctataacgtcaacaacaactgaaactacagatggaagagctaggcatatccataggggatagccatagaaaactttgattgacgcagcaaatcgagacgttaacctgaagctgccataacttcaggaacttgaacacaagagtggcattaaaattcctatgttgcatttcagaaataccacaagtaaactgagcaattaacttgttcataccgacactacaagttaactacaaaccgagacccttatatgcagactgataatacagaatgatacgtatcactcctaagactaggaacatacagagcttttgagcttttggtattttcaagttatttgaaaaattaaatctttatactagggacgaggttcgtgacaaaagaagacaatcatgagactcaccgtcttgtcatctacaagtttcaataagcttccaacttgagtaacctagccatgtgattagtaactttcgaagcatgattcagaacgttctgctctgccgcgtcaaaaagggcagctacttgaagttaaaagaaagatcaaaaattcctgagcattgttcaaacagctataaatcaaaacagaataagaaataaggccatcacttaccaaggaaaaacaaaaaagttcgagagatcaacaatgcacttcagggacgcacctgaacctttacattagtgtaaaaataaattattagattgttcggttcgagcgttagccatatacaaaatgcggccttcaaagtcattggaaagagctgctttaggacctctccaaaagctaaattgacaaacaggctacctcatatctccatcatctttgccaaccttttcagtaacaaaaaacaaaagcaatatcgtttatttgtccattggacgatgaagaaaagatcaaacttagtctaggtgaggctaatcaataaagtttgttgatcctactaattacgatcctatcctagccttagcatttacatctaaaggtctggtttcacattctaattgtgaaatgatctagcaagctctccctagcataattattcacctccaaatttgtgtacttggttcatcgttcactgaatgagaatgagaaagattcaaaattgagaagtgtgttcgaacacttcaactgtcgcactgttgctttctacagatatacgcataacaaatacattagttgcttaaagggcagcttcttttaatgtgtatcagtatagtcaaggactcagctacgaacaactcacaatagtctttttcaccacgacccacaagttcgcacaagaattctaaataccttgacgaatagtaaatatcggatgttcagccgtcagcgaaatgatttttgtgctaatacaatttctaaggcaattgacacaggttggaaaagtgtgggtattcggctgatattgtggaactgtaatcatcgagctaccagttctggtttgttacttgttcgtctcattgtcgtggataagaacaaataatgtaccggtttattcgaccaccaatcagcgcgtaaagcacgcagcagaccaagcaagtagccggatgaagcagctacgttacccaaccgtttatcctacccaagtaggcgagttcaatttgtctcacgcgagatgcactcagtgctacaagattgaatgggtaattagaagcatgttagtttctacaatgatcgattcatagtatccggcgacaattgatcttactgaagaaaggaggcagacacgcttaccgaaaaaagtgcctacctagcaatatttgatgatcgtccaatggacaaataaacgatattgcttttgttttttgttactgaaaaggttggcaaagatgatggagatatgaggtagcctgtttgtcaatttagcttttggagaggtcctaaagcagctcttttccaatgactttgaaggccgcattttgtatatggctaacgctcgaaccgaacaatctaataatttatttttacactaatgtaaaggttcaggtgcgtccctgaagtgcattgttgatctctcgaacttttttgtttttccttggtaagtgatggccttatttcttattctgttttgatttatagctgtttgaacaatgctcaggaatttttgatctttcttttaacttcaagtagctgccctttttgacgcggcagagcagaacgttctgaatcatgcttcgaaagttactaatcacatggctaggttactcaagttggaagcttattgaaacttgtagatgacaagacggtgagtctcatgattgtcttcttttgtcacgaacctcgtccctagtataaagatttaatttttcaaataacttgaaaataccaaaagctcaaaagctctgtatgttcctagtcttaggagtgatacgtatcattctgtattatcagtctgcatataagggtctcggtttgtagttaacttgtagtgtcggtatgaacaagttaattgctcagtttacttgtggtatttctgaaatgcaacataggaattttaatgccactcttgtgttcaagttcctgaagttatggcagcttcaggttaacgtctcgatttgctgcgtcaatcaaagttttctatggctatcccctatggatatgcctagctcttccatctgtagtttcagttgttgttgacgttataggtactatcttctggttctatccagctcacctcaggaagcttcaggaactatagtcgattgtctttggggaaaggaccaattttagcttccgaatgatccagcaattatcctacatgtactttccgtttcaagaaactaacgagatttgtttattccatcgtattctagagagcagcctcttaacgtctttttgaagtcctatctacgtgagcaagcttatggcaatcggctcgcttttaaatggagcagagtaggcagatggaggaaactgaccccttgctctaaatgctcacactagcaaacttatcctcccctatctactattaattgtatggatgcaatgtcatgtcatgaagggcacgcctcaaaagaaacaaaagataattgagctatccagaagtagttgttctttttaataaacgttttgtatgctctctgccttatttggtttgagctccgtaacaatcctaagccgaaaacttaagcccccttacgtggttttggtttcaaagaaacatggcatatttgcaaacggcaatcaagcgtctaataggtactattagatatggcttaacattccccctgctcagacttcactgctacaaatttagacttcaactttaaaattatcaatcaataagaacattgcttggtctttgaagactcttgaaaaatatatcacttgaagttcattggattgatttccagtgtttcgatggtagaccacaacgacacctagagatgattgtgtctcttaaccgttttagactattgattgagagaggctgccagcctaatagagttcttcaaatcgatttactggtatgctatgttgccaggttccgtgttttatgaacttgtaaactcactgaaaccttgttcttattatctagtacccaccgggctccatgcttttgagttgttgtacaacgctagcttacttctttacactttcttctctgtgcccttaacacgatcccttactttttgtagcggtttcaatttatagagctcaggtcattactggcaagttatctgattcggttattacatattactacaagaaaaaaagaaacatgaaattctgactcgttgacttgtttatcgaacattgccgaagctatgcctctaactatcgttctcagtttatgacgtatctctctgtatcgaaattttcgtggattatgctttaaaatagattaagagccacttgaggcgcacctgaacatgattgcctcgaaacatattagctgtacatccgttttagatcttctagttagcctctgagagtggttctatcttttgaagaaatagtacttactctacattatcaatgtcttttatttcgcgctgccgtgtaagcaagtattggtcgatctttaggtccagtttctacatcatactaaaagcgcgcgtggaggttgcttatgaagaaattggtttagctttggaaattggagcctctcaccgtaaaccaaaaaatctgtcagtactataatttctggtggagaaccacatctagtgcaactggctgttgctagaagattccgttggaccaatgtggaatcagagggtatacaagtacgtgaatattaagacattgattgattcgaacatcagaatctatgtattctaatcctatattgtatcaacaataatattattattgatcctgtatgtagaaggttatgagacttaaaatcacacggtttaagataccgaaatgctaagcaaaaacgtttcgtcgagagaaacaaaaacattgagttgtgaatgggacaatttgaattaaagaatgtatggaaagataacgctgactggtgtaccacttgtcggtgggacttgatgcttggcaagctcaagaagtcacttgatctacggtacccgggatgtgctgcaaggcgattaagttgggtaacgccagggttttcccagtcacgacgttgtaaaacgacggccagtgaattcgagctcggtacccggggattccgaagagatgagatgagctgataagatgagattgtacaatcggaagacaggatacgacccgctctcaagttgttaaaacatgtaataattcggtattactagcagcagtattgtagtcttgaaaaaaggttttattctacaggctaaacgcctagagaatgtaaagtaaggaagtgggaaagcagcatcgaaatccgtgatagtttagtggtagaaccaccgcttgtcgcgcggtagaccggggttcaattccccgtcgcggaggtctttttttaaattgtggaaaggcctagaatacagttggagcgccgaatcccctctagagtcgacctgcaggttgagttggcgaaggtgcgtgagcaacttaggttaacagtactaagattgagcttacttctagctttagacgttagctgtttctaagtttaacttactagtttgaagcttacgtacacttactttagtcttagcagttgtttttaactcttgtagctcagcttacccacctaagtctgctaactaggtctgaactaacttaaaaggacaagttgaacttagccaatctaacccttaagtgaagtttggaacaggactgccgtgtaacgtctgtacagcagactgtacagttttttaactgtaaccctacactcagttgtctctcgttgttttagtctaacaagaaaagactgttctacttaaaagagattctactatttggtttgtggtgcaagcacagacagtgtaagtaggtctaatactgtgcagaaactactagtacggtagagtttaaaaactgaaagttggtcaagtctggaatattgaaggctgaggtggaacttacgaaacttaggtggcagttctgctagtagccttaacactgttacacacagacctattgaaactcgtttttctgcagacttagtcctttgttcttaggcaactagctagatcccacaactaatagacctcgtttgttgttattctaatactaaaagcaagcctagttctagttagctgttcacagaccttttagtttccaaactcacagacttcagcttctgctcttaaactagctgcttgtctcaatattgagtagacactacggtttagaaaagttactctttctacgtttaacggtttcttgacagctaacaactatcccacacaacagttaagtttaactttagccgtcaaatacagacctaactcactcaatctattagaaagtaagcagaaaagacaagattcacaacaaacgagaataccagtttgtagacctacacggttggctcacttaatcttctgtactctgaagaggagtgggaaataccaagaaaaacatcaaactcgaatgattttcccaaacccctaccacaagatattcatcagctgcgagataggctgatcaggagcaagctcgtacgagaagaaacaaaatgacaaaaaaaatcctatactatataggttacaaataaaaaagtatcaaaaatgaagcctgcatctctcaggcaaatggcattctgacatcctcttgattagaatctagcaagaccggtcttctcgtaagtgcccaacttgaactgaggaacagtcatgtctaaggctaaaactcaatgatgatgatgatgatggtcgacggcgctattcagatcctcttctgagatgagtttttgttcgggcccaagctggcggccgccgcggttatttcaggaaagtttcggaggagatagtgttcggcagtttgtacatcatctgcgggatcaggtacggtttgatcaggttgtagaagatcaggtaagacatagaatcgatgtagatgatcggtttgtttttgttgatttttacgtaacagttcagttggaatttgttacgcagacccttaaccaggtattctacttcttcgaaagtgaaagactgggtgttcagtacgatcgatttgttggtagagtttttgttgtaatcccatttaccaccatcatccatgaaccagtatgccagagacatcggggtcaggtagttttcaaccaggttgttcgggatggtttttttgttgttaacgatgaacaggttagccagtttgttgaaagcttggtgtttgaaagtctgggcgccccaggtgattaccaggttacccaggtggttaacacgttcttttttgtgcggcggggacagtacccactgatcgtacagcagacatacgtggtccatgtatgctttgtttttccactcgaactgcatacagtaggttttaccttcatcacgagaacggatgtaagcatcacccaggatcagaccgatacctgcttcgaactgttcgatgttcagttcgatcagctgggatttgtattctttcagcagtttagagttcggacccaggttcattacctggttttttttgatgtttttcatacccgcatagtcaggaacatcgtatgggtacataaatcgtgggtcttctacttttcgtttttttttggtggtgctgcatgctgatcggcacgtaagaggttccaactttcaccataatgaaataagatcactaccgggcgtattttttgagttatcgagattttcaggagctaaggaagctaaaatgagccatattcaacgggaaacgtcttgctcgaggccgcgattaaattccaacatggatgctgatttatatgggtataaatgggctcgcgataatgtcgggcaatcaggtgcgacaatctatcgattgtatgggaagcccgatgcgccagagttgtttctgaaacatggcaaaggtagcgttgccaatgatgttacagatgagatggtcaggctaaactggctgacggaatttatgcctcttccgaccatcaagcattttatccgtactcctgatgatgcatggttactcaccactgcgatcccagggaaaacagcattccaggtattagaagaatatcctgattcaggtgaaaatattgttgatgcgctggcagtgttcctgcgccggttgcattcgattcctgtttgtaattgtccttttaacggcgatcgcgtatttcgtctcgctcaggcgcaatcacgaatgaataacggtttggttggtgcgagtgattttgatgacgagcgtaatggctggcctgttgaacaagtctggaaagaaatgcataagcttttgccattctcaccggattcagtcgtcactcatggtgatttctcacttgataaccttatttttgacgaggggaaattaataggttgtattgatgttggacgagtcggaatcgcagaccgataccaggatcttgccatcctatggaactgcctcggtgagttttctccttcattacagaaacggctttttcaaaaatatggtattgataatcctgatatgaataaattgcagtttcacttgatgctcgatgagtttttctaatgagggcccaaatgtaatcacctggctcaccttcgggtgggcctttctgcggcatgcaagcttggcgtaatcatggtcatagctgtttcctgtgtgaaattgttatccgctcacaattcgcggcctttttacggttcctggccttttgctggccttttgctcacatgttggtctccagcttgcaaattaaagccttcgagcgtcccaaaaccttctcaagcaaggttttcagtataatgttacatgcgtacacgcgtctgtacagaaaaaaaagaaaaatttgaaatataaataacgttcttaatactaacataactataaaaaaataaatagggacctagacttcaggttgtctaactccttccttttcggttagagcggatgtgggggagggcgtgaatgtaagcgtgacataactaattacatgatatcgacaaaggaaaagggggacggatctccgaggcctcggacccgtcgggccgcgtcggacgtgtcagtcctgctcctcggccacgaagtgcacgcagttgccggccgggtcgcgcagggcgaactcccgcccccacggctgctcgccgatctcggtcatggccggcccggaggcgtcccggaagttcgtggacacgacctccgaccactcggcgtacagctcgtccaggccgcgcacccacacccaggccagggtgttgtccggcaccacctggtcctggaccgcgctgatgaacagggtcacgtcgtcccggaccacaccggcgaagtcgtcctccacgaagtcccgggagaacccgagccggtcggtccagaactcgaccgctccggcgacgtcgcgcgcggtgagcaccggaacggcactggtcaacttggccatggtttagttcctcaccttgtcgtattatactatgccgatatactatgccgatgattaattgtcaacaccgcccttagattagattgctatgctttctttctaatgagcaagaagtaaaaaaagttgtaatagaacaagaaaaatgaaactgaaacttgagaaattgaagaccgtttattaacttaaatatcaatgggaggtcatcgaaagagaaaaaaatcaaaaaaaaaattttcaagaaaaagaaacgtgataaaaatttttattgcctttttcgacgaagaaaaagaaacgaggcggtctcttttttcttttccaaacctttagtacgggtaattaacgacaccctagaggaagaaagaggggaaatttagtatgctgtgcttgggtgttttgaagtggtacggcgatgcgcggagtccgagaaaatctggaagagtaaaaaaggagtagaaacattttgaagctatggtgtgtgggggatccgcacaaacgaaggtctcacttaatcttctgtactctgaagaggagtgggaaataccaagaaaaacatcaaactcgaatgattttcccaaacccctaccacaagatattcatcagctgcaaactaagattggactattgctatccaacgtaaaaggactaactagaatagacttcttgtggcaatctttacttttgttttcacacagactcagttgtgagccaacctctatcctgtaaggtcaaagcaagactaggcttcagacttgcaacaactaatagttgtggaatagtacagaggattgaagttagtttttttgggaggcctgacacaagtcagttagttttttctgaacaggctaagattcaaactagtctggtttttcacacactactgtctacttaagtcaaatctgagaaacactgtctgaagttagacttgaataactgagcctactttattctaaactagacttgtctgttgcagtgagttaggcaagtttcgaataagcagcaagaacagttgtactcagtctaggtcaaaacagctacactgtttctaaaccaaaggcagaggattgagattctaactgacttaggtttaaaactgccagacactcaatctttctactaaaagtacaagtccacgaagtagacgcttactagtttgttaacgtaaactgttgaggcacaaaccgtcagtaatacagaggtgaagtaaactactagttgtctttttaaacgtttaaacctagacaactagcacgaacgcagctacttgttgaactcttgtgtaacacagacaaatctattgaatctttctgcttttcagacttttcgtactcacagttctaacttggtaactagagtttctagttggttaactgaggtttcgacttgtaacagtctagtggctagttctgaatacgattgcagtagactgtgggttgacttaaagacggtttaacacagctgcgagataggctgatcaggagcaagctcgtacgagaagaaacaaaatgacaactgcattaatgaatcggccaacgcgcggggagaggcggtttgcgtattgggcgctcttccgcttcctcgctcactgactcgctgcgctcggtcgttcggctgcggcgagcggtatcagctcactcaaaggcggtaatacggttatccacagaatcaggggataacgcaggaaagaacatgtgagcaaaaggccagcaaaaggccaggaaccgtaaaaaggccgcgttgctggcgtttttccataggctccgcccccctgacgagcatcacaaaaatcgacgctcaagtcagaggtggcgaaacccgacaggactataaagataccaggcgtttccccctggaagctccctcgtgcgctctcctgttccgaccctgccgcttaccggatacctgtccgcctttctcccttcgggaagcgtggcgctttctcatagctcacgctgtaggtatctcagttcggtgtaggtcgttcgctccaagctgggctgtgtgcacgaaccccccgttcagcccgaccgctgcgccttatccggtaactatcgtcttgagtccaacccggtaagacacgacttatcgccactggcagcagccactggtaacaggattagcagagcgaggtatgtaggcggtgctacagagttcttgaagtggtggcctaactacggctacactagaagaacagtatttggtatctgcgctctgctgaagccagttaccttcggaaaaagagttggtagctcttgatccggcaaacaaaccaccgctggtagcggtggtttttttgtttgcaagcagcagattacgcgcagaaaaggatctcaagaagatcctttgatcttttctacggggtctgacgctcagtggaacgaaaactcacgttaagggattttggtcatgagattatcaaaaaggatcttcacctagatccttttaaattaaaaatgaagttttaaatcaatctaaagtatatatgagtaaacttggtctgacagttaccaatgcttaatcagtgaggcacctatctcagcgatctgtctatttcgttcatccatagttgcctgactccccgtcgtgtagataactacgatacgggagggcttaccatctggccccagtgctgcaatgataccgcgagacccacgctcaccggctccagatttatcagcaataaaccagccagccggaagggccgagcgcagaagtggtcctgcaactttatccgcctccatccagtctattaattgttgccgggaagctagagtaagtagttcgccagttaatagtttgcgcaacgttgttgccattgctacaggcatcgtggtgtcacgctcgtcgtttggtatggcttcattcagctccggttcccaacgatcaaggcgagttacatgatcccccatgttgtgcaaaaaagcggttagctccttcggtcctccgatcgtggatgctggatgctggatgctggatgctggatgctggatgctggatgctggatgctggatgctggatgctggatgctggatgctggatgctggatgctggatgctggatgctggatgctggatgctggatgctggatgctggatgctggatgctggatgctggatgctggatgctggatgctggatgctggatgctggatgct
